# Identification of key genes and biological pathways in chronic obstructive pulmonary disease using bioinformatics and next generation sequencing data analysis

**DOI:** 10.1101/2024.02.10.579759

**Authors:** Basavaraj Vastrad, Chanabasayya Vastrad

## Abstract

Identification of accurate biomarkers is still particularly urgent for improving the poor survival of chronic obstructive pulmonary disease (COPD) patients. In this investigation, we aimed to identity the potential biomarkers in COPD via bioinformatics and next generation sequencing (NGS) data analysis. In this investigation, the differentially expressed genes (DEGs) in COPD were identified using NGS dataset (GSE239897) from Gene Expression Omnibus (GEO) database. Subsequently, gene ontology (GO) and pathway enrichment analysis was conducted to evaluate the underlying molecular mechanisms involved in progression of COPD. Protein-protein interaction (PPI), modules, miRNA-hub gene regulatory network and TF-hub gene regulatory network analysis were performed to determine the hub genes, miRNAs and TFs. The receiver operating characteristic (ROC) analysis was performed to determine the diagnostic value of hub genes. A total of 956 overlapping DEGs (478 up regulated and 478 down regulated genes) were identified in the NGS dataset. DEGs were mainly associated with GO functional terms and pathways in cellular response to stimulus. response to stimulus, immune system and neutrophil degranulation. There were 10 hub genes (MYC, LMNA, VCAM1, MAPK6, DDX3X, SHMT2, PHGDH, S100A9, FKBP5 and RPS6KA2) identified by PPI, modules, miRNA-hub gene regulatory network and TF-hub gene regulatory network analysis. In conclusion, the DEGs, relative GO terms, pathways and hub genes identified in the present investigation might aid in understanding of the molecular mechanisms underlying COPD progression and provide potential molecular targets and biomarkers for COPD.

## Introduction

Chronic obstructive pulmonary disease (COPD) represents the third leading cause of death in 2030 according to WHO prediction [1]. The clinical incidence of COPD is high, and its main features include expiratory airflow limitation that is not fully reversible, deregulated chronic airway inflammation, and emphysematous destruction of the lungs [2]. It might affect many other organs and cause other diseases include pneumonia [3], respiratory viral infections [4], pneumothorax [5], heart problems [6], osteoporosis [7], depression and anxiety [8], lung cancer [9], pulmonary hypertension [10], secondary polycythemia [11], idiopathic pulmonary fibrosis [12], obesity [13] and diabetes mellitus [14]. Airway inflammation is triggered by both genetic susceptibility [15] and environmental factors [16]. However, the etiology and symptoms of COPD are complex in clinical practice, making its diagnosis challenging.

Correct early diagnosis assessment of COPD is very difficult, even though much disease related genes and cellular pathways related to COPD have appeared [17]. The common treatments of COPD are bronchodilator and antiinflammatory treatments, such as phosphodiesterase (PDE)-4, p38 mitogen-activated protein kinase (MAPK), and nuclear factor (NF)-κB inhibitors [18], but there are no valid treatment tactics available to treat COPD. Therefore, it is vitally important to explore potential diagnostic and prognostic biomarkers, and therapeutic targets of COPD.

In recent years, next generation sequencing (NGS) technology has been applied to the research on various bioinformatics, which can screen and identify genes and signalizing pathways of various diseases [19–20]. NGS is an emerging molecular biology technology based on a high-throughput platform, which is widely used in COPD [21]. Aberrant expression of genes plays an essential role in the initiation and progression of COPD, so mastering the modification in the characteristics of essential genes promotes to comprehensively understand COPD progression and screen related molecular markers [22]. Recently, investigation shows expression of genes include SERPINE2 [23], TNFα, IL1β, and IL1RN [24], TGFB1 [25], ADRB2 [26] and PTX3 [27] in COPD. Indeed, some researchers found signaling pathways include HIFL1 signaling pathway [28], Wnt signal pathway [29], PI3K signaling in pathway [30], TLR4/NF-kB signaling pathway [31] and FoxO1/MuRF1/Atrogin-1 signaling pathway [32] were responsible for advancement of COPD. However, the use of bioinformatics analysis methods to identify the relevant genes and pathways of COPD has not yet been confirmed.

Our main purpose is to explore the connection between COPD and its associated complications. First, we download the GSE239897 [33] NGS dataset file in the NCBI Gene Expression Omnibus (GEO) [https://www.ncbi.nlm.nih.gov/geo/] [34] database for analysis, then use limma package in R software to draw the differentially expressed genes (DEGs) distribution map of the COPD and normal control samples in the dataset. Then, based on gene ontology (GO) terms and the REACTOME pathways related to COPD are analyzed. Immediately afterward, the protein-protein interaction (PPI) network, modules, miRNA-hub gene regulatory network and TF-hub gene regulatory network are drawn and the hub genes, miRNAs and TFs are identified. Hub genes were further validated by receiver operating characteristic (ROC) curve analysis. The final results will help us obtain novel treatment targets for COPD.

## Materials and Methods

### Next generation sequencing data source

The GEO database is an open source platform for the storage of NGS data. NGS dataset [GSE239897 (GPL17303 Ion Torrent Proton (Homo sapiens)] [33] was downloaded. The GSE239897 dataset includes 43 COPD samples and 39 normal control samples.

### Identification of DEGs

The DEGs in the samples were identified by the limma package of R software [35]. The Benjamini & Hochberg False Discovery Rate correction method [36] was used to correct the P values. Fold change > 0.45 for up regulated genes, Fold change < - 0.407 for down regulated genes and adj. PLvalue <0.05 were considered to be statistically significant. ggplot2 packages of R software was applied to generate volcano plot. Hierarchical clustering analysis was performed and the gplot packages in R software was used for visualization of heatmaps.

### GO and pathway enrichment analyses of DEGs

To better investigate the biological functions of DEGs, we performed functional enrichment analysis of COPD DEGs using the g:Profiler (http://biit.cs.ut.ee/gprofiler/) [37], which includes GO (http://www.geneontology.org) [38] terms and REACTOME (https://reactome.org/) [39] pathway enrichment analysis, with P < 0.05 being statistically significant. GO analysis includes biological processes (BP), cellular components (CC), and molecular functions (MF).

### Construction of the PPI network and module analysis

We constructed PPI networks using pickle interactome (http://pickle.gr/) [40] database and Cytoscape visualization software version 3.10.1 (http://www.cytoscape.org/) [41]. The Network Analyzer plugin was used to obtain hub genes according to the node degree [42], betweenness [43], stress [44] and closeness [45] methods. The highest scoring modules were screened using the PEWCC [46] plugin of Cytoscape software, and the genes in the modules were defined as hub genes.

### Construction of the miRNA-hub gene regulatory network

miRNAs can play a role in maintaining physiological stability by regulating the expression of hub genes. miRNA-hub gene regulatory network was constructed using the online tool of miRNet database (https://www.mirnet.ca/) [47]. We used TarBase, miRTarBase, miRecords, miRanda (S mansoni only), miR2Disease, HMDD, PhenomiR, SM2miR, PharmacomiR, EpimiR, starBase, TransmiR, ADmiRE and TAM 2 databases to find miRNAs regulating hub genes, taking the intersection of the results of these fourteen databases. We visualized miRNA-hub gene regulatory network with Cytoscape software [41].

### Construction of the TF-hub gene regulatory network

TFs can play a role in maintaining physiological stability by regulating the expression of hub genes. TF-hub gene regulatory network was constructed using the online tool of NetworkAnalyst database (https://www.networkanalyst.ca/) [48]. We used Jasper database to find TFs regulating hub genes, taking the intersection of the results of this database. We visualized TF-hub gene regulatory network with Cytoscape software [41].

### Receiver operating characteristic curve (ROC) analysis

ROC analysis was performed to predict the diagnostic effectiveness of COPD. We plotted ROC curves for each hub gene using the pROC package in R software [49]. The area under the ROC curve (AUC) value was utilized to determine the diagnostic effectiveness in discriminating COPD from normal control samples.

## Results

### Identification of DEGs

NGS dataset (GSE239897) including 43 COPD and 39 normal control samples. A total of 956 DEGs were obtained and included 478 up regulated and 478 down regulated genes (Table 1). The DEGs were visualized by the volcano plot (Fig.1) and heatmap (Fig.2).

**Table 1.**
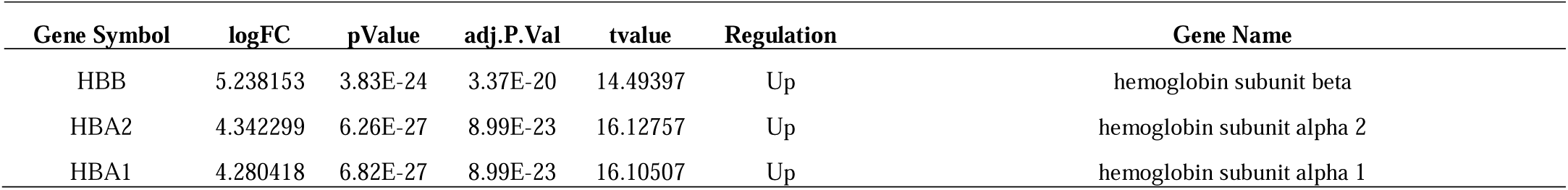
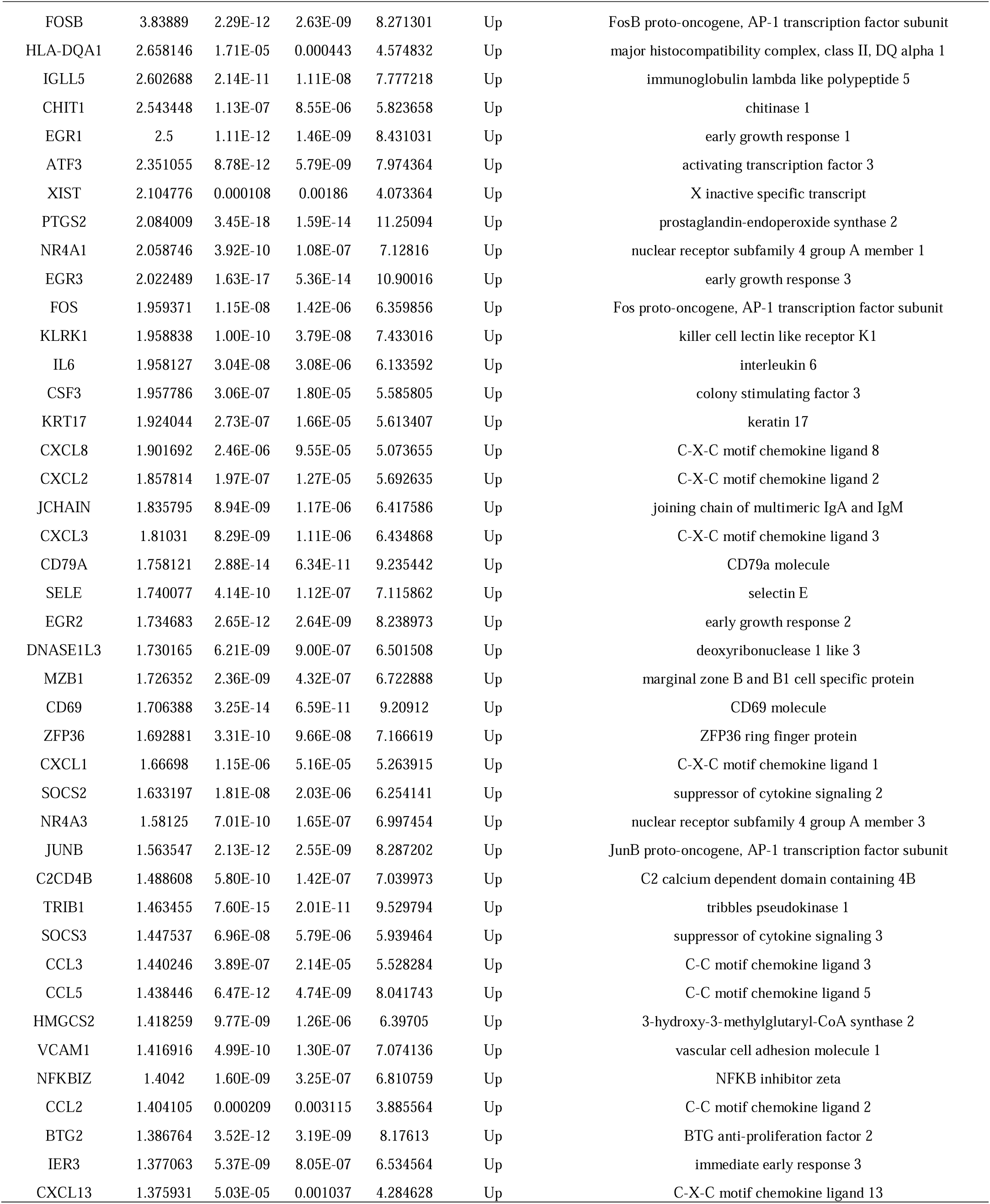
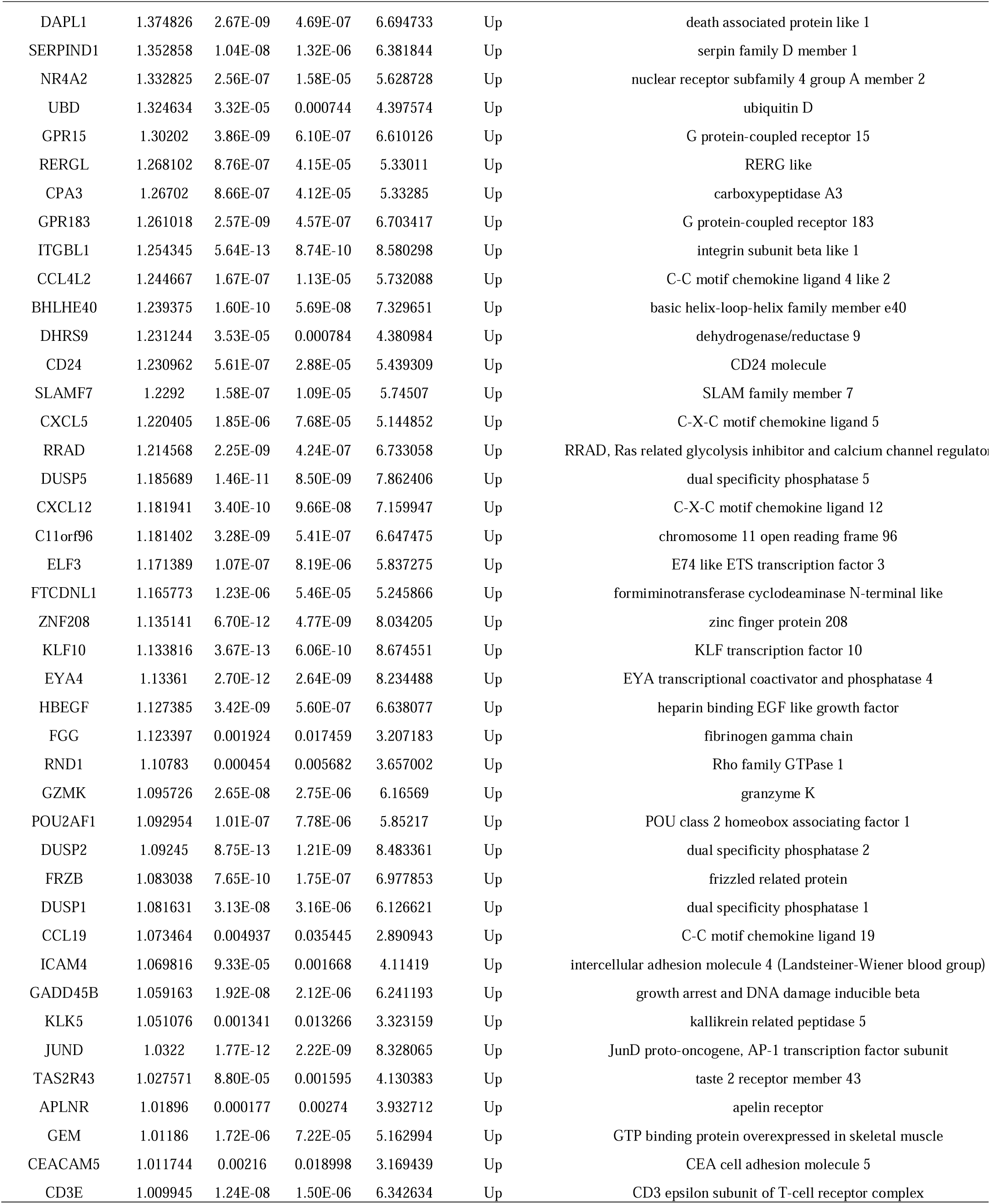
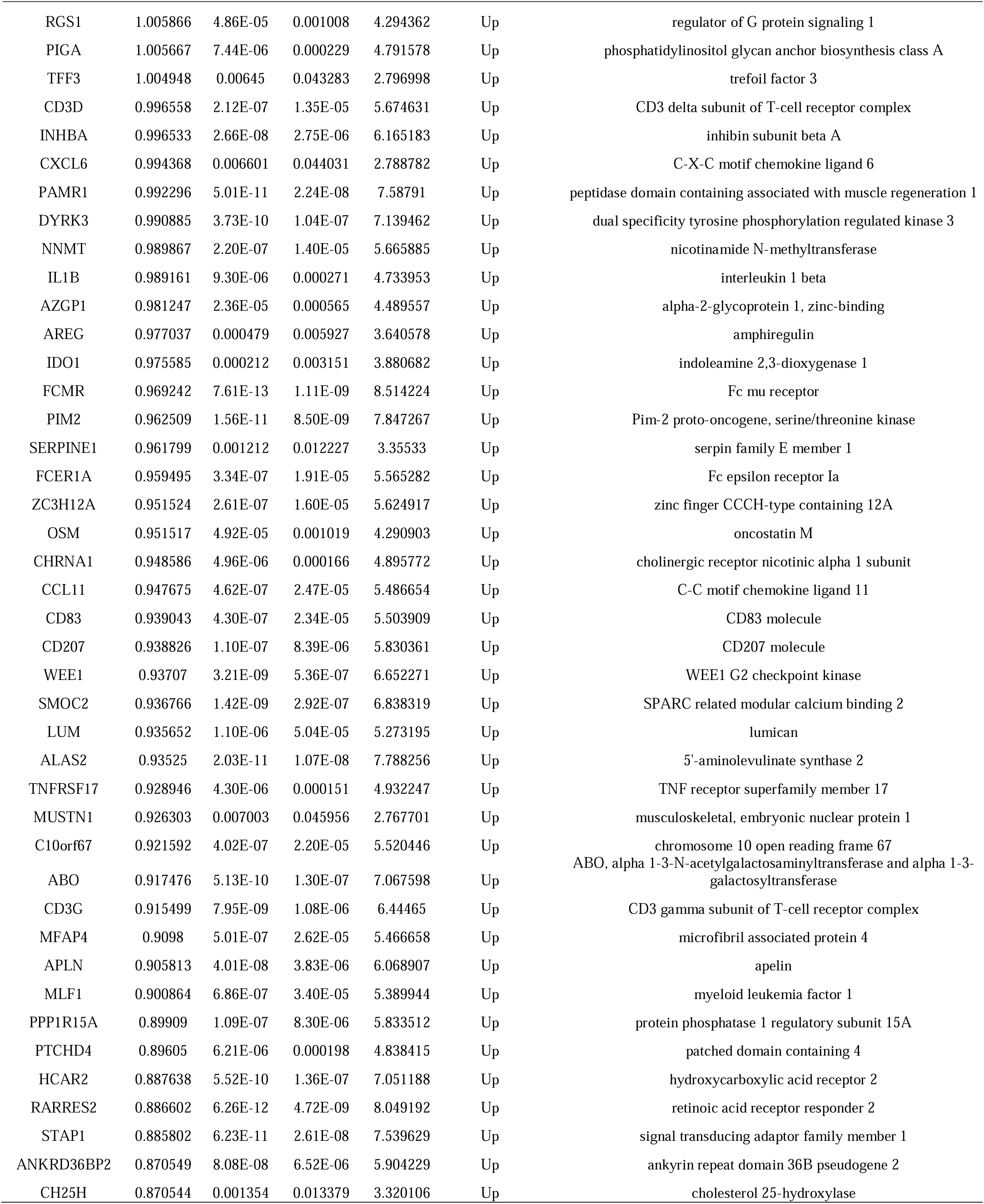
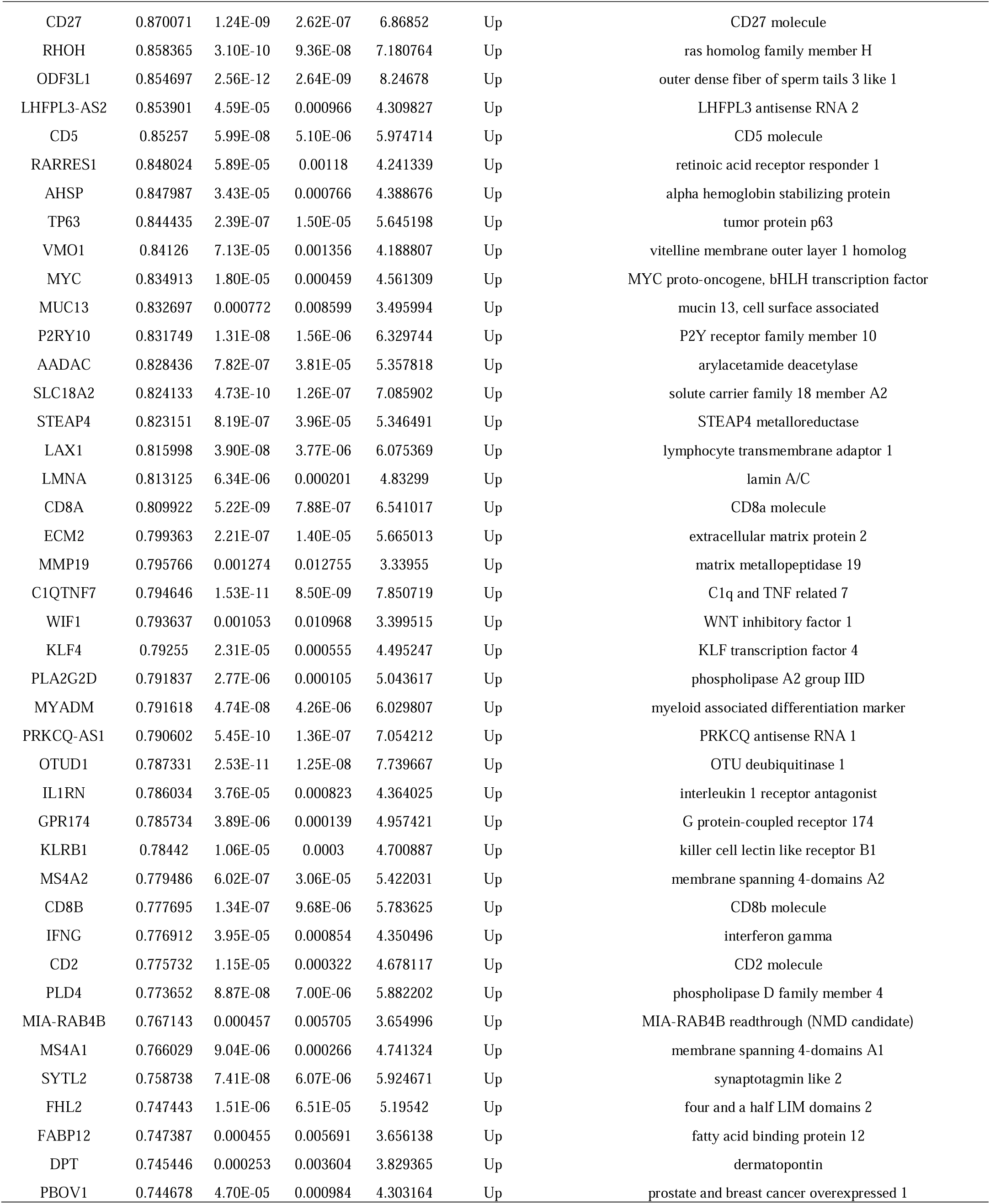
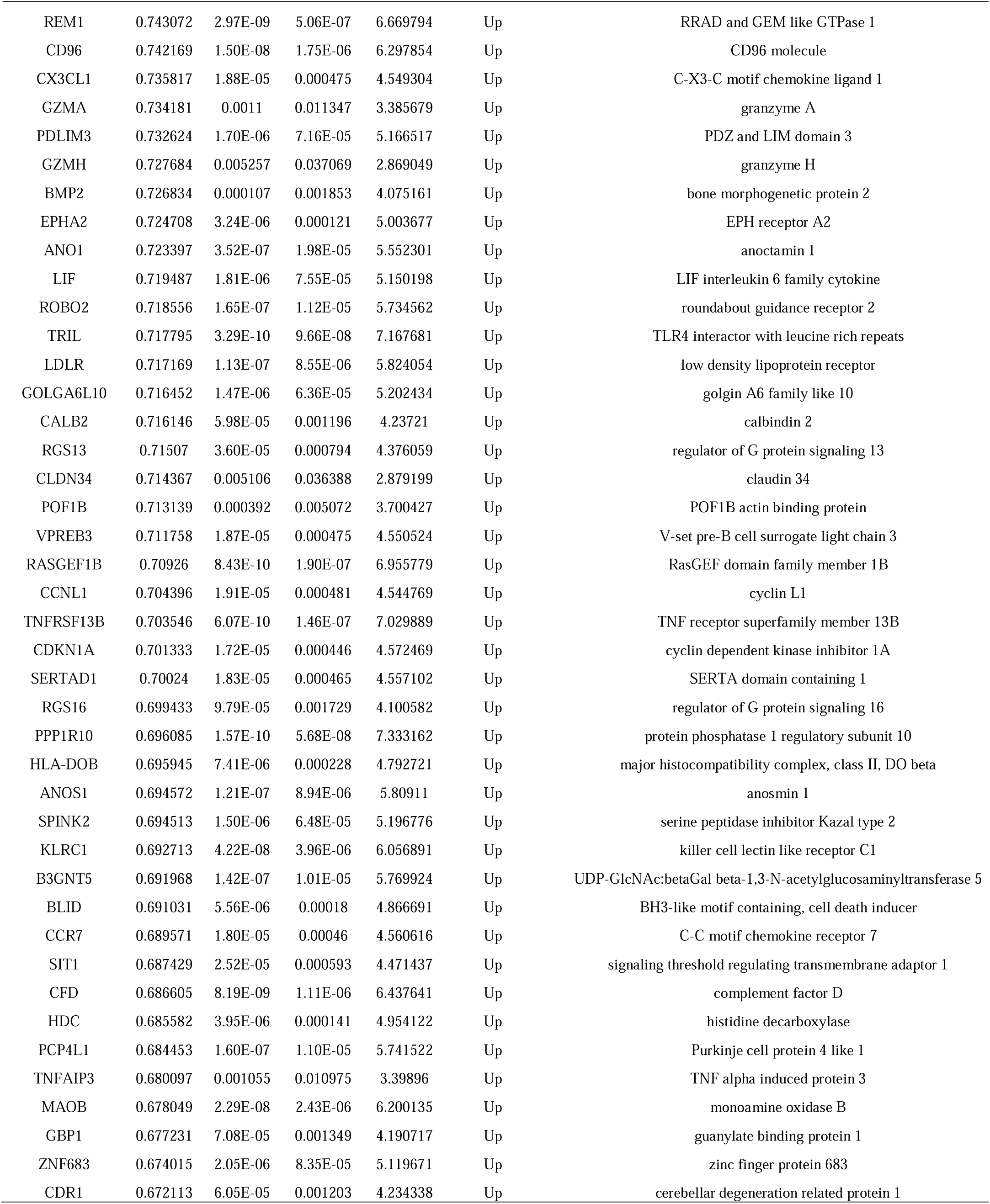
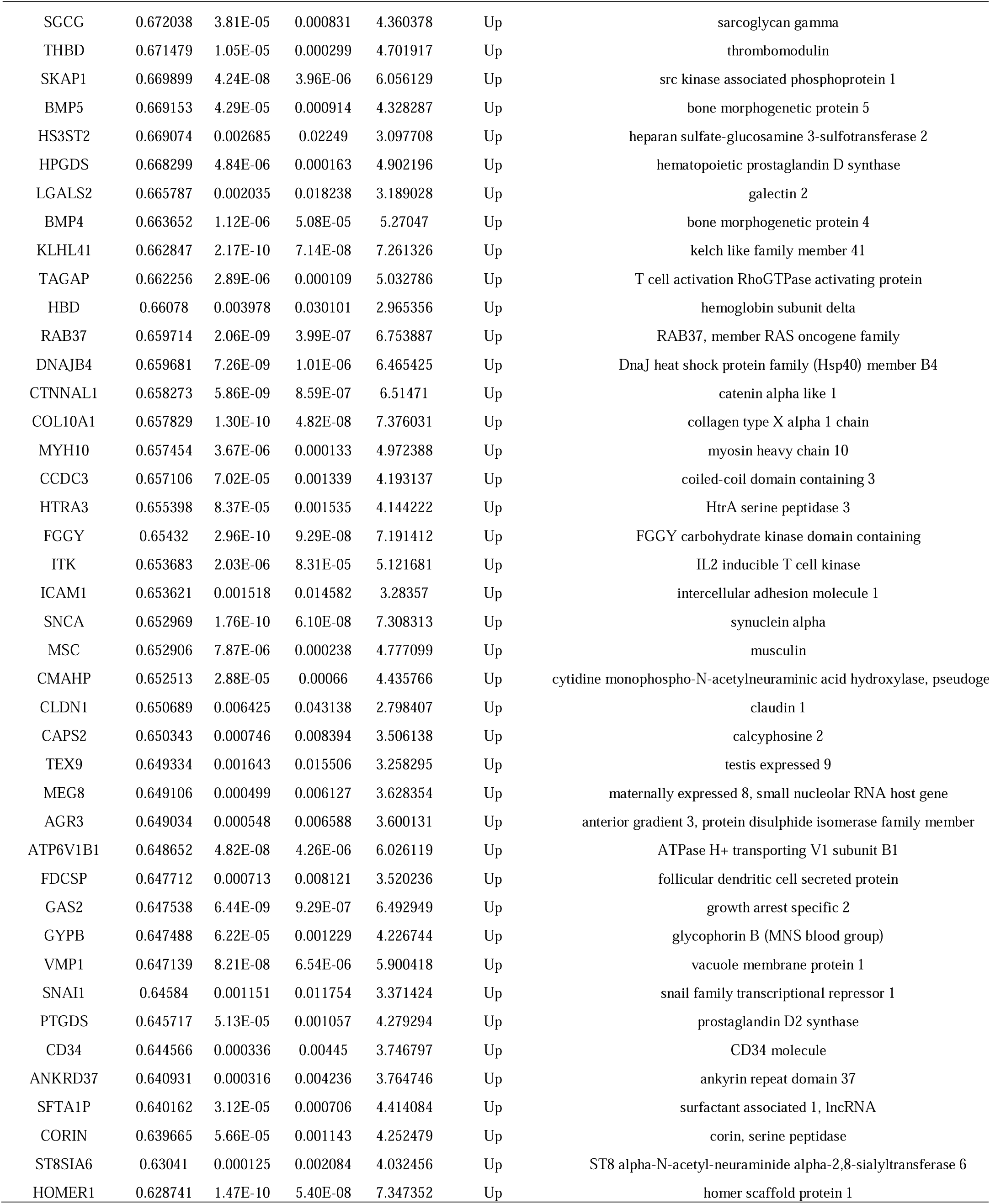
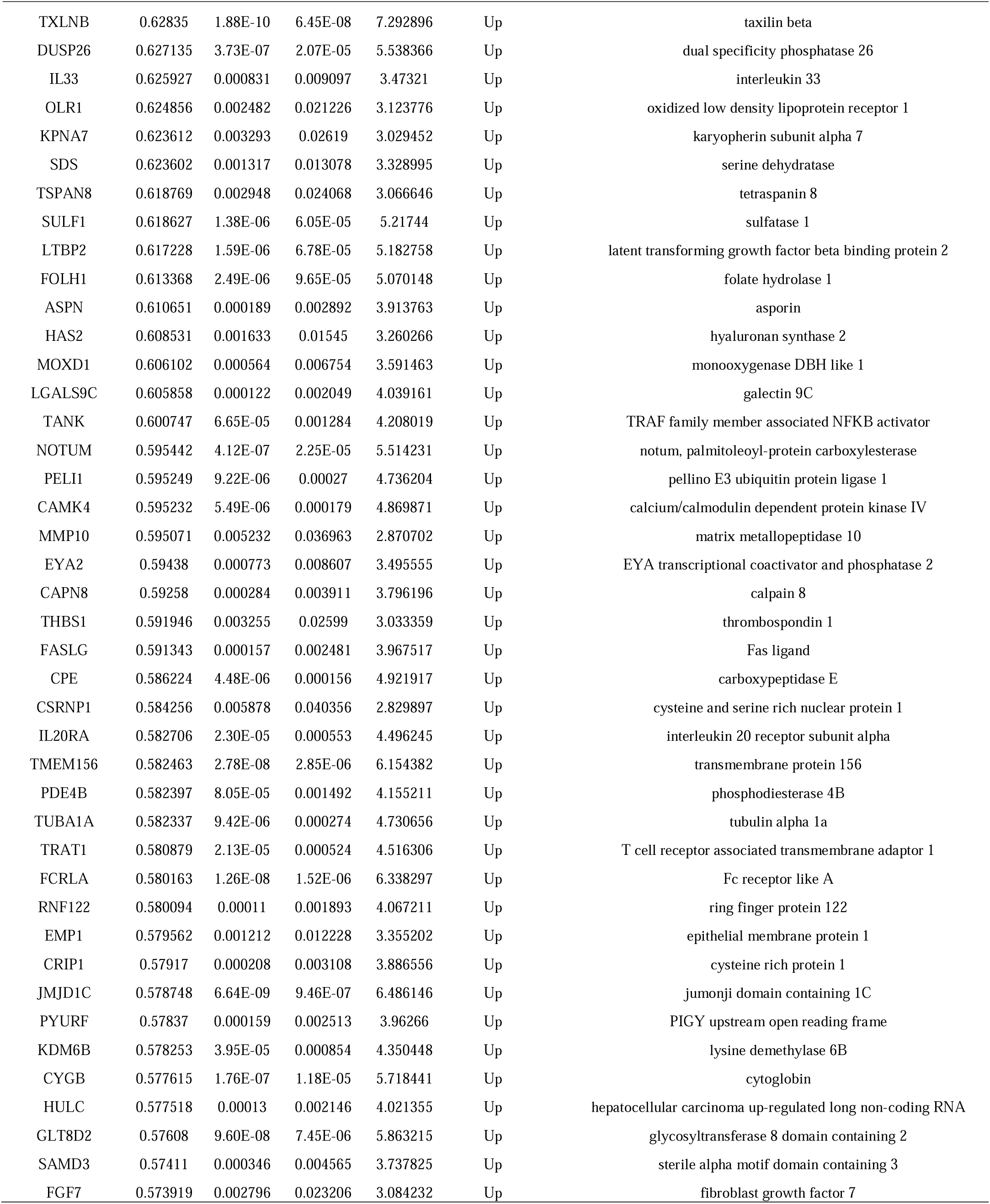
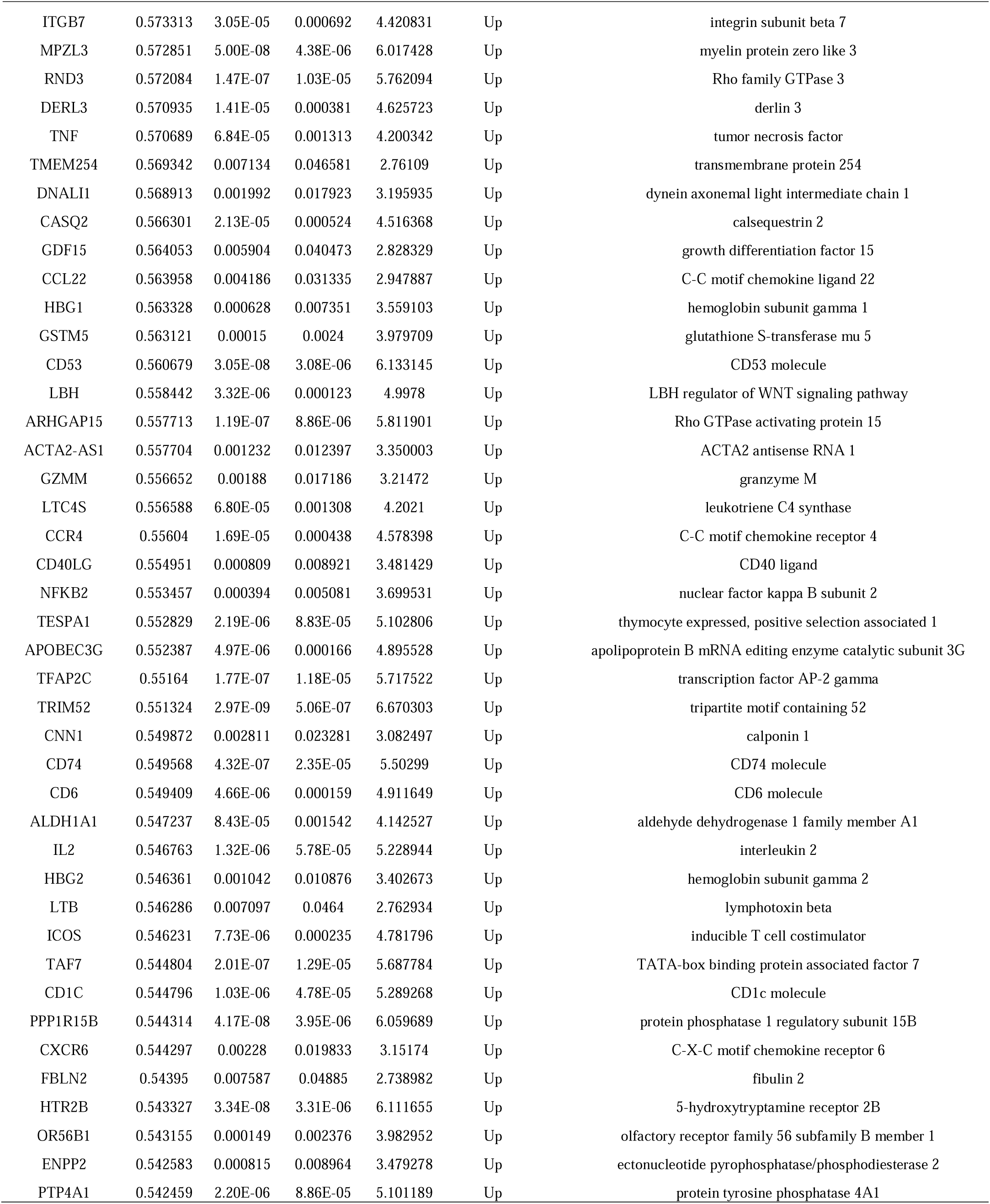
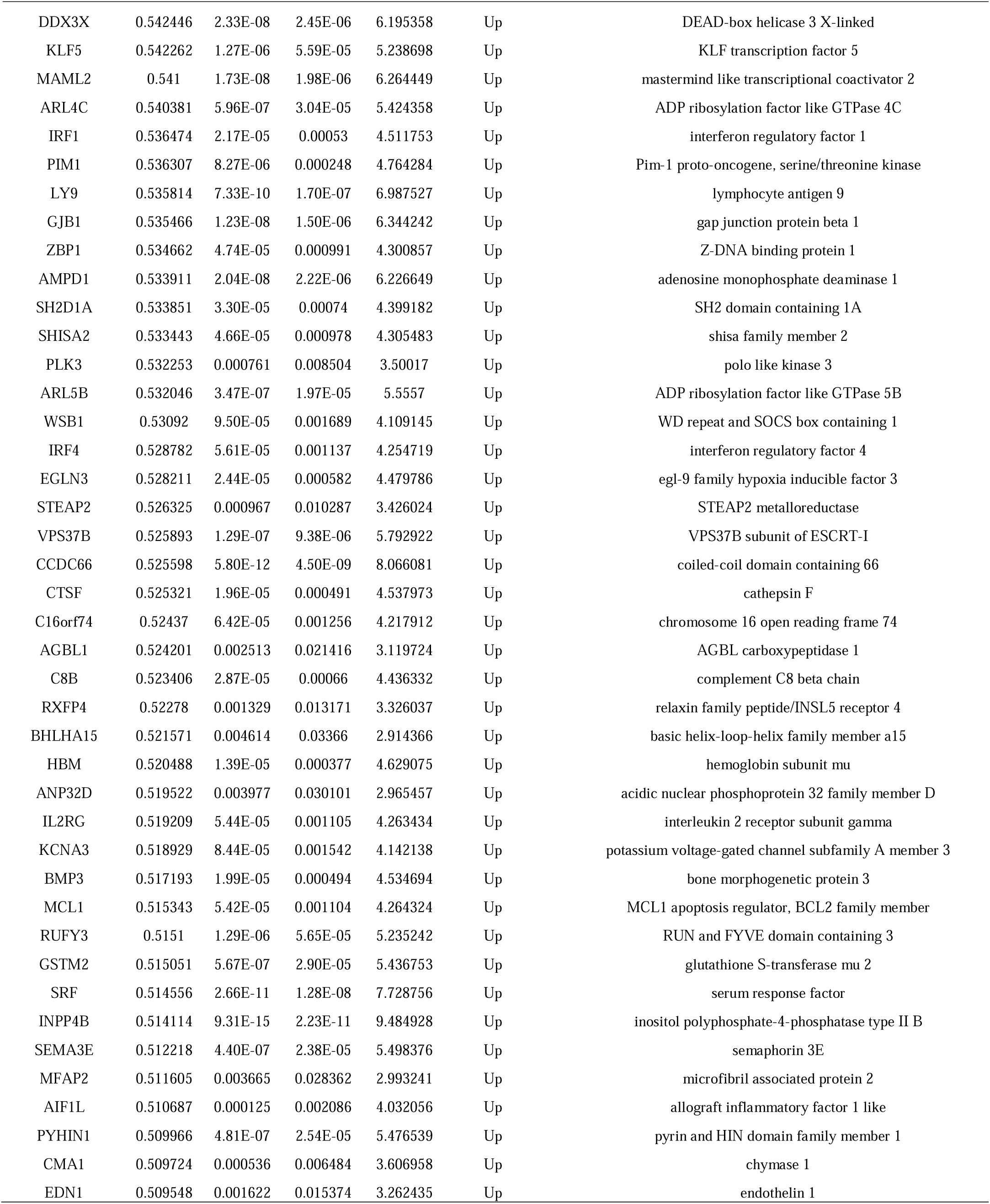
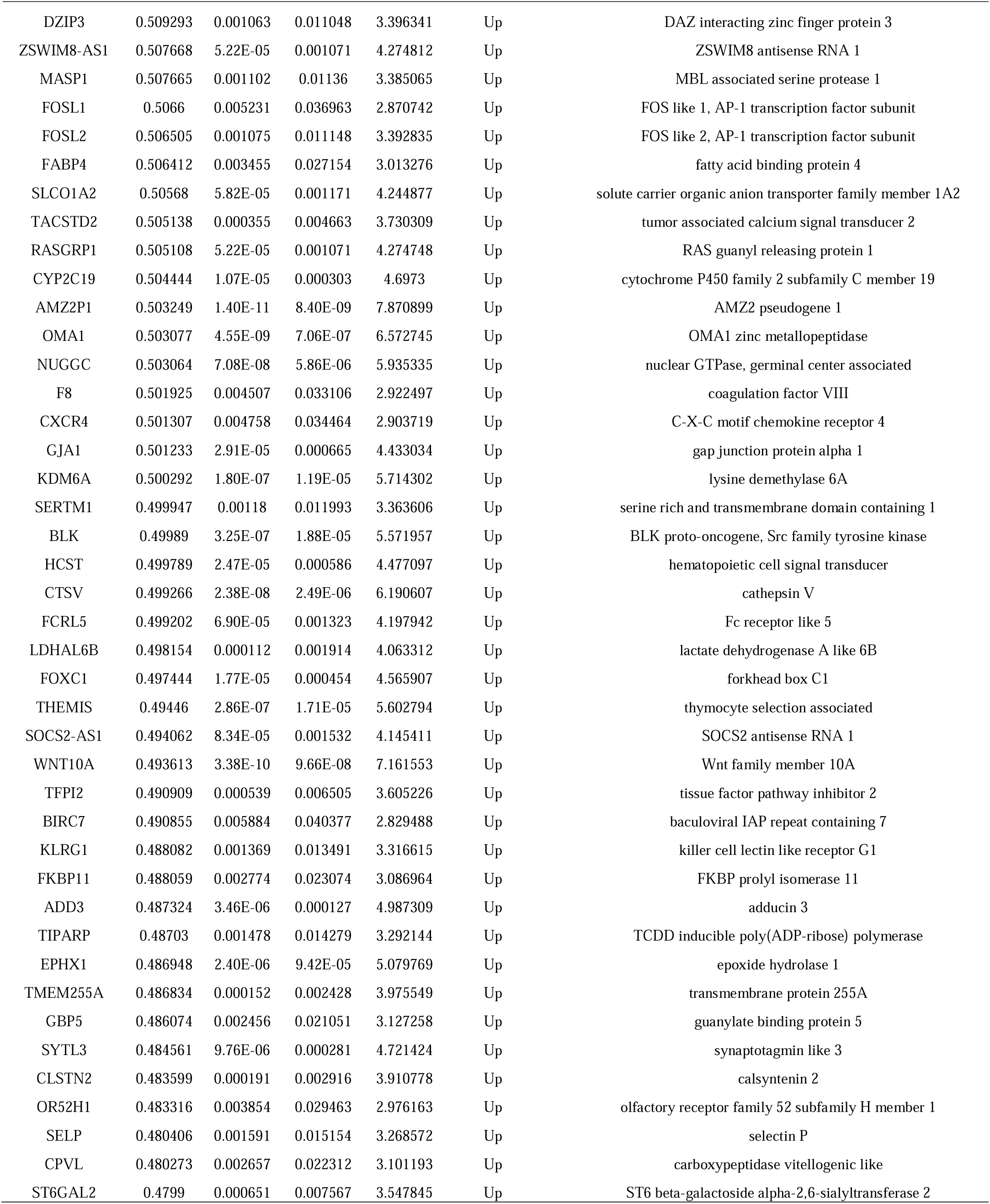
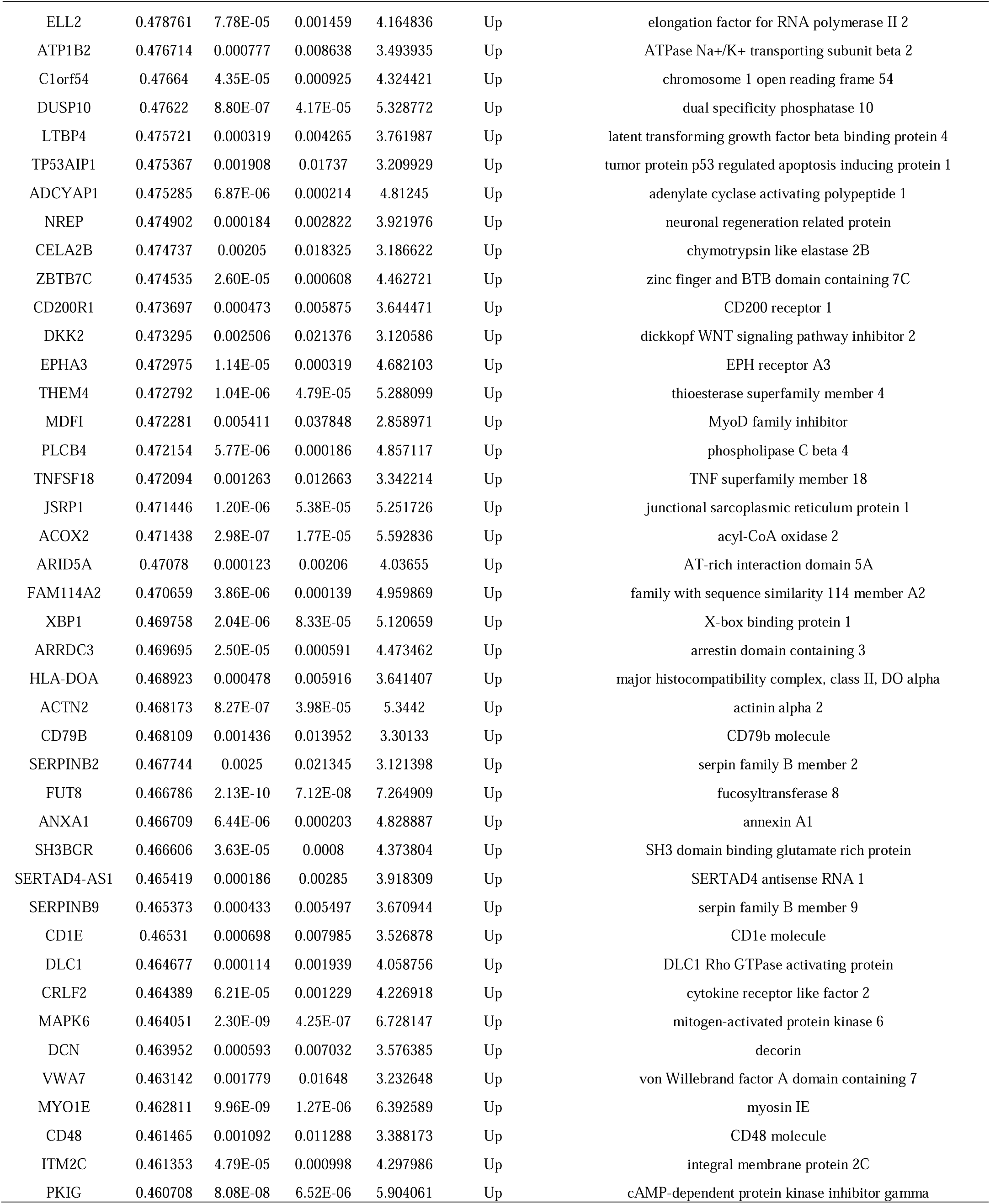
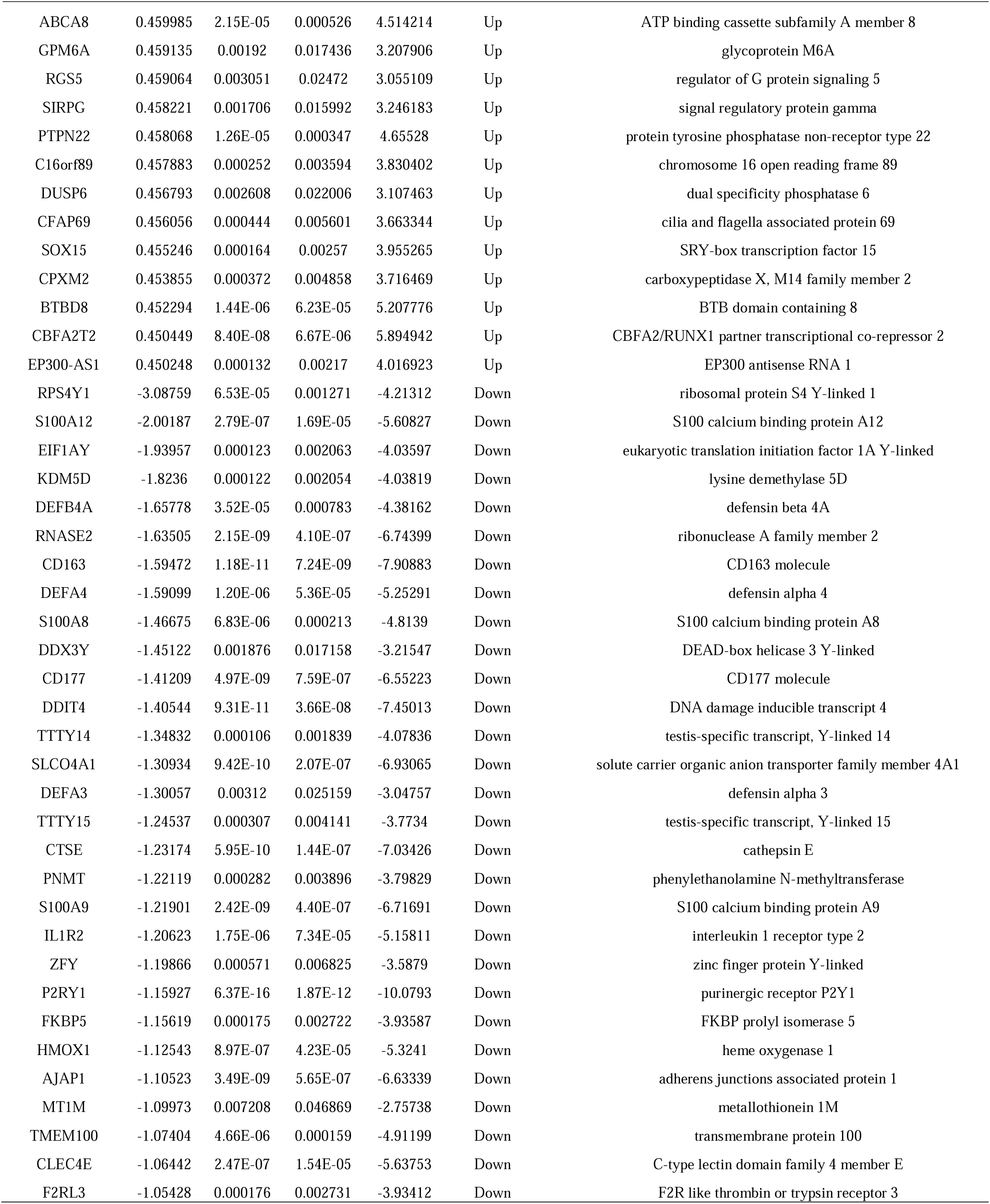
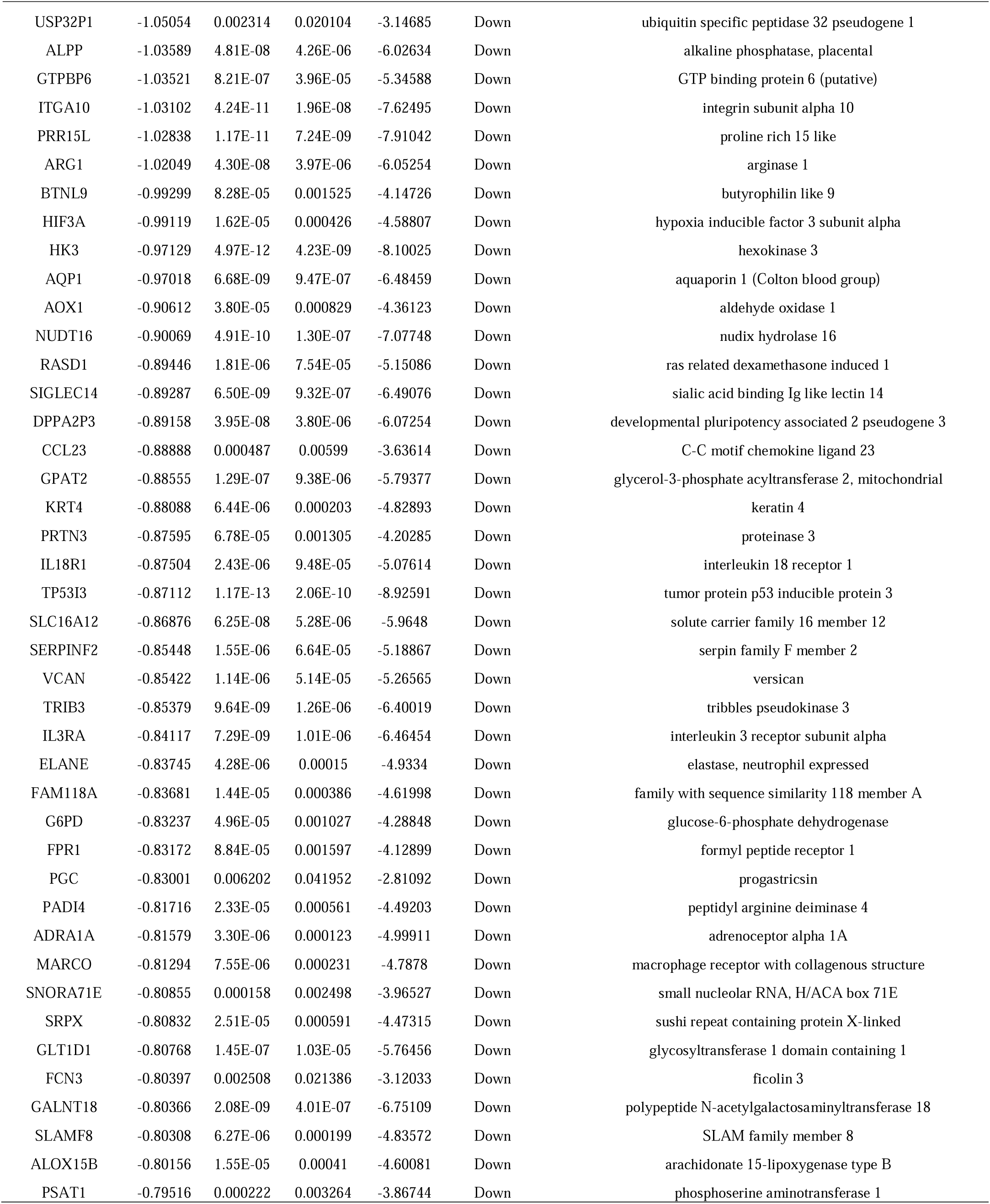
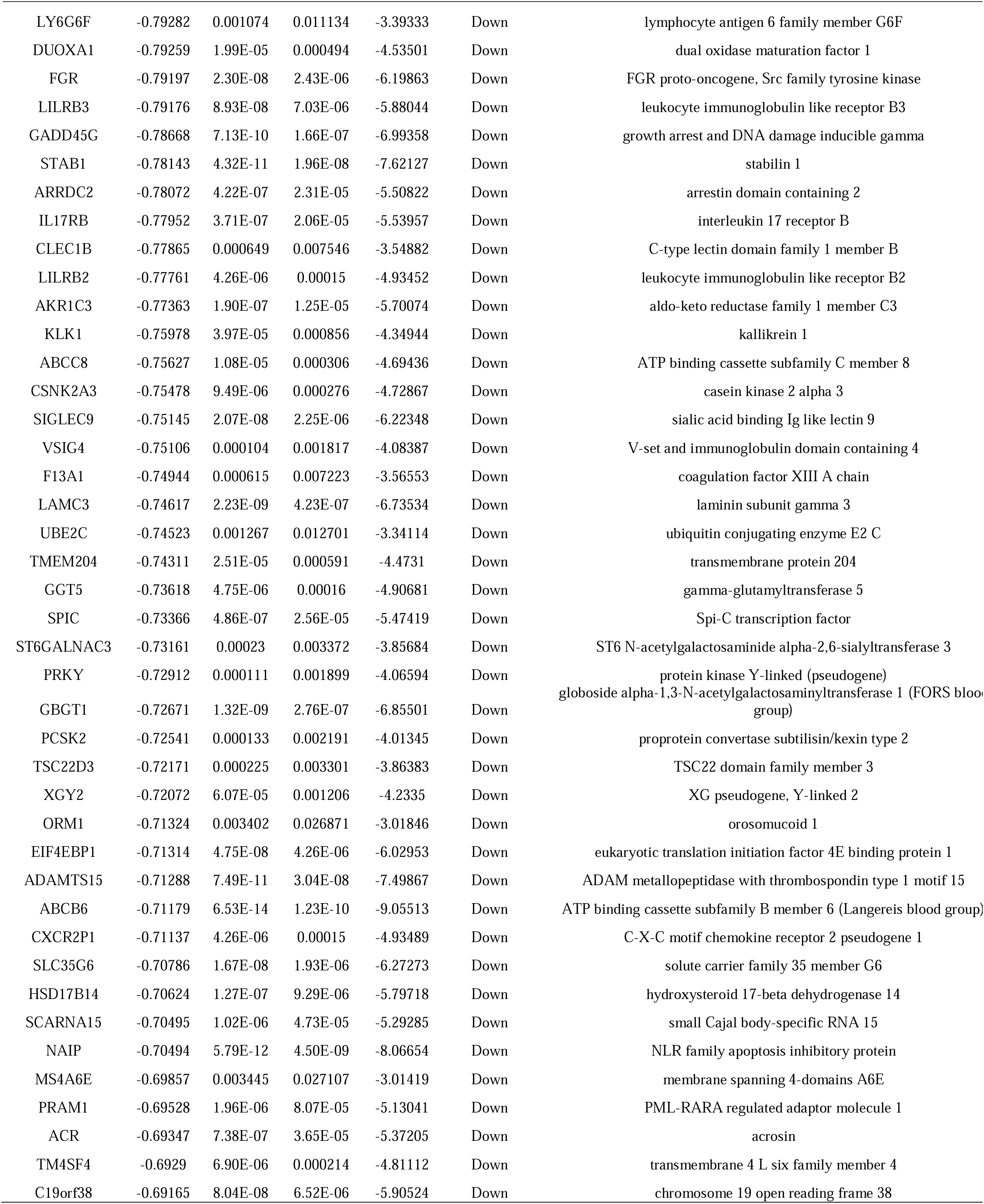
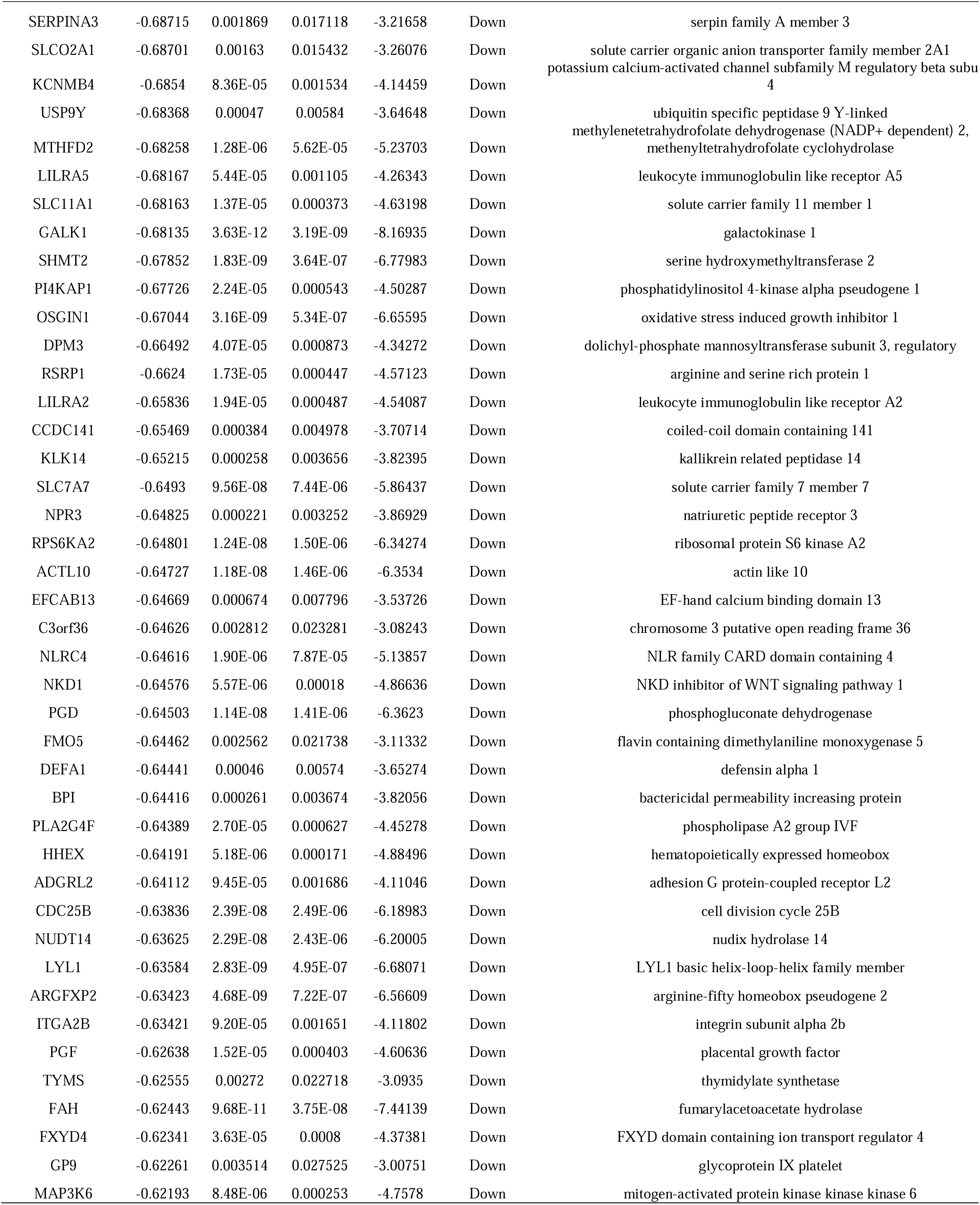
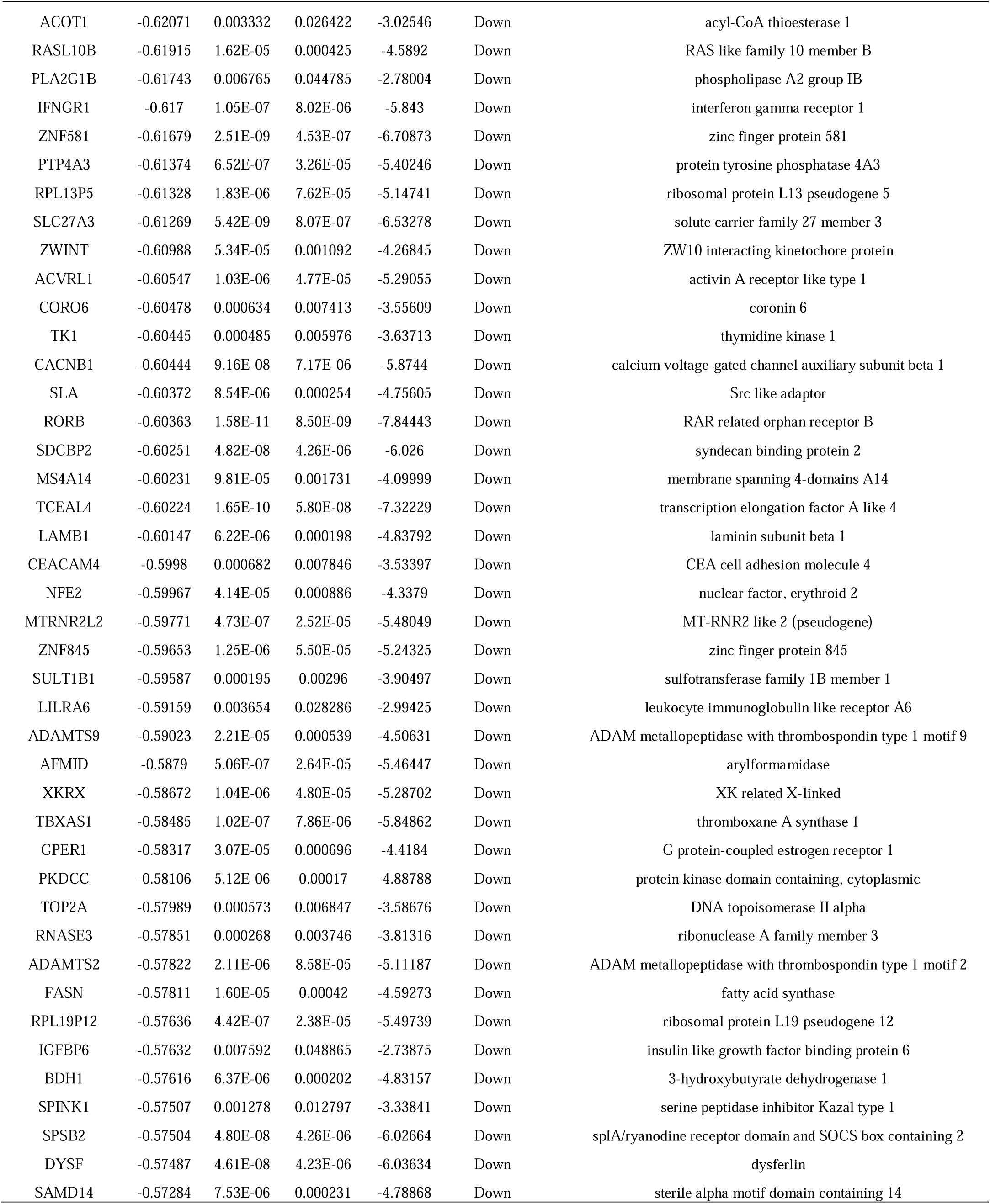
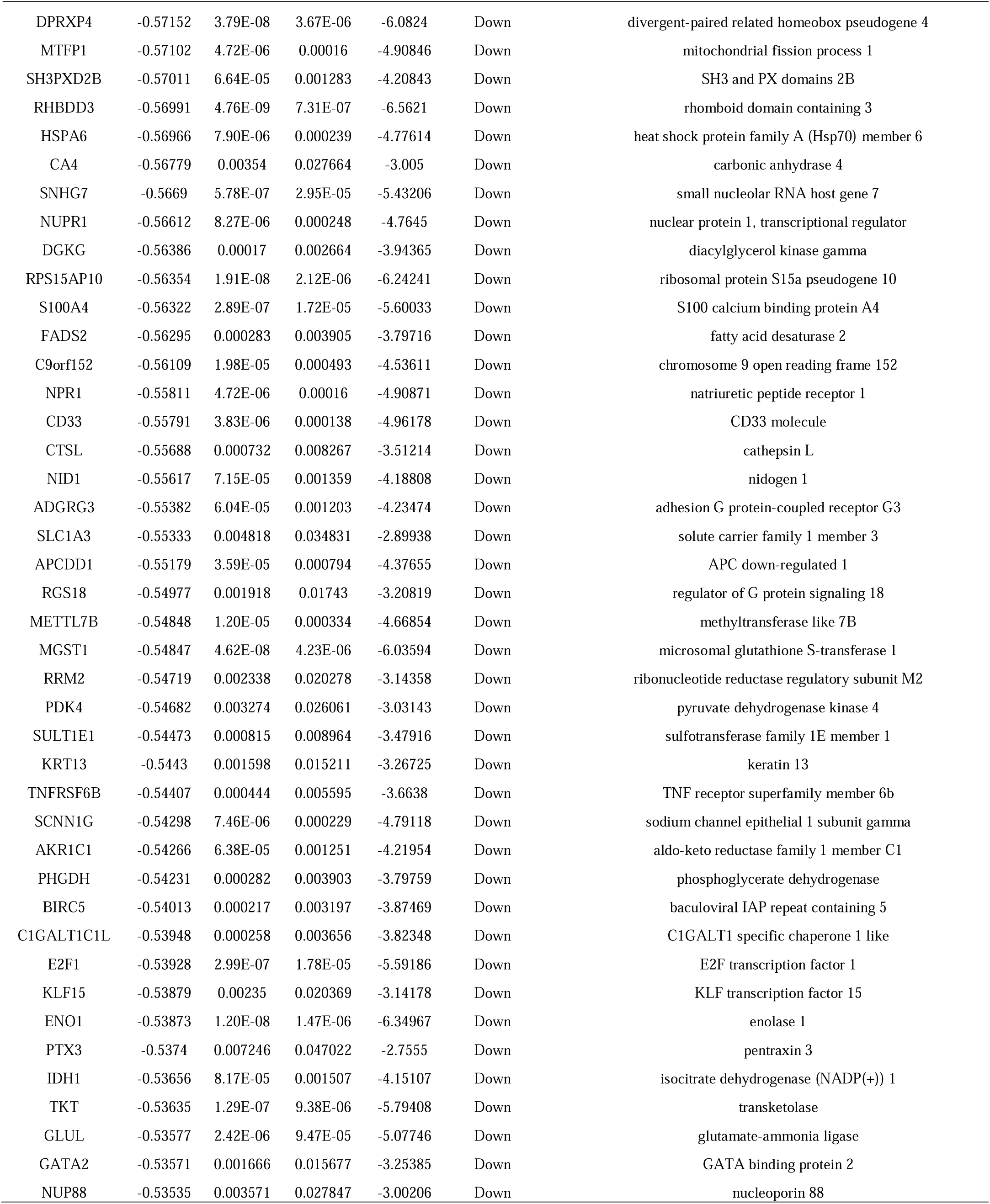
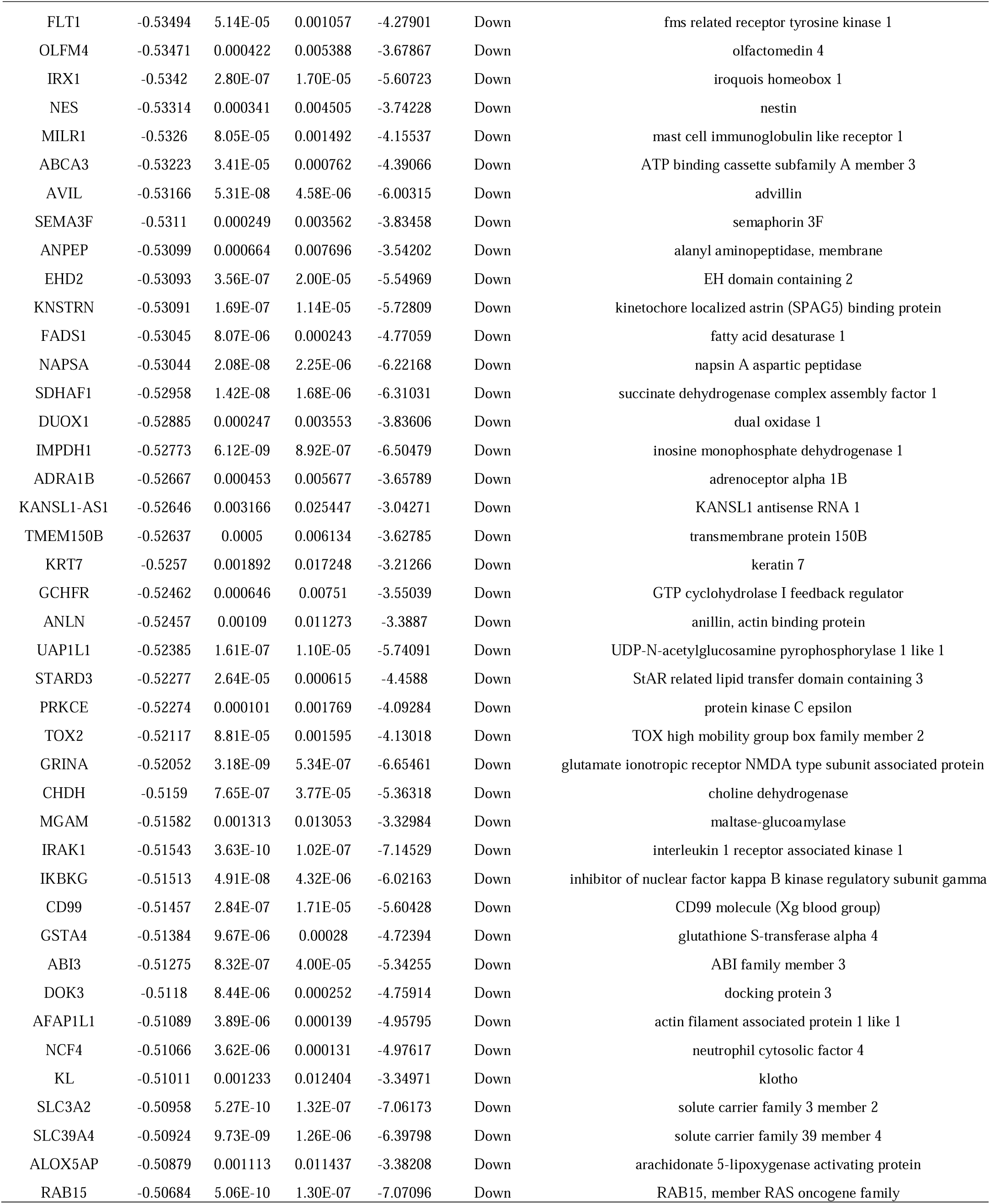
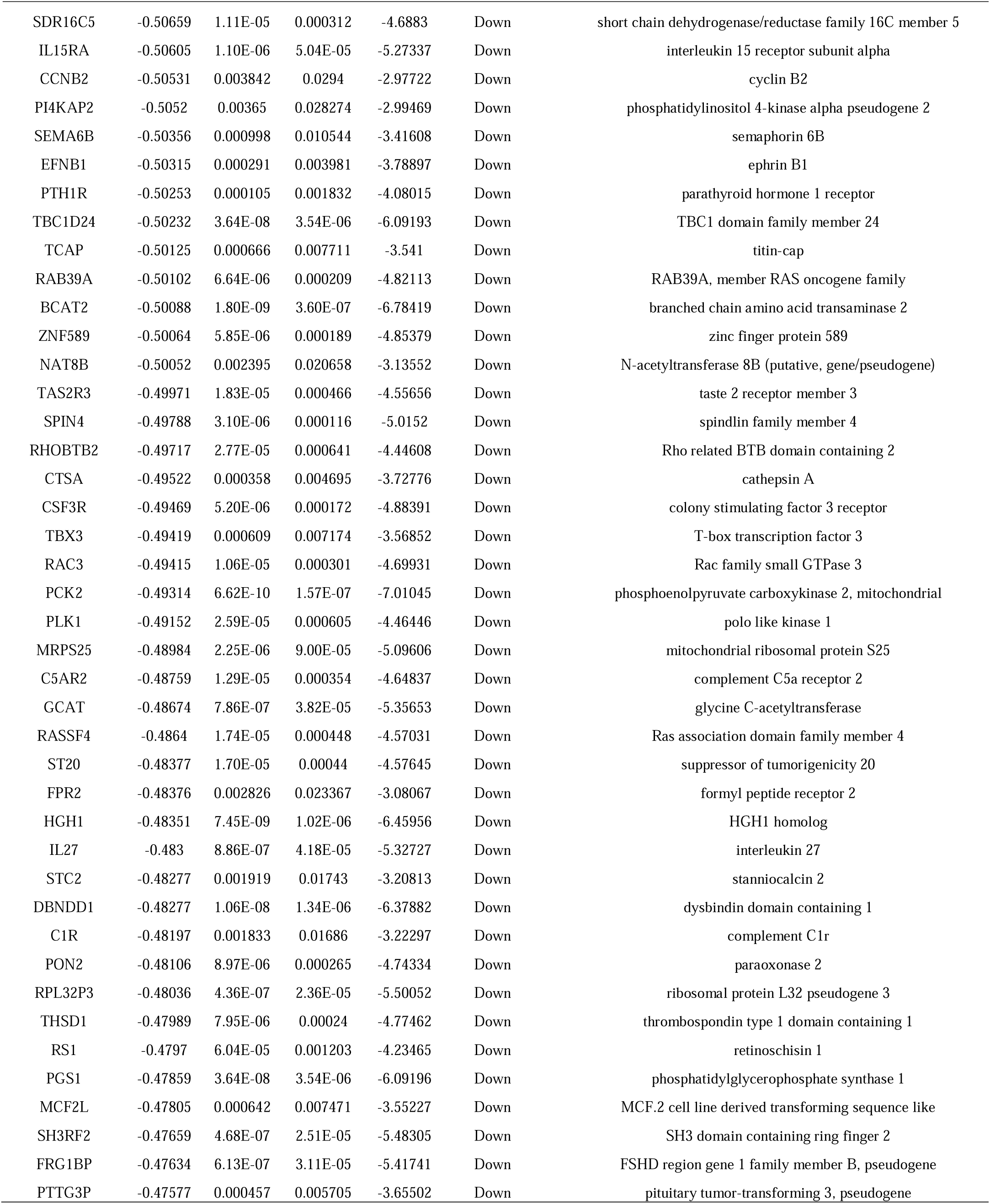
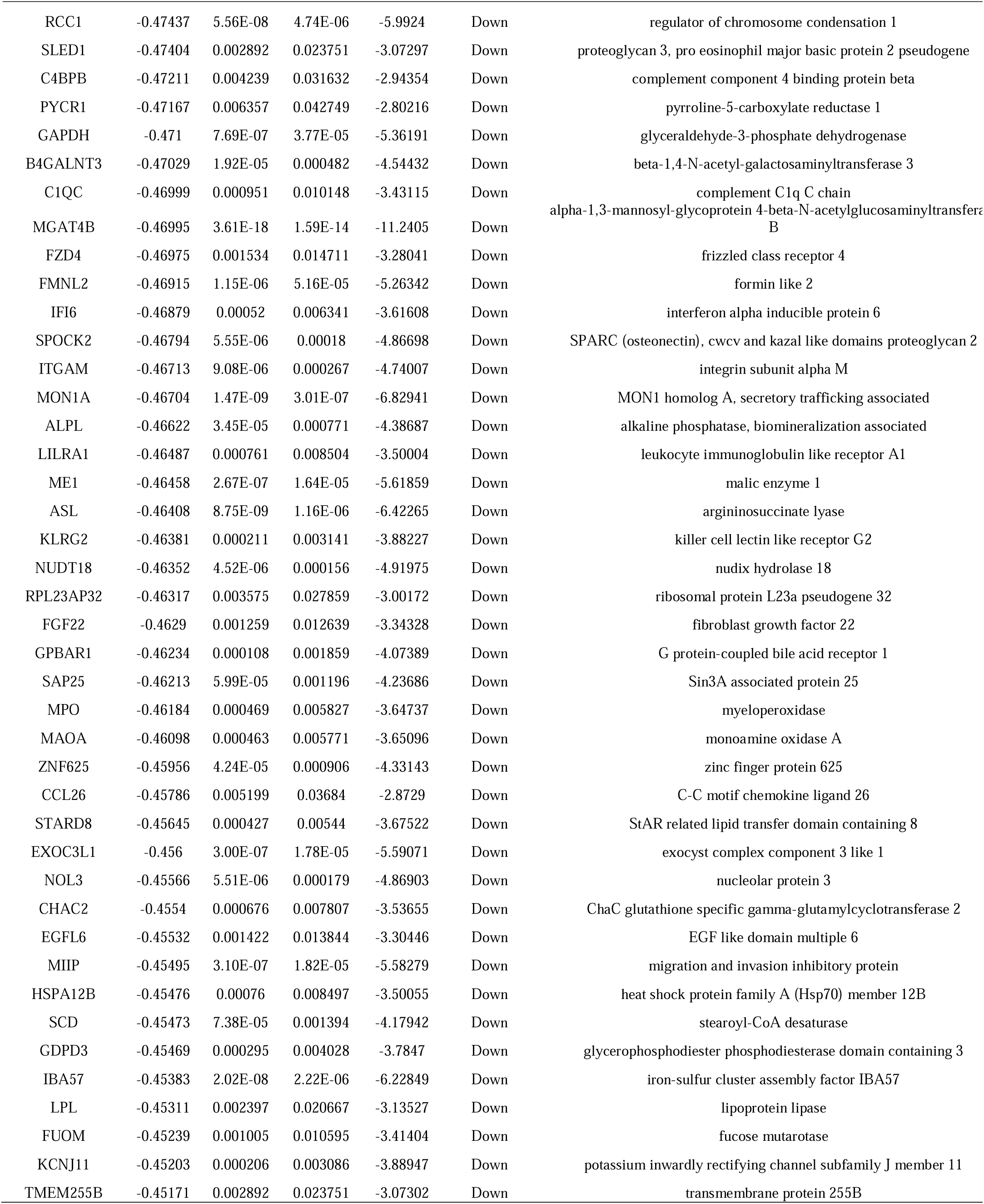
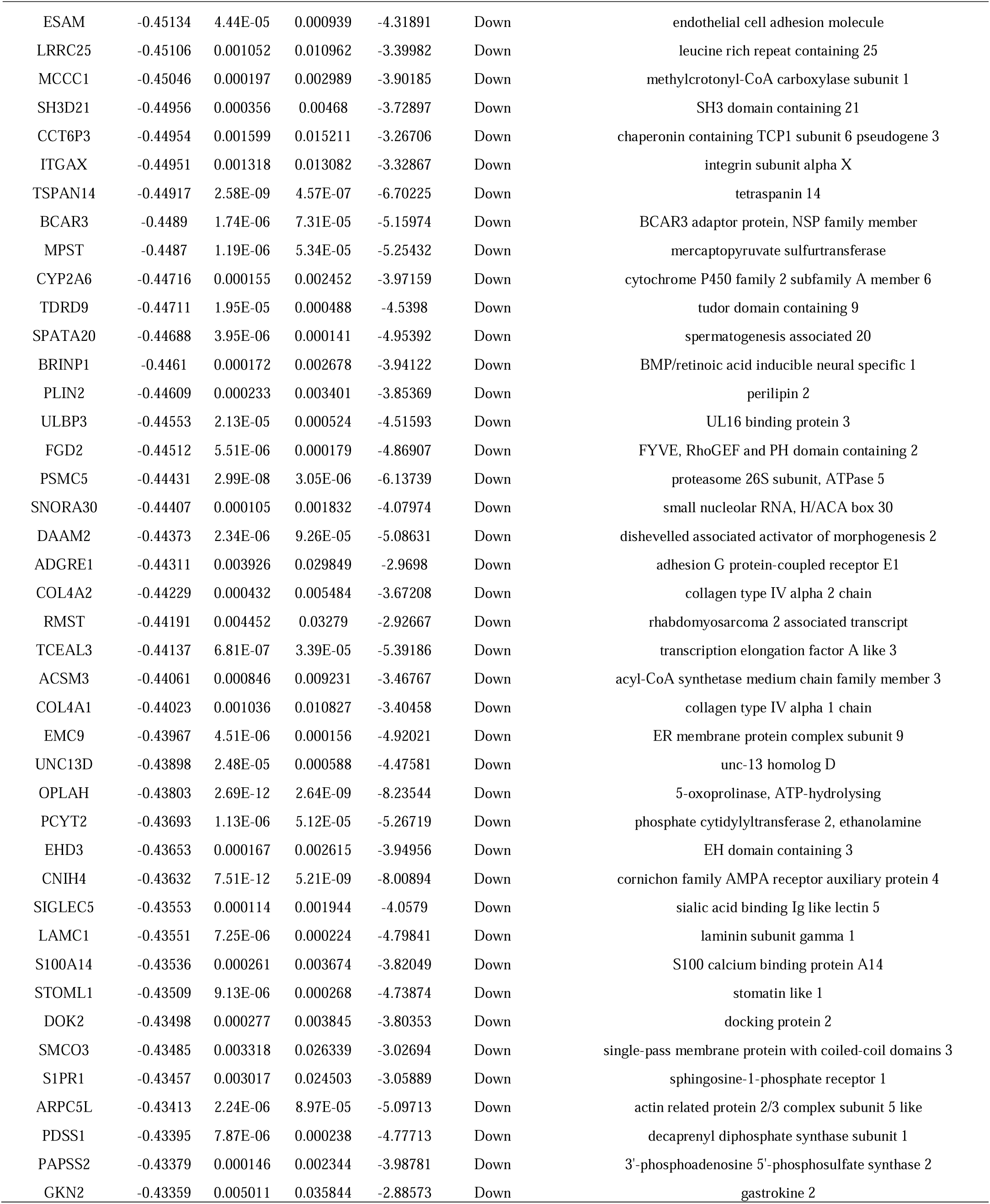
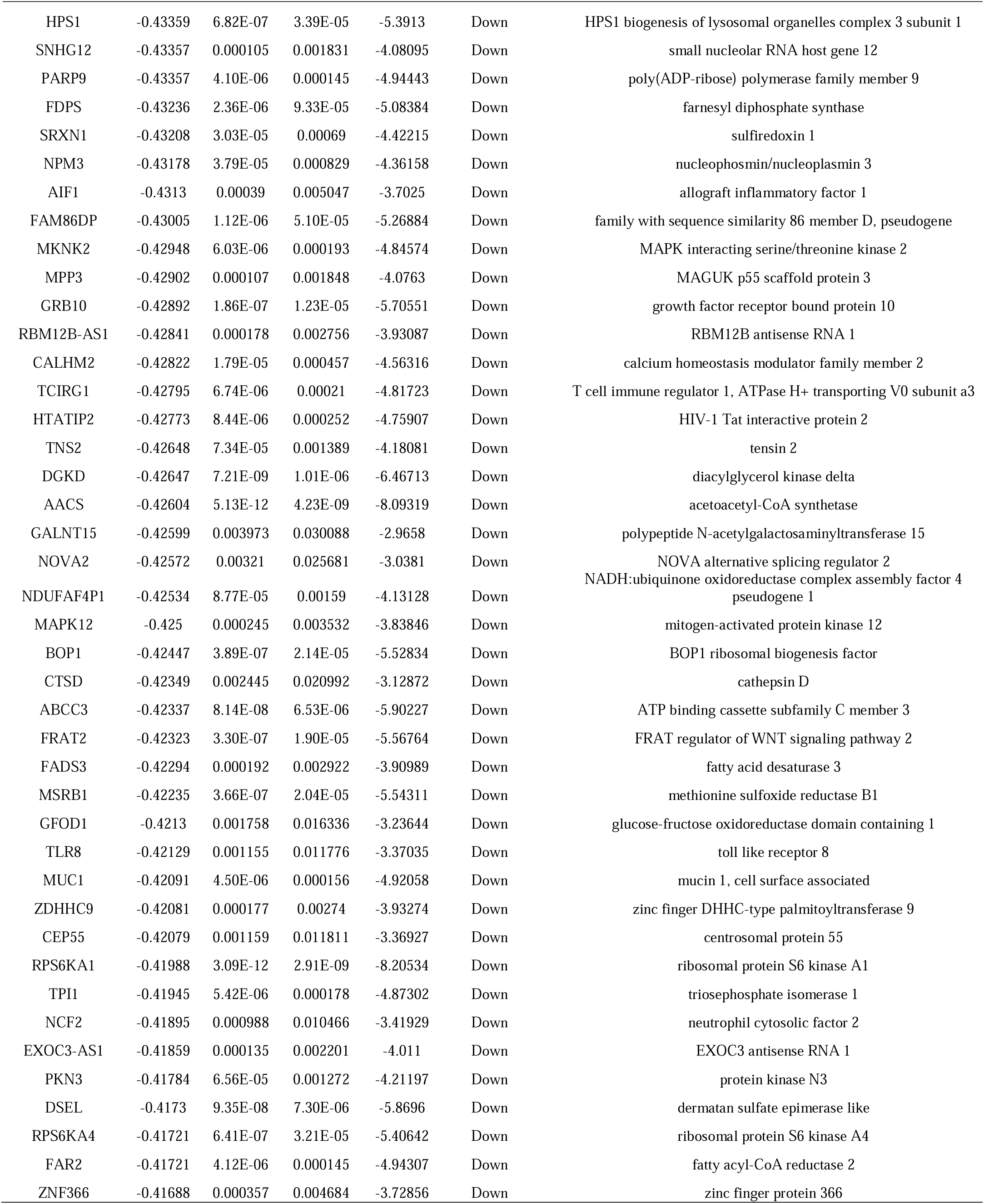
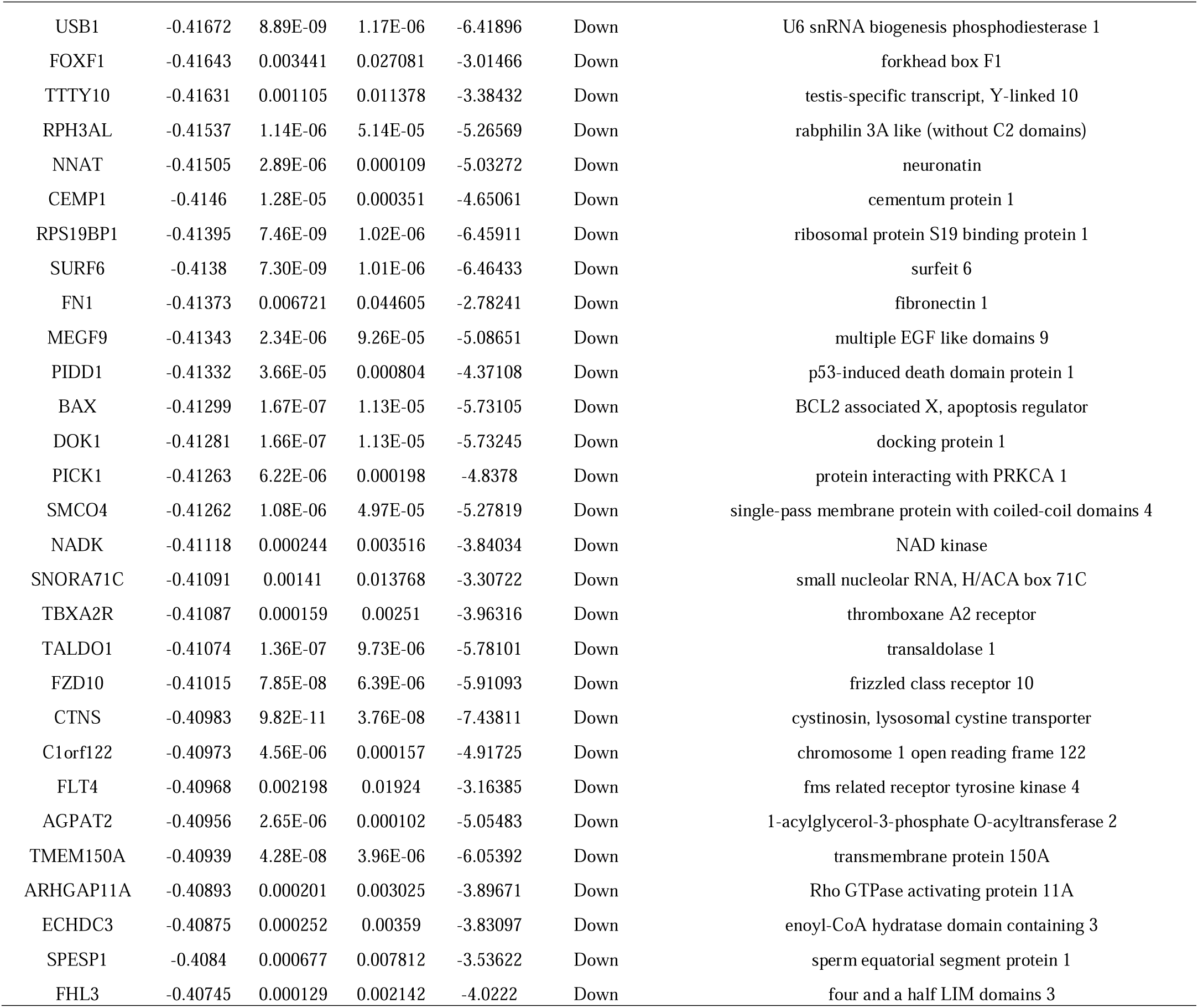
The statistical metrics for key differentially expressed genes (DEGs)

**Fig. 1.**
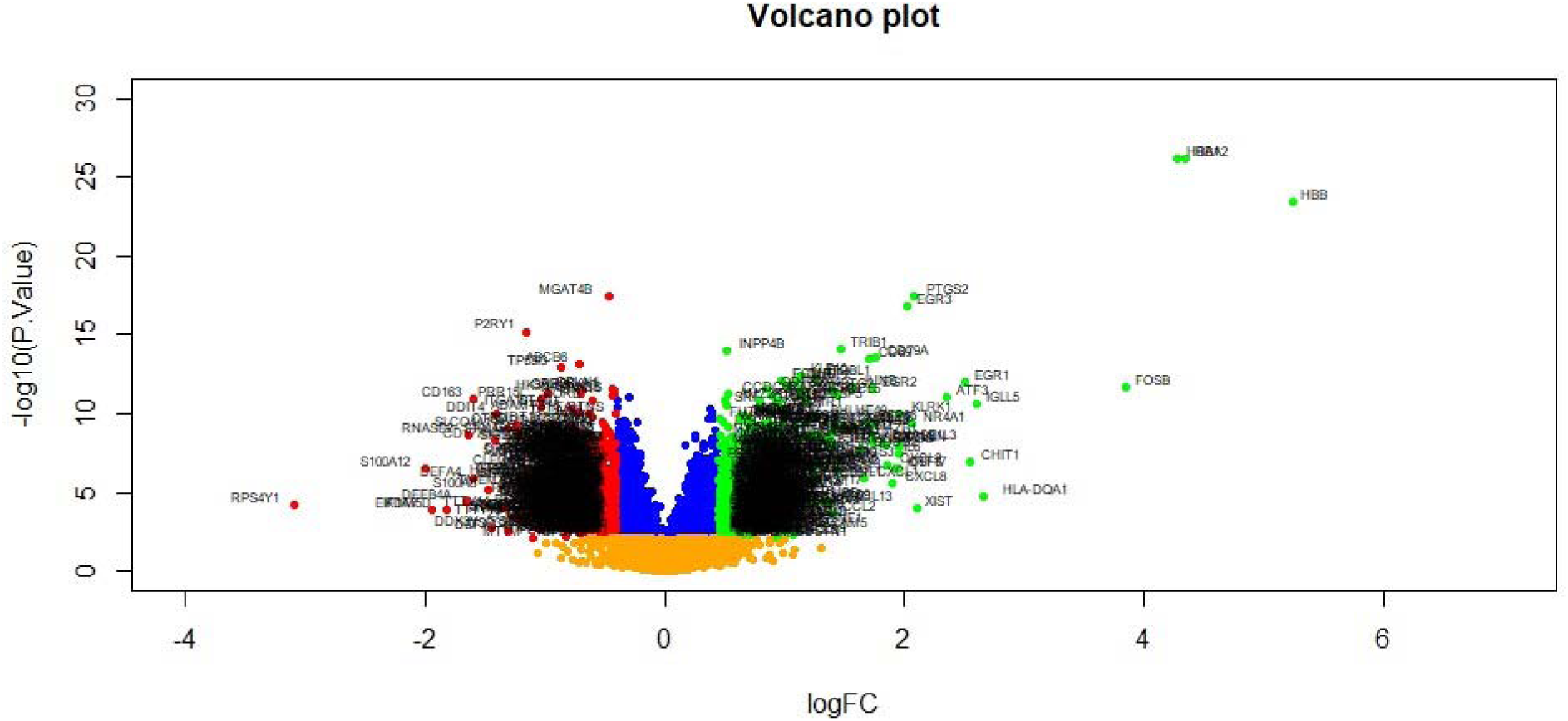
Volcano plot of differentially expressed genes. Genes with a significant change of more than two-fold were selected. Green dot represented up regulated significant genes and red dot represented down regulated significant genes.

**Fig. 2.**
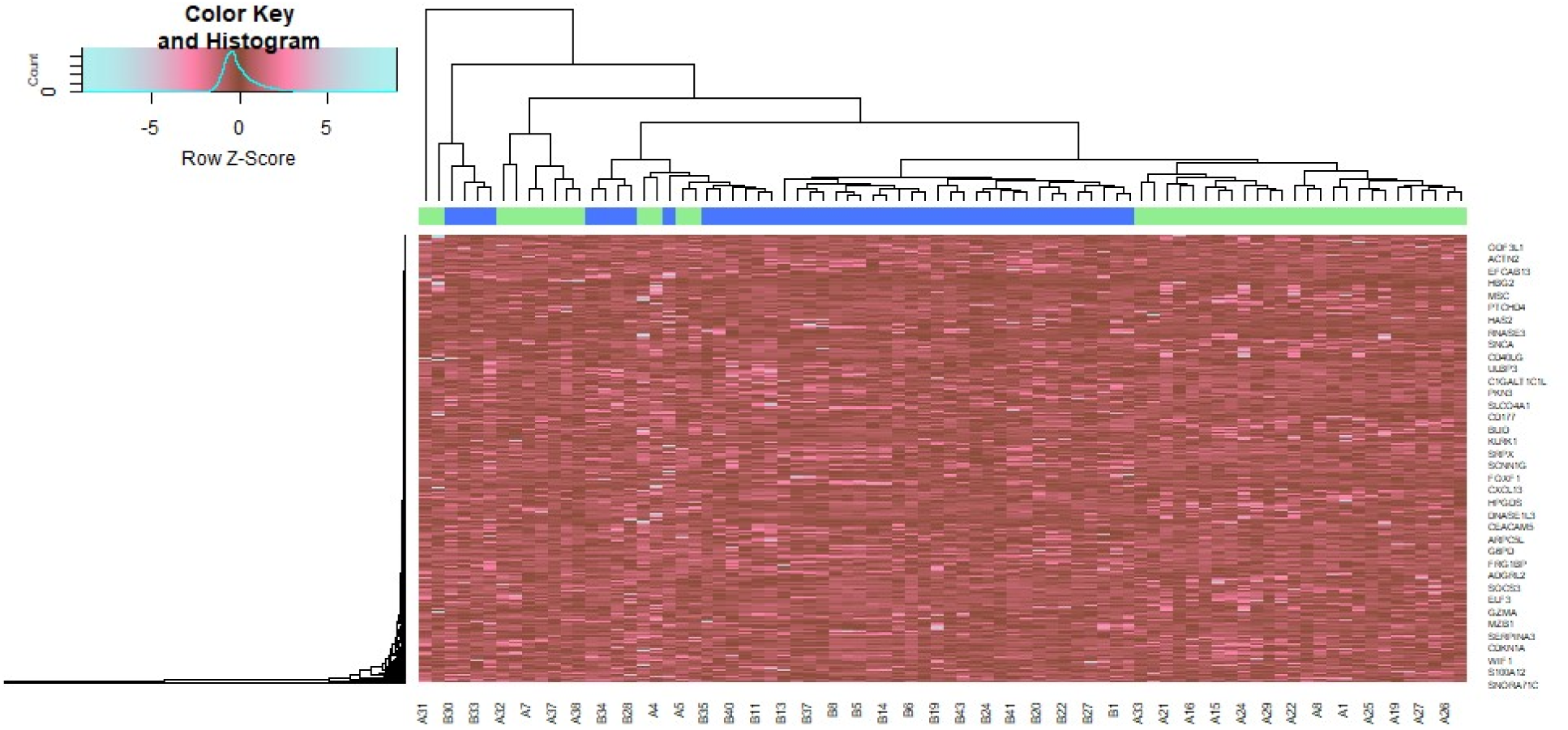
Heat map of differentially expressed genes. Legend on the top left indicate log fold change of genes. (A1 – A39 = COPD samples; B1 – B 43 = Normal control samples)

### GO and pathway enrichment analyses of DEGs

We performed GO terms and REACTOME pathway enrichment analysis of DEGs in the COPD samples and normal control samples. In GO BP analysis, we found that DEGs were rich in cellular response to stimulus, signaling, response to stimulus and cell communication (Table 2). In GO CC analysis, DEGs were rich in extracellular region, cell periphery, endomembrane system and membrane (Table 2). In GO MF analysis, DEGs were mainly rich in signaling receptor binding, molecular function regulator activity, catalytic activity and small molecule binding (Table 2). In REACTOME pathway enrichment analysis, DEGs were primarily rich in immune system, signaling by interleukins, neutrophil degranulation and innate immune system (Table 3).

**Table 2.**
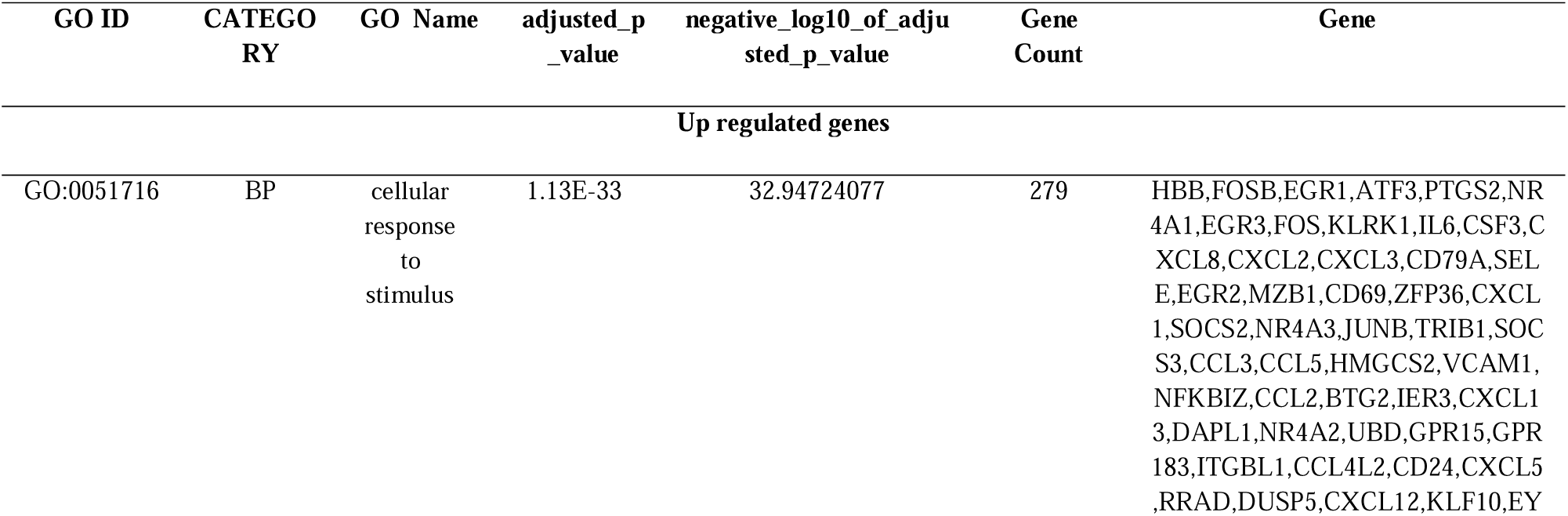
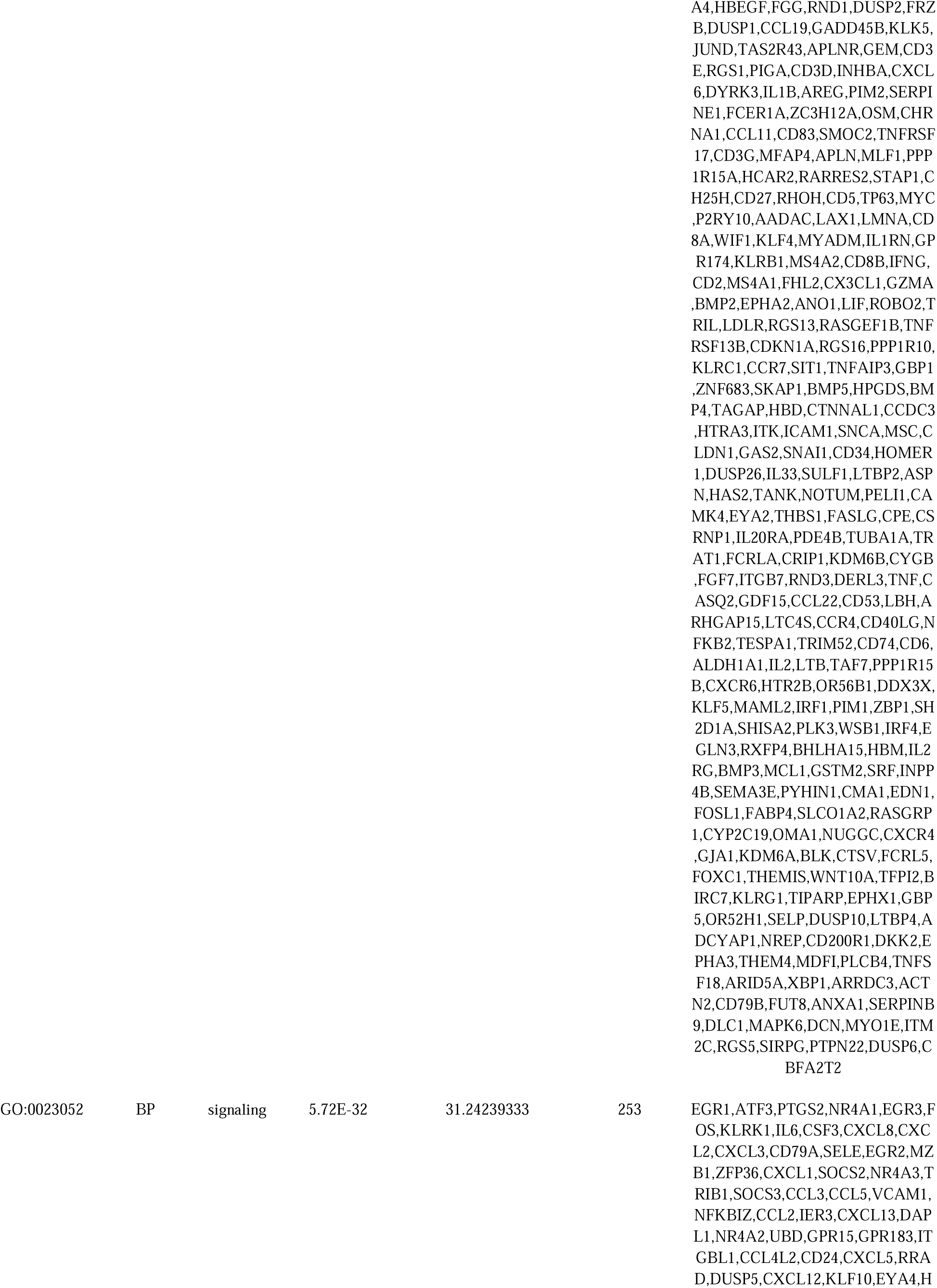
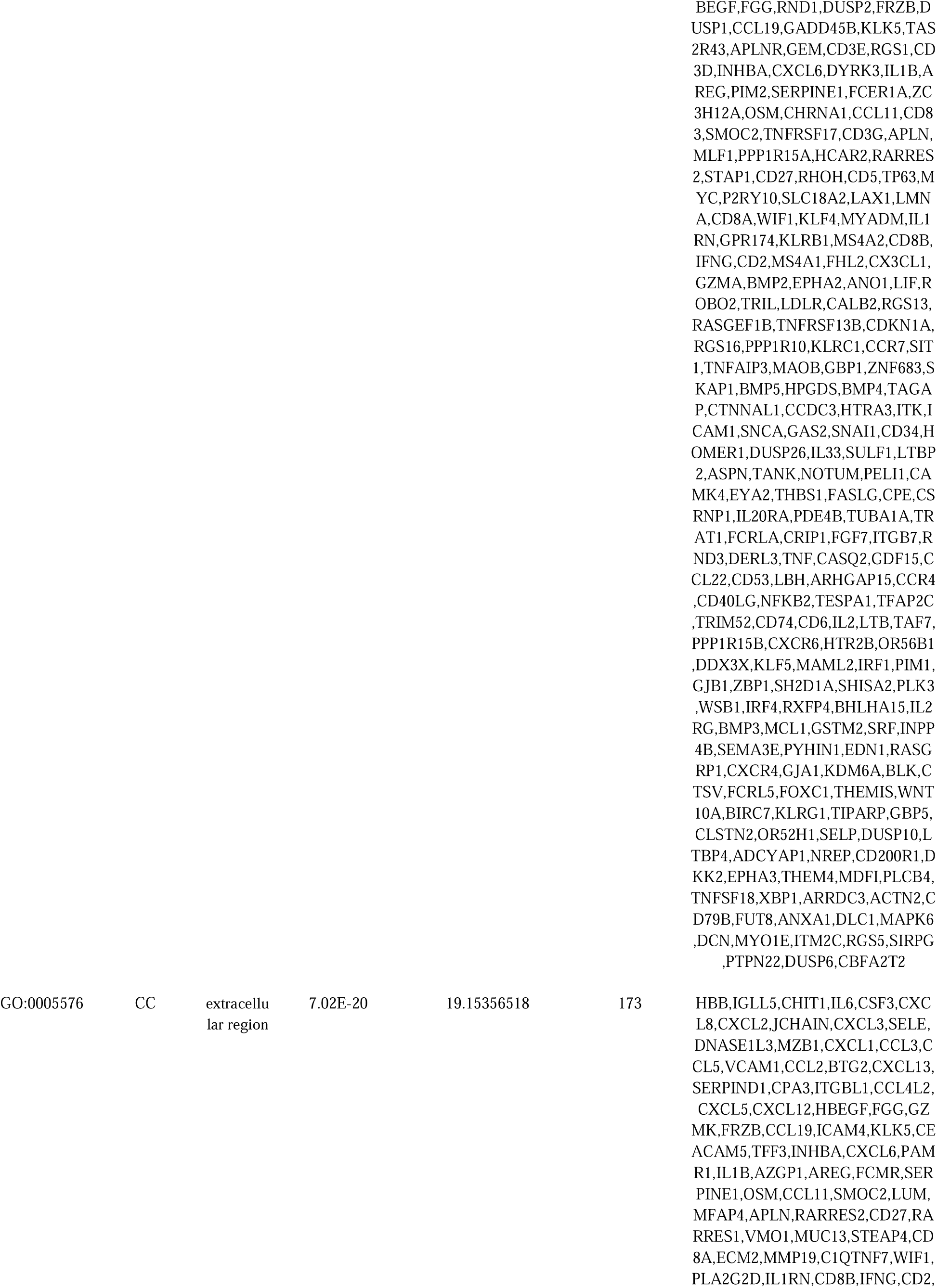
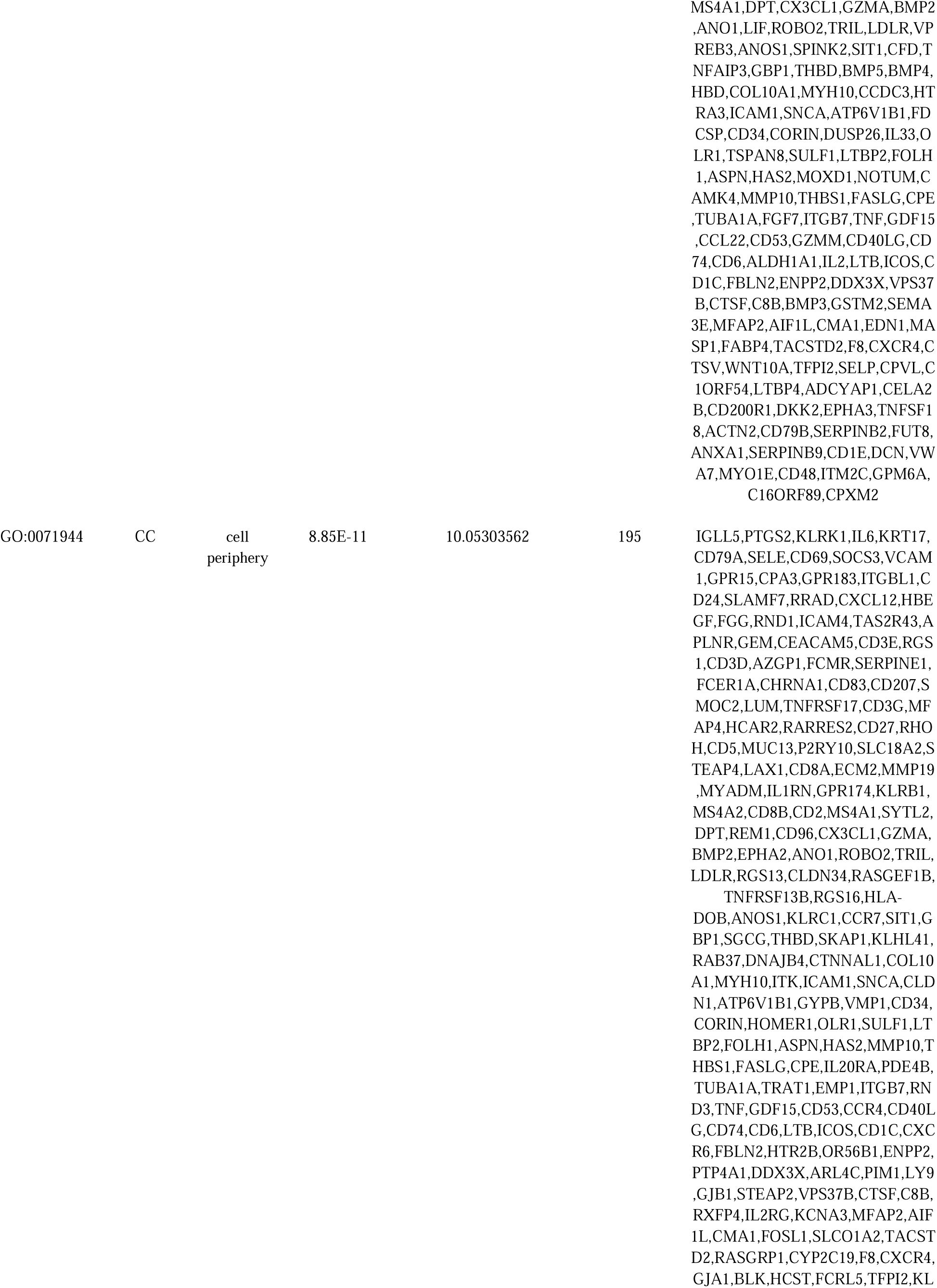
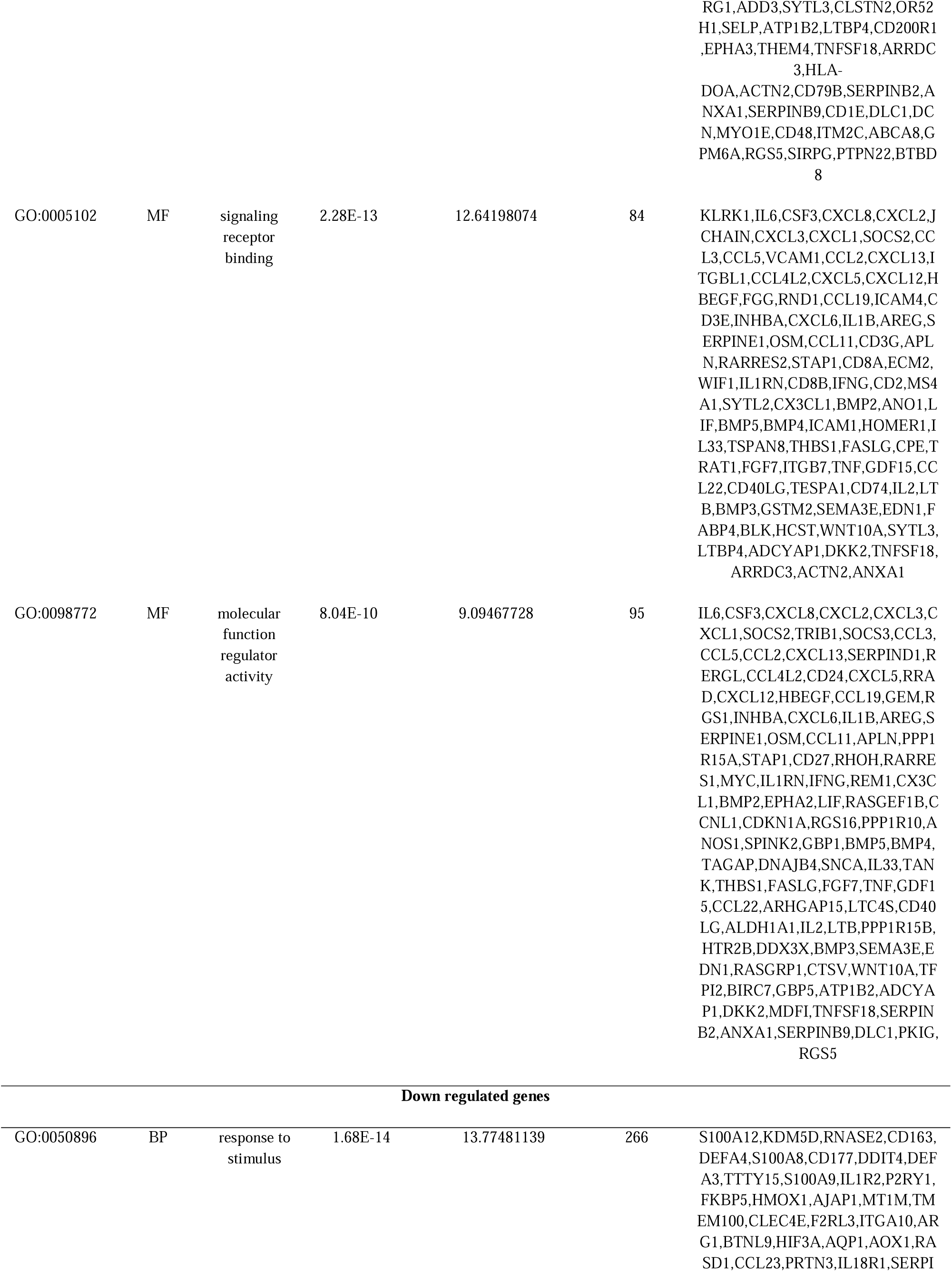
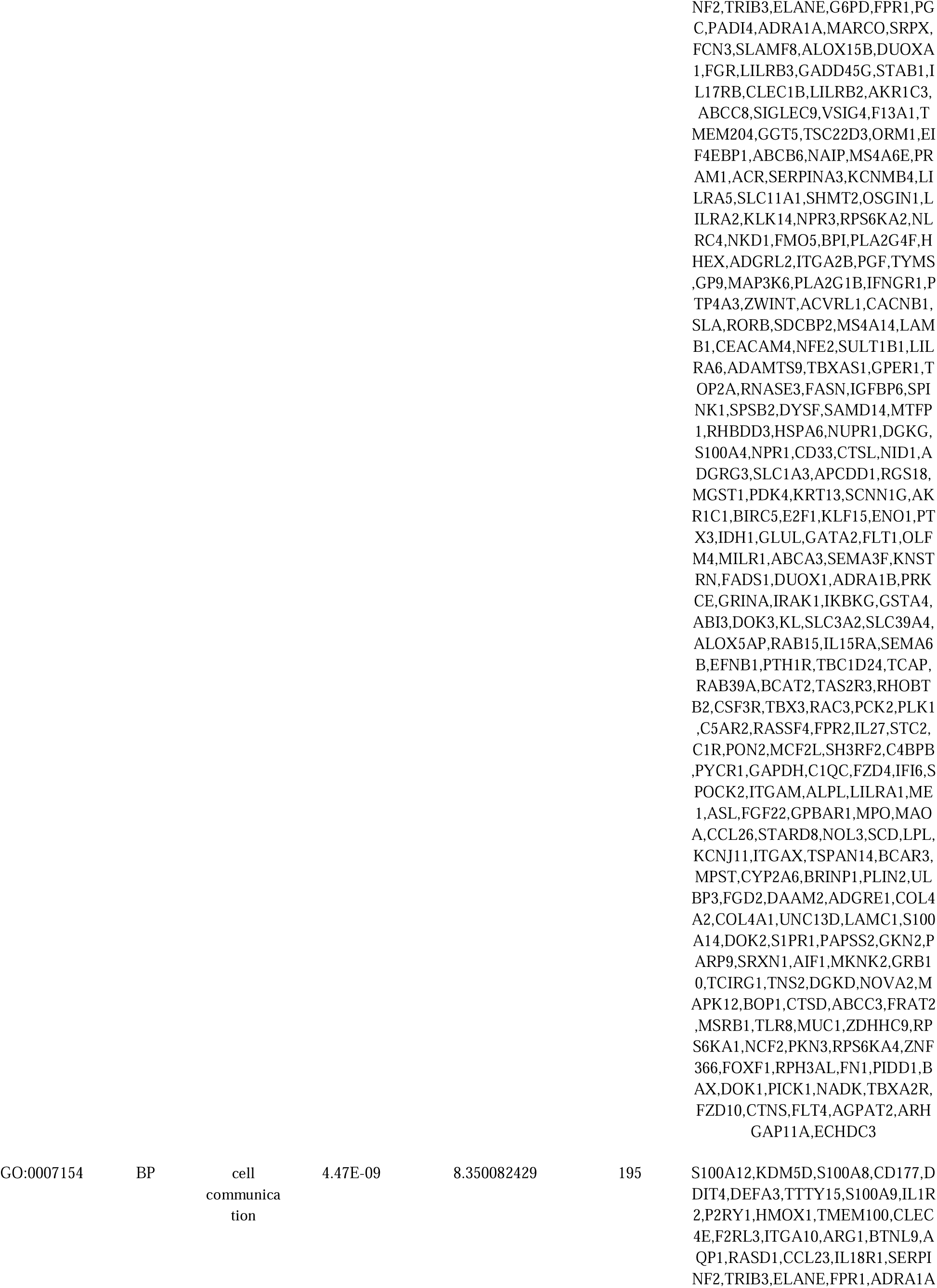
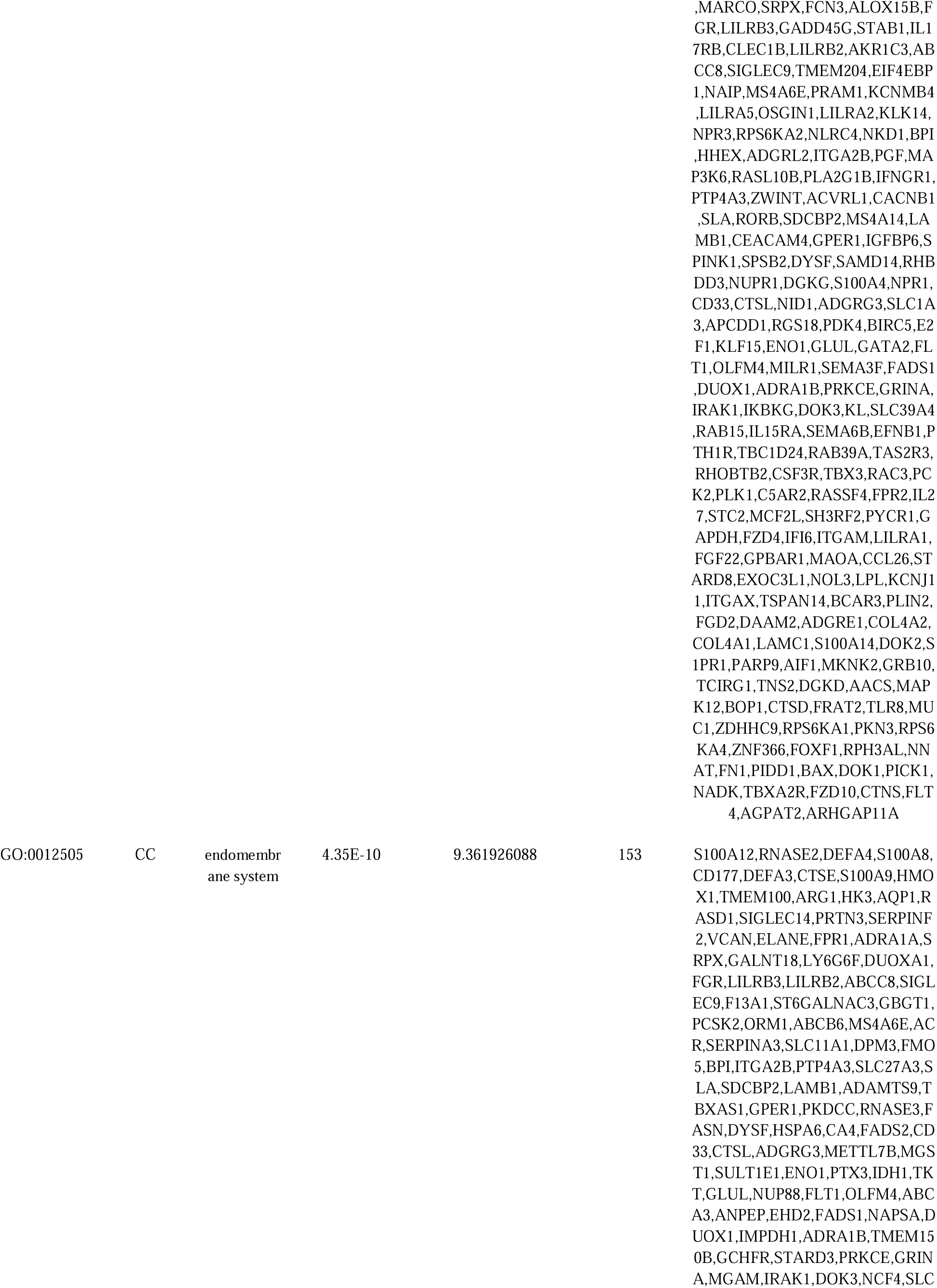
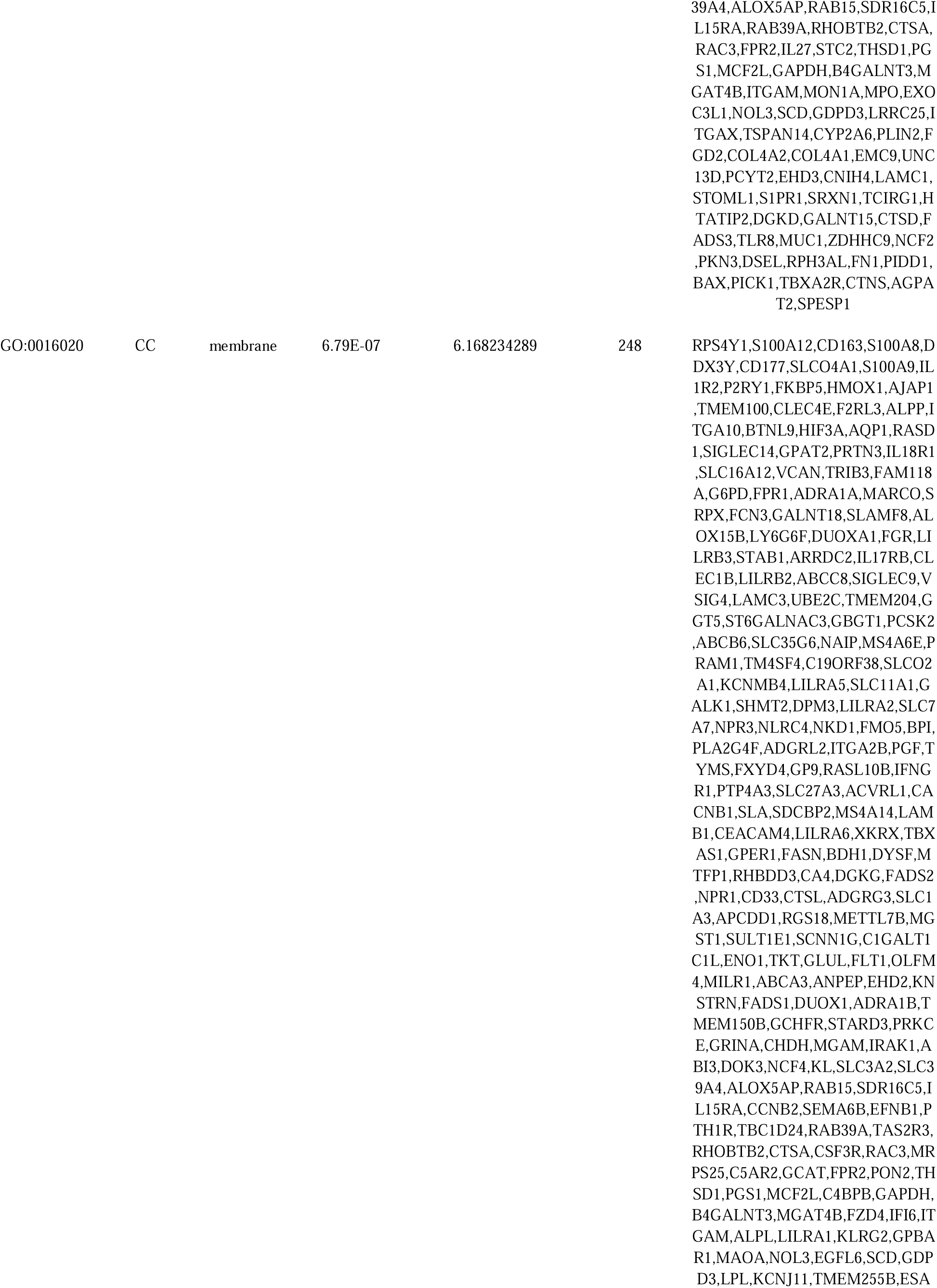
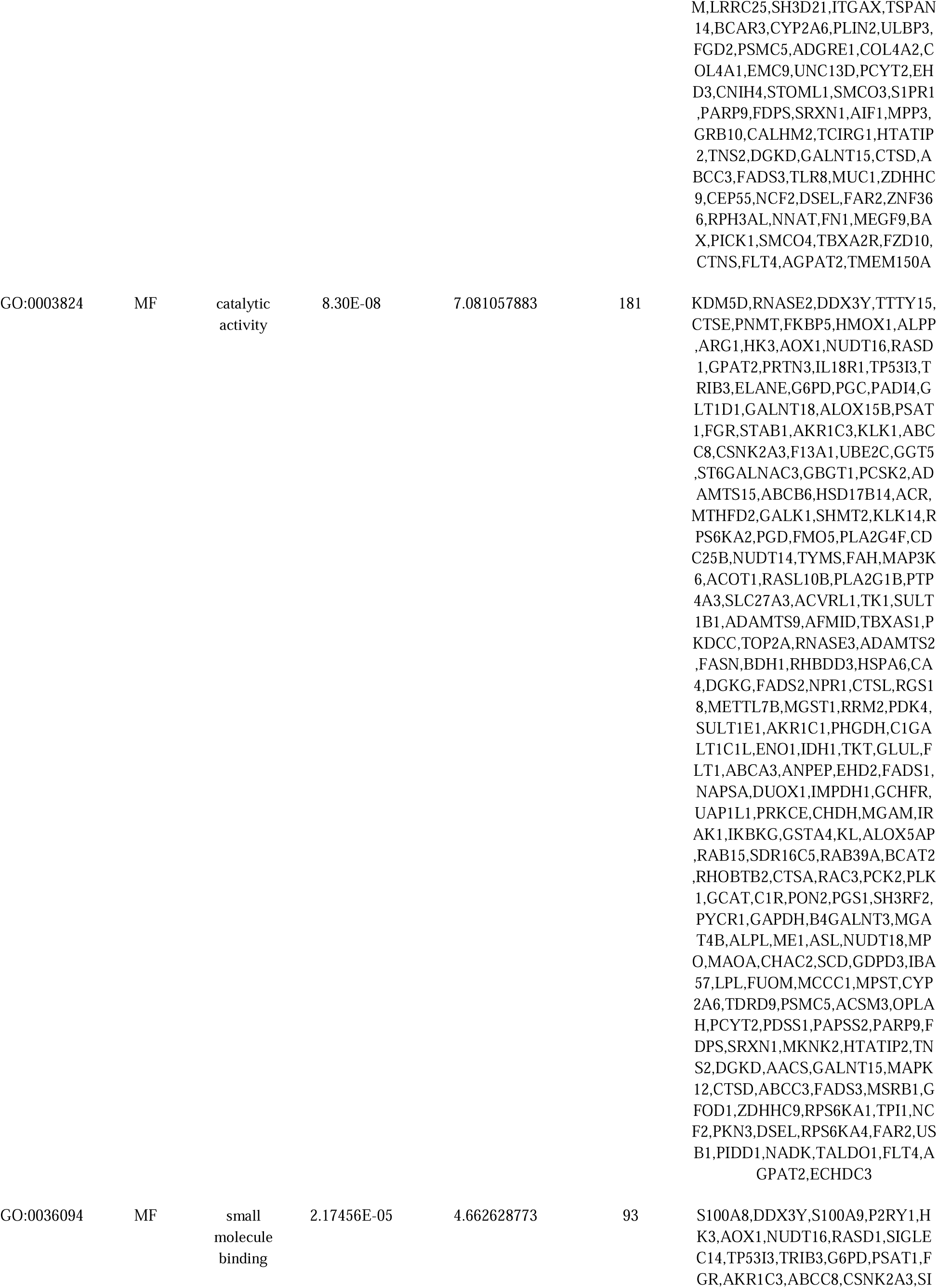
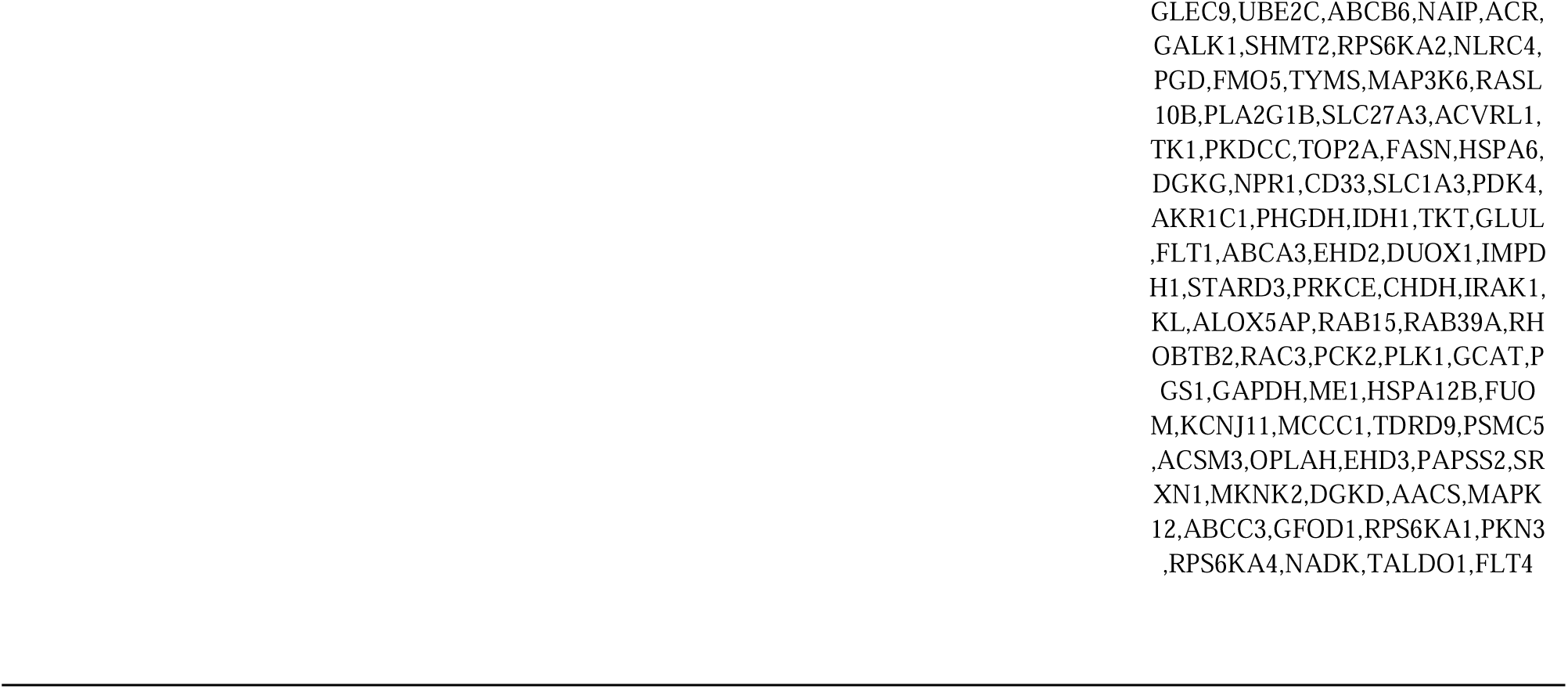
The enriched GO terms of the up and down regulated differentially expressed genes.

**Table 3.**
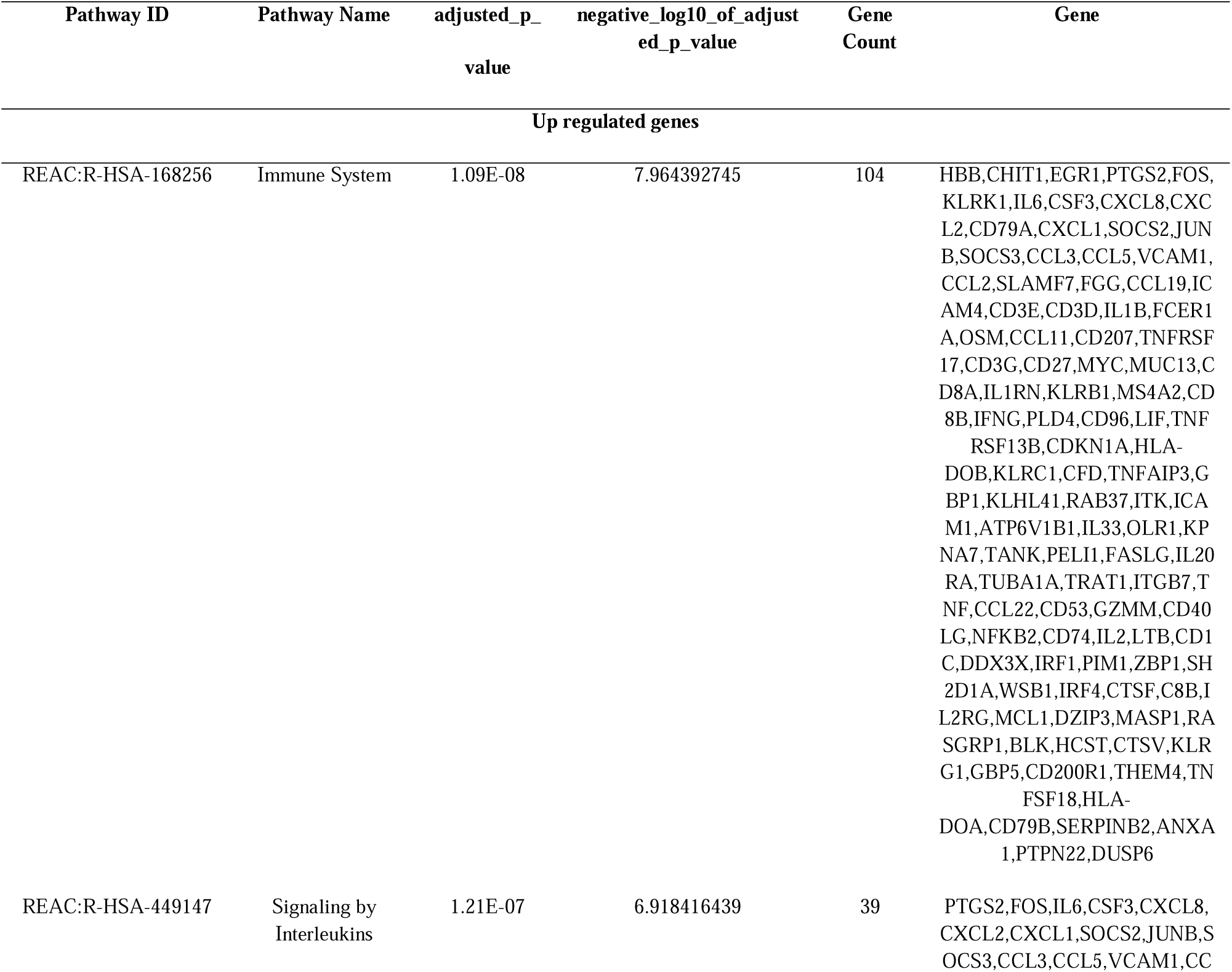
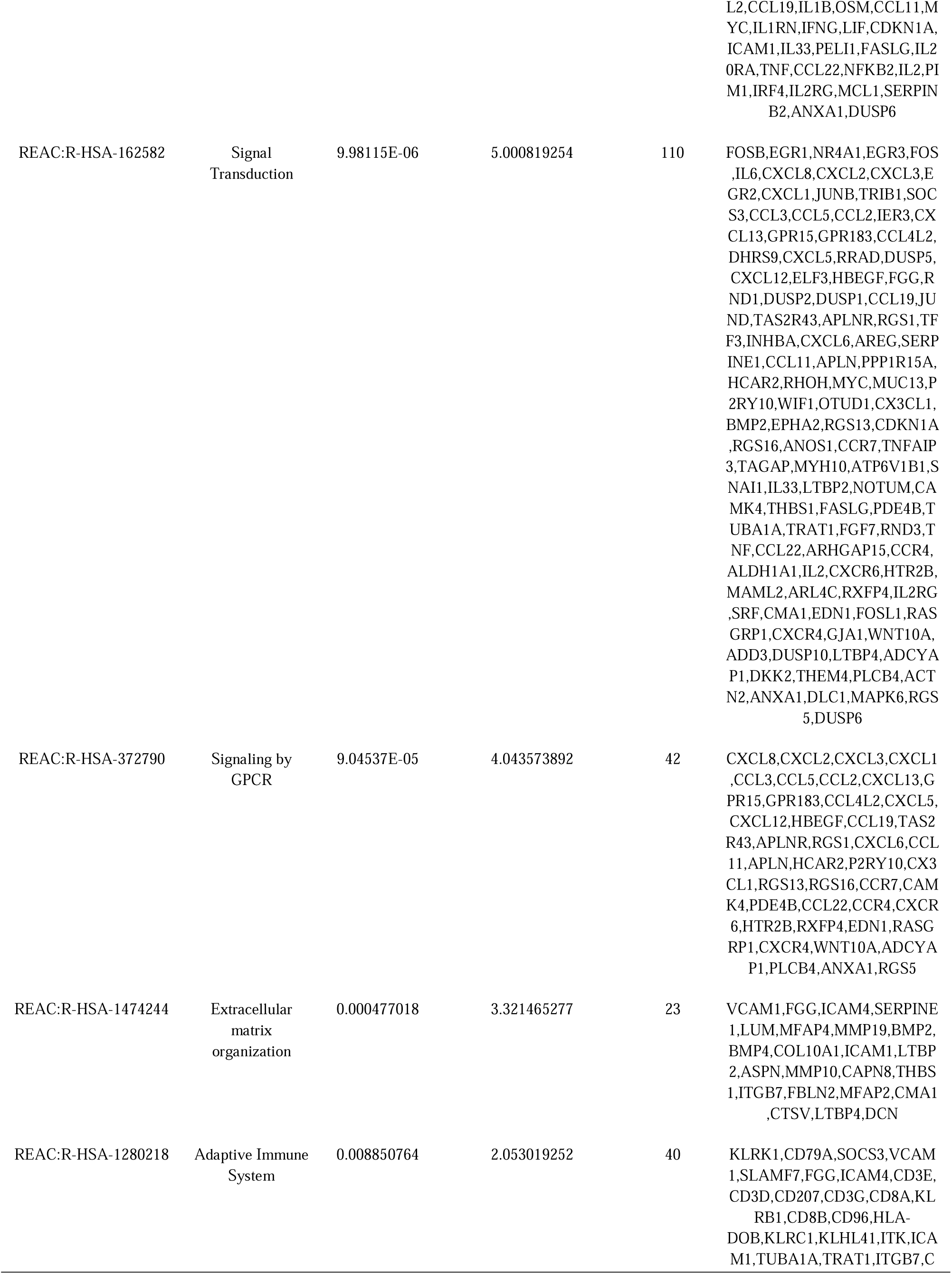
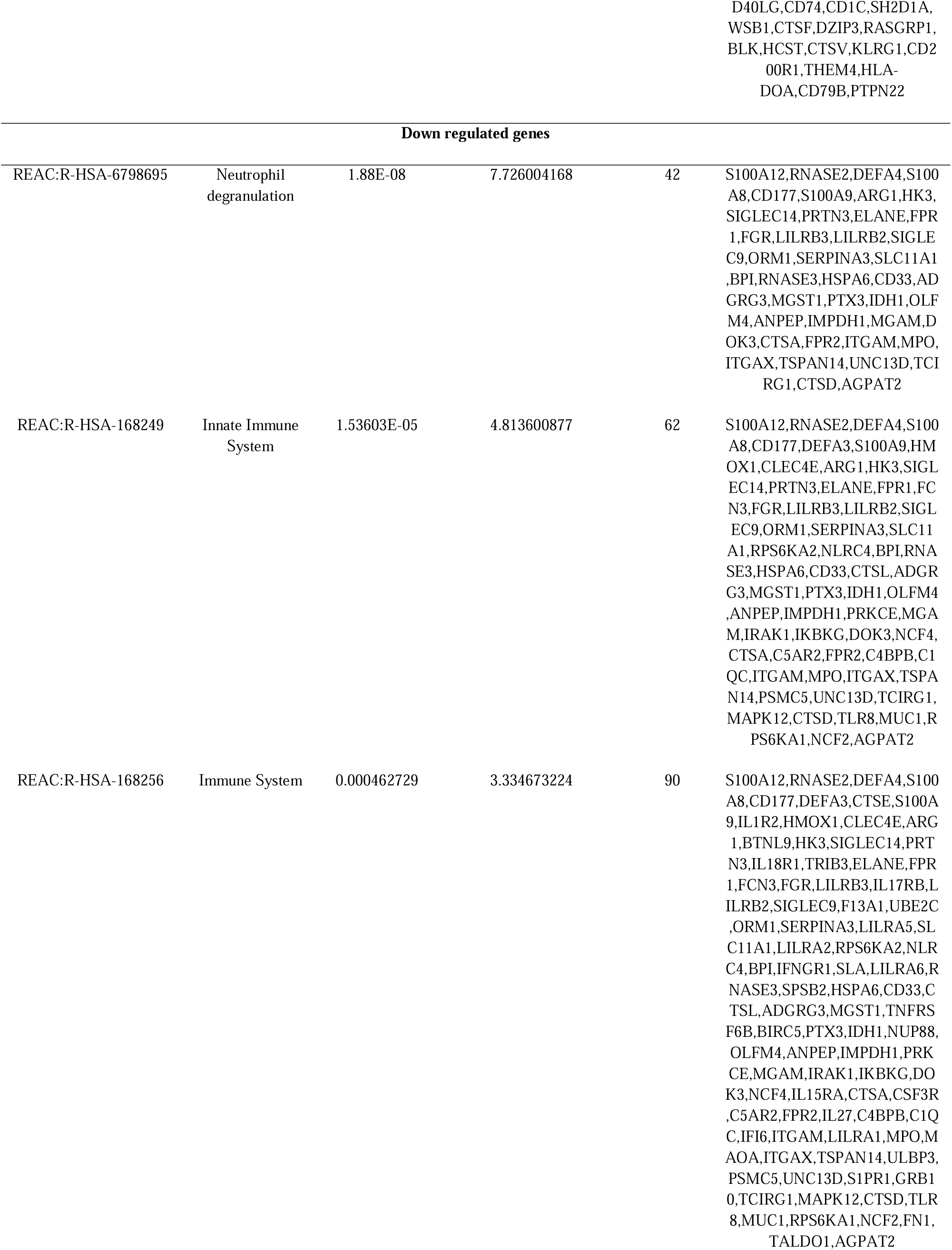
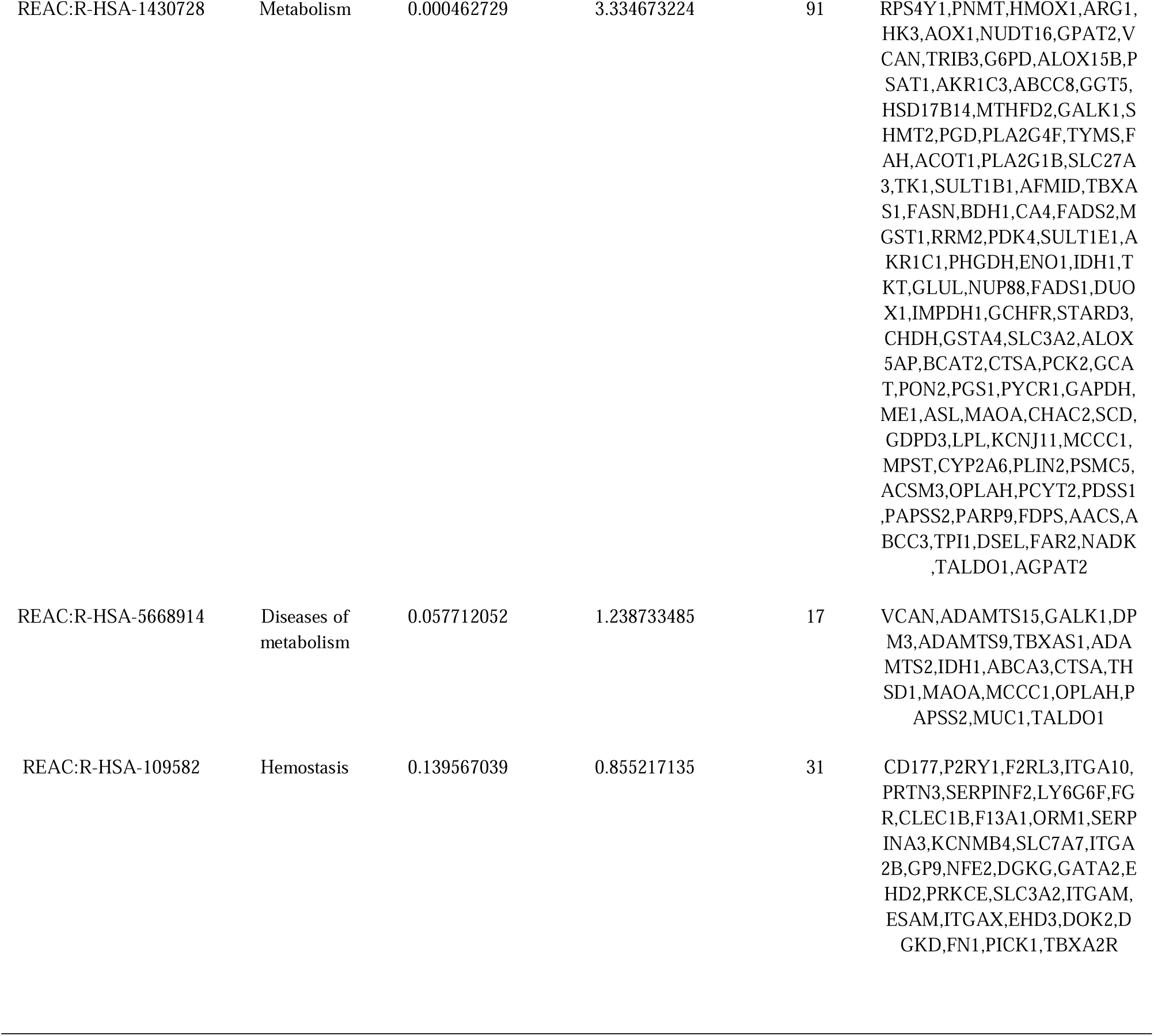
The enriched pathway terms of the up and down regulated differentially expressed genes.

### Construction of the PPI network and module analysis

Using the pickle platform, PPI analysis of these DEGs identified 6497 nodes and 13223 interactions. MYC, LMNA, VCAM1, MAPK6, DDX3X, SHMT2, PHGDH, S100A9, FKBP5 and RPS6KA2 were hub nodes in PPI network (Table 4). And the most significant module 1 consisting of 88 nodes and 228 edges (Fig. 4A) and module 2 consisting of 101 nodes and 107 edges (Fig 4B) were found using the PEWCC plug-in of Cytoscape software. In GO terms and REACTOME pathway enrichment analysis, hub genes were rich in the cellular response to stimulus, signaling, immune system, signaling by interleukins, signal transduction, molecular function regulator activity, cell periphery, neutrophil degranulation, innate immune system, response to stimulus, cell communication, endomembrane system, catalytic activity, metabolism, small molecule binding and hemostasis.

**Fig. 3.**
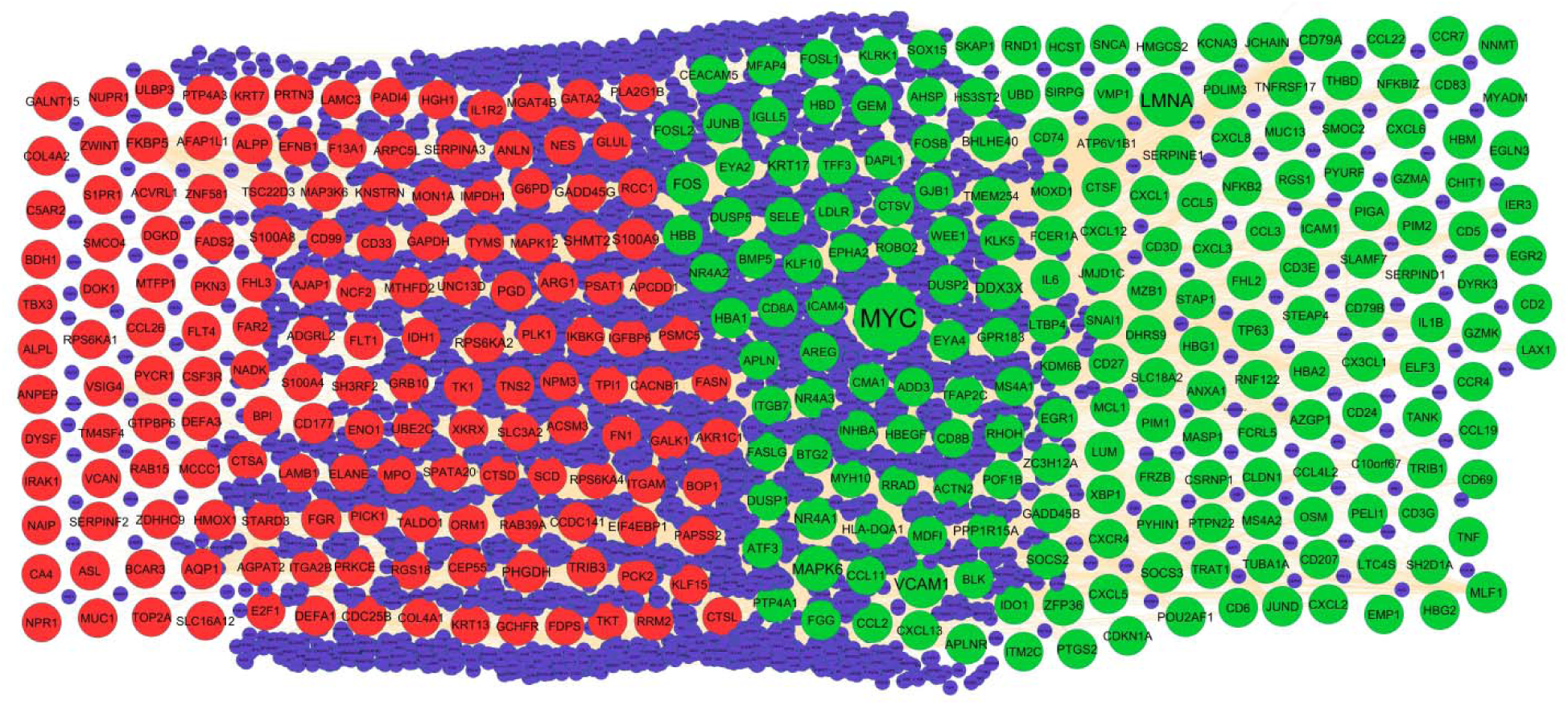
PPI network of DEGs. Up regulated genes are marked in parrot green; down regulated genes are marked in red.

**Fig. 4.**
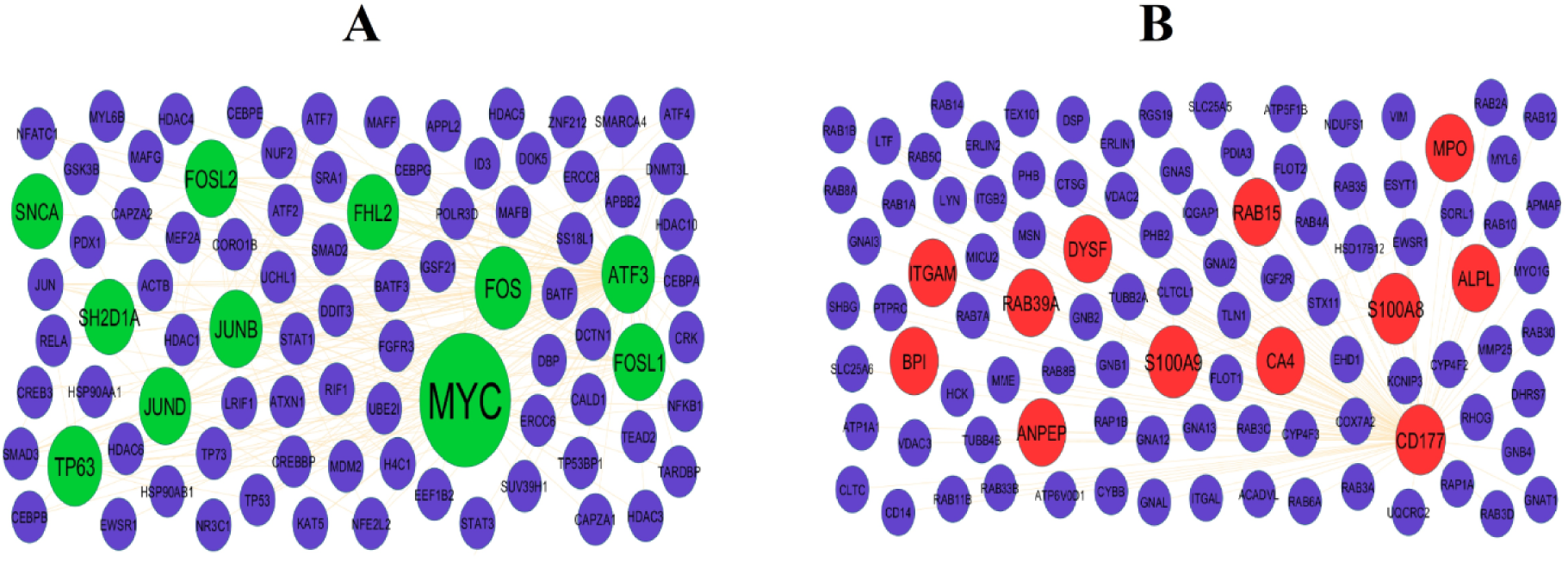
Modules selected from the PPI network. (A) The most significant module was obtained from PPI network with 88 nodes and 228 edges for up regulated genes (B) The most significant module was obtained from PPI network with 101 nodes and 107 edges for down regulated genes. Up regulated genes are marked in parrot green; down regulated genes are marked in red.

**Table 4.**
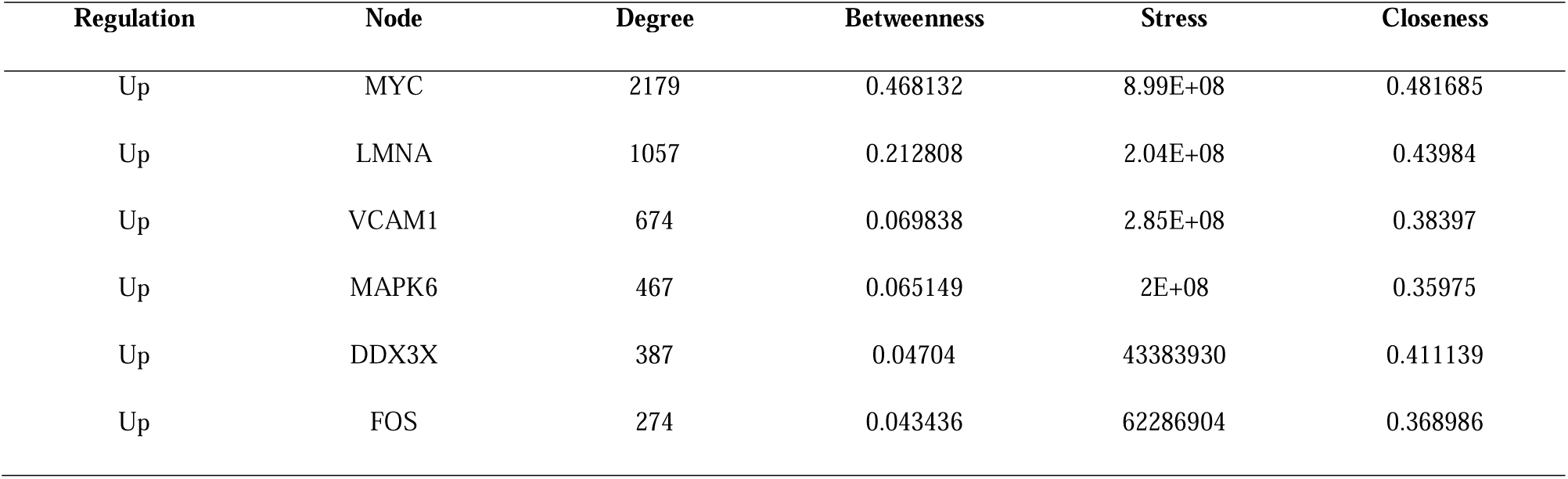
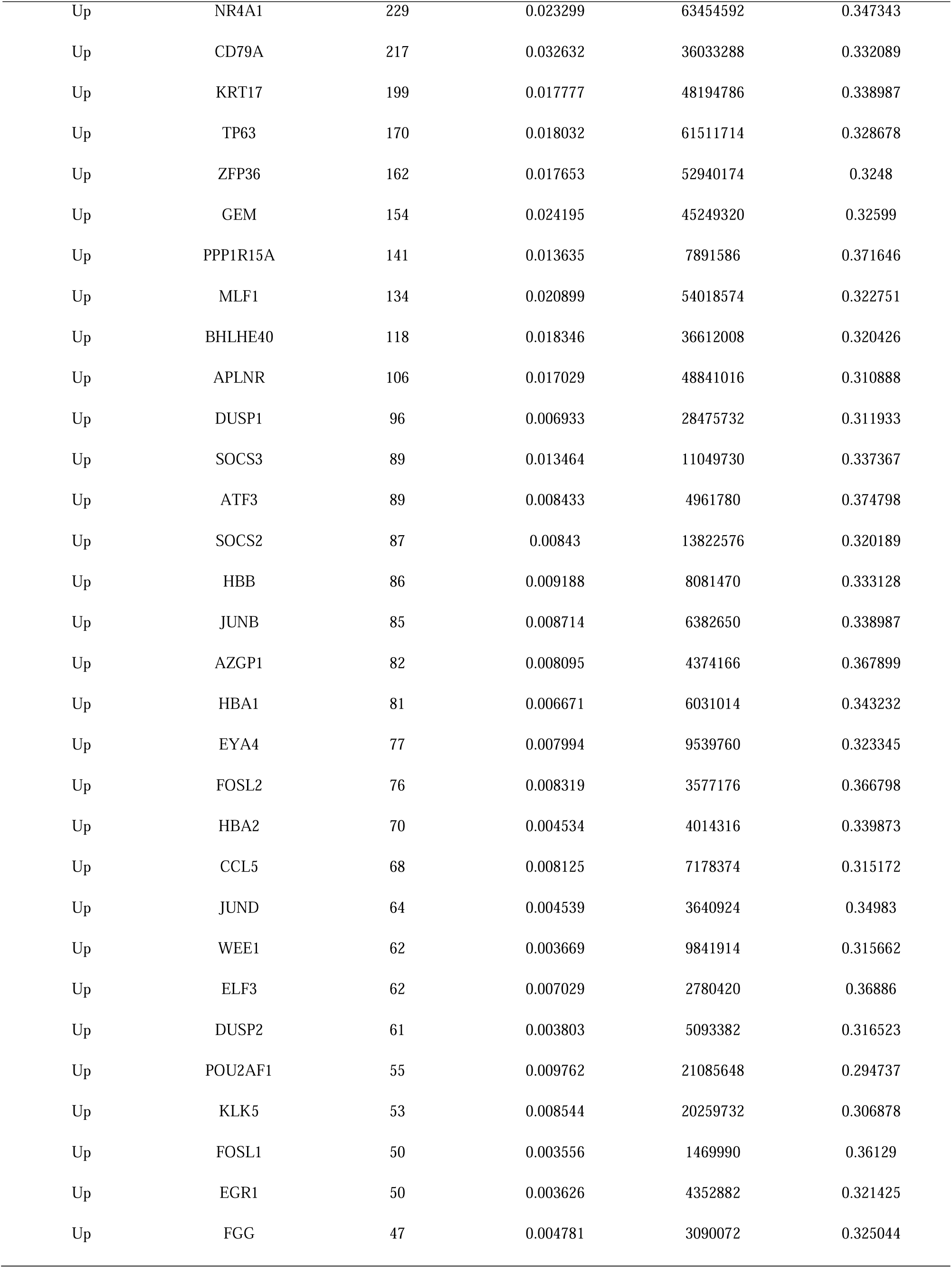
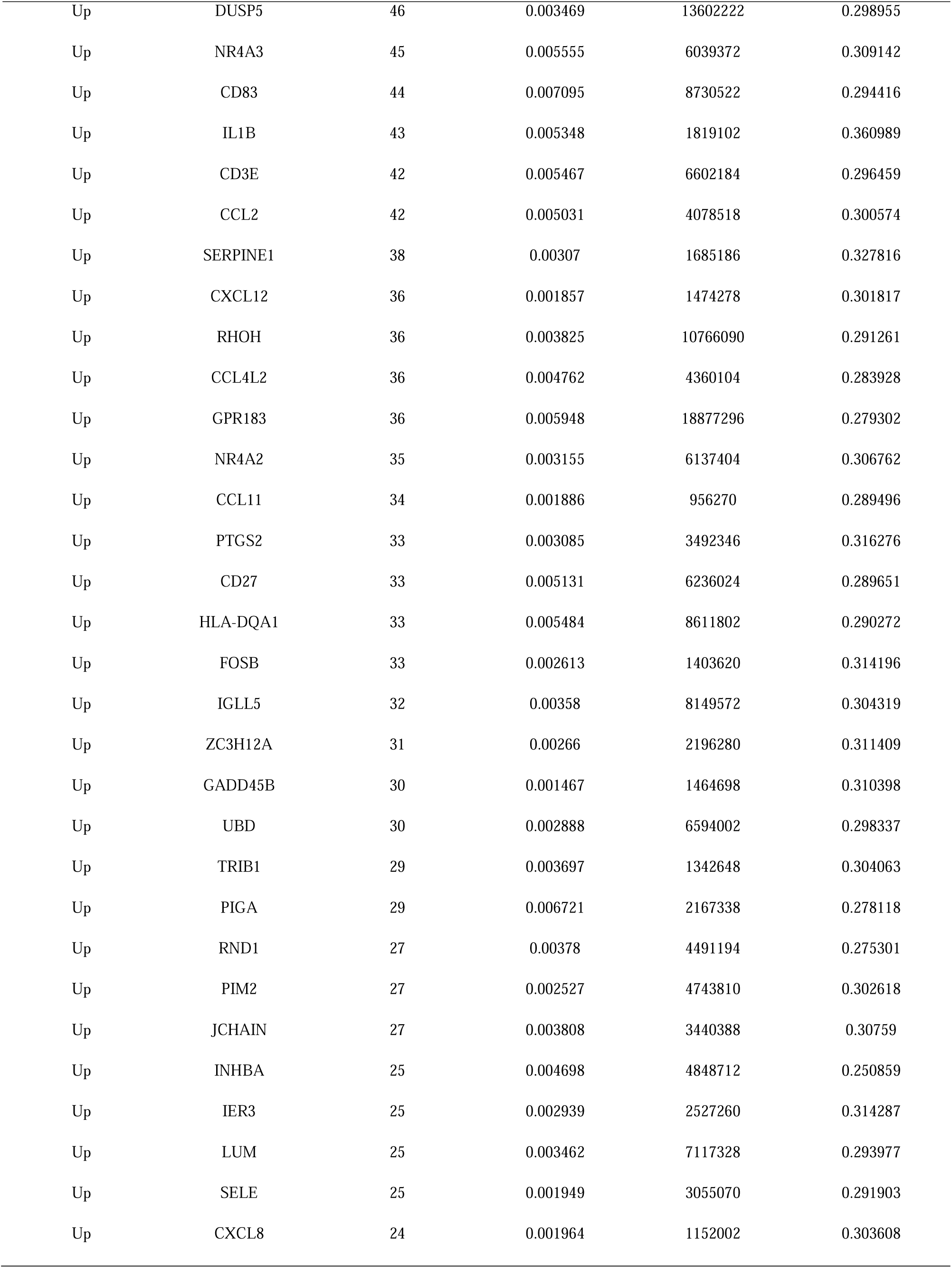
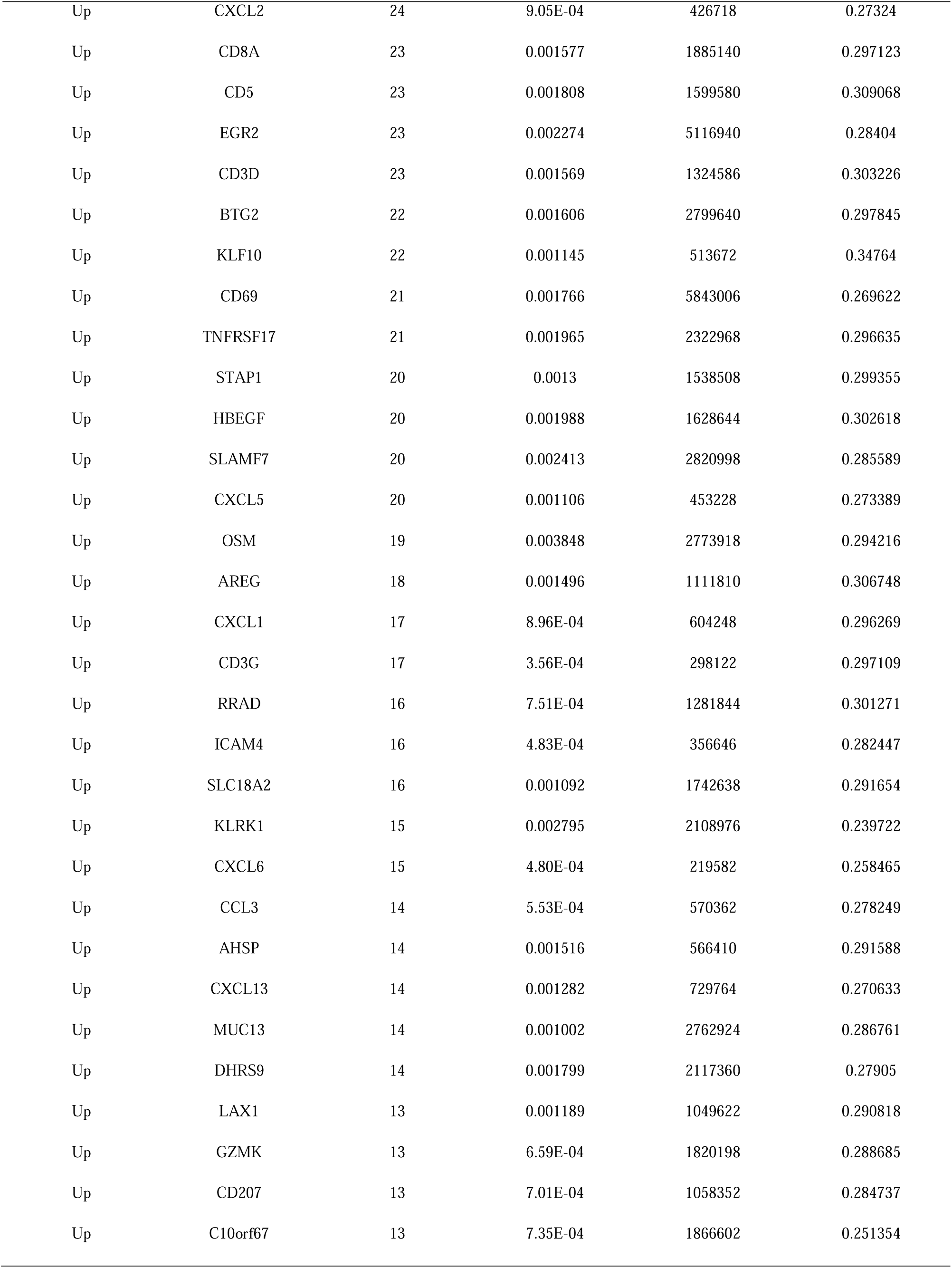
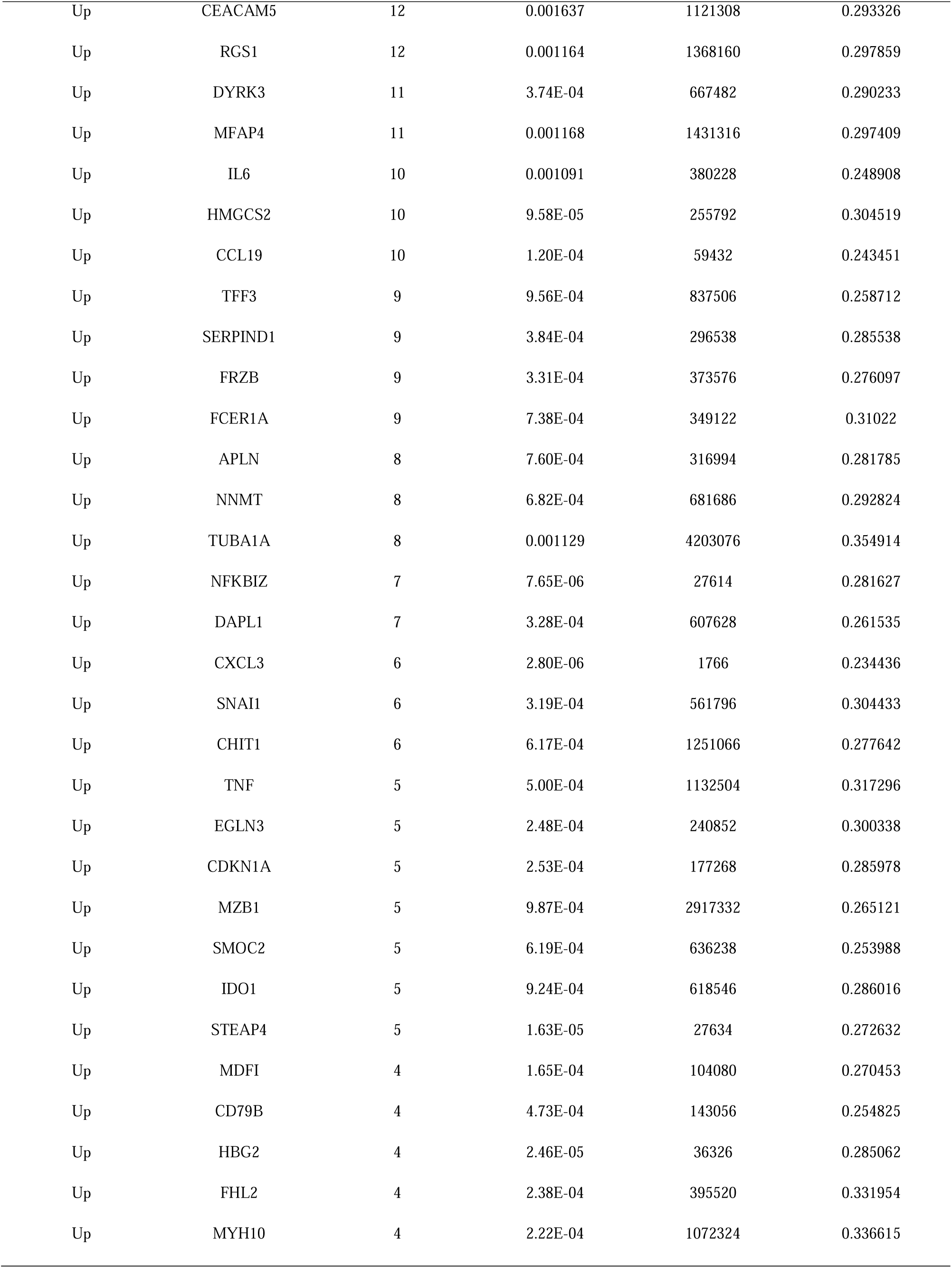
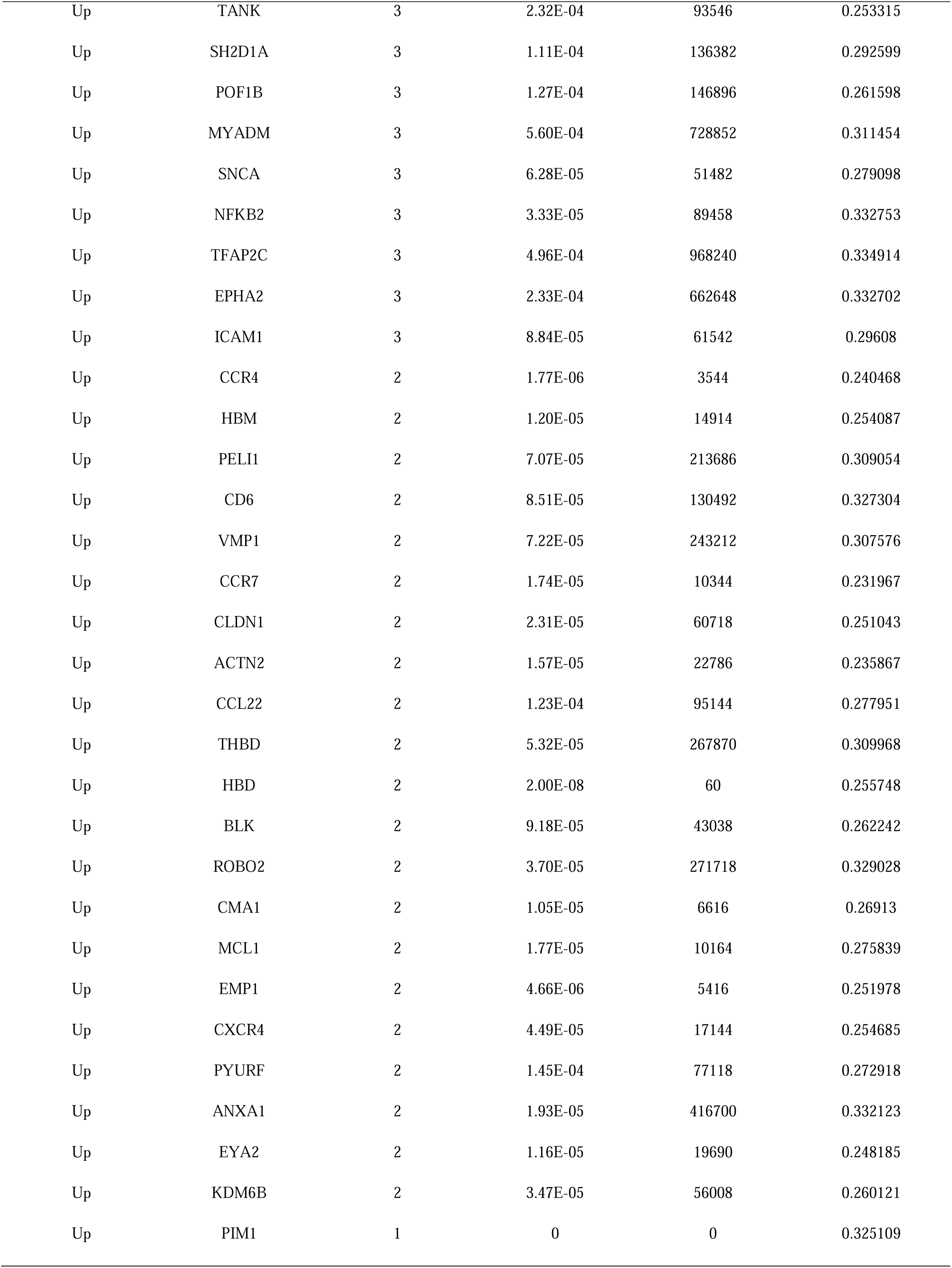
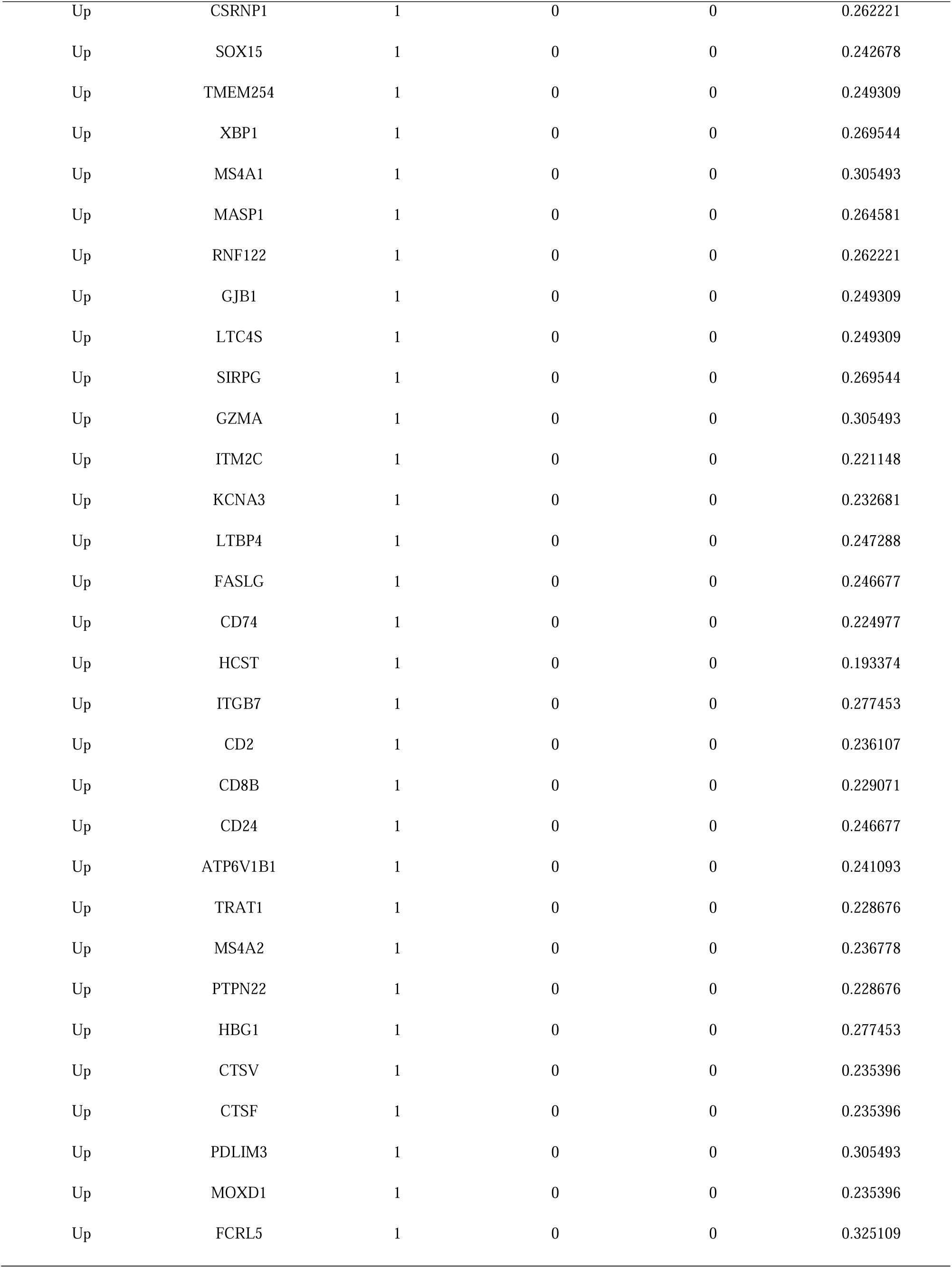
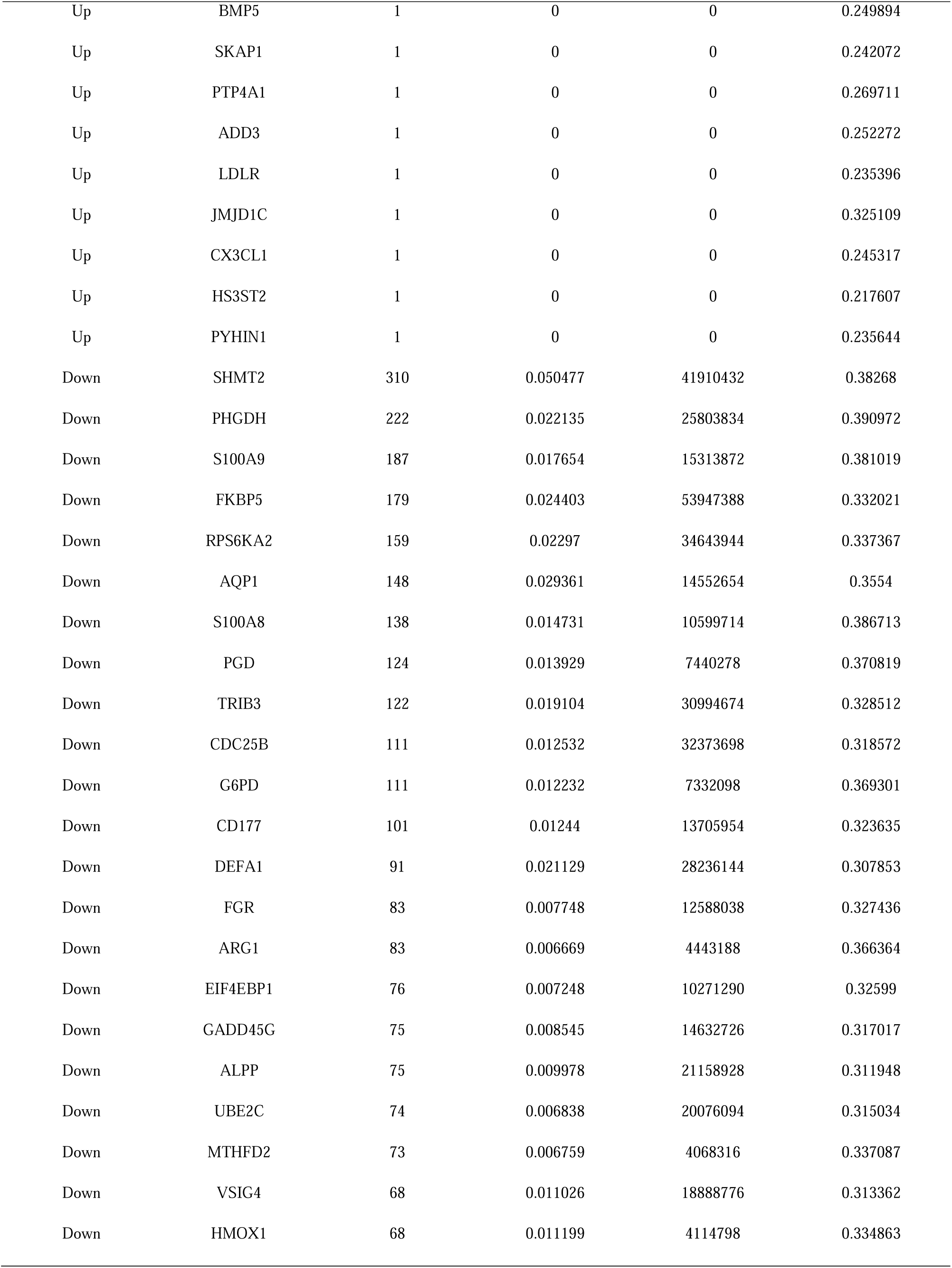
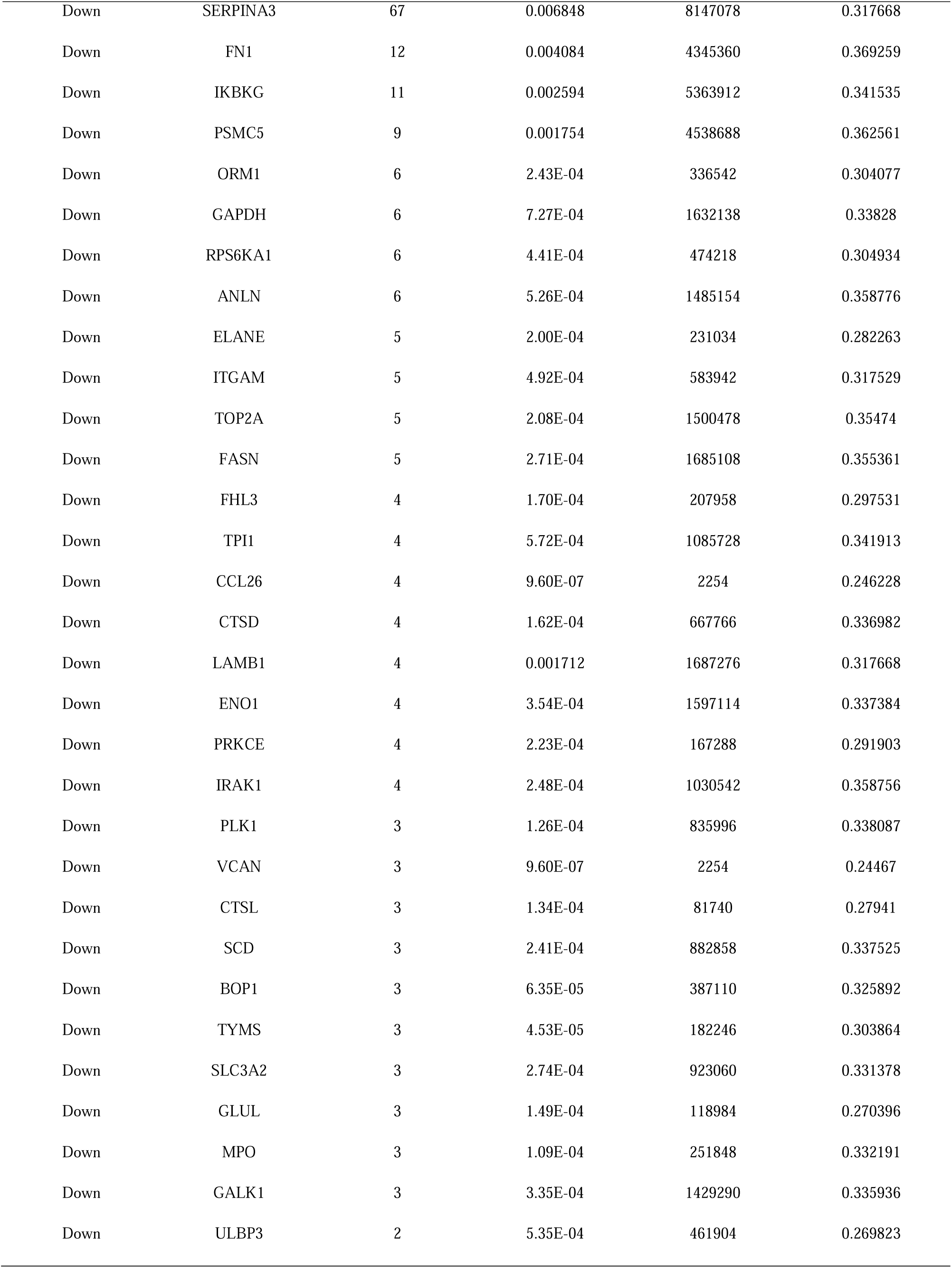
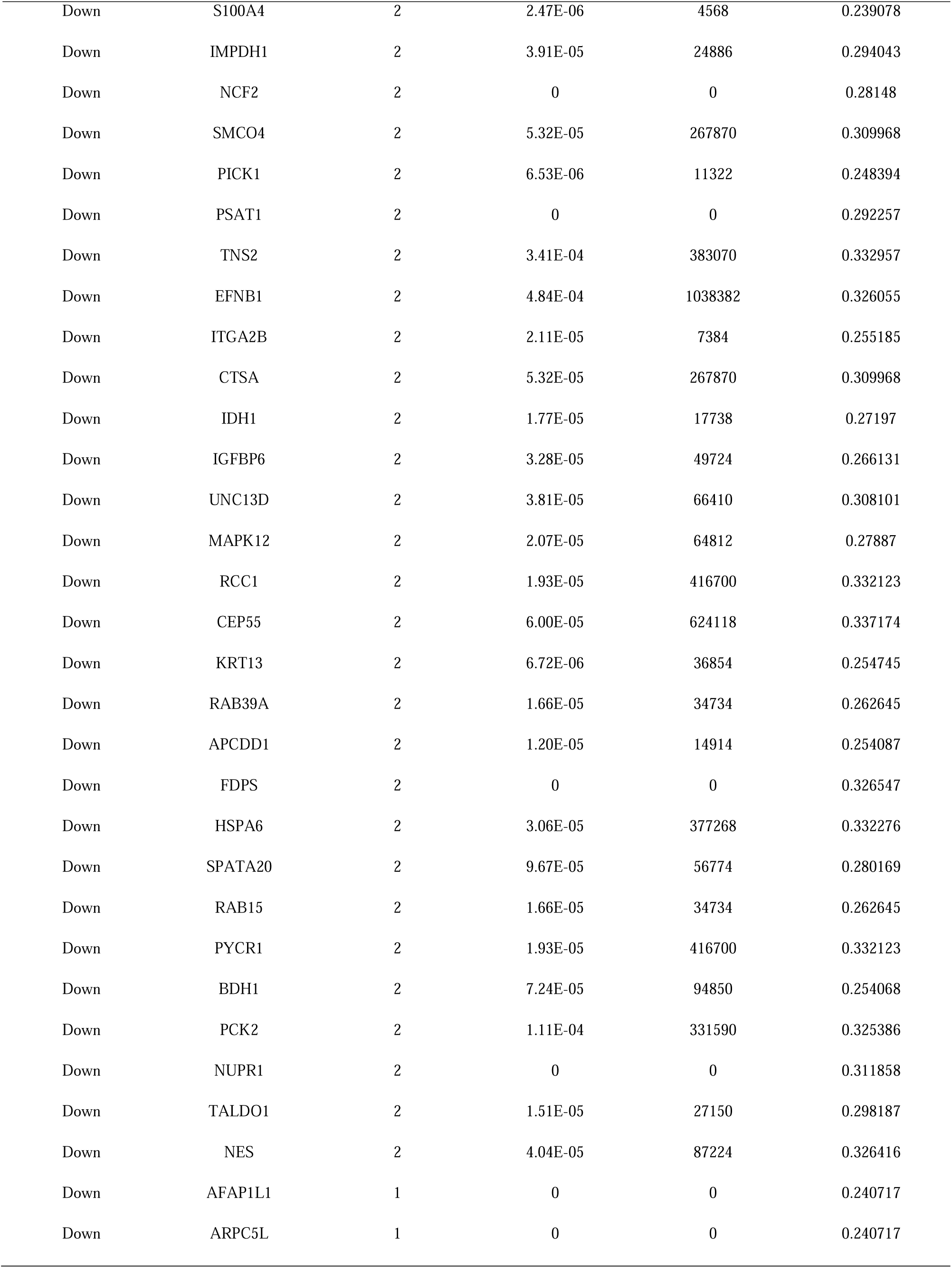
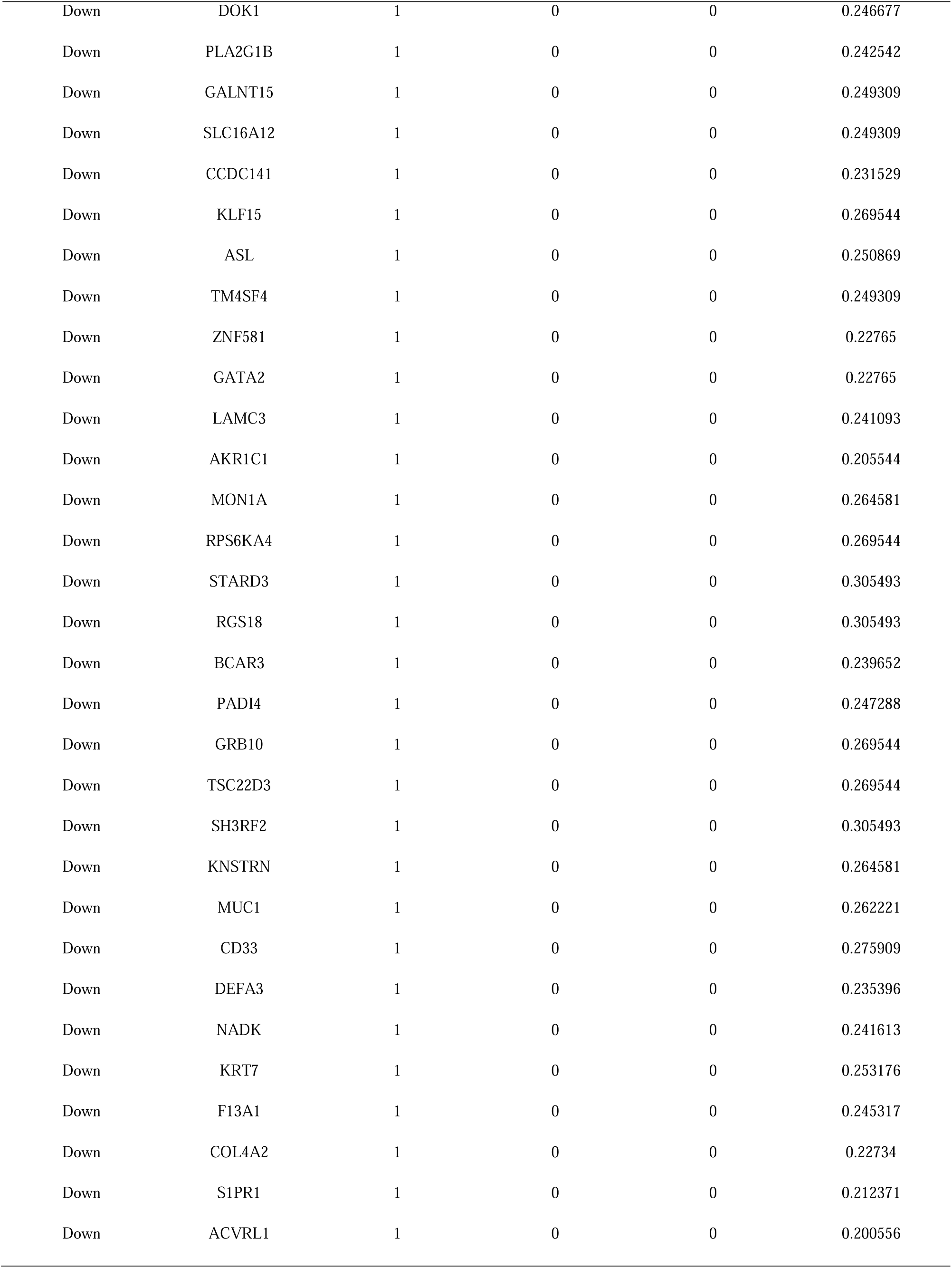
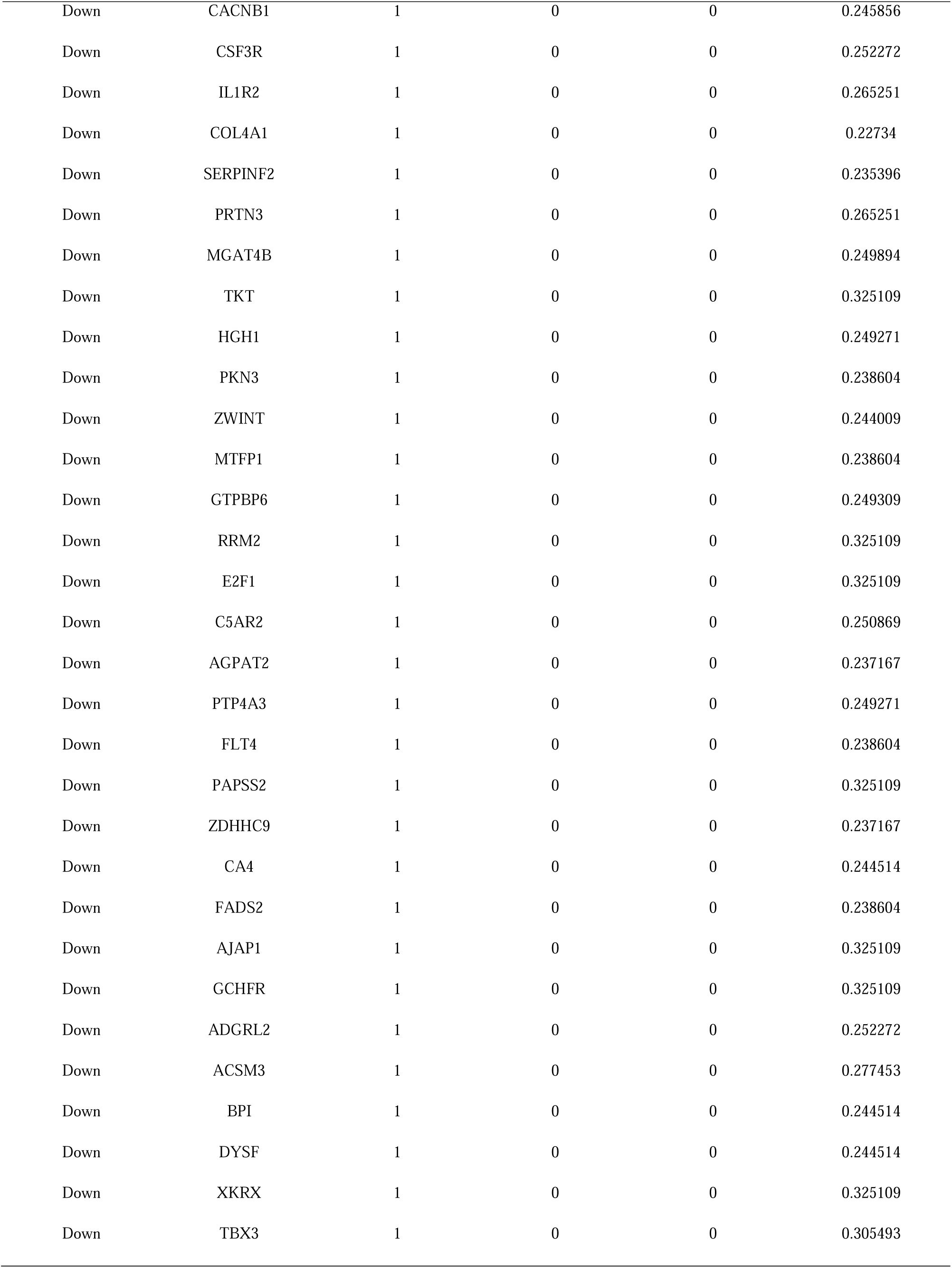
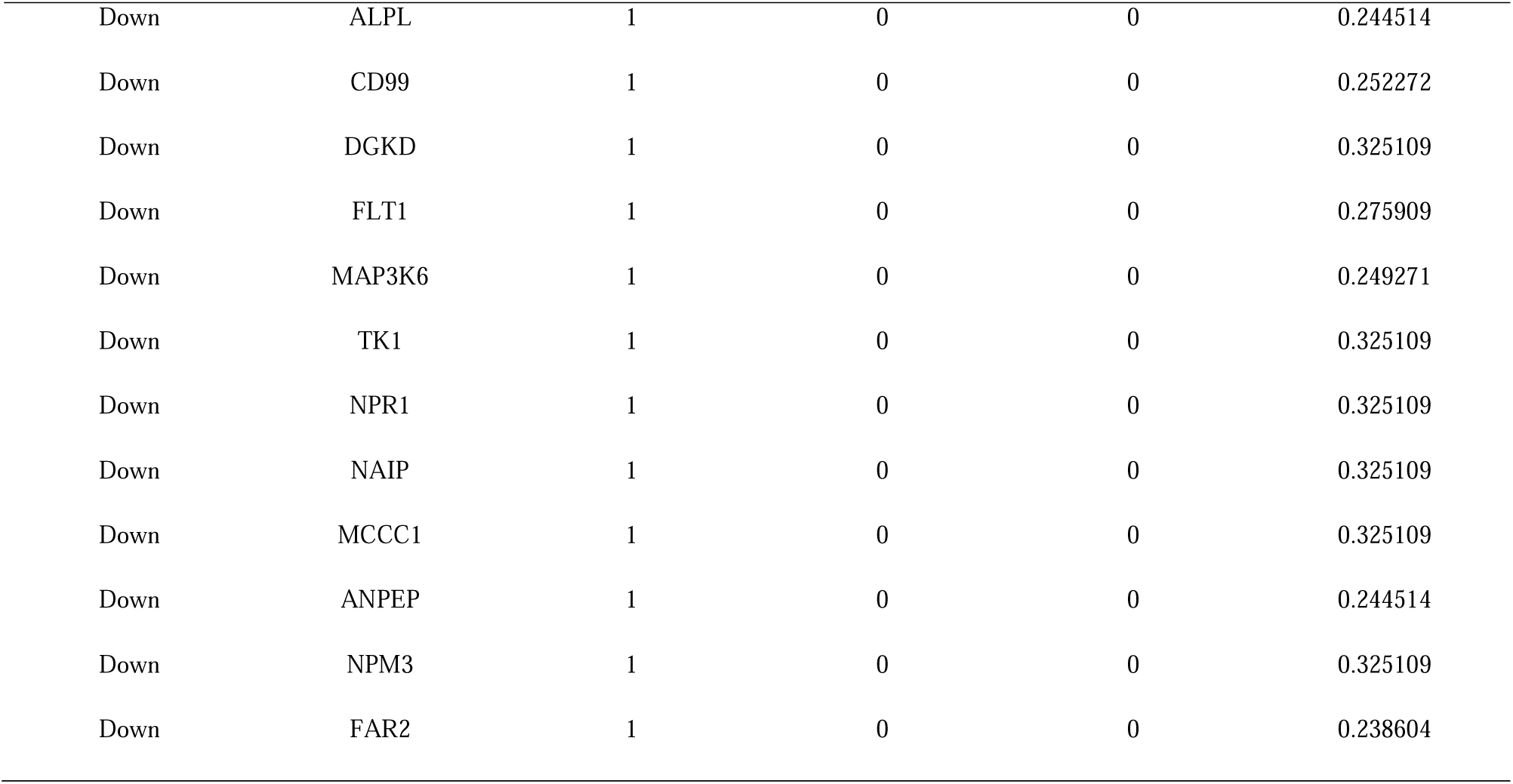
Topology table for up and down regulated genes.

### Construction of the miRNA-hub gene regulatory network

To understand the potential regulation of hub genes, we performed a comprehensive network analysis of miRNA. The miRNA-hub gene regulatory network consists of 2630 (miRNA: 2279; Hub Gene: 351) nodes and 17158 edges (Fig. 5). We found 275 miRNAs (ex: hsa-mir-410-3p) regulating DDX3X, 194 miRNAs (ex: hsa-mir-186-5p) regulating MYC, 138 miRNAs (ex: hsa-mir-1277-5p) regulating MAPK6, 130 miRNAs (ex: hsa-mir-4723-5p) regulating ZFP36, 105 miRNAs (ex: hsa-mir-432-5p) regulating BHLHE40, 116 miRNAs (ex: hsa-mir-539-5p) regulating FKBP5, 96 miRNAs (ex: hsa-mir-1268a) regulating CDC25B, 89 miRNAs (ex: hsa-mir-2861) regulating SHMT2, 80 miRNAs (ex: hsa-mir-4533) regulating TRIB3 and 57 miRNAs (ex: hsa-mir-1229-3p) regulating PHGDH (Table 5).

**Fig. 5.**
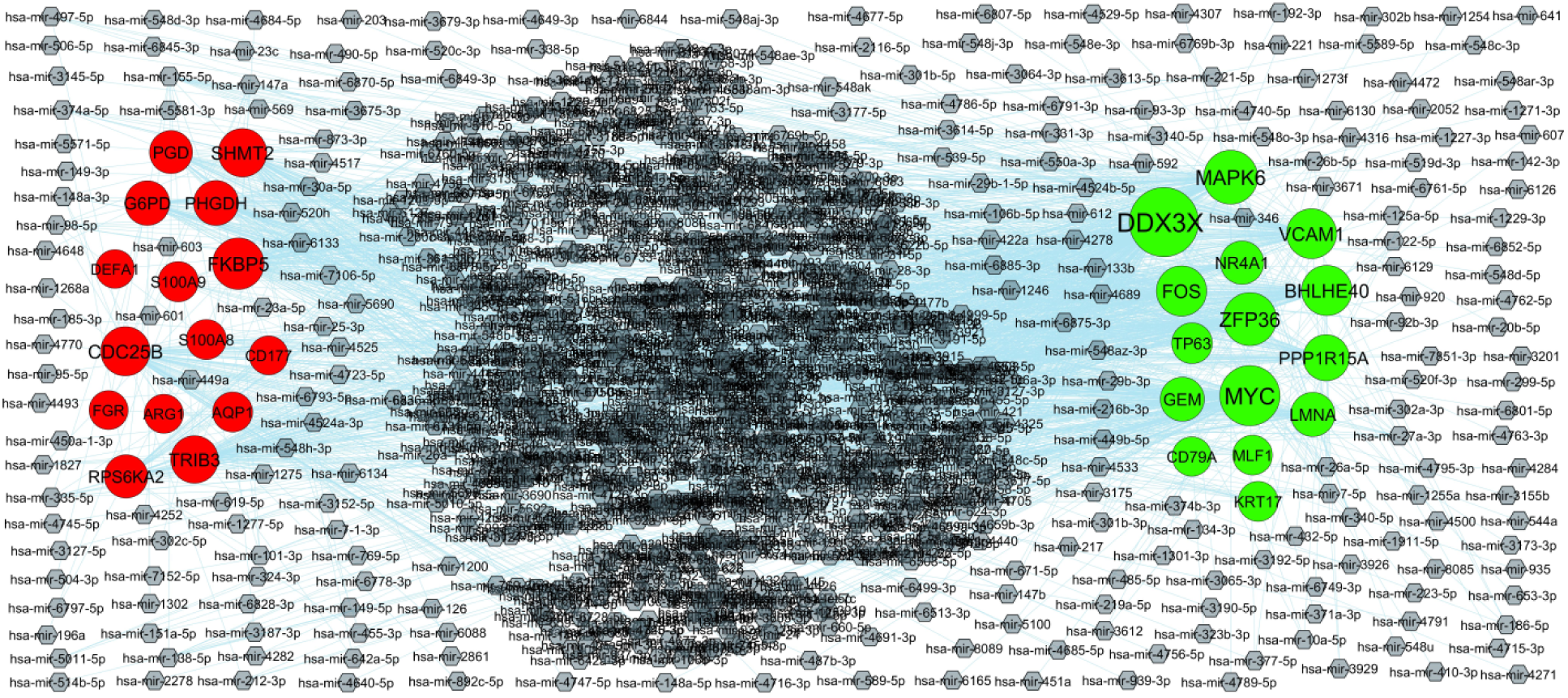
Hub gene - miRNA regulatory network. The ash color diamond nodes represent the key miRNAs; up regulated genes are marked in green; down regulated genes are marked in red.

**Table 5.** MiRNA - hub gene and TF – hub gene topology table.

| Regulation | Hub Genes | Degree | MicroRNA | Regulation | Hub Genes | Degree | TF |
| --- | --- | --- | --- | --- | --- | --- | --- |
| Up | DDX3X | 275 | hsa-mir-410-3p | Up | TP63 | 22 | BRCA1 |
| Up | MYC | 194 | hsa-mir-186-5p | Up | FOS | 14 | FOXA1 |
| Up | MAPK6 | 138 | hsa-mir-1277-5p | Up | DDX3X | 12 | CREB1 |
| Up | ZFP36 | 130 | hsa-mir-4723-5p | Up | KRT17 | 12 | SREBF2 |
| Up | BHLHE40 | 105 | hsa-mir-432-5p | Up | BHLHE40 | 12 | MAX |
| Up | FOS | 105 | hsa-mir-612 | Up | PPP1R15A | 11 | NFKB1 |
| Up | VCAM1 | 102 | hsa-mir-6769b-3p | Up | CD79A | 10 | GATA3 |
| Up | PPP1R15A | 68 | hsa-mir-539-5p | Up | VCAM1 | 9 | SRF |
| Up | LMNA | 50 | hsa-mir-340-5p | Up | GEM | 9 | PAX2 |
| Up | GEM | 49 | hsa-mir-3671 | Up | NR4A1 | 9 | HINFP |
| Up | NR4A1 | 43 | hsa-mir-497-5p | Up | MLF1 | 7 | TEAD1 |
| Up | TP63 | 20 | hsa-mir-1246 | Up | MYC | 7 | FOXC1 |
| Up | KRT17 | 12 | hsa-mir-122-5p | Up | MAPK6 | 6 | TP53 |
| Up | MLF1 | 11 | hsa-mir-422a | Up | ZFP36 | 5 | TFAP2C |
| Up | CD79A | 7 | hsa-mir-520f-3p | Up | LMNA | 4 | YY1 |
| Down | FKBP5 | 116 | hsa-mir-539-5p | Down | FKBP5 | 17 | ESR1 |
| Down | CDC25B | 96 | hsa-mir-1268a | Down | FGR | 16 | PPARG |
| Down | SHMT2 | 89 | hsa-mir-2861 | Down | CDC25B | 13 | NFIC |
| Down | TRIB3 | 80 | hsa-mir-4533 | Down | SHMT2 | 13 | STAT3 |
| Down | PHGDH | 57 | hsa-mir-1229-3p | Down | S100A8 | 13 | NOBOX |
| Down | G6PD | 54 | hsa-mir-1302 | Down | TRIB3 | 10 | ARID3A |
| Down | RPS6KA2 | 43 | hsa-mir-3175 | Down | G6PD | 10 | ELK4 |
| Down | PGD | 40 | hsa-mir-1227-3p | Down | ARG1 | 9 | MYB |
| Down | S100A9 | 15 | hsa-mir-4252 | Down | AQP1 | 9 | RELA |
| Down | AQP1 | 13 | hsa-mir-3187-3p | Down | RPS6KA2 | 9 | KLF5 |
| Down | S100A8 | 12 | hsa-mir-671-5p | Down | PGD | 8 | HOXA5 |
| Down | ARG1 | 4 | hsa-mir-30a-5p | Down | PHGDH | 6 | GATA2 |
| Down | CD177 | 3 | hsa-mir-374b-3p | Down | DEFA1 | 5 | RUNX2 |
| Down | FGR | 3 | hsa-mir-147a | Down | CD177 | 2 | NKX3-2 |
| Down | DEFA1 | 1 | hsa-mir-26b-5p | Down | S100A9 | 2 | USF2 |

### Construction of the TF-hub gene regulatory network

To understand the potential regulation of hub genes, we performed a comprehensive network analysis of TF. The TF-hub gene regulatory network consists of 449 (TF: 96; Hub Gene: 353) nodes and 2990 edges (Fig. 6). We found 22 TFs (ex: BRCA1) regulating TP63, 14 TFs (ex: FOXA1) regulating FOS, 12 TFs (ex: CREB1) regulating DDX3X, 12 TFs (ex: SREBF2) regulating KRT17, 12 TFs (ex: MAX) regulating BHLHE40, 17 TFs (ex: ESR1) regulating FKBP5, 16 TFs (ex: PPARG) regulating FGR, 13 TFs (ex: NFIC) regulating CDC25B, 13 TFs (ex: STAT3) regulating SHMT2 and 13 TFs (ex: NOBOX) regulating S100A8 (Table 5).

**Fig. 6.**
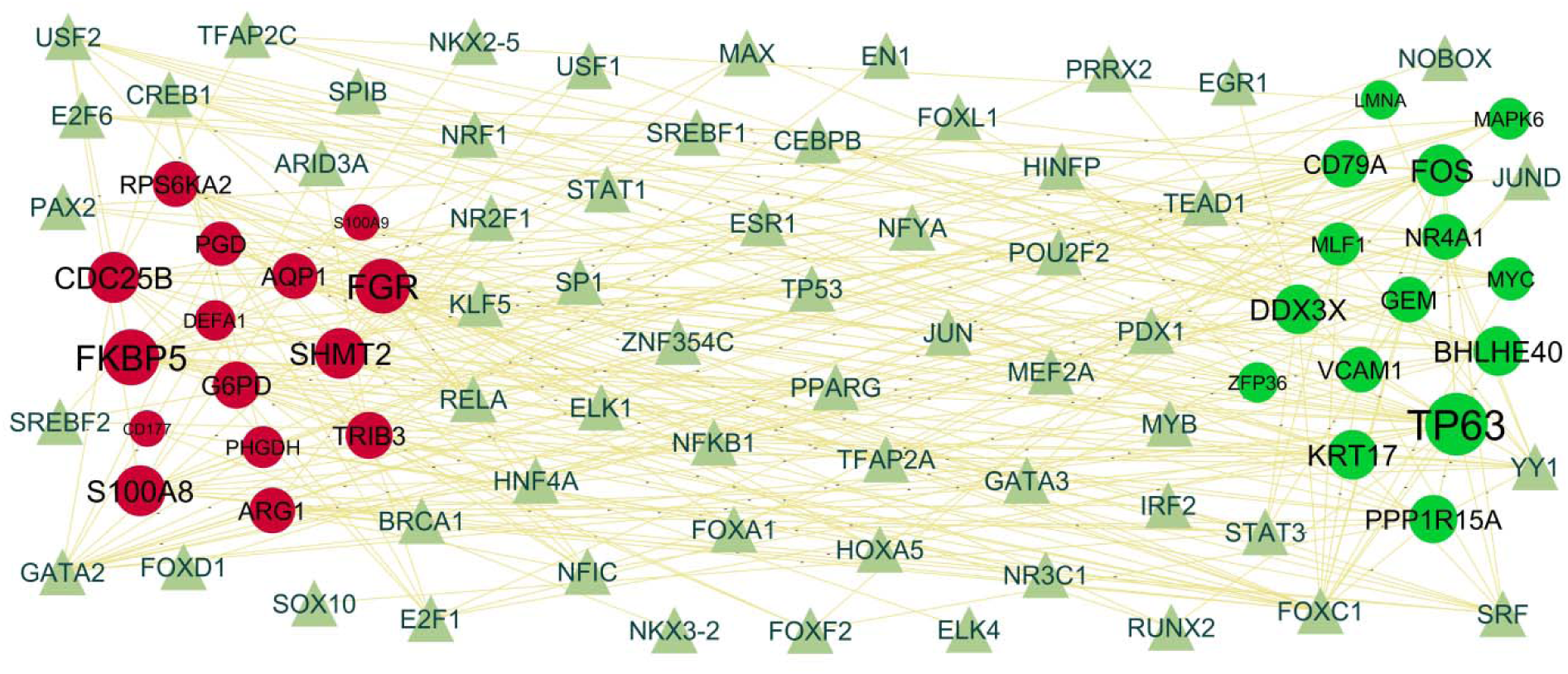
Hub gene - TF regulatory network. The light green color triangle nodes represent the key TFs; up regulated genes are marked in dark green; down regulated genes are marked in dark red.

### Receiver operating characteristic curve (ROC) analysis

ROC curve analysis was performed on the hub genes. The results indicated that all 10 hub gene had good diagnostic value. The AUC value (>□0.8) of hub genes are as follows: MYC (0.889), LMNA (0.925), VCAM1 (0.906), MAPK6 (0.913), DDX3X (0.905), SHMT2 (0.908), PHGDH (0.922), S100A9 (0.918), FKBP5 (0.901) and RPS6KA2 (0.894) (Fig.7).

**Fig. 7.**
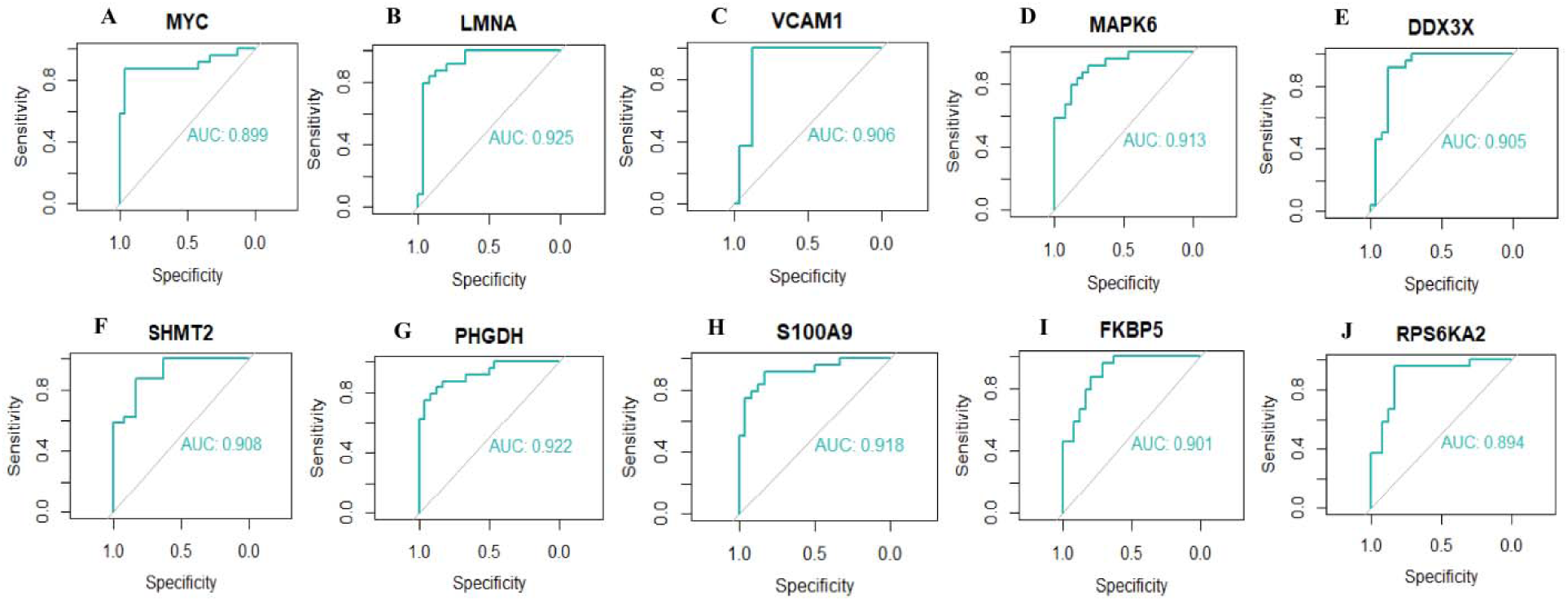
ROC curve analyses of hub genes. A) MYC B) LMNA C) VCAM1 D) MAPK6 E) DDX3X F) SHMT2 G) PHGDH H) S100A9 I) FKBP5 J) RPS6KA2

## Discussion

In recent years, due to the lack of reliable early diagnostic tools and methods, most COPD patients have suffered breathing difficulty, which would require bronchodilator and antiinflammatory treatments. This causes leading to severe physical, mental, and economic burdens to the patients. Previous investigations have shown that the immune microenvironment might play a key role in the occurrence and development of COPD [50–51]. However, the specific targets and therapeutic mechanisms of COPD remain unclear and require further investigation. With the advancement of NGS technology, it has started to be widely applied to find the key genes of diseases. The analysis of NGS data provides a convenient and comprehensive platform to reveal the molecular pathogenesis of COPD initiation and find effective therapeutic target for the treatment of COPD.

In this investigation, we analyzed the COPD expression profile GSE239897 screened from the GEO database. It includes 43 COPD and 39 normal control samples, we found 956 DEGs (including 478 up regulated genes and 478 down regulated genes). HBB (hemoglobin subunit beta) [52], HLA-DQA1 [53], EGR1 [54], ATF3 [55], XIST (X inactive specific transcript) [56], CD163 [57] and S100A8 [58] plays a crucial role in the pathogenesis of lung cancer and considered a promising target for pharmacological based therapies. HBB (hemoglobin subunit beta) [59], HBA1 [60], HLA-DQA1 [61], EGR1 [62], ATF3 [63], XIST (X inactive specific transcript) [64], S100A12 [65], CD163 [66] and S100A8 [67] are closely related to the progression of diabetes mellitus. HBA2 [68], HBA1 [68], EGR1 [69], XIST (X inactive specific transcript) [70], S100A12 [71] and CD163 [72] reported to play a central role in pulmonary hypertension. CHIT1 [73], EGR1 [74], S100A12 [75], CD163 [76] and S100A8 [77] have a critical role in initiating COPD. CHIT1 [78], EGR1 [79], ATF3 [80], S100A12 [81], CD163 [82] and S100A8 [81] were significantly associated with idiopathic pulmonary fibrosis. CHIT1 [83], EGR1 [84], ATF3 [85], XIST (X inactive specific transcript) [86], CD163 [66] and S100A8 [87] level were significantly altered in obesity. Increasing evidence demonstrated that EGR1 [88], RNASE2 [89], CD163 [90] and S100A8 [91], and its protein product has a function in respiratory viral infections. EGR1 [92], ATF3 [93], XIST (X inactive specific transcript) [94], S100A12 [95] and S100A8 [96] were identified as a candidate genes of heart problems. EGR1 [97], ATF3 [63] and XIST (X inactive specific transcript) [98] are involved in the development of osteoporosis. A recent research suggested that EGR1 [99] and S100A8 [100] contributed to the advancement of depression and anxiety. ATF3 [101], S100A12 [102], CD163 [103] and S100A8 [104] could act as diagnosis and prognosis biomarker for airway inflammation. ATF3 [105], S100A12 [106], CD163 [107] and S100A8 [108] played an important role in the progression of pneumonia. All these findings suggested that significant DEGs act as key factors in the pathological process of COPD and deserve more attention.

Next, the DEGs were annotated by performing GO term and pathway enrichment analysis, and we observed that these genes were closely related to signaling pathways. Signaling pathways include immune system [109], signal transduction [110], extracellular matrix organization [111], adaptive immune system [112], neutrophil degranulation [113], innate immune system [114], metabolism [115], diseases of metabolism [116] and Hemostasis [117] were responsible for advancement of COPD. PTGS2 [118], IL6 [119], CSF3 [120], CXCL8 [121], CXCL1 [122], SOCS3 [123], CCL3 [124], CCL5 [125], VCAM1 [126], CXCL13 [127], CXCL5 [128], CXCL12 [129], FGG (fibrinogen gamma chain) [130], IL1B [131], AREG (amphiregulin) [132], SERPINE1 [133], OSM (oncostatin M) [134], CCL11 [135], MFAP4 [136], APLN (apelin) [137], IL1RN [138], IFNG (interferon gamma) [139], CX3CL1 [140], CDKN1A [141], ICAM1 [142], SNAI1 [143], CD34 [144], IL33 [145], FASLG (Fas ligand) [146], FGF7 [147], TNF (tumor necrosis factor) [148], GDF15 [149], CCL22 [150], IL2 [151], CXCR6 [152], KLF5 [153], MCL1 [154], SRF (serum response factor) [155], FABP4 [156], CYP2C19 [157], CXCR4 [158], FOXC1 [159], EPHX1 [160], ANXA1 [161], DUSP6 [162], CPA3 [163], MYH10 [164], ICOS (inducible T cell costimulator) [165], CD1C [166], CD96 [167], S100A9 [168], FKBP5 [169], HMOX1 [170], PRTN3 [171], MARCO (macrophage receptor with collagenous structure) [172], STAB1 [173], SIGLEC9 [174], SLC11A1 [175], BPI (bactericidal permeability increasing protein) [176], PGF (placental growth factor) [177], FASN (fatty acid synthase) [178], S100A4 [179], CD33 [180], PTX3 [181], FLT1 [182], ABCA3 [183], DUOX1 [184], KL (klotho) [185], ALOX5AP [186], FPR2 [187], IL27 [188], FZD4 [189], MPO (myeloperoxidase) [190], CTSD (cathepsin D) [191], MUC1 [192], SIGLEC14 [193], VCAN (versican) [194] and KLK1 [195] have been identified as a target for COPD. PTGS2 [196], NR4A1 [197], IL6 [198], CXCL1 [199], SOCS3 [200], CCL3 [201], CCL5 [202], CCL2 [203], NR4A2 [204], CXCL12 [205], DUSP1 [206], GADD45B [207], APLNR (apelin receptor) [208], IL1B [209], SERPINE1 [210], CCL11 [211], APLN (apelin) [212], HCAR2 [213], IFNG (interferon gamma) [214], CX3CL1 [215], LDLR (low density lipoprotein receptor) [216], TNFAIP3 [217], BMP5 [218], ITK (IL2 inducible T cell kinase) [219], ICAM1 [220], SNCA (synuclein alpha) [221], CD34 [222], HOMER1 [223], IL33 [224], CPE (carboxypeptidase E) [225], KDM6B [226], TNF (tumor necrosis factor) [227], GDF15 [228], CCL22 [229], CCR4 [230], IL2 [231], IRF1 [196], FABP4 [232], CYP2C19 [233], CXCR4 [234], TFPI2 [235], ANXA1 [236], RGS5 [237], TFF3 [238], S100A9 [239], FKBP5 [240], ARG1 [241], RASD1 [242], ADRA1A [243], STAB1 [244], ABCB6 [245], IFNGR1 [246], GPER1 [247], SLC1A3 [248], GLUL (glutamate-ammonia ligase) [249], GATA2 [250], OLFM4 [251], FADS1 [252], KL (klotho) [253], FPR2 [254], FGF22 [255], MPO (myeloperoxidase) [256], MAOA (monoamine oxidase A) [257], BRINP1 [258], CTSD (cathepsin D) [259], BAX (BCL2 associated X, apoptosis regulator) [260], FADS2 [261], EHD3 [262] and PSAT1 [263] are a major mediators of depression and anxiety. PTGS2 [264], NR4A1 [265], EGR3 [266], KLRK1 [267], IL6 [268], CXCL8 [269], CXCL2 [269], SELE (selectin E) [270], MZB1 [271], CD69 [272], ZFP36 [273], CXCL1 [274], SOCS2 [275], NR4A3 [276], TRIB1 [277], SOCS3 [278], CCL3 [279], CCL5 [280], VCAM1 [281], CCL2 [282], BTG2 [283], IER3 [284], CXCL13 [285], ITGBL1 [286], CD24 [287], CXCL5 [288], CXCL12 [289], EYA4 [290], FGG (fibrinogen gamma chain) [291], RND1 [292], DUSP1 [293], CCL19 [294], GADD45B [295], KLK5 [296], CD3E [297], INHBA (inhibin subunit beta A) [298], CXCL6 [299], IL1B [300], AREG (amphiregulin) [301], PIM2 [302], SERPINE1 [303], OSM (oncostatin M) [304], CHRNA1 [305], CCL11 [306], MFAP4 [307], MLF1 [308], CH25H [309], CD27 [310], RHOH (ras homolog family member H) [311], CD5 [312], TP63 [313], LMNA (lamin A/C) [314], CD8A [315], WIF1 [316], KLF4 [317], IL1RN [318], IFNG (interferon gamma) [319], MS4A1 [320], FHL2 [321], CX3CL1 [322], BMP2 [323], EPHA2 [324], ANO1 [325], LDLR (low density lipoprotein receptor) [326], CDKN1A [327], CCR7 [328], GBP1 [329], BMP5 [330], BMP4 [331], CTNNAL1 [332], HTRA3 [333], ICAM1 [334], SNCA (synuclein alpha) [335], MSC (musculin) [336], CLDN1 [337], SNAI1 [338], CD34 [339], DUSP26 [340], IL33 [341], SULF1 [342], HAS2 [343], EYA2 [344], THBS1 [345], FASLG (Fas ligand) [346], CPE (carboxypeptidase E) [347], TRAT1 [348], KDM6B [349], CYGB (cytoglobin) [350], RND3 [351], DERL3 [352], TNF (tumor necrosis factor) [353], GDF15 [354], CCL22 [355], ARHGAP15 [356], CCR4 [357], NFKB2 [358], TESPA1 [359], TRIM52 [360], CD74 [361], CD6 [362], ALDH1A1 [363], IL2 [364], CXCR6 [365], DDX3X[366], KLF5 [367], IRF1 [368], PIM1 [369], PLK3 [370], IRF4 [371], EGLN3 [372], IL2RG [373], MCL1 [374], GSTM2 [375], SRF (serum response factor) [376], INPP4B [377], EDN1 [378], FOSL1 [379], FABP4 [380], CYP2C19 [381], CXCR4 [382], GJA1 [383], KDM6A [384], CTSV (cathepsin V) [385], FOXC1 [386], TFPI2 [387], KLRG1 [388], TIPARP (TCDD inducible poly(ADP-ribose) polymerase) [389], EPHX1 [390], GBP5 [391], DUSP10 [392], CD200R1 [393], DKK2 [394], EPHA3 [395], MDFI (MyoD family inhibitor) [396], ARID5A [397], XBP1 [398], ARRDC3 [399], FUT8 [400], ANXA1 [401], DLC1 [402], MAPK6 [403], DCN (decorin) [404], MYO1E [405], RGS5 [406], DUSP6 [407], CEACAM5 [408], TFF3 [409], AZGP1 [410], LUM (lumican) [411], MUC13 [412], MMP19 [413], PLA2G2D [414], COL10A1 [415], CORIN (corin, serine peptidase) [416], OLR1 [417], TSPAN8 [418], MMP10 [419], ICOS (inducible T cell costimulator) [420], CD1C [421], FBLN2 [422], CTSF (cathepsin F) [423], MASP1 [424], F8 [425], SERPINB2 [426], CD1E [427], CD48 [428], HCST (hematopoietic cell signal transducer) [429], CD96 [430], DDIT4 [431], TTTY15 [432], S100A9 [433], IL1R2 [434], HMOX1 [435], MT1M [436], TMEM100 [437], CLEC4E [438], F2RL3 [439], ARG1 [440], BTNL9 [441], HIF3A [442], AQP1 [443], TRIB3 [444], G6PD [445], FPR1 [446], PGC (progastricsin) [447], PADI4 [448], ADRA1A [449], MARCO (macrophage receptor with collagenous structure) [450], FCN3 [451], LILRB2 [452], AKR1C3 [453], VSIG4 [454], F13A1 [455], EIF4EBP1 [456], ABCB6 [457], SERPINA3 [458], SHMT2 [459], NLRC4 [460], NKD1 [461], ITGA2B [462], PGF (placental growth factor) [463], TYMS (thymidylate synthetase) [464], PTP4A3 [465], ZWINT (ZW10 interacting kinetochore protein) [466], ACVRL1 [467], SDCBP2 [468], ADAMTS9 [469], GPER1 [470], TOP2A [471], FASN (fatty acid synthase) [472], IGFBP6 [473], SPINK1 [474], DYSF (dysferlin) [475], HSPA6 [476], NUPR1 [477], S100A4 [478], CD33 [479], CTSL (cathepsin L) [480], RGS18 [481], MGST1 [482], PDK4 [483], KRT13 [484], AKR1C1 [485], BIRC5 [486], E2F1 [487], KLF15 [488], ENO1 [489], PTX3 [490], IDH1 [491], GATA2 [492], FLT1 [493], OLFM4 [494], ABCA3 [495], SEMA3F [496], KNSTRN (kinetochore localized astrin (SPAG5) binding protein) [497], FADS1 [498], DUOX1 [499], IRAK1 [500], DOK3 [501], KL (klotho) [502], SLC3A2 [503], SLC39A4 [504], TBX3 [505], RAC3 [506], PCK2 [507], PLK1 [508], RASSF4 [509], IL27 [510], STC2 [511], C1R [512], PON2 [513], MCF2L [514], PYCR1 [515], GAPDH (glyceraldehyde- 3-phosphate dehydrogenase) [516], FZD4 [517], SPOCK2 [518], ME1 [519], FGF22 [520], MPO (myeloperoxidase) [521], MAOA (monoamine oxidase A) [522], SCD (stearoyl-CoA desaturase) [523], LPL (lipoprotein lipase) [524], TSPAN14 [525], CYP2A6 [526], PLIN2 [527], S100A14 [528], DOK2 [529], S1PR1 [530], GKN2 [531], SRXN1 [532], AIF1 [533], NOVA2 [534], CTSD (cathepsin D) [535], ABCC3 [536], TLR8 [537], MUC1 [538], ZDHHC9 [539], NCF2 [540], FOXF1 [541], FN1 [542], BAX (BCL2 associated X, apoptosis regulator) [543], FLT4 [544], ARHGAP11A [545], VCAN (versican) [546], CA4 [547], METTL7B [548], TKT (transketolase) [549], EHD2 [550], HTATIP2 [551], TP53I3 [552], PSAT1 [553], UBE2C [554], MTHFD2 [555], CDC25B [556], TK1 [557], BDH1 [558], RRM2 [559], PHGDH (phosphoglycerate dehydrogenase) [560], CHAC2 [561], TDRD9 [562] and PSMC5 [563] expression was significantly altered in lung cancer. PTGS2 [564], IL6 [565], CXCL8 [566], NR4A3 [567], SOCS3 [568], CCL5 [569], CCL2 [570], CXCL13 [571], DUSP5 [572], CXCL12 [573], HBEGF (heparin binding EGF like growth factor) [574], AREG (amphiregulin) [575], APLN (apelin) [576], WIF1 [577], KLF4 [578], IFNG (interferon gamma) [579], CX3CL1 [580], BMP2 [581], ANO1 [582], LDLR (low density lipoprotein receptor) [583], TNFRSF13B [584], CCR7 [585], TNFAIP3 [586], HPGDS (hematopoietic prostaglandin D synthase) [587], ICAM1 [588], CD34 [589], IL33 [590], ASPN (asporin) [591], HAS2 [592], CAMK4 [593], THBS1 [594], FASLG (Fas ligand) [595], PDE4B [596], FGF7 [597], TNF (tumor necrosis factor) [598], GDF15 [149], CD74 [599], HTR2B [600], KLF5 [601], PIM1 [602], SRF (serum response factor) [603], EDN1 [604], CXCR4 [605], KDM6A [606], FOXC1 [607], SELP (selectin P) [608], FUT8 [609], ANXA1 [610], RGS5 [611], ECM2 [612], THBD (thrombomodulin) [613], MMP10 [614], ICOS (inducible T cell costimulator) [615], FBLN2 [616], F8 [617], HMOX1 [618], HIF3A [619], AQP1 [620], TRIB3 [621], G6PD [622], ABCC8 [623], EIF4EBP1 [624], SHMT2 [625], PGF (placental growth factor) [626], ACVRL1 [627], FASN (fatty acid synthase) [628], S100A4 [629], PDK4 [630], KLF15 [631], ENO1 [632], PTX3 [633], GATA2 [634], FLT1 [635], ABCA3 [636], KL (klotho) [637], PLK1 [638], GAPDH (glyceraldehyde-3- phosphate dehydrogenase) [639], MPO (myeloperoxidase) [640], LPL (lipoprotein lipase) [641], TLR8 [642], MUC1 [643], FOXF1 [644], VCAN (versican) [645] and FDPS (farnesyl diphosphate synthase) [646] have been proposed as the most promising biomarkers for pulmonary hypertension. PTGS2 [647], NR4A1 [648], IL6 [649], CXCL8 [650], CD69 [651], CXCL1 [652], SOCS2 [653], SOCS3 [654], CCL3 [655], CCL5 [656], VCAM1 [657], CCL2 [658], CXCL5 [659], DUSP5 [660], CXCL12 [661], KLF10 [662], KLK5 [663], CD3E [664], CD3D [665], IL1B [666], AREG (amphiregulin) [667], SERPINE1 [668], OSM (oncostatin M) [669], CD83 [670], MFAP4 [671], APLN (apelin) [672], RARRES2 [673], CH25H [674], CD27 [675], LMNA (lamin A/C) [676], KLF4 [677], IL1RN [678], IFNG(interferon gamma) [679], FHL2 [680], CX3CL1 [681], GZMA (granzyme A) [682], BMP2 [683], LDLR (low density lipoprotein receptor) [684], CDKN1A [685], RGS16 [686], TNFAIP3 [687], BMP4 [688], ICAM1 [689], SNCA (synuclein alpha) [690], CD34 [691], IL33 [692], FASLG (Fas ligand) [693], CPE (carboxypeptidase E) [694], PDE4B [695], FGF7 [696], ITGB7 [697], RND3 [698], TNF (tumor necrosis factor) [699], CASQ2 [700], GDF15 [701], CCR4 [702], CD74 [703], IL2 [704], PPP1R15B [705], CXCR6 [706], HTR2B [707], DDX3X [708], KLF5 [709], IRF1 [710], PIM1 [711], PLK3 [712], IRF4 [713], MCL1 [714], SRF (serum response factor) [715], SEMA3E [716], EDN1 [717], FOSL1 [718], FABP4 [719], RASGRP1 [720], CYP2C19 [721], CXCR4 [722], TFPI2 [723], EPHX1 [724], SELP (selectin P) [725], LTBP4 [726], ADCYAP1 [727], XBP1 [728], ANXA1 [729], DCN (decorin) [730], RGS5 [731], SIRPG (signal regulatory protein gamma) [732], PTPN22 [733], LUM (lumican) [734], STEAP4 [735], CORIN (corin, serine peptidase) [736], TSPAN8 [737], MMP10 [738], ICOS (inducible T cell costimulator) [739], MASP1 [740], F8 [741], CD48 [742], HLA-DOA [743], DDIT4 [744], S100A9 [745], P2RY1 [746], FKBP5 [747], HMOX1 [748], ITGA10 [749], ARG1 [750], HIF3A [751], AQP1 [752], TRIB3 [753], G6PD [754], ABCC8 [755], SLC11A1 [756], NPR3 [757], NLRC4 [758], HHEX (hematopoietically expressed homeobox) [759], ADAMTS9 [760], FASN (fatty acid synthase) [761], SPINK1 [762], S100A4 [763], CD33 [764], CTSL (cathepsin L) [765], NID1 [766], PDK4 [767], E2F1 [768], KLF15 [769], PTX3 [770], GLUL (glutamate-ammonia ligase) [249], FLT1 [771], OLFM4 [772], FADS1 [773], IRAK1 [774], KL (klotho) [775], ALOX5AP [776], IL15RA [777], EFNB1 [778], PCK2 [779], PLK1 [780], FPR2 [781], IL27 [782], PON2 [783], GAPDH (glyceraldehyde-3-phosphate dehydrogenase) [784], FZD4 [785], GPBAR1 [786], MPO (myeloperoxidase) [787], MAOA (monoamine oxidase A) [788], SCD (stearoyl-CoA desaturase) [789], LPL (lipoprotein lipase) [790], KCNJ11 [791], CYP2A6 [792], COL4A1 [793], AIF1 [794], GRB10 [795], DGKD (diacylglycerol kinase delta) [796], CTSD (cathepsin D) [797], TLR8 [798], MUC1 [799], BAX (BCL2 associated X, apoptosis regulator) [800], DOK1 [801], ECHDC3 [802], VCAN (versican) [803], PCSK2 [804], FADS2 [805], TKT (transketolase) [806], NCF4 [807], HTATIP2 [808], PNMT (phenylethanolamine N-methyltransferase) [809], KLK1 [810], HSD17B14 [811], ACOT1 [812], TK1 [813] and AFMID (arylformamidase) [814] have been identified as high-risk genes in diabetes mellitus. NR4A1 [815], IL6 [816], CXCL8 [817], CD69 [818], CXCL1 [819], SOCS3 [820], CCL3 [821], CCL5 [822], VCAM1 [823], CCL2 [824], CXCL13 [825], CD24 [826], CXCL5 [827], CXCL12 [828], OSM (oncostatin M) [829], CH25H [830], CD27 [831], KLF4 [832], IFNG (interferon gamma) [833], FHL2 [834], CX3CL1 [835], GZMA (granzyme A) [836], CCR7 [837], ICAM1 [838], CD34 [823], IL33 [839], FASLG (Fas ligand) [840], CPE (carboxypeptidase E) [841], KDM6B [842], TNF (tumor necrosis factor) [843], CCR4 [844], CD40LG [845], IL2 [846], IRF1 [847], IL2RG [848], FABP4 [849], CXCR4 [850], SELP (selectin P) [851], XBP1 [852], ANXA1 [853], MAPK6 [854], THBD (thrombomodulin) [855], ICOS (inducible T cell costimulator) [856], F8 [857], DDIT4 [858], S100A9 [108], ARG1 [859], ELANE (elastase, neutrophil expressed) [860], G6PD [861], MARCO (macrophage receptor with collagenous structure) [862], NLRC4 [863], BPI (bactericidal permeability increasing protein) [864], PGF (placental growth factor) [865], S100A4 [866], PTX3 [867], GATA2 [868], ABCA3 [869], FPR2 [870], IL27 [871], GAPDH (glyceraldehyde-3- phosphate dehydrogenase) [872], MPO (myeloperoxidase) [873], FOXF1 [874], VCAN (versican) [875] and FUOM (fucose mutarotase) [876] were revealed to serve an important role in pneumonia. NR4A1 [877], IL6 [878], CSF3 [879], CXCL8 [880], EGR2 [881], ZFP36 [882], CXCL1 [883], SOCS3 [884], CCL3 [885], CCL5 [886], CXCL13 [887], NR4A2 [888], GPR183 [889], CD24 [890], CXCL5 [891], DUSP5 [892], DUSP1 [892], CCL19 [893], CXCL6 [894], IL1B [895], SERPINE1 [896], OSM (oncostatin M) [897], CCL11 [898], CD83 [899], APLN (apelin) [900], RARRES2 [901], CH25H [902], IL1RN [903], IFNG (interferon gamma) [833], FHL2 [904], CX3CL1 [905], EPHA2 [906], ANO1 [907], LDLR (low density lipoprotein receptor) [908], CCR7 [909], SIT1 [910], TNFAIP3 [911], GBP1 [912], HPGDS (hematopoietic prostaglandin D synthase) [913], ICAM1 [914], CLDN1 [915], IL33 [916], THBS1 [917], FASLG (Fas ligand) [918], PDE4B [919], TNF (tumor necrosis factor) [843], GDF15 [920],D74 [921], IL2 [846], CXCR6 [922], DDX3X [923], IRF1 [924], PIM1 [925], ZBP1 [926], PLK3 [927], IRF4 [928], MCL1 [929], CYP2C19 [930], CXCR4 [931], GBP5 [932], SELP (selectin P) [933], DUSP10 [934], XBP1 [852], ANXA1 [935], DUSP6 [936], MMP19 [937], CFD (complement factor D) [938], TSPAN8 [939], MMP10 [940], F8 [857], HLA-DOB [941], CD177 [942], S100A9 [943], FKBP5 [944], HMOX1 [945], ARG1 [946], IL18R1 [947], TRIB3 [948], ELANE (elastase, neutrophil expressed) [949], G6PD [861], MARCO (macrophage receptor with collagenous structure) [950], IL17RB [951], F13A1 [952], NLRC4 [953], BPI (bactericidal permeability increasing protein) [954], GPER1 [955], FASN (fatty acid synthase) [956], HSPA6 [957], CD33 [958], CTSL (cathepsin L) [959], PTX3 [960], OLFM4 [961], ABCA3 [962], IRAK1 [963], DOK3 [964], PLK1 [965], FPR2 [966], IL27 [967], STC2 [968], GAPDH (glyceraldehyde-3-phosphate dehydrogenase) [969], ITGAM (integrin subunit alpha M) [970], MPO (myeloperoxidase) [971], TLR8 [972], MUC1 [973], FLT4 [974], VCAN (versican) [975] and CDC25B [976] have been reported to act as a potential biomarkers for respiratory viral infections treatment. NR4A1 [977], IL6 [978], CXCL8 [979], CD69 [980], ZFP36 [981], CXCL1 [982], SOCS2 [983], SOCS3 [984], CCL5 [985], VCAM1 [986], CCL2 [987], BTG2 [988], CD24 [989], CXCL5 [982], KLF10 [990], FGG (fibrinogen gamma chain) [991], DUSP2 [992], DUSP1 [993], CCL19 [994], APLNR (apelin receptor) [995], IL1B [996], SERPINE1 [997], OSM (oncostatin M) [998], APLN (apelin) [999], CH25H [674], CD27 [675], LMNA (lamin A/C) [676], WIF1 [1000], KLF4 [1001], IL1RN [996], IFNG (interferon gamma) [1002], FHL2 [1003], CX3CL1 [1004], BMP2 [1005], LDLR (low density lipoprotein receptor) [1006], CCR7 [1007], BMP4 [1008], CCDC3 [1009], ICAM1 [1010], CD34 [1011], IL33 [1012], THBS1 [1013], FASLG (Fas ligand) [1014], CPE (carboxypeptidase E) [694], PDE4B [1015], TNF (tumor necrosis factor) [1016], GDF15 [1017], CCL22 [1018], CD74 [1019], ALDH1A1 [1020], IL2 [1021], HTR2B [707], PIM1 [1022], IRF4 [1023], MCL1 [1024], SEMA3E [716], FABP4 [1025], CYP2C19 [1026], OMA1 [1027], CXCR4 [1028], KDM6A [1029], SELP (selectin P) [1030], EPHA3 [1031], XBP1 [1032], ANXA1 [1033], RGS5 [731], DUSP6 [1034], LUM (lumican) [734], STEAP4 [1035], MMP19 [1036], DPT (dermatopontin) [1037], CFD (complement factor D) [1038], CORIN (corin, serine peptidase) [1039], OLR1 [1040], MMP10 [1041], ENPP2 [1042], AIF1L [1043], DDIT4 [1044], S100A9 [1045], FKBP5 [1046], HMOX1 [1047], ARG1 [1048], HIF3A [1049], PRTN3 [1050], TRIB3 [1051],ELANE (elastase, neutrophil expressed) [1052], G6PD [1053], ABCC8 [1054], F13A1 [1055], NPR3 [1056], NLRC4 [1057], PGF (placental growth factor) [1058], TOP2A [1059], FASN (fatty acid synthase) [1060], SPINK1 [1061], NUPR1 [1062], S100A4 [1063], CTSL (cathepsin L) [1064], NID1 [1065], APCDD1 [1066], PDK4 [1067], BIRC5 [1059], E2F1 [1068], KLF15 [1069], PTX3 [1070], GATA2 [1071], OLFM4 [1072], FADS1 [1073], DUOX1 [1074], IRAK1 [1075], ABI3 [1076], ALOX5AP [1077], BCAT2 [1078], PCK2 [779], PLK1 [1079], IL27 [1080], PON2 [1081], ME1 [1082], GPBAR1 [1083], MPO (myeloperoxidase) [1084], MAOA (monoamine oxidase A) [1085], SCD (stearoyl-CoA desaturase) [789], LPL (lipoprotein lipase) [1086], MPST (mercaptopyruvate sulfurtransferase) [1087], CYP2A6 [1088], PLIN2 [1089], LAMC1 [1090], AIF1 [1091], CTSD (cathepsin D) [1092], TLR8 [798], DOK1 [1093], PICK1 [1094], FLT4 [1095], VCAN (versican) [1096], FADS2 [1097], TKT (transketolase) [1098], PNMT (phenylethanolamine N-methyltransferase) [1099], PHGDH (phosphoglycerate dehydrogenase) [1100] and CHDH (choline dehydrogenase) [1101] have been previously reported to be an effective diagnostic biomarkers for obesity. NR4A1 [648], IL6 [1102], CXCL2 [1103], MZB1 [1104], CD69 [92], CXCL1 [1105], SOCS2 [1106], TRIB1 [1107], SOCS3 [1108], CCL3 [655], CCL5 [1109], VCAM1 [1110], NFKBIZ (NFKB inhibitor zeta) [1111], CCL2 [1112], IER3 [1113], NR4A2 [1114], ITGBL1 [1115], DUSP5 [572], CXCL12 [1116], EYA4 [1117], JUND (JunD proto-oncogene, AP-1 transcription factor subunit) [1118], APLNR (apelin receptor) [208], IL1B [1119], AREG (amphiregulin) [1120], OSM (oncostatin M) [669], SMOC2 [1121], MFAP4 [1122], APLN (apelin) [212], RARRES2 [1123], CH25H [1124], TP63 [1125], AADAC (arylacetamide deacetylase) [1126], LMNA (lamin A/C) [1127], KLF4 [1128], IL1RN [1129], GPR174 [1130], IFNG (interferon gamma) [1131], FHL2 [1132], EPHA2 [1133], LDLR (low density lipoprotein receptor) [684], TNFAIP3 [1134], ICAM1 [1110], CD34 [1135], HOMER1 [1136], DUSP26 [1137], IL33 [1138], ASPN (asporin) [1139], FASLG (Fas ligand) [1140], CPE (carboxypeptidase E) [1141], RND3 [698], TNF (tumor necrosis factor) [1142], CASQ2 [700], GDF15 [1143], CCR4 [1144], IL2 [1145], KLF5 [709], PIM1 [711], MCL1 [714], SRF (serum response factor) [715], FABP4 [1146], CYP2C19 [1147], OMA1 [1148], CXCR4 [234], GJA1 [1149], FOXC1 [1150], TFPI2 [723], LTBP4 [1151], TNFSF18 [1152], XBP1 [1153], ACTN2 [1154], ANXA1 [1155], DCN (decorin) [730], RGS5 [1156], DUSP6 [1157], TFF3 [1158], CFD (complement factor D) [1159], THBD (thrombomodulin) [1160], MYH10 [1161], CORIN (corin, serine peptidase) [736], OLR1 [1162], F8 [1163], CPXM2 [1164], SYTL3 [1165], SLAMF7 [1166], DEFA3 [1167], TTTY15 [1168], S100A9 [239], IL1R2 [1169], P2RY1 [1170], HMOX1 [748], CLEC4E [1171], F2RL3 [1072], ARG1 [1173], HIF3A [751], AQP1 [752], ELANE (elastase, neutrophil expressed) [1174], G6PD [1175], VSIG4 [1176], F13A1 [1177], BPI (bactericidal permeability increasing protein) [1178], ADAMTS9 [1179], DYSF (dysferlin) [1180], S100A4 [629], CTSL (cathepsin L) [1181], BIRC5 [1182], KLF15 [769], PTX3 [1183], GATA2 [1071], FLT1 [1184], SEMA3F [1185], FADS1 [773], PRKCE (protein kinase C epsilon) [1186], KL (klotho) [1187], ALOX5AP [1188], TCAP (titin-cap) [1189], BCAT2 [1190], FPR2 [1191], IL27 [1192], PON2 [1193], ALPL (alkaline phosphatase, biomineralization associated) [1194], MPO (myeloperoxidase) [1195], MAOA (monoamine oxidase A) [1196], LPL (lipoprotein lipase) [1197], KCNJ11 [1198], CYP2A6 [1199], COL4A2 [1200], COL4A1 [1201], S1PR1 [1203], GRB10 [795], CTSD (cathepsin D) [1203], FOXF1 [1204], BAX (BCL2 associated X, apoptosis regulator) [1205], VCAN (versican) [1206], TKT (transketolase) [1207], NCF4 [807], LRRC25 [1208], KLK1 [1209], ACOT1 [812], ADAMTS2 [1210] and BDH1 [1211] have been reported to be associated with heart problems. Studies have shown that IL6 [1212], CSF3 [1213], CXCL8 [1214], CXCL2 [1215], CD69 [1216], CXCL1 [1217], SOCS2 [1218], SOCS3 [1219], CCL5 [1220], VCAM1 [1221], CCL2 [1222], CXCL12 [1223], KLF10 [1224], CCL19 [1225], CXCL6 [1226], IL1B [1227], AREG (amphiregulin) [1228], OSM (oncostatin M) [1229], CCL11 [1230], APLN (apelin) [1231], CH25H [1232], KLF4 [1233], IFNG (interferon gamma) [1234], FHL2 [1235], CX3CL1 [1236], EPHA2 [1237], LIF (LIF interleukin 6 family cytokine) [1238], RGS16 [1239], CCR7 [1240], TNFAIP3 [1241], BMP4 [1242], ITK (IL2 inducible T cell kinase) [1243], ICAM1 [1244], CD34 [1245], IL33 [1246], SULF1 [1247], HAS2 [1248], FASLG (Fas ligand) [1249], CPE (carboxypeptidase E) [1250], TNF (tumor necrosis factor) [1251], GDF15 [1252], CCL22 [1253], CCR4 [1254], CD74 [1255], CXCR6 [1256], HTR2B [1257], IRF1 [1258], PIM1 [1259], ZBP1 [926], MCL1 [1260], FABP4 [1261], CXCR4 [1262], KLRG1 [1263], XBP1 [1264], ANXA1 [1265], RGS5 [1266], MMP19 [1267], THBD (thrombomodulin) [1268], MYH10 [164], CD48 [1269], S100A9 [1270], ARG1 [1271], AQP1 [1272], MARCO (macrophage receptor with collagenous structure) [1273], FCN3 [1274], IL17RB [1275], SIGLEC9 [1276], IGFBP6 [473], S100A4 [1277], PTX3 [1278], ABCA3 [1279], DUOX1 [1074], PRKCE (protein kinase C epsilon) [1280], KL (klotho) [1281], SLC3A2 [1282], FPR2 [1283], IL27 [1284], MPO (myeloperoxidase) [1285], CCL26 [1286], CTSD (cathepsin D) [1287], MUC1 [1288], FOXF1 [1289], CTSE (cathepsin E) [1290] and VCAN (versican) [975] are associated with airway inflammation. Recent studies have indicated that IL6 [1291], CXCL8 [1292], CXCL2 [1293], MZB1 [1294], CD69 [1295], ZFP36 [1296], CCL3 [1297], CCL5 [1298], VCAM1 [1299], CCL2 [1300], CXCL13 [1301], ITGBL1 [1302], CXCL12 [1303], CXCL6 [1304], IL1B [1305], AREG (amphiregulin) [1306], SERPINE1 [1307], OSM (oncostatin M) [1308], CCL11 [1298], SMOC2 [1309], MFAP4 [1310], APLN (apelin) [1231], PPP1R15A [1311], TP63 [1312], LMNA (lamin A/C) [1313], WIF1 [1314], KLF4 [1315], IL1RN [1316], IFNG (interferon gamma) [1317], FHL2 [1318], CX3CL1 [1319], EPHA2 [1320], ANO1 [1321], LDLR (low density lipoprotein receptor) [1322], CCR7 [1323], BMP4 [1324], HTRA3 [1325], ICAM1 [1326], SNAI1 [1327], CD34 [1328], IL33 [1329], LTBP2 [1330], HAS2 [1331], FASLG (Fas ligand) [1332], PDE4B [1333], KDM6B [1334], RND3 [1335], TNF (tumor necrosis factor) [1336], GDF15 [1337], CCL22 [1338], CCR4 [1339], HTR2B [1340], IRF1 [1341], PIM1 [1342], IRF4 [1343], BMP3 [1344], SRF (serum response factor) [1345], EDN1 [1346], FABP4 [1347], CXCR4 [1348], WNT10A [1349], DUSP10 [1350], NREP (neuronal regeneration related protein) [1351], ANXA1 [1352], CPA3 [163], MMP19 [1353], MMP10 [1354], ICOS (inducible T cell costimulator) [1355], CD1C [1356], S100A9 [81], HMOX1 [1357], AQP1 [1358], IL18R1 [1359], TRIB3 [1360], FPR1 [1361], MARCO (macrophage receptor with collagenous structure) [1362], PGF (placental growth factor) [1363], FASN (fatty acid synthase) [1364], S100A4 [1365], CTSL (cathepsin L) [1366], E2F1 [1367], PTX3 [1368], GATA2 [634], FLT1 [1369], ABCA3 [636], DUOX1 [1370], IRAK1 [1371], KL (klotho) [1372], FPR2 [1373], IL27 [1374], MPO (myeloperoxidase) [1375], S1PR1 [1376], AIF1 [1377], MUC1 [1378], FOXF1 [1379], BAX (BCL2 associated X, apoptosis regulator) [1380], PSAT1 [1381] and PHGDH (phosphoglycerate dehydrogenase) [1382] are involved in the development of idiopathic pulmonary fibrosis. IL6 [1383], CSF3 [1384], SOCS3 [1385], CCL3 [1386], CCL5 [1387], VCAM1 [1388], CCL2 [1387], CXCL12 [1389], KLF10 [1390], FRZB (frizzled related protein) [1391], DUSP1 [1392], IL1B [1393], CCL11 [1394], LMNA (lamin A/C) [1395], WIF1 [1396], IL1RN [1397], IFNG (interferon gamma) [1398], CX3CL1 [1399], BMP2 [1400], CDKN1A [1401], BMP4 [1402], ICAM1 [1403], CD34 [1404], IL33 [1405], FASLG (Fas ligand) [1406], KDM6B [1407], TNF (tumor necrosis factor) [1408], GDF15 [1409], IL2 [1410], INPP4B [1411], FABP4 [1412], CYP2C19 [1413], CXCR4 [1414], DKK2 [1415], PLCB4 [1416], ANXA1 [1417], DUSP6 [1418], CORIN (corin, serine peptidase) [1419], F8 [1420], DDIT4 [1421], IL1R2 [1422], S100A4 [1417], CTSL (cathepsin L) [1423], PTX3 [1424], EFNB1 [1425], MCF2L [1426], FZD4 [1427], CCL26 [1428], PHGDH (phosphoglycerate dehydrogenase) [1429] and FDPS (farnesyl diphosphate synthase) [1430] have been identified as predictive biomarkers of osteoporosis. IL6 [1431], SOCS2 [1432], SOCS3 [1432], DUSP1 [1433], MLF1 [1434], EPHA2 [1435], CD34 [1436], IL33 [1437], GDF15 [1438], IL2 [1439], CD177 [1440], G6PD [1441], NFE2 [1442], PTX3 [1443], IDH1 [1444], MPO (myeloperoxidase) [1445] and KLK1 [1446] might be associated with the pathogenesis of polycythemia. The abnormal expression of IL33 [1447] and TNF (tumor necrosis factor) [1448] contributes to the pneumothorax. The above results suggest that enriched genes maight influence the progression of COPD.

Using the DEGs genes, a PPI network was constructed and significant modules were isolated on understanding the molecular pathogenesis of COPD, and predicting prognostic and diagnostic biomarkers as well as therapeutic targets. Here, we identified hub genes based on the topological properties (i.e., degree, betweenness, stress and closeness), which can be key biomarkers in COPD and linked with numerous molecular mechanisms. The top hub genes indicate most risk factors for the COPD. Research has shown that LMNA (lamin A/C) [314], VCAM1 [281], MAPK6 [403], DDX3X [366], SHMT2 [459], PHGDH (phosphoglycerate dehydrogenase) [560], S100A9 [433], SNCA (synuclein alpha) [335], TP63 [313], FOSL2 [1449], FHL2 [321], ATF3 [55], FOSL1 [379], DYSF (dysferlin) [475], CA4 [547], S100A8 [58] and MPO (myeloperoxidase) [521] plays an important role in the pathogenesis of lung cancer. LMNA (lamin A/C) [676], VCAM1 [657], DDX3X [708], S100A9 [745], FKBP5 [747], SNCA (synuclein alpha) [690], FOSL2 [1450], FHL2 [680], ATF3 [63], FOSL1 [718], S100A8 [67] and MPO (myeloperoxidase) [787] altered expression is associated with poor prognosis of diabetes mellitus. LMNA (lamin A/C) [676], VCAM1 [986], PHGDH (phosphoglycerate dehydrogenase) [1100], S100A9 [1045], FKBP5 [1046], FHL2 [1003], ATF3 [85] and MPO (myeloperoxidase) [1084] plays an important role in the development of obesity. LMNA (lamin A/C) [1127], VCAM1 [1110], S100A9 [239], TP63 [1125], JUND (JunD proto-oncogene, AP-1 transcription factor subunit) [1118], FHL2 [1132], ATF3 [93], BPI (bactericidal permeability increasing protein) [1178], DYSF (dysferlin) [1180], MPO (myeloperoxidase) [1195] and ALPL (alkaline phosphatase, biomineralization associated) [1194] were reported to be associated with the prognosis of heart problems. LMNA (lamin A/C) [1313], VCAM1 [1299], PHGDH (phosphoglycerate dehydrogenase) [1382], S100A9 [81], TP63 [1312], FHL2 [1318], ATF3 [80] and MPO (myeloperoxidase) [1375] have been identified as a key genes in idiopathic pulmonary fibrosis. LMNA (lamin A/C) [1395], VCAM1 [1388], PHGDH (phosphoglycerate dehydrogenase) [1429] and ATF3 [63] might be a potential targets for osteoporosis. VCAM1 [126], S100A9 [168], FKBP5 [169], BPI (bactericidal permeability increasing protein) [176] and MPO (myeloperoxidase) [190] are mainly associated with COPD. VCAM1 [823], MAPK6 [854], S100A9 [108], FHL2 [834], ATF3 [105], BPI (bactericidal permeability increasing protein) [864] and MPO (myeloperoxidase) [873] have a significant prognostic potential in pneumonia. VCAM1 [1221], S100A9 [1270], FHL2 [1235], ATF3 [101] and MPO (myeloperoxidase) [1285] are molecular markers for the diagnosis and prognosis of airway inflammation. DDX3X [923], S100A9 [943], FKBP5 [944], FHL2 [904], ITGAM (integrin subunit alpha M) [970], BPI (bactericidal permeability increasing protein) [954], MPO (myeloperoxidase) [971] and CD177 [942] have a significant prognostic potential in respiratory viral infections. Research reported that increased level of SHMT2 [625] and MPO (myeloperoxidase) [640] are related to pulmonary hypertension, as a novel target for pulmonary hypertension diagnosis and treatment. A recent study found that the S100A9 [239], FKBP5 [240], SNCA (synuclein alpha) [221] and MPO (myeloperoxidase) [256] are crucial for the progression of depression and anxiety. Regulation of MPO (myeloperoxidase) [1445] and CD177 [1440] levels might be a novel treatment option against polycythemia. The results of this investigation suggest that novel biomarkers include MYC (MYC proto-oncogene, bHLH transcription factor), RPS6KA2, SH2D1A, JUNB (JunB proto-oncogene, AP-1 transcription factor subunit), FOS (Fos proto-oncogene, AP-1 transcription factor subunit), RAB39A and RAB15 might play a key role in the pathogenesis of COPD. This investigation has been proved to be a useful approach to identify novel biomarkers in other diseases.

A miRNA-hub gene regulatory network and TF-hub gene regulatory network were constructed for the identified hub genes, miRNAs and TFs were defined by the degree rank. DDX3X [366], MAPK6 [403], ZFP36 [273], BHLHE40 [1451], TP63 [313], KRT17 [1452], CDC25B [556], SHMT2 [459], TRIB3 [444], PHGDH (phosphoglycerate dehydrogenase) [560], S100A8 [58], hsa-mir-410-3p [1453], hsa-mir-186-5p [1454], hsa-mir-432-5p [1455], BRCA1 [1456], FOXA1 [1457], CREB1 [1458], MAX (MYC associated factor X) [1459], ESR1 [1460], NFIC (nuclear factor I C) [1461] and STAT3 [1462] expression might be regarded as an indicator of susceptibility to lung cancer. DDX3X [708], BHLHE40 [1463], FKBP5 [747], TRIB3 [753], S100A8 [67], FOXA1 [1464], CREB1 [1465], SREBF2 [1466], ESR1 [1467] and STAT3 [1468] are a potential markers for the detection and prognosis of diabetes mellitus. A previous study reported that DDX3X [923], ZFP36 [882], FKBP5 [944], CDC25B [976], TRIB3 [948], hsa-mir-1277-5p [1469], MAX (MYC associated factor X) [1470] and STAT3 [1471] are altered expressed in respiratory viral infections. MAPK6 [854], BRCA1 [1472] and STAT3 [1473] are found to be associated with pneumonia. ZFP36 [981], FKBP5 [1046], TRIB3 [1051], PHGDH (phosphoglycerate dehydrogenase) [1100], BRCA1 [1474], SREBF2 [1475], ESR1 [1476] and STAT3 [1477] plays a vital role in the development of obesity. ZFP36 [1296], TP63 [1312], KRT17 [1478], TRIB3 [1360], PHGDH (phosphoglycerate dehydrogenase) [1382], FOXA1 [1479], CREB1 [1480], SREBF2 [1481] and STAT3 [1482] have been observed to be expressed in idiopathic pulmonary fibrosis patients. TP63 [1125], hsa-mir-410-3p [1483], hsa-mir-432-5p [1484], BRCA1 [1485], ESR1 [1486], NFIC (nuclear factor I C) [1487] and STAT3 [1488] might serve as therapeutic targets for heart problems. The expression levels of FKBP5 [169] and STAT3 [1489] have been proved to be altered in COPD patients. Recent studies have shown that altered expression of FKBP5 [240], BRCA1 [1490], CREB1 [1491], MAX (MYC associated factor X) [1492], ESR1 [1493] and STAT3 [1494] might be associated with the progression of depression and anxiety. Studies have found that SHMT2 [625], TRIB3 [621], BRCA1 [1495] and STAT3 [1496] expression was associated with pulmonary hypertension. PHGDH (phosphoglycerate dehydrogenase) [1429], BRCA1 [1497], FOXA1 [1498], CREB1 [1499], ESR1 [1500] and STAT3 [1501] might be a prognostic biomarker and potential therapeutic target for patients with osteoporosis. BRCA1 [1502], ESR1 [1503] and STAT3 [1504] have been linked to the start of airway inflammation. STAT3 [1505] is implicated in polycythemia. MYC (MYC proto-oncogene, bHLH transcription factor), FOS (Fos proto-oncogene, AP-1 transcription factor subunit), FGR (FGR proto-oncogene, Src family tyrosine kinase), hsa-mir-4723-5p, hsa-mir-539-5p, hsa-mir-1268a, hsa-mir-2861, hsa-mir-4533, hsa-mir-1229-3p, PPARG (peroxisome proliferator activated receptor gamma) and NOBOX (NOBOX oogenesis homeobox) migt be considered to be a novel biomarkers for COPD. These findings suggest that the hub genes, miRNAs and TFs can serve as candidate diagnostic biomarkers or drug targets for the clinical treatment of COPD.

In conclusion, we filtrated a total of 956 DEGs from GEO NGS dataset and further validated 10 hub genes (MYC, LMNA, VCAM1, MAPK6, DDX3X, SHMT2, PHGDH, S100A9, FKBP5 and RPS6KA2). The 10 hub genes were might associated with the prognosis of COPD. The GO functional terms and pathways identified in the investigation migt contribute to understand the molecular mechanisms of COPD. Our findings might provide novel therapeutic targets for COPD patients.

## Acknowledgement

I thanks very much to John McDonough, Yale School of Medicine, New Haven, CT, USA, the author who deposited their NGS dataset GSE239897, into the public GEO database.

## Conflict of interest

The authors declare that they have no conflict of interest.

## Ethical approval

This article does not contain any studies with human participants or animals performed by any of the authors.

## Informed consent

No informed consent because this study does not contain human or animals participants.

## Availability of data and materials

The datasets supporting the conclusions of this article are available in the GEO (Gene Expression Omnibus) (https://www.ncbi.nlm.nih.gov/geo/) repository. [(GSE239897) https://www.ncbi.nlm.nih.gov/geo/query/acc.cgi?acc=GSE239897]

## Consent for publication

Not applicable.

## Competing interests

The authors declare that they have no competing interests.

## Author Contributions

B. V. - Writing original draft, and review and editing

C. V. - Software and investigation

## Notes

### Competing Interest Statement

The authors have declared no competing interest.

## References

1. Kahnert K, Jörres RA, Behr J, Welte T. The Diagnosis and Treatment of COPD and Its Comorbidities. Dtsch Arztebl Int. 2023;120(25):434–444. doi:10.3238/arztebl.m2023.027

2. Bagdonas E, Raudoniute J, Bruzauskaite I, Aldonyte R. Novel aspects of pathogenesis and regeneration mechanisms in COPD. Int J Chron Obstruct Pulmon Dis. 2015;10:995–1013. doi:10.2147/COPD.S82518

3. Festic E, Scanlon PD. Incident pneumonia and mortality in patients with chronic obstructive pulmonary disease. A double effect of inhaled corticosteroids?. Am J Respir Crit Care Med. 2015;191(2):141–148. doi:10.1164/rccm.201409-1654PP

4. Zwaans WA, Mallia P, van Winden ME, Rohde GG. The relevance of respiratory viral infections in the exacerbations of chronic obstructive pulmonary disease—a systematic review. J Clin Virol. 2014;61(2):181–188. doi:10.1016/j.jcv.2014.06.025

5. Hobbs BD, Foreman MG, Bowler R, Jacobson F, Make BJ, Castaldi PJ, San José Estépar R, Silverman EK, Hersh CP; COPD Gene Investigators. Pneumothorax risk factors in smokers with and without chronic obstructive pulmonary disease. Ann Am Thorac Soc. 2014;11(9):1387-1394. doi:10.1513/AnnalsATS.201405-224OC

6. Pellicori P, Cleland JGF, Clark AL. Chronic Obstructive Pulmonary Disease and Heart Failure: A Breathless Conspiracy. Cardiol Clin. 2022;40(2):171–182. doi:10.1016/j.ccl.2021.12.005

7. Chen YW, Ramsook AH, Coxson HO, Bon J, Reid WD. Prevalence and Risk Factors for Osteoporosis in Individuals With COPD: A Systematic Review and Meta-analysis. Chest. 2019;156(6):1092–1110. doi:10.1016/j.chest.2019.06.036

8. Panagioti M, Scott C, Blakemore A, Coventry PA. Overview of the prevalence, impact, and management of depression and anxiety in chronic obstructive pulmonary disease. Int J Chron Obstruct Pulmon Dis. 2014;9:1289–1306. doi:10.2147/COPD.S72073

9. Eapen MS, Hansbro PM, Larsson-Callerfelt AK, Jolly MK, Myers S, Sharma P, Jones B, Rahman MA, Markos J, Chia C, et al. Chronic Obstructive Pulmonary Disease and Lung Cancer: Underlying Pathophysiology and New Therapeutic Modalities. Drugs. 2018;78(16):1717–1740. doi:10.1007/s40265-018-1001-8

10. Piccari L, Aguilar-Colindres R, Rodríguez-Chiaradía DA. Pulmonary hypertension in interstitial lung disease and in chronic obstructive pulmonary disease: different entities?. Curr Opin Pulm Med. 2023;29(5):370–379. doi:10.1097/MCP.0000000000000984

11. Zhang J, DeMeo DL, Silverman EK, Make BJ, Wade RC, Wells JM, Cho MH, Hobbs BD. Secondary polycythemia in chronic obstructive pulmonary disease: prevalence and risk factors. BMC Pulm Med. 2021;21(1):235. doi:10.1186/s12890-021-01585-5

12. Siddhuraj P, Jönsson J, Alyamani M, Prabhala P, Magnusson M, Lindstedt S, Erjefält JS. Dynamically upregulated mast cell CPA3 patterns in chronic obstructive pulmonary disease and idiopathic pulmonary fibrosis. Front Immunol. 2022;13:924244. doi:10.3389/fimmu.2022.924244

13. Goto T, Hirayama A, Faridi MK, Camargo CA Jr, Hasegawa K. Obesity and Severity of Acute Exacerbation of Chronic Obstructive Pulmonary Disease. Ann Am Thorac Soc. 2018;15(2):184–191. doi:10.1513/AnnalsATS.201706-485OC

14. Hsu IL, Lu CL, Li CC, Tsai SH, Chen CZ, Hu SC, Li CY. Population-based cohort study suggesting a significantly increased risk of developing chronic obstructive pulmonary disease in people with type 2 diabetes mellitus. Diabetes Res Clin Pract. 2018;138:66–74. doi:10.1016/j.diabres.2018.01.037

15. Steiling K, Lenburg ME, Spira A. Airway gene expression in chronic obstructive pulmonary disease. Proc Am Thorac Soc. 2009;6(8):697–700. doi:10.1513/pats.200907-076DP

16. Hong Y, Lim MN, Kim WJ, Rhee CK, Yoo KH, Lee JH, Yoon HI, Kim TH, Lee JH, Lim SY, et al. Influence of environmental exposures on patients with chronic obstructive pulmonary disease in Korea. Tuberc Respir Dis (Seoul). 2014;76(5):226–232. doi:10.4046/trd.2014.76.5.226

17. Huang X, Li Y, Guo X, Zhu Z, Kong X, Yu F, Wang Q. Identification of differentially expressed genes and signaling pathways in chronic obstructive pulmonary disease via bioinformatic analysis. FEBS Open Bio. 2019;9(11):1880–1899. doi:10.1002/2211-5463.12719

18. Barnes PJ. Future treatments for chronic obstructive pulmonary disease and its comorbidities. Proc Am Thorac Soc. 2008;5(8):857–864. doi:10.1513/pats.200807-069TH

19. Ganekal P, Vastrad B, Kavatagimath S, Vastrad C, Kotrashetti S. Bioinformatics and Next-Generation Data Analysis for Identification of Genes and Molecular Pathways Involved in Subjects with Diabetes and Obesity. Medicina (Kaunas). 2023;59(2):309. doi:10.3390/medicina59020309

20. Alur V, Raju V, Vastrad B, Vastrad C, Kavatagimath S, Kotturshetti S. Bioinformatics Analysis of Next Generation Sequencing Data Identifies Molecular Biomarkers Associated With Type 2 Diabetes Mellitus. Clin Med Insights Endocrinol Diabetes. 2023;16:11795514231155635. doi:10.1177/11795514231155635

21. Lin S, Liu C, Sun J, Guan Y. RNA-Sequencing and Bioinformatics Analysis of Exosomal Long Noncoding RNAs Revealed a Novel ceRNA Network in Stable COPD. Int J Chron Obstruct Pulmon Dis. 2023;18:1995–2007. doi:10.2147/COPD.S414901

22. Gosselink JV, Hayashi S, Elliott WM, Xing L, Chan B, Yang L, Wright C, Sin D, Paré PD, Pierce JA, et al. Differential expression of tissue repair genes in the pathogenesis of chronic obstructive pulmonary disease. Am J Respir Crit Care Med. 2010;181(12):1329–1335. doi:10.1164/rccm.200812-1902OC

23. DeMeo D, Mariani T, Lange C, Lake S, Litonjua A, Celedón J, Reilly J, Chapman HA, Sparrow D, Spira A, et al. The SERPINE2 gene is associated with chronic obstructive pulmonary disease. Proc Am Thorac Soc. 2006;3(6):502. doi:10.1513/pats.200603-070MS

24. Hegab AE, Sakamoto T, Saitoh W, Nomura A, Ishii Y, Morishima Y, Iizuka T, Kiwamoto T, Matsuno Y, Massoud HH, Massoud HM, Hassanein KM, Sekizawa K., et al. Polymorphisms of TNFalpha, IL1beta, and IL1RN genes in chronic obstructive pulmonary disease. Biochem Biophys Res Commun. 2005;329(4):1246–1252. doi:10.1016/j.bbrc.2005.02.099

25. Celedón JC, Lange C, Raby BA, Litonjua AA, Palmer LJ, DeMeo DL, Reilly JJ, Kwiatkowski DJ, Chapman HA, Laird N, et al. The transforming growth factor-beta1 (TGFB1) gene is associated with chronic obstructive pulmonary disease (COPD). Hum Mol Genet. 2004;13(15):1649–1656. doi:10.1093/hmg/ddh171

26. Vacca G, Schwabe K, Dück R, Hlawa HP, Westphal A, Pabst S, Grohé C, Gillissen A. Polymorphisms of the beta2 adrenoreceptor gene in chronic obstructive pulmonary disease. Ther Adv Respir Dis. 2009;3(1):3–10. doi:10.1177/1753465809102553

27. He Q, Li H, Rui Y, Liu L, He B, Shi Y, Su X. Pentraxin 3 Gene Polymorphisms and Pulmonary Aspergillosis in Chronic Obstructive Pulmonary Disease Patients. Clin Infect Dis. 2018;66(2):261–267. doi:10.1093/cid/cix749

28. Fu X, Zhang F. Role of the HIF-1 signaling pathway in chronic obstructive pulmonary disease. Exp Ther Med. 2018;16(6):4553–4561. doi:10.3892/etm.2018.6785

29. Qu J, Yue L, Gao J, Yao H. Perspectives on Wnt Signal Pathway in the Pathogenesis and Therapeutics of Chronic Obstructive Pulmonary Disease. J Pharmacol Exp Ther. 2019;369(3):473–480. doi:10.1124/jpet.118.256222

30. Moradi S, Jarrahi E, Ahmadi A, Salimian J, Karimi M, Zarei A, Azimzadeh Jamalkandi S, Ghanei M. PI3K signalling in chronic obstructive pulmonary disease and opportunities for therapy. J Pathol. 2021;254(5):505–518. doi:10.1002/path.5696

31. Hu X, Hong B, Sun M. Peitu Shengjin Recipe Attenuates Airway Inflammation via the TLR4/NF-kB Signaling Pathway on Chronic Obstructive Pulmonary Disease. Evid Based Complement Alternat Med. 2022;2022:2090478. doi:10.1155/2022/2090478

32. Pomiès P, Blaquière M, Maury J, Mercier J, Gouzi F, Hayot M. Involvement of the FoxO1/MuRF1/Atrogin-1 Signaling Pathway in the Oxidative Stress-Induced Atrophy of Cultured Chronic Obstructive Pulmonary Disease Myotubes. PLoS One. 2016;11(8):e0160092. doi:10.1371/journal.pone.0160092

33. de Fays C, Geudens V, Gyselinck I, Kerckhof P, Vermaut A, Goos T, Vermant M, Beeckmans H, Kaes J, Van Slambrouck J, et al. Mucosal immune alterations at the early onset of tissue destruction in chronic obstructive pulmonary disease. Front Immunol. 2023;14:1275845. doi:10.3389/fimmu.2023.1275845

34. Clough E, Barrett T. The Gene Expression Omnibus Database. Methods Mol Biol. 2016;1418:93–110. doi:10.1007/978-1-4939-3578-9_5

35. Ritchie ME, Phipson B, Wu D, Hu Y, Law CW, Shi W, Smyth GK. limma powers differential expression analyses for RNA-sequencing and microarray studies. Nucleic Acids Res. 2015;43(7):e47. doi:10.1093/nar/gkv007

36. Solari A, Goeman JJ. Minimally adaptive BH: A tiny but uniform improvement of the procedure of Benjamini and Hochberg. Biom J. 2017;59(4):776–780. doi:10.1002/bimj.201500253

37. Reimand J, Kull M, Peterson H, Hansen J, Vilo J. g:Profiler--a web-based toolset for functional profiling of gene lists from large-scale experiments. Nucleic Acids Res. 2007;35(Web Server issue):W193–W200. doi:10.1093/nar/gkm226

38. Thomas PD. The Gene Ontology and the Meaning of Biological Function. Methods Mol Biol. 2017;1446:15L24. doi:10.1007/978-1-4939-3743-1_2

39. Fabregat A, Jupe S, Matthews L, Sidiropoulos K, Gillespie M, Garapati P, Haw R, Jassal B, Korninger F, May B et al The Reactome Pathway Knowledgebase. Nucleic Acids Res. 2018;46(D1):D649–D655. doi:10.1093/nar/gkx1132

40. Dimitrakopoulos GN, Klapa MI, Moschonas NK. PICKLE 3.0: enriching the human meta-database with the mouse protein interactome extended via mouse-human orthology. Bioinformatics. 2021;37(1):145–146. doi:10.1093/bioinformatics/btaa1070

41. Shannon P, Markiel A, Ozier O, Baliga NS, Wang JT, Ramage D, Amin N, Schwikowski B, Ideker T Cytoscape: a software environment for integrated models of biomolecular interaction networks. Genome Res 2003;13(11):2498–2504. doi:10.1101/gr.1239303

42. Luo X, Guo L, Dai XJ, Wang Q, Zhu W, Miao X, Gong H. Abnormal intrinsic functional hubs in alcohol dependence: evidence from a voxelwise degree centrality analysis. Neuropsychiatr Dis Treat. 2017;13:2011–2020. doi:10.2147/NDT.S142742

43. Li Y, Li W, Tan Y, Liu F, Cao Y, Lee KY. Hierarchical Decomposition for Betweenness Centrality Measure of Complex Networks. Sci Rep. 2017;7:46491. doi:10.1038/srep46491

44. Gilbert M, Li Z, Wu XN, Rohr L, Gombos S, Harter K, Schulze WX. Comparison of path-based centrality measures in protein-protein interaction networks revealed proteins with phenotypic relevance during adaptation to changing nitrogen environments. J Proteomics. 2021;235:104114. doi:10.1016/j.jprot.2021.104114

45. Li G, Li M, Wang J, Li Y, Pan Y. United Neighborhood Closeness Centrality and Orthology for Predicting Essential Proteins. IEEE/ACM Trans Comput Biol Bioinform. 2020;17(4):1451–1458. doi:10.1109/TCBB.2018.2889978

46. Zaki N, Efimov D, Berengueres J. Protein complex detection using interaction reliability assessment and weighted clustering coefficient. BMC Bioinformatics. 2013;14:163. doi:10.1186/1471-2105-14

47. Fan Y, Xia J (2018) miRNet-Functional Analysis and Visual Exploration of miRNA-Target Interactions in a Network Context. Methods Mol Biol 1819:215–233. doi:10.1007/978-1-4939-8618-7_10

48. Zhou G, Soufan O, Ewald J, Hancock REW, Basu N, Xia J (2019) NetworkAnalyst 3.0: a visual analytics platform for comprehensive gene expression profiling and meta-analysis. Nucleic Acids Res 47:W234–W241. doi:10.1093/nar/gkz240

49. Robin X, Turck N, Hainard A, Tiberti N, Lisacek F, Sanchez JC, Müller M. pROC: an open-source package for R and S+ to analyze and compare ROC curves. BMC Bioinformatics 2011;12:77. doi:10.1186/1471-2105-12-77

50. Zhang Q, Feng X, Hu W, Li C, Sun D, Peng Z, Wang S, Li H, Zhou M. Chronic obstructive pulmonary disease alters the genetic landscape and tumor immune microenvironment in lung cancer patients. Front Oncol. 2023;13:1169874. doi:10.3389/fonc.2023.1169874

51. van Eeden SF, Hogg JC. Immune-Modulation in Chronic Obstructive Pulmonary Disease: Current Concepts and Future Strategies. Respiration. 2020;99(7):550–565. doi:10.1159/000502261

52. Kang N, Qiu WJ, Wang B, Tang DF, Shen XY. Role of hemoglobin alpha and hemoglobin beta in non-small-cell lung cancer based on bioinformatics analysis. Mol Carcinog. 2022;61(6):587–602. doi:10.1002/mc.23404

53. Kohno T, Kunitoh H, Mimaki S, Shiraishi K, Kuchiba A, Yamamoto S, Yokota J. Contribution of the TP53, OGG1, CHRNA3, and HLA-DQA1 genes to the risk for lung squamous cell carcinoma. J Thorac Oncol. 2011;6(4):813–817. doi:10.1097/JTO.0b013e3181ee80ef

54. Su T, Liu P, Ti X, Wu S, Xue X, Wang Z, Dioum E, Zhang Q. EGR1 and SP1 co-regulate the erythropoietin receptor expression under hypoxia: an essential role in the growth of non-small cell lung cancer cells. Cell Commun Signal. 2019;17(1):152. doi:10.1186/s12964-019-0458-8

55. Qian X, Zhu L, Xu M, Liu H, Yu X, Shao Q, Qin J. Shikonin suppresses small cell lung cancer growth via inducing ATF3-mediated ferroptosis to promote ROS accumulation. Chem Biol Interact. 2023;382:110588. doi:10.1016/j.cbi.2023.110588

56. Du R, Jiang F, Yin Y, Xu J, Li X, Hu L, Wang X. Knockdown of lncRNA X inactive specific transcript (XIST) radiosensitizes non-small cell lung cancer (NSCLC) cells through regulation of miR-16-5p/WEE1 G2 checkpoint kinase (WEE1) axis. Int J Immunopathol Pharmacol. 2021;35:2058738420966087. doi:10.1177/2058738420966087

57. Matsubara E, Komohara Y, Shinchi Y, Mito R, Fujiwara Y, Ikeda K, Shima T, Shimoda M, Kanai Y, Sakagami T, et al. CD163-positive cancer cells are a predictor of a worse clinical course in lung adenocarcinoma. Pathol Int. 2021;71(10):666–673. doi:10.1111/pin.13144

58. Wong SW, McCarroll J, Hsu K, Geczy CL, Tedla N. Intranasal Delivery of Recombinant S100A8 Protein Delays Lung Cancer Growth by Remodeling the Lung Immune Microenvironment. Front Immunol. 2022;13:826391. doi:10.3389/fimmu.2022.826391

59. Liu Y, Liu B, Qiao YC, Niu WY. A New Case of Hb Headington (HBB: c.217A>C) Due to a New DNA Transversion, Found in a Patient with Type 2 Diabetes Mellitus. Hemoglobin. 2022;46(3):180–183. doi:10.1080/03630269.2022.2067044

60. Bartley PC, Hambling TE. Haemoglobin A1 in diabetes mellitus. Aust N Z J Med. 1979;9(1):49–53. doi:10.1111/j.1445-5994.1979.tb04112.x

61. Soetjipto, Rochmah N, Faizi M, Hisbiyah Y, Endaryanto A. HLA-DQA1 and HLA-DQB1 Gene Polymorphism in Indonesian Children with Type I Diabetes Mellitus. Appl Clin Genet. 2022;15:11–17. doi:10.2147/TACG.S348115

62. Shen N, Jiang S, Lu JM, Yu X, Lai SS, Zhang JZ, Zhang JL, Tao WW, Wang XX, Xu N, et al. The constitutive activation of Egr-1/C/EBPa mediates the development of type 2 diabetes mellitus by enhancing hepatic gluconeogenesis. Am J Pathol. 2015;185(2):513–523. doi:10.1016/j.ajpath.2014.10.016

63. Zhao Y, Du Y, Gao Y, Xu Z, Zhao D, Yang M. ATF3 Regulates Osteogenic Function by Mediating Osteoblast Ferroptosis in Type 2 Diabetic Osteoporosis. Dis Markers. 2022;2022:9872243. doi:10.1155/2022/9872243

64. Sohrabifar N, Ghaderian SMH, Alipour Parsa S, Ghaedi H, Jafari H. Variation in the expression level of MALAT1, MIAT and XIST lncRNAs in coronary artery disease patients with and without type 2 diabetes mellitus. Arch Physiol Biochem. 2022;128(5):1308–1315. doi:10.1080/13813455.2020.1768410

65. Tzouvelekis A, Herazo-Maya JD, Ryu C, Chu JH, Zhang Y, Gibson KF, Adonteng-Boateng PK, Li Q, Pan H, Cherry B, et al. S100A12 as a marker of worse cardiac output and mortality in pulmonary hypertension. Respirology. 2018;23(8):771–779. doi:10.1111/resp.13302

66. Sørensen LP, Parkner T, Søndergaard E, Bibby BM, Møller HJ, Nielsen S. Visceral obesity is associated with increased soluble CD163 concentration in men with type 2 diabetes mellitus. Endocr Connect. 2015;4(1):27–36. doi:10.1530/EC-14-0107

67. Lim RR, Vaidya T, Gadde SG, Yadav NK, Sethu S, Hainsworth DP, Mohan RR, Ghosh A, Chaurasia SS. Correlation between systemic S100A8 and S100A9 levels and severity of diabetic retinopathy in patients with type 2 diabetes mellitus. Diabetes Metab Syndr. 2019;13(2):1581–1589. doi:10.1016/j.dsx.2019.03.014

68. Hill QA, Farrar L, Lordan J, Gallienne A, Henderson S. A combination of two novel alpha globin variants Hb Bridlington (HBA1) and Hb Taybe (HBA2) resulting in severe hemolysis, pulmonary hypertension, and death. Hematology. 2015;20(1):50–52. doi:10.1179/1607845414Y.0000000164

69. Laggner M, Oberndorfer F, Golabi B, Bauer J, Zuckermann A, Hacker P, Lang I, Skoro-Sajer N, Gerges C, Taghavi S, et al. EGR1 Is Implicated in Right Ventricular Cardiac Remodeling Associated with Pulmonary Hypertension. Biology (Basel). 2022;11(5):677. doi:10.3390/biology11050677

70. Qin S, Predescu D, Carman B, Patel P, Chen J, Kim M, Lahm T, Geraci M, Predescu SA. Up-Regulation of the Long Noncoding RNA X-Inactive-Specific Transcript and the Sex Bias in Pulmonary Arterial Hypertension. Am J Pathol. 2021;191(6):1135–1150. doi:10.1016/j.ajpath.2021.03.009

71. Tzouvelekis A, Herazo-Maya JD, Ryu C, Chu JH, Zhang Y, Gibson KF, Adonteng-Boateng PK, Li Q, Pan H, Cherry B, et al. S100A12 as a marker of worse cardiac output and mortality in pulmonary hypertension. Respirology. 2018;23(8):771–779. doi:10.1111/resp.13302

72. Jasiewicz M, Kowal K, Kowal-Bielecka O, Knapp M, Skiepko R, Bodzenta-Lukaszyk A, Sobkowicz B, Musial WJ, Kaminski KA. Serum levels of CD163 and TWEAK in patients with pulmonary arterial hypertension. Cytokine. 2014;66(1):40–45. doi:10.1016/j.cyto.2013.12.013

73. Agapov E, Battaile JT, Tidwell R, Hachem R, Patterson GA, Pierce RA, Atkinson JJ, Holtzman MJ. Macrophage chitinase 1 stratifies chronic obstructive lung disease. Am J Respir Cell Mol Biol. 2009;41(4):379–384. doi:10.1165/rcmb.2009-0122rc

74. Hattori N, Nakagawa T, Yoneda M, Hayashida H, Nakagawa K, Yamamoto K, Htun MW, Shibata Y, Koji T, Ito T. Compounds in cigarette smoke induce EGR1 expression via the AHR, resulting in apoptosis and COPD. J Biochem. 2022;172(6):365–376. doi:10.1093/jb/mvac077

75. Lorenz E, Muhlebach MS, Tessier PA, Alexis NE, Duncan Hite R, Seeds MC, Peden DB, Meredith W. Different expression ratio of S100A8/A9 and S100A12 in acute and chronic lung diseases. Respir Med. 2008;102(4):567–573. doi:10.1016/j.rmed.2007.11.011

76. Higham A, Baker JM, Jackson N, Shah R, Lea S, Singh D. Dysregulation of the CD163-Haptoglobin Axis in the Airways of COPD Patients. Cells. 2021;11(1):2. doi:10.3390/cells11010002

77. Huang SJ, Ding ZN, Xiang HX, Fu L, Fei J. Association Between Serum S100A8/S100A9 Heterodimer and Pulmonary Function in Patients with Acute Exacerbation of Chronic Obstructive Pulmonary Disease. Lung. 2020;198(4):645–652. doi:10.1007/s00408-020-00376-9

78. Sklepkiewicz P, Dymek BA, Mlacki M, Koralewski R, Mazur M, Nejman-Gryz P, Korur S, Zagozdzon A, Rymaszewska A, von der Thüsen JH, et al. Inhibition of CHIT1 as a novel therapeutic approach in idiopathic pulmonary fibrosis. Eur J Pharmacol. 2022;919:174792. doi:10.1016/j.ejphar.2022.174792

79. Ghatak S, Markwald RR, Hascall VC, Dowling W, Lottes RG, Baatz JE, Beeson G, Beeson CC, Perrella MA, Thannickal VJ, et al. Transforming growth factor β1 (TGFβ1) regulates CD44V6 expression and activity through extracellular signal-regulated kinase (ERK)-induced EGR1 in pulmonary fibrogenic fibroblasts. J Biol Chem. 2017;292(25):10465–10489. doi:10.1074/jbc.M116.752451

80. Wei Y, Sun L, Liu C, Li L. Naringin regulates endoplasmic reticulum stress and mitophagy through the ATF3/PINK1 signaling axis to alleviate pulmonary fibrosis. Naunyn Schmiedebergs Arch Pharmacol. 2023;396(6):1155–1169. doi:10.1007/s00210-023-02390-z

81. Ding D, Luan R, Xue Q, Yang J. Prognostic significance of peripheral blood S100A12, S100A8, and S100A9 concentrations in idiopathic pulmonary fibrosis. Cytokine. 2023;172:156387. doi:10.1016/j.cyto.2023.156387

82. Chauvin P, Morzadec C, de Latour B, Llamas-Gutierrez F, Luque-Paz D, Jouneau S, Vernhet L. Soluble CD163 is produced by monocyte-derived and alveolar macrophages, and is not associated with the severity of idiopathic pulmonary fibrosis. Innate Immun. 2022;28(3-4):138–151. doi:10.1177/17534259221097835

83. Laranu I, Răcătăianu N, Drugan C, Cătană CS, Mirea AM, Miclea D, Bolboacă SD. Exploratory Longitudinal Analysis of the Circulating CHIT1 Activity in Pediatric Patients with Obesity. Children (Basel). 2023;10(1):124. doi:10.3390/children10010124

84. Zhang J, Zhang Y, Sun T, Guo F, Huang S, Chandalia M, Abate N, Fan D, Xin HB, Chen YE, et al. Dietary obesity-induced Egr-1 in adipocytes facilitates energy storage via suppression of FOXC2. Sci Rep. 2013;3:1476. doi:10.1038/srep01476

85. Ku HC, Chan TY, Chung JF, Kao YH, Cheng CF. The ATF3 inducer protects against diet-induced obesity via suppressing adipocyte adipogenesis and promoting lipolysis and browning. Biomed Pharmacother. 2022;145:112440. doi:10.1016/j.biopha.2021.112440

86. Wu C, Fang S, Zhang H, Li X, Du Y, Zhang Y, Lin X, Wang L, Ma X, et al. Long noncoding RNA XIST regulates brown preadipocytes differentiation and combats high-fat diet induced obesity by targeting C/EBPα. Mol Med. 2022;28(1):6. doi:10.1186/s10020-022-00434-3

87. Pan X, Yang L, Wang S, Liu Y, Yue L, Chen S. Semaglutide ameliorates obesity-induced cardiac inflammation and oxidative stress mediated via reduction of neutrophil Cxcl2, S100a8, and S100a9 expression. Mol Cell Biochem. 2023. doi:10.1007/s11010-023-04784-2

88. Zhao Y, Sui L, Wu P, Li L, Liu L, Ma B, Wang W, Chi H, Wang ZD, Wei Z, et al. EGR1 functions as a new host restriction factor for SARS-CoV-2 to inhibit virus replication through the E3 ubiquitin ligase MARCH8. J Virol. 2023. doi:10.1128/jvi.01028-23

89. Domachowske JB, Dyer KD, Bonville CA, Rosenberg HF. Recombinant human eosinophil-derived neurotoxin/RNase 2 functions as an effective antiviral agent against respiratory syncytial virus. J Infect Dis. 1998;177(6):1458–1464. doi:10.1086/515322

90. Wang TY, Liu YG, Li L, Wang G, Wang HM, Zhang HL, Zhao SF, Gao JC, An TQ, Tian ZJ, et al. Porcine alveolar macrophage CD163 abundance is a pivotal switch for porcine reproductive and respiratory syndrome virus infection. Oncotarget. 2018;9(15):12174–12185. doi:10.18632/oncotarget.24040

91. Mellett L, Khader SA. S100A8/A9 in COVID-19 pathogenesis: Impact on clinical outcomes. Cytokine Growth Factor Rev. 2022;63:90–97. doi:10.1016/j.cytogfr.2021.10.004

92. Xiang Y, Peng J, Liu Q, Guo C. The association between CD69 and EGR1 levels, and CHD patients without reflow after PCI. Exp Ther Med. 2019;17(5):3913–3920. doi:10.3892/etm.2019.7411

93. Huang Y, Dai H. ATF3 affects myocardial fibrosis remodeling after myocardial infarction by regulating autophagy and its mechanism of action. Gene. 2023;885:147705. doi:10.1016/j.gene.2023.147705

94. Xiao L, Gu Y, Sun Y, Chen J, Wang X, Zhang Y, Gao L, Li L. The long noncoding RNA XIST regulates cardiac hypertrophy by targeting miR-101. J Cell Physiol. 2019;234(8):13680–13692. doi:10.1002/jcp.28047

95. He YY, Yan W, Liu CL, Li X, Li RJ, Mu Y, Jia Q, Wu FF, Wang LL, He KL. Usefulness of S100A12 as a prognostic biomarker for adverse events in patients with heart failure. Clin Biochem. 2015;48(4-5):329–333. doi:10.1016/j.clinbiochem.2014.11.016

96. Bonora BM, Palano MT, Testa G, Fadini GP, Sangalli E, Madotto F, Persico G, Casciaro F, Vono R, Colpani O, et al. Hematopoietic progenitor cell liabilities and alarmins S100A8/A9-related inflammaging associate with frailty and predict poor cardiovascular outcomes in older adults. Aging Cell. 2022;21(3):e13545. doi:10.1111/acel.13545

97. Wang C, Zhang X, Chen R, Zhu X, Lian N. EGR1 mediates METTL3/m6A/CHI3L1 to promote osteoclastogenesis in osteoporosis. Genomics. 2023;115(5):110696. doi:10.1016/j.ygeno.2023.110696

98. Chen X, Ma F, Zhai N, Gao F, Cao G. Long nonLcoding RNA XIST inhibits osteoblast differentiation and promotes osteoporosis via Nrf2 hyperactivation by targeting CUL3. Int J Mol Med. 2021;48(1):137. doi:10.3892/ijmm.2021.4970

99. Gallo FT, Katche C, Morici JF, Medina JH, Weisstaub NV. Immediate Early Genes, Memory and Psychiatric Disorders: Focus on c-Fos, Egr1 and Arc. Front Behav Neurosci. 2018;12:79. doi:10.3389/fnbeh.2018.00079

100. Gamboa-Sánchez C, Becerril-Villanueva E, Alvarez-Herrera S, Leyva-Mascareño G, González-López SL, Estudillo E, Fernández-Molina AE, Elizalde-Contreras JM, Ruiz-May E, Segura-Cabrera A, et al. Upregulation of S100A8 in peripheral blood mononuclear cells from patients with depression treated with SSRIs: a pilot study. Proteome Sci. 2023;21(1):23. doi:10.1186/s12953-023-00224-7

101. Yan F, Wu Y, Liu H, Wu Y, Shen H, Li W. ATF3 is positively involved in particulate matter-induced airway inflammation in vitro and in vivo. Toxicol Lett. 2018;287:113–121. doi:10.1016/j.toxlet.2018.01.022

102. Camoretti-Mercado B, Karrar E, Nuñez L, Bowman MA. S100A12 and the Airway Smooth Muscle: Beyond Inflammation and Constriction. J Allergy Ther. 2012;3(Suppl 1):S1–007. doi:10.4172/2155-6121.S1-007

103. Dai C, Yao X, Gordon EM, Barochia A, Cuento RA, Kaler M, Meyer KS, Keeran KJ, Nugent GZ, Jeffries KR, et al. A CCL24-dependent pathway augments eosinophilic airway inflammation in house dust mite-challenged Cd163(-/-) mice. Mucosal Immunol. 2016;9(3):702–717. doi:10.1038/mi.2015.94

104. Van Crombruggen K, Vogl T, Pérez-Novo C, Holtappels G, Bachert C. Differential release and deposition of S100A8/A9 proteins in inflamed upper airway tissue. Eur Respir J. 2016;47(1):264–274. doi:10.1183/13993003.00159-2015

105. Wang J, Cheng W, Wang Z, Xin L, Zhang W. ATF3 inhibits the inflammation induced by Mycoplasma pneumonia in vitro and in vivo. Pediatr Pulmonol. 2017;52(9):1163–1170. doi:10.1002/ppul.23705

106. Jiang X, Huang CM, Feng CM, Xu Z, Fu L, Wang XM. Associations of Serum S100A12 With Severity and Prognosis in Patients With Community-Acquired Pneumonia: A Prospective Cohort Study. Front Immunol. 2021;12:714026. doi:10.3389/fimmu.2021.714026

107. Taira M, Matsumura T, Sumita Y, Moriyama M, Kondo M, Ishikawa N, Wada Y, Nagase M, Nishikawa E, Tsubata Y, et al. A Rare Case of Acute Fibrinous and Organizing Pneumonia Associated with Systemic Lupus Erythematosus and Autoimmune-associated Hemophagocytic Syndrome: The Involvement of CD163-positive Macrophages. Intern Med. 2022;61(4):559–565. doi:10.2169/internalmedicine.7184-21

108. Xie S, Wang J, Tuo W, Zhuang S, Cai Q, Yao C, Han F, Zhu H, Xiang Y, Yuan C. Serum level of S100A8/A9 as a biomarker for establishing the diagnosis and severity of community-acquired pneumonia in children. Front Cell Infect Microbiol. 2023;13:1139556. doi:10.3389/fcimb.2023.1139556

109. McGrath JJC, Stampfli MR. The immune system as a victim and aggressor in chronic obstructive pulmonary disease. J Thorac Dis. 2018;10(Suppl 17):S2011–S2017. doi:10.21037/jtd.2018.05.63

110. Murugan V, Peck MJ. Signal transduction pathways linking the activation of alveolar macrophages with the recruitment of neutrophils to lungs in chronic obstructive pulmonary disease. Exp Lung Res. 2009;35(6):439–485. doi:10.1080/01902140902759290

111. Åhrman E, Hallgren O, Malmström L, Hedström U, Malmström A, Bjermer L, Zhou XH, Westergren-Thorsson G, Malmström J. Quantitative proteomic characterization of the lung extracellular matrix in chronic obstructive pulmonary disease and idiopathic pulmonary fibrosis. J Proteomics. 2018;189:23–33. doi:10.1016/j.jprot.2018.02.027

112. Nurwidya F, Damayanti T, Yunus F. The Role of Innate and Adaptive Immune Cells in the Immunopathogenesis of Chronic Obstructive Pulmonary Disease. Tuberc Respir Dis (Seoul). 2016;79(1):5–13. doi:10.4046/trd.2016.79.1.5

113. Lodge KM, Vassallo A, Liu B, Long M, Tong Z, Newby PR, Agha-Jaffar D, Paschalaki K, Green CE, Belchamber KBR, et al. Hypoxia Increases the Potential for Neutrophil-mediated Endothelial Damage in Chronic Obstructive Pulmonary Disease. Am J Respir Crit Care Med. 2022;205(8):903–916. doi:10.1164/rccm.202006-2467OC

114. Kotlyarov S. Involvement of the Innate Immune System in the Pathogenesis of Chronic Obstructive Pulmonary Disease. Int J Mol Sci. 2022;23(2):985. doi:10.3390/ijms23020985

115. Engelen MPKJ, Kirschner SK, Coyle KS, Argyelan D, Neal G, Dasarathy S, Deutz NEP. Sex related differences in muscle health and metabolism in chronic obstructive pulmonary disease. Clin Nutr. 2023;42(9):1737–1746. doi:10.1016/j.clnu.2023.06.031

116. Chan SMH, Selemidis S, Bozinovski S, Vlahos R. Pathobiological mechanisms underlying metabolic syndrome (MetS) in chronic obstructive pulmonary disease (COPD): clinical significance and therapeutic strategies. Pharmacol Ther. 2019;198:160–188. doi:10.1016/j.pharmthera.2019.02.013

117. Kunter E, Ilvan A, Ozmen N, Demirer E, Ozturk A, Avsar K, Sayan O, Kartaloğlu Z. Effect of corticosteroids on hemostasis and pulmonary arterial pressure during chronic obstructive pulmonary disease exacerbation. Respiration. 2008;75(2):145–154. doi:10.1159/000097748

118. Mani S, Norel X, Varret M, Bchir S, Ben Anes A, Garrouch A, Tabka Z, Longrois D, Chahed K. Polymorphisms rs2745557 in PTGS2 and rs2075797 in PTGER2 are associated with the risk of chronic obstructive pulmonary disease development in a Tunisian cohort. Prostaglandins Leukot Essent Fatty Acids. 2021;166:102252. doi:10.1016/j.plefa.2021.102252

119. Yi E, Zhang J, Zheng M, Zhang Y, Liang C, Hao B, Hong W, Lin B, Pu J, Lin Z, et al. Long noncoding RNA IL6-AS1 is highly expressed in chronic obstructive pulmonary disease and is associated with interleukin 6 by targeting miR-149-5p and early B-cell factor 1. Clin Transl Med. 2021;11(7):e479. doi:10.1002/ctm2.479

120. He JQ, Shumansky K, Connett JE, Anthonisen NR, Paré PD, Sandford AJ. Association of genetic variations in the CSF2 and CSF3 genes with lung function in smoking-induced COPD. Eur Respir J. 2008;32(1):25–34. doi:10.1183/09031936.00040307

121. Castellucci M, Rossato M, Calzetti F, Tamassia N, Zeminian S, Cassatella MA, Bazzoni F. IL-10 disrupts the Brd4-docking sites to inhibit LPS-induced CXCL8 and TNF-α expression in monocytes: Implications for chronic obstructive pulmonary disease. J Allergy Clin Immunol. 2015;136(3):781–791.e9. doi:10.1016/j.jaci.2015.04.023

122. Inui T, Watanabe M, Nakamoto K, Sada M, Hirata A, Nakamura M, Honda K, Ogawa Y, Takata S, Yokoyama T, et al. Bronchial epithelial cells produce CXCL1 in response to LPS and TNFα: A potential role in the pathogenesis of COPD. Exp Lung Res. 2018;44(7):323–331. doi:10.1080/01902148.2018.1520936

123. Zhang Y, Wang L, Yan F, Yang M, Gao H, Zeng Y. Mettl3 Mediated m6A Methylation Involved in Epithelial-Mesenchymal Transition by Targeting SOCS3/STAT3/SNAI1 in Cigarette Smoking-Induced COPD. Int J Chron Obstruct Pulmon Dis. 2023;18:1007–1017. doi:10.2147/COPD.S398289

124. Yu W, Ye T, Ding J, Huang Y, Peng Y, Xia Q, Cuntai Z. miR-4456/CCL3/CCR5 Pathway in the Pathogenesis of Tight Junction Impairment in Chronic Obstructive Pulmonary Disease. Front Pharmacol. 2021;12:551839. doi:10.3389/fphar.2021.551839

125. Di Stefano A, Caramori G, Gnemmi I, Contoli M, Bristot L, Capelli A, Ricciardolo FL, Magno F, D’Anna SE, Zanini A, et al. Association of increased CCL5 and CXCL7 chemokine expression with neutrophil activation in severe stable COPD. Thorax. 2009;64(11):968–975. doi:10.1136/thx.2009.113647

126. Li J, Wang Q, Zhang Q, Wang Z, Wan X, Miao C, Zeng X. Higher Blood Vascular Cell Adhesion Molecule-1 is Related to the Increased Risk of Cardiovascular Events in Chronic Obstructive Pulmonary Disease. Int J Chron Obstruct Pulmon Dis. 2020;15:2289–2295. doi:10.2147/COPD.S264889

127. Bracke KR, Verhamme FM, Seys LJ, Bantsimba-Malanda C, Cunoosamy DM, Herbst R, Hammad H, Lambrecht BN, Joos GF, Brusselle GG. Role of CXCL13 in cigarette smoke-induced lymphoid follicle formation and chronic obstructive pulmonary disease. Am J Respir Crit Care Med. 2013;188(3):343–355. doi:10.1164/rccm.201211-2055OC

128. Chen J, Dai L, Wang T, He J, Wang Y, Wen F. The elevated CXCL5 levels in circulation are associated with lung function decline in COPD patients and cigarette smoking-induced mouse model of COPD. Ann Med. 2019;51(5-6):314–329. doi:10.1080/07853890.2019.1639809

129. Shen W, Weng Z, Fan M, Wang S, Wang R, Zhang Y, Tian H, Wang X, Wu X, Yang X, et al. Mechanisms by Which the MBD2/miR-301a-5p/CXCL12/CXCR4 Pathway Regulates Acute Exacerbations of Chronic Obstructive Pulmonary Disease. Int J Chron Obstruct Pulmon Dis. 2020;15:2561–2572. doi:10.2147/COPD.S261522

130. Zhang H, Li C, Song X, Cheng L, Liu Q, Zhang N, Wei L, Chung K, Adcock IM, Ling C, et al. Integrated analysis reveals lung fibrinogen gamma chain as a biomarker for chronic obstructive pulmonary disease. Ann Transl Med. 2021;9(24):1765. doi:10.21037/atm-21-5974

131. Yi G, Liang M, Li M, Fang X, Liu J, Lai Y, Chen J, Yao W, Feng X, Hu L, et al. A large lung gene expression study identifying IL1B as a novel player in airway inflammation in COPD airway epithelial cells. Inflamm Res. 2018;67(6):539–551. doi:10.1007/s00011-018-1145-8

132. Stolarczyk M, Amatngalim GD, Yu X, Veltman M, Hiemstra PS, Scholte BJ. ADAM17 and EGFR regulate IL-6 receptor and amphiregulin mRNA expression and release in cigarette smoke-exposed primary bronchial epithelial cells from patients with chronic obstructive pulmonary disease (COPD). Physiol Rep. 2016;4(16):e12878. doi:10.14814/phy2.12878

133. Xu X, Wang H, Li H, Cui X, Zhang H. SERPINE1 -844 and -675 polymorphisms and chronic obstructive pulmonary disease in a Chinese Han population. J Int Med Res. 2016;44(6):1292–1301. doi:10.1177/0300060516664270

134. Richards CD, Botelho F. Oncostatin M in the Regulation of Connective Tissue Cells and Macrophages in Pulmonary Disease. Biomedicines. 2019;7(4):95. doi:10.3390/biomedicines7040095

135. Daldegan MB, Teixeira MM, Talvani A. Concentration of CCL11, CXCL8 and TNF-alpha in sputum and plasma of patients undergoing asthma or chronic obstructive pulmonary disease exacerbation. Braz J Med Biol Res. 2005;38(9):1359–1365. doi:10.1590/s0100-879x2005000900010

136. Johansson SL, Roberts NB, Schlosser A, Andersen CB, Carlsen J, Wulf-Johansson H, Sækmose SG, Titlestad IL, Tornoe I, Miller B, et al. Microfibrillar-associated protein 4: a potential biomarker of chronic obstructive pulmonary disease. Respir Med. 2014;108(9):1336–1344. doi:10.1016/j.rmed.2014.06.003

137. Yan J, Wang A, Cao J, Chen L. Apelin/APJ system: an emerging therapeutic target for respiratory diseases. Cell Mol Life Sci. 2020;77(15):2919–2930. doi:10.1007/s00018-020-03461-7

138. Issac MS, Ashur W, Mousa H. Genetic polymorphisms of surfactant protein D rs2243639, Interleukin (IL)-1β rs16944 and IL-1RN rs2234663 in chronic obstructive pulmonary disease, healthy smokers, and non-smokers. Mol Diagn Ther. 2014;18(3):343–354. doi:10.1007/s40291-014-0084-5

139. Xu WH, Hu XL, Liu XF, Bai P, Sun YC. Peripheral Tc17 and Tc17/Interferon-γ Cells are Increased and Associated with Lung Function in Patients with Chronic Obstructive Pulmonary Disease. Chin Med J (Engl). 2016;129(8):909–916. doi:10.4103/0366-6999.179798

140. Hao W, Li M, Zhang C, Zhang Y, Du W. Increased levels of inflammatory biomarker CX3CL1 in patients with chronic obstructive pulmonary disease. Cytokine. 2020;126:154881. doi:10.1016/j.cyto.2019.154881

141. Yang J, Zhang MY, Du YM, Ji XL, Qu YQ. Identification and Validation of CDKN1A and HDAC1 as Senescence-Related Hub Genes in Chronic Obstructive Pulmonary Disease. Int J Chron Obstruct Pulmon Dis. 2022;17:1811–1825. doi:10.2147/COPD.S374684

142. Wang YX, Ji ML, Jiang CY, Qian ZB. Upregulation of ICAM-1 and IL-1β protein expression promotes lung injury in chronic obstructive pulmonary disease. Genet Mol Res. 2016;15(3):10.4238/gmr.15037971. doi:10.4238/gmr.15037971

143. Zhang Y, Wang L, Yan F, Yang M, Gao H, Zeng Y. Mettl3 Mediated m6A Methylation Involved in Epithelial-Mesenchymal Transition by Targeting SOCS3/STAT3/SNAI1 in Cigarette Smoking-Induced COPD. Int J Chron Obstruct Pulmon Dis. 2023;18:1007–1017. doi:10.2147/COPD.S398289

144. Liu Y, Liu X, Lin G, Sun L, Li H, Xie C. Decreased CD34+ cell number is correlated with cardiac dysfunction in patients with acute exacerbation of COPD. Heart Lung Circ. 2014;23(9):875–882. doi:10.1016/j.hlc.2014.03.008

145. Sun BB, Ma LJ, Qi Y, Zhang GJ. Correlation of IL-33 gene polymorphism with chronic obstructive pulmonary disease. Eur Rev Med Pharmacol Sci. 2019;23(14):6277–6282. doi:10.26355/eurrev_201907_18449

146. Takabatake N, Arao T, Sata M, Inoue S, Abe S, Shibata Y, Kubota I. Circulating levels of soluble Fas ligand in cachexic patients with COPD are higher than those in non-cachexic patients with COPD. Intern Med. 2005;44(11):1137–1143. doi:10.2169/internalmedicine.44.1137

147. Xu SC, Kuang JY, Liu J, Ma CL, Feng YL, Su ZG. Association between fibroblast growth factor 7 and the risk of chronic obstructive pulmonary disease. Acta Pharmacol Sin. 2012;33(8):998–1003. doi:10.1038/aps.2012.69

148. Yao Y, Zhou J, Diao X, Wang S. Association between tumor necrosis factor-α and chronic obstructive pulmonary disease: a systematic review and meta-analysis. Ther Adv Respir Dis. 2019;13:1753466619866096. doi:10.1177/1753466619866096

149. Lv Z, Liang G, Cheng M. Predictive Value of GDF-15 and sST2 for Pulmonary Hypertension in Acute Exacerbation of Chronic Obstructive Pulmonary Disease. Int J Chron Obstruct Pulmon Dis. 2023;18:2431–2438. doi:10.2147/COPD.S429334

150. Gao HX, Su Y, Zhang AL, Xu JW, Fu Q, Yan L. MiR-34c-5p plays a protective role in chronic obstructive pulmonary disease via targeting CCL22. Exp Lung Res. 2019;45(1-2):1–12. doi:10.1080/01902148.2018.1563925

151. Rybka J, Korte SM, Czajkowska-Malinowska M, Wiese M, Kędziora-Kornatowska K, Kędziora J. The links between chronic obstructive pulmonary disease and comorbid depressive symptoms: role of IL-2 and IFN-γ. Clin Exp Med. 2016;16(4):493–502. doi:10.1007/s10238-015-0391-0

152. Marques P, Collado A, Escudero P, Rius C, González C, Servera E, Piqueras L, Sanz MJ. Cigarette Smoke Increases Endothelial CXCL16-Leukocyte CXCR6 Adhesion In Vitro and In Vivo. Potential Consequences in Chronic Obstructive Pulmonary Disease. Front Immunol. 2017;8:1766. doi:10.3389/fimmu.2017.01766

153. Dang X, Yang L, Guo J, Hu H, Li F, Liu Y, Pang Y. miR-145-5p is associated with smoke-related chronic obstructive pulmonary disease via targeting KLF5. Chem Biol Interact. 2019;300:82–90. doi:10.1016/j.cbi.2019.01.011

154. Zhang J, He J, Xia J, Chen Z, Chen X. Delayed apoptosis by neutrophils from COPD patients is associated with altered Bak, Bcl-xl, and Mcl-1 mRNA expression. Diagn Pathol. 2012;7:65. doi:10.1186/1746-1596-7-65

155. Lewis A, Riddoch-Contreras J, Natanek SA, Donaldson A, Man WD, Moxham J, Hopkinson NS, Polkey MI, Kemp PR. Downregulation of the serum response factor/miR-1 axis in the quadriceps of patients with COPD. Thorax. 2012;67(1):26–34. doi:10.1136/thoraxjnl-2011-200309

156. Perea L, Rodrigo-Troyano A, Cantó E, Domínguez-Álvarez M, Giner J, Sanchez-Reus F, Villar-García J, Quero S, García-Núñez M, Marín A, et al. Reduced airway levels of fatty-acid binding protein 4 in COPD: relationship with airway infection and disease severity. Respir Res. 2020;21(1):21. doi:10.1186/s12931-020-1278-5

157. Lu H, Xu D, Yang Y, Feng Q, Sun J, Li Q, Zhao J, Zhou X, Niu H, Liu J, et al. Genetic Polymorphisms of CYP2C9/CYP2C19 in Chronic Obstructive Pulmonary Disease. COPD. 2020;17(5):595–600. doi:10.1080/15412555.2020.1780577

158. Zhang X, Li X, Ma W, Liu F, Huang P, Wei L, Li L, Qian Y. Astragaloside IV restores Th17/Treg balance via inhibiting CXCR4 to improve chronic obstructive pulmonary disease. Immunopharmacol Immunotoxicol. 2023;45(6):682–691. doi:10.1080/08923973.2023.2228479

159. Xia S, Qu J, Jia H, He W, Li J, Zhao L, Mao M, Zhao Y. Overexpression of Forkhead box C1 attenuates oxidative stress, inflammation and apoptosis in chronic obstructive pulmonary disease. Life Sci. 2019;216:75–84. doi:10.1016/j.lfs.2018.11.023

160. Yang Q, Huang W, Yin D, Zhang L, Gao Y, Tong J, Li Z. EPHX1 and GSTP1 polymorphisms are associated with COPD risk: a systematic review and meta-analysis. Front Genet. 2023;14:1128985. doi:10.3389/fgene.2023.1128985

161. Kamel AA, Hashem MK, AbdulKareem ES, Ali AH, Mahmoud EA, Abd-Elkader AS, Abdellatif H, Abdelbadea A, Abdel-Rady NM, Al Anany MGE, et al. Significant Interrelations among Serum Annexin A1, Soluble Receptor for Advanced Glycation End Products (sRAGE) and rs2070600 in Chronic Obstructive Pulmonary Disease. Biology (Basel). 2022;11(12):1707. doi:10.3390/biology11121707

162. Gu W, Yuan Y, Wang L, Yang H, Li S, Tang Z, Li Q. Long non-coding RNA TUG1 promotes airway remodelling by suppressing the miR-145-5p/DUSP6 axis in cigarette smoke-induced COPD. J Cell Mol Med. 2019;23(11):7200–7209. doi:10.1111/jcmm.14389

163. Siddhuraj P, Jönsson J, Alyamani M, Prabhala P, Magnusson M, Lindstedt S, Erjefält JS.Dynamically upregulated mast cell CPA3 patterns in chronic obstructive pulmonary disease and idiopathic pulmonary fibrosis. Front Immunol. 2022;13:924244. doi:10.3389/fimmu.2022.924244

164. Kim HT, Yin W, Jin YJ, Panza P, Gunawan F, Grohmann B, Buettner C, Sokol AM, Preussner J, Guenther S, et al. Myh10 deficiency leads to defective extracellular matrix remodeling and pulmonary disease. Nat Commun. 2018;9(1):4600. doi:10.1038/s41467-018-06833-7

165. Li DY, Chen L, Miao SY, Zhou M, Wu JH, Sun SW, Liu LL, Qi C, Xiong XZ. Inducible Costimulator-C-X-C Motif Chemokine Receptor 3 Signaling is Involved in Chronic Obstructive Pulmonary Disease Pathogenesis. Int J Chron Obstruct Pulmon Dis. 2022;17:1847–1861. doi:10.2147/COPD.S371801

166. Tsoumakidou M, Tousa S, Semitekolou M, Panagiotou P, Panagiotou A, Morianos I, Litsiou E, Trochoutsou AI, Konstantinou M, Potaris K, et al. Tolerogenic signaling by pulmonary CD1c+ dendritic cells induces regulatory T cells in patients with chronic obstructive pulmonary disease by IL-27/IL-10/inducible costimulator ligand. J Allergy Clin Immunol. 2014;134(4):944–954.e8. doi:10.1016/j.jaci.2014.05.045

167. Zhao X, Feng X, Liu P, Ye J, Tao R, Li R, Shen B, Zhang X, Wang X, Zhao D. Abnormal expression of CD96 on natural killer cell in peripheral blood of patients with chronic obstructive pulmonary disease. Clin Respir J. 2022;16(8):546–554. doi:10.1111/crj.13523

168. Jiang J, Wang M, Shen W, Wu J, Ma Q, Wang Z, Chen Z, Bian T, Ji N, Huang M, et al. CD146 deficiency aggravates chronic obstructive pulmonary disease via the increased production of S100A9 and MMP-9 in macrophages. Int Immunopharmacol. 2023. doi:10.1016/j.intimp.2023.111410

169. Marcolongo F, Scarlata S, Tomino C, De Dominicis C, Giacconi R, Malavolta M, Bonassi S, Russo P, Prinzi G. Psycho-cognitive assessment and quality of life in older adults with chronic obstructive pulmonary disease-carrying the rs4713916 gene polymorphism (G/A) of gene FKBP5 and response to pulmonary rehabilitation: a proof of concept study. Psychiatr Genet. 2022;32(3):116–124. doi:10.1097/YPG.0000000000000308

170. Zhou H, Ying X, Liu Y, Ye S, Yan J, Li Y. Genetic polymorphism of heme oxygenase 1 promoter in the occurrence and severity of chronic obstructive pulmonary disease: a meta-analysis. J Cell Mol Med. 2017;21(5):894–903. doi:10.1111/jcmm.13028

171. Crisford H, Sapey E, Stockley RA. Proteinase 3; a potential target in chronic obstructive pulmonary disease and other chronic inflammatory diseases. Respir Res. 2018;19(1):180. doi:10.1186/s12931-018-0883-z

172. Thomsen M, Nordestgaard BG, Kobzik L, Dahl M. Genetic variation in the scavenger receptor MARCO and its association with chronic obstructive pulmonary disease and lung infection in 10,604 individuals. Respiration. 2013;85(2):144–153. doi:10.1159/000342354

173. Kim RY, Sunkara KP, Bracke KR, Jarnicki AG, Donovan C, Hsu AC, Ieni A, Beckett EL, Galvão I, Wijnant S, et al. A microRNA-21-mediated SATB1/S100A9/NF-κB axis promotes chronic obstructive pulmonary disease pathogenesis. Sci Transl Med. 2021;13(621):eaav7223. doi:10.1126/scitranslmed.aav7223

174. Zeng Z, Li M, Wang M, Wu X, Li Q, Ning Q, Zhao J, Xu Y, Xie J. Increased expression of Siglec-9 in chronic obstructive pulmonary disease. Sci Rep. 2017;7(1):10116. doi:10.1038/s41598-017-09120-5

175. Kim EJ, Kim KM, Park SH, Kim JS, Lee WK, Cha SI, Kim CH, Kang YM, Han SB, Jung TH, et al. SLC11A1 polymorphisms are associated with the risk of chronic obstructive pulmonary disease in a Korean population. Biochem Genet. 2008;46(7-8):506–519. doi:10.1007/s10528-008-9166-6

176. Chen CZ, Ou CY, Wang RH, Lee CH, Lin CC, Chang HY, Hsiue TR. The role of bactericidal/permeability-increasing protein in men with chronic obstructive pulmonary disease. COPD. 2012;9(2):197–202. doi:10.3109/15412555.2011.654143

177. Cheng SL, Wang HC, Cheng SJ, Yu CJ. Elevated placenta growth factor predicts pneumonia in patients with chronic obstructive pulmonary disease under inhaled corticosteroids therapy. BMC Pulm Med. 2011;11:46. doi:10.1186/1471-2466-11-46

178. Fan LC, McConn K, Plataki M, Kenny S, Williams NC, Kim K, Quirke JA, Chen Y, Sauler M, Möbius ME, et al. Alveolar type II epithelial cell FASN maintains lipid homeostasis in experimental COPD. JCI Insight. 2023;8(16):e163403. doi:10.1172/jci.insight.163403

179. Qin HY, Li MD, Xie GF, Cao W, Xu DX, Zhao H, Fu L. Associations among S100A4, Sphingosine-1-Phosphate, and Pulmonary Function in Patients with Chronic Obstructive Pulmonary Disease. Oxid Med Cell Longev. 2022;2022:6041471. doi:10.1155/2022/6041471

180. Zhang LX, Ye J, Chen YB, Peng HL, Chen X, Liu L, Jiang AG, Huang JX. The effect of CD33 expression on inflammatory response in chronic obstructive pulmonary disease. Immunol Invest. 2013;42(8):701–710. doi:10.3109/08820139.2013.806542

181. Poznański M, Brzeziańska-Lasota E, Kiszałkiewicz J, Kurnatowska I, Kroczyńska-Bednarek J, Pękala-Wojciechowska A, Pietras T, Antczak A. Serum levels and gene expression of pentraxin 3 are elevated in COPD. Adv Med Sci. 2019;64(1):85–89. doi:10.1016/j.advms.2018.08.006

182. Kranenburg AR, de Boer WI, Alagappan VK, Sterk PJ, Sharma HS. Enhanced bronchial expression of vascular endothelial growth factor and receptors (Flk-1 and Flt-1) in patients with chronic obstructive pulmonary disease. Thorax. 2005;60(2):106–113. doi:10.1136/thx.2004.023986

183. Bækvad-Hansen M, Nordestgaard BG, Dahl M. Heterozygosity for E292V in ABCA3, lung function and COPD in 64,000 individuals. Respir Res. 2012;13(1):67. doi:10.1186/1465-9921-13-67

184. Schiffers C, van de Wetering C, Bauer RA, Habibovic A, Hristova M, Dustin CM, Lambrichts S, Vacek PM, Wouters EF, Reynaert NL, et al. Downregulation of epithelial DUOX1 in chronic obstructive pulmonary disease. JCI Insight. 2021;6(2):e142189. doi:10.1172/jci.insight.142189

185. Qiu J, Zhang YN, Zheng X, Zhang P, Ma G, Tan H. Notch promotes DNMT-mediated hypermethylation of Klotho leads to COPD-related inflammation. Exp Lung Res. 2018;44(7):368–377. doi:10.1080/01902148.2018.1556749

186. Tulah AS, Parker SG, Moffatt MF, Wardlaw AJ, Connolly MJ, Sayers I. The role of ALOX5AP, LTA4H and LTB4R polymorphisms in determining baseline lung function and COPD susceptibility in UK smokers. BMC Med Genet. 2011;12:173. doi:10.1186/1471-2350-12-173

187. Chen YC, Lin MC, Lee CH, Liu SF, Wang CC, Fang WF, Chao TY, Wu CC, Wei YF, Chang HC, et al. Defective formyl peptide receptor 2/3 and annexin A1 expressions associated with M2a polarization of blood immune cells in patients with chronic obstructive pulmonary disease. J Transl Med. 2018;16(1):69. doi:10.1186/s12967-018-1435-5

188. Hobbs BD, Parker MM, Chen H, Lao T, Hardin M, Qiao D, Hawrylkiewicz I, Sliwinski P, Yim JJ, Kim WJ, et al. Exome Array Analysis Identifies a Common Variant in IL27 Associated with Chronic Obstructive Pulmonary Disease. Am J Respir Crit Care Med. 2016;194(1):48–57. doi:10.1164/rccm.201510-2053OC

189. Skronska-Wasek W, Mutze K, Baarsma HA, Bracke KR, Alsafadi HN, Lehmann M, Costa R, Stornaiuolo M, Novellino E, Brusselle GG, et al. Reduced Frizzled Receptor 4 Expression Prevents WNT/β-Catenin-driven Alveolar Lung Repair in Chronic Obstructive Pulmonary Disease. Am J Respir Crit Care Med. 2017;196(2):172–185. doi:10.1164/rccm.201605-0904OC

190. Zhu A, Ge D, Zhang J, Teng Y, Yuan C, Huang M, Adcock IM, Barnes PJ, Yao X. Sputum myeloperoxidase in chronic obstructive pulmonary disease. Eur J Med Res. 2014;19(1):12. doi:10.1186/2047-783X-19-12

191. Bchir S, Boumiza S, Ben Nasr H, Garrouch A, Kallel I, Tabka Z, Chahed K. Impact of cathepsin D activity and C224T polymorphism (rs17571) on chronic obstructive pulmonary disease: correlations with oxidative and inflammatory markers. Clin Exp Med. 2021;21(3):457–465. doi:10.1007/s10238-021-00692-1

192. Milara J, Díaz-Platas L, Contreras S, Ribera P, Roger I, Ballester B, Montero P, Cogolludo Á, Morcillo E, Cortijo J. MUC1 deficiency mediates corticosteroid resistance in chronic obstructive pulmonary disease. Respir Res. 2018;19(1):226. doi:10.1186/s12931-018-0927-4

193. Angata T, Ishii T, Motegi T, Oka R, Taylor RE, Soto PC, Chang YC, Secundino I, Gao CX, Ohtsubo K, et al. Loss of Siglec-14 reduces the risk of chronic obstructive pulmonary disease exacerbation. Cell Mol Life Sci. 2013;70(17):3199–3210. doi:10.1007/s00018-013-1311-7

194. Merrilees MJ, Ching PS, Beaumont B, Hinek A, Wight TN, Black PN. Changes in elastin, elastin binding protein and versican in alveoli in chronic obstructive pulmonary disease. Respir Res. 2008;9(1):41. doi:10.1186/1465-9921-9-41

195. Tang L, Zhong X, Gong H, Tuerxun M, Ma T, Ren J, Xie C, Zheng A, Abudureheman Z, Abudukadeer A, et al. nalysis of the association of ANO3/MUC15, COL4A4, RRBP1, and KLK1 polymorphisms with COPD susceptibility in the Kashi population. BMC Pulm Med. 2022;22(1):178. doi:10.1186/s12890-022-01975-3

196. Bialek K, Czarny P, Watala C, Wigner P, Talarowska M, Galecki P, Szemraj J, Sliwinski T. Novel association between TGFA, TGFB1, IRF1, PTGS2 and IKBKB single-nucleotide polymorphisms and occurrence, severity and treatment response of major depressive disorder. PeerJ. 2020;8:e8676. doi:10.7717/peerj.8676

197. Nakayama T, Hirano F, Okushi Y, Matsuura K, Ohashi M, Matsumiya A, Yoshimura T. Orphan nuclear receptor nr4a1 regulates winter depression-like behavior in medaka. Neurosci Lett. 2023;814:137469. doi:10.1016/j.neulet.2023.137469

198. Ting EY, Yang AC, Tsai SJ. Role of Interleukin-6 in Depressive Disorder. Int J Mol Sci. 2020;21(6):2194. doi:10.3390/ijms21062194

199. Fanelli G, Benedetti F, Wang SM, Lee SJ, Jun TY, Masand PS, Patkar AA, Han C, Serretti A, Pae CU, et al. Reduced CXCL1/GRO chemokine plasma levels are a possible biomarker of elderly depression. J Affect Disord. 2019;249:410–417. doi:10.1016/j.jad.2019.02.042

200. Ni C, Jiang W, Wang Z, Wang Z, Zhang J, Zheng X, Liu Z, Ou H, Jiang T, Liang W. et al. LncRNA-AC006129.1 reactivates a SOCS3- mediated anti-inflammatory response through DNA methylation-mediated CIC downregulation in schizophrenia. Mol Psychiatry. 2021;26(8):4511–4528. doi:10.1038/s41380-020-0662-3

201. Huckans M, Wilhelm CJ, Phillips TJ, Huang ET, Hudson R, Loftis JM. Parallel Effects of Methamphetamine on Anxiety and CCL3 in Humans and a Genetic Mouse Model of High Methamphetamine Intake. Neuropsychobiology. 2017;75(4):169–177. doi:10.1159/000485129

202. Shi DD, Zhang YD, Zhang S, Liao BB, Chu MY, Su S, Zhuo K, Hu H, Zhang C, Wang Z. Stress-induced red nucleus attenuation induces anxiety-like behavior and lymph node CCL5 secretion. Nat Commun. 2023;14(1):6923. doi:10.1038/s41467-023-42814-1

203. Zhu X, Meng J, Han C, Wu Q, Du Y, Qi J, Wei L, Li H, He W, Zhang K, et al. CCL2-mediated inflammatory pathogenesis underlies high myopia-related anxiety. Cell Discov. 2023;9(1):94. doi:10.1038/s41421-023-00588-2

204. Song X, Sun N, Zhang A, Lei L, Li X, Liu Z, Wang Y, Yang C, Zhang K. Association Between NR4A2 Gene Polymorphism and Depressive Symptoms and Antidepressant Effect. Neuropsychiatr Dis Treat. 2021;17:2613–2623. doi:10.2147/NDT.S319548

205. Yang L, Wang M, Guo YY, Sun T, Li YJ, Yang Q, Zhang K, Liu SB, Zhao MG, Wu YM. Systemic inflammation induces anxiety disorder through CXCL12/CXCR4 pathway. Brain Behav Immun. 2016;56:352–362. doi:10.1016/j.bbi.2016.03.001

206. Zhao Y, Wang S, Chu Z, Dang Y, Zhu J, Su X. MicroRNA-101 in the ventrolateral orbital cortex (VLO) modulates depressive-like behaviors in rats and targets dual-specificity phosphatase 1 (DUSP1). Brain Res. 2017;1669:55–62. doi:10.1016/j.brainres.2017.05.020

207. Yin Q, Du T, Yang C, Li X, Zhao Z, Liu R, Yang B, Liu B. Gadd45b is a novel mediator of depression-like behaviors and neuroinflammation after cerebral ischemia. Biochem Biophys Res Commun. 2021;554:107–113. doi:10.1016/j.bbrc.2021.03.104

208. Wang Y, Liu W, Xiao Y, Yuan H, Wang F, Jiang P, Luo Z. Association of Apelin and Apelin Receptor Polymorphisms With the Risk of Comorbid Depression and Anxiety in Coronary Heart Disease Patients. Front Genet. 2020;11:893. doi:10.3389/fgene.2020.00893

209. Kovacs D, Eszlari N, Petschner P, Pap D, Vas S, Kovacs P, Gonda X, Juhasz G, Bagdy G. Effects of IL1B single nucleotide polymorphisms on depressive and anxiety symptoms are determined by severity and type of life stress. Brain Behav Immun. 2016;56:96–104. doi:10.1016/j.bbi.2016.02.012

210. Fang Y, Zhang L, Zeng Z, Lian Y, Jia Y, Zhu H, Xu Y. Promoter polymorphisms of SERPINE1 are associated with the antidepressant response to depression in Alzheimer’s disease. Neurosci Lett. 2012;516(2):217–220. doi:10.1016/j.neulet.2012.03.090

211. Teixeira AL, Reis HJ, Nicolato R, Brito-Melo G, Correa H, Teixeira MM, Romano-Silva MA. Increased serum levels of CCL11/eotaxin in schizophrenia. Prog Neuropsychopharmacol Biol Psychiatry. 2008;32(3):710–714. doi:10.1016/j.pnpbp.2007.11.019

212. Wang Y, Liu W, Xiao Y, Yuan H, Wang F, Jiang P, Luo Z. Association of Apelin and Apelin Receptor Polymorphisms With the Risk of Comorbid Depression and Anxiety in Coronary Heart Disease Patients. Front Genet. 2020;11:893.doi:10.3389/fgene.2020.00893

213. de Souza AG, Lopes IS, Filho AJMC, Cavalcante TMB, Oliveira JVS, de Carvalho MAJ, de Lima KA, Jucá PM, Mendonça SS, Mottin M, et al. Neuroprotective effects of dimethyl fumarate against depression-like behaviors via astrocytes and microglia modulation in mice: possible involvement of the HCAR2/Nrf2 signaling pathway. Naunyn Schmiedebergs Arch Pharmacol. 2022;395(9):1029–1045. doi:10.1007/s00210-022-02247-x

214. Chen S, Zhang Y, Yuan Y. The Combination of Serum BDNF, Cortisol and IFN-Gamma Can Assist the Diagnosis of Major Depressive Disorder. Neuropsychiatr Dis Treat. 2021;17:2819–2829. doi:10.2147/NDT.S322078

215. Liu Y, Zhang T, Meng D, Sun L, Yang G, He Y, Zhang C. Involvement of CX3CL1/CX3CR1 in depression and cognitive impairment induced by chronic unpredictable stress and relevant underlying mechanism. Behav Brain Res. 2020;381:112371. doi:10.1016/j.bbr.2019.112371

216. Engel DF, de Oliveira J, Lopes JB, Santos DB, Moreira ELG, Farina M, Rodrigues ALS, de Souza Brocardo P, de Bem AF. Is there an association between hypercholesterolemia and depression? Behavioral evidence from the LDLr(-/-) mouse experimental model. Behav Brain Res. 2016;311:31–38. doi:10.1016/j.bbr.2016.05.029

217. Tseng CC, Wang SC, Yang YC, Fu HC, Chou CK, Kang HY, Hung YY. Aberrant Histone Modification of TNFAIP3, TLR4, TNIP2, miR-146a, and miR-155 in Major Depressive Disorder. Mol Neurobiol. 2023;60(8):4753–4760. doi:10.1007/s12035-023-03374-z

218. Tammiste A, Jiang T, Fischer K, Mägi R, Krjutškov K, Pettai K, Esko T, Li Y, Tansey KE, Carroll LS, et al. Whole-exome sequencing identifies a polymorphism in the BMP5 gene associated with SSRI treatment response in major depression. J Psychopharmacol. 2013;27(10):915–920. doi:10.1177/0269881113499829

219. Algahtani MM, Alshehri S, Alqarni SS, Ahmad SF, Al-Harbi NO, Alqarni SA, Alfardan AS, Ibrahim KE, Attia SM, Nadeem A. Inhibition of ITK Signaling Causes Amelioration in Sepsis-Associated Neuroinflammation and Depression-like State in Mice. Int J Mol Sci. 2023;24(9):8101. doi:10.3390/ijms24098101

220. Schaefer M, Horn M, Schmidt F, Schmid-Wendtner MH, Volkenandt M, Ackenheil M, Mueller N, Schwarz MJ. Correlation between sICAM-1 and depressive symptoms during adjuvant treatment of melanoma with interferon-alpha. Brain Behav Immun. 2004;18(6):555–562. doi:10.1016/j.bbi.2004.02.002

221. Kim K, Wi S, Seo JH, Pyo S, Cho SR. Reduced Interaction of Aggregated α-Synuclein and VAMP2 by Environmental Enrichment Alleviates Hyperactivity and Anxiety in a Model of Parkinson’s Disease. Genes (Basel). 2021;12(3):392. doi:10.3390/genes12030392

222. Larijani TT, Mohammadi S, Kasaeian A, Malek Mohammadi A, Mostafaei S, Alimoghaddam K, Ghavamzadeh A. How do anxiety affect CD34 and CD3 cells in allogeneic peripheral blood stem cell transplantation?. Transfus Apher Sci. 2018;57(1):107–110. doi:10.1016/j.transci.2018.01.006

223. Yu S, Wang G, Yao B, Xiao L, Tuo H. Arc and Homer1 are involved in comorbid epilepsy and depression: A microarray data analysis. Epilepsy Behav. 2022;132:108738. doi:10.1016/j.yebeh.2022.108738

224. Liu R, Liu L, Ren S, Wei C, Wang Y, Li D, Zhang W. The role of IL-33 in depression: a systematic review and meta-analysis. Front Psychiatry. 2023;14:1242367. doi:10.3389/fpsyt.2023.1242367

225. Xiao L, Loh YP. Neurotrophic Factor-α1/Carboxypeptidase E Functions in Neuroprotection and Alleviates Depression. Front Mol Neurosci. 2022;15:918852. doi:10.3389/fnmol.2022.918852

226. Shentu Y, Tian Q, Yang J, Liu X, Han Y, Yang D, Zhang N, Fan X, Wang P, Ma J, et al. Upregulation of KDM6B contributes to lipopolysaccharide-induced anxiety-like behavior via modulation of VGLL4 in mice. Behav Brain Res. 2021;408:113305. doi:10.1016/j.bbr.2021.113305

227. Yao L, Pan L, Qian M, Sun W, Gu C, Chen L, Tang X, Hu Y, Xu L, Wei Y, et al. Tumor Necrosis Factor-α Variations in Patients With Major Depressive Disorder Before and After Antidepressant Treatment. Front Psychiatry. 2020;11:518837. doi:10.3389/fpsyt.2020.518837

228. Li Y, Mei T, Sun T, Xiao X, Peng R. Altered circulating GDF-15 level predicts sex hormone imbalance in males with major depressive disorder. BMC Psychiatry. 2023;23(1):28. doi:10.1186/s12888-023-04527-z

229. Laurikainen H, Vuorela A, Toivonen A, Reinert-Hartwall L, Trontti K, Lindgren M, Keinänen J, Mäntylä T, Paju J, Ilonen T, Armio RL, et al. Elevated serum chemokine CCL22 levels in first-episode psychosis: associations with symptoms, peripheral immune state and in vivo brain glial cell function. Transl Psychiatry. 2020;10(1):94. doi:10.1038/s41398-020-0776-z

230. Freff J, Beins EC, Bröker L, Schwarte K, Leite Dantas R, Maj C, Arolt V, Dannlowski U, Nöthen MM, Baune BT, et al. Chemokine receptor 4 expression on blood T lymphocytes predicts severity of major depressive disorder. J Affect Disord. 2022;310:343–353. doi:10.1016/j.jad.2022.05.003

231. Huang C, Zhang F, Li P, Song C. Low-Dose IL-2 Attenuated Depression-like Behaviors and Pathological Changes through Restoring the Balances between IL-6 and TGF-β and between Th17 and Treg in a Chronic Stress-Induced Mouse Model of Depression. Int J Mol Sci. 2022;23(22):13856. doi:10.3390/ijms232213856

232. Hung WC, Yu TH, Wu CC, Lee TL, Tsai IT, Hsuan CF, Chen CY, Chung FM, Lee YJ, Tang WH. FABP3, FABP4, and heart rate variability among patients with chronic schizophrenia. Front Endocrinol (Lausanne). 2023;14:1165621. doi:10.3389/fendo.2023.1165621

233. Arnone D, Omar O, Arora T, Östlundh L, Ramaraj R, Javaid S, Govender RD, Ali BR, Patrinos GP, Young AH, et al. Effectiveness of pharmacogenomic tests including CYP2D6 and CYP2C19 genomic variants for guiding the treatment of depressive disorders: Systematic review and meta-analysis of randomised controlled trials. Neurosci Biobehav Rev. 2023;144:104965. doi:10.1016/j.neubiorev.2022.104965

234. Hou J, Wang C, Ma D, Chen Y, Jin H, An Y, Jia J, Huang L, Zhao H. The cardioprotective and anxiolytic effects of Chaihujialonggumuli granule on rats with anxiety after acute myocardial infarction is partly mediated by suppression of CXCR4/NF-κB/GSDMD pathway. Biomed Pharmacother. 2021;133:111015. doi:10.1016/j.biopha.2020.11101

235. Xu Z, Zhu X, Mu S, Fan R, Wang B, Gao W, Kang T. FTO overexpression expedites wound healing and alleviates depression in burn rats through facilitating keratinocyte migration and angiogenesis via mediating TFPI-2 demethylation. Mol Cell Biochem. 2023. doi:10.1007/s11010-023-04719-x

236. Peritore AF, Crupi R, Scuto M, Gugliandolo E, Siracusa R, Impellizzeri D, Cordaro M, D’amico R, Fusco R, Di Paola R, et al. The Role of Annexin A1 and Formyl Peptide Receptor 2/3 Signaling in Chronic Corticosterone-Induced Depression-Like behaviors and Impairment in Hippocampal-Dependent Memory. CNS Neurol Disord Drug Targets. 2020;19(1):27–43. doi:10.2174/1871527319666200107094732

237. DLSouza MS, Guisinger TC, Norman H, Seeley SL, Chrissobolis S. Regulator of G-protein signaling 5 protein protects against anxiety- and depression-like behavior. Behav Pharmacol. 2019;30(8):712–721. doi:10.1097/FBP.0000000000000506

238. Li J, Luo Y, Zhang R, Shi H, Zhu W, Shi J. Neuropeptide Trefoil Factor 3 Reverses Depressive-Like Behaviors by Activation of BDNF-ERK-CREB Signaling in Olfactory Bulbectomized Rats. Int J Mol Sci. 2015;16(12):28386–28400. doi:10.3390/ijms161226105

239. Sun Y, Wang Z, Hou J, Shi J, Tang Z, Wang C, Zhao H. huangxinfang Prevents S100A9-Induced Macrophage/Microglial Inflammation to Improve Cardiac Function and Depression-Like Behavior in Rats After Acute Myocardial Infarction. Front Pharmacol. 2022;13:832590. doi:10.3389/fphar.2022.832590

240. Lou QY, Li Z, Teng Y, Xie QM, Zhang M, Huang SW, Li WF, Chen YF, Pan FM, Xu SQ, et al. Associations of FKBP4 and FKBP5 gene polymorphisms with disease susceptibility, glucocorticoid efficacy, anxiety, depression, and health-related quality of life in systemic lupus erythematosus patients. Clin Rheumatol. 2021;40(1):167–179. doi:10.1007/s10067-020-05195-0

241. Zhang J, Li L, Liu Q, Zhao Z, Su D, Xiao C, Jin T, Chen L, Xu C, You Z, et al. Gastrodin programs an Arg-1+ microglial phenotype in hippocampus to ameliorate depression- and anxiety-like behaviors via the Nrf2 pathway in mice. Phytomedicine. 2023;113:154725. doi:10.1016/j.phymed.2023.154725

242. Durmaz CD, Karabulut HG, Saka MC, Sucularlı C, Gümüş Akay G, Atbaşoğlu C, Ilgın Ruhi H. Genetic Analysis of RASD1 as a Candidate Gene for Schizophrenia. Balkan Med J. 2022;39(6):422–428. doi:10.4274/balkanmedj.galenos.2022.2022-5-90

243. Cheng C, Chiu HJ, Lohel W, Chan CH, Hwu TM, Liu YR, Lan TH. Association of the ADRA1A gene and the severity of metabolic abnormalities in patients with schizophrenia. Prog Neuropsychopharmacol Biol Psychiatry. 2012;36(1):205–210. doi:10.1016/j.pnpbp.2011.10.011

244. Witt SH, Juraeva D, Sticht C, Strohmaier J, Meier S, Treutlein J, Dukal H, Frank J, Lang M, Deuschle M, et al. Investigation of manic and euthymic episodes identifies state- and trait-specific gene expression and STAB1 as a new candidate gene for bipolar disorder. Transl Psychiatry. 2014;4(8):e426. doi:10.1038/tp.2014.71

245. Huang X, Yu T, Li X, Cao Y, Li X, Liu B, Yang F, Li W, Zhao X, Feng G, et al. ABCB6, ABCB1 and ABCG1 genetic polymorphisms and antidepressant response of SSRIs in Chinese depressive patients. Pharmacogenomics. 2013;14(14):1723–1730. doi:10.2217/pgs.13.151

246. Lan B, Lv D, Sun X, Yang M, Zhang L, Ma F. Genetic Variations in IFNGR1, BDNF and IL-10 May Predict the Susceptibility to Depression and Anxiety in Chinese Women With Breast Cancer. Clin Breast Cancer. 2022;22(7):674–680. doi:10.1016/j.clbc.2022.07.002

247. Findikli E, Kurutas EB, Camkurt MA, Karaaslan MF, Izci F, Fındıklı HA, Kardaş S, Dag B, Altun H. Increased Serum G Protein-coupled Estrogen Receptor 1 Levels and Its Diagnostic Value in Drug Naïve Patients with Major Depressive Disorder. Clin Psychopharmacol Neurosci. 2017;15(4):337–342. doi:10.9758/cpn.2017.15.4.337

248. Ghosh M, Ali A, Joshi S, Srivastava AS, Tapadia MG. SLC1A3 C3590T but not BDNF G196A is a predisposition factor for stress as well as depression, in an adolescent eastern Indian population. BMC Med Genet. 2020;21(1):53. doi:10.1186/s12881-020-0993-6

249. Griffin JWD, Liu Y, Bradshaw PC, Wang K. In Silico Preliminary Association of Ammonia Metabolism Genes GLS, CPS1, and GLUL with Risk of Alzheimer’s Disease, Major Depressive Disorder, and Type 2 Diabetes. J Mol Neurosci. 2018;64(3):385–396. doi:10.1007/s12031-018-1035-0

250. Choi M, Wang SE, Ko SY, Kang HJ, Chae SY, Lee SH, Kim YS, Duman RS, Son H. Overexpression of human GATA-1 and GATA-2 interferes with spine formation and produces depressive behavior in rats. PLoS One. 2014;9(10):e109253. doi:10.1371/journal.pone.0109253

251. Xu K, Zheng P, Zhao S, Wang J, Feng J, Ren Y, Zhong Q, Zhang H, Chen X, Chen J, et al. LRFN5 and OLFM4 as novel potential biomarkers for major depressive disorder: a pilot study. Transl Psychiatry. 2023;13(1):188. doi:10.1038/s41398-023-02490-7

252. Sublette ME, Vaquero C, Baca-Garcia E, Pachano G, Huang YY, Oquendo MA, Mann JJ. Lack of association of SNPs from the FADS1-FADS2 gene cluster with major depression or suicidal behavior. Psychiatr Genet. 2016;26(2):81–86. doi:10.1097/YPG.0000000000000111

253. Paroni G, Panza F, De Cosmo S, Greco A, Seripa D, Mazzoccoli G. Klotho at the Edge of Alzheimer’s Disease and Senile Depression. Mol Neurobiol. 2019;56(3):1908–1920. doi:10.1007/s12035-018-1200-z

254. Peritore AF, Crupi R, Scuto M, Gugliandolo E, Siracusa R, Impellizzeri D, Cordaro M, D’amico R, Fusco R, Di Paola R, et al. The Role of Annexin A1 and Formyl Peptide Receptor 2/3 Signaling in Chronic Corticosterone-Induced Depression-Like behaviors and Impairment in Hippocampal-Dependent Memory. CNS Neurol Disord Drug Targets. 2020;19(1):27–43. doi:10.2174/1871527319666200107094732

255. Williams AJ, Yee P, Smith MC, Murphy GG, Umemori H. Deletion of fibroblast growth factor 22 (FGF22) causes a depression-like phenotype in adult mice. Behav Brain Res. 2016;307:11–17. doi:10.1016/j.bbr.2016.03.047

256. Baranyi A, Enko D, Meinitzer A, Von Lewinski D, Rothenhäusler HB, Harpf L, Traninger H, Obermayer-Pietsch B, Harb BM, Schweinzer M, et al. Myeloperoxidase as a Potential Biomarker of Acute-Myocardial-Infarction-Induced Depression and Suppression of the Innate Immune System. Antioxidants (Basel). 2022;11(11):2083. doi:10.3390/antiox11112083

257. Naoi M, Maruyama W, Shamoto-Nagai M. Type A monoamine oxidase and serotonin are coordinately involved in depressive disorders: from neurotransmitter imbalance to impaired neurogenesis. J Neural Transm (Vienna). 2018;125(1):53–66. doi:10.1007/s00702-017-1709-8

258. Kobayashi M, Nakatani T, Koda T, Matsumoto K, Ozaki R, Mochida N, Takao K, Miyakawa T, Matsuoka I. Absence of BRINP1 in mice causes increase of hippocampal neurogenesis and behavioral alterations relevant to human psychiatric disorders. Mol Brain. 2014;7:12. doi:10.1186/1756-6606-7-12

259. Zhou R, Lu Y, Han Y, Li X, Lou H, Zhu L, Zhen X, Duan S. Mice heterozygous for cathepsin D deficiency exhibit mania-related behavior and stress-induced depression. Prog Neuropsychopharmacol Biol Psychiatry. 2015;63:110–118. doi:10.1016/j.pnpbp.2015.06.007

260. Wang X, Xie Y, Zhang T, Bo S, Bai X, Liu H, Li T, Liu S, Zhou Y, Cong X, et al. Resveratrol reverses chronic restraint stress-induced depression-like behaviour: Involvement of BDNF level, ERK phosphorylation and expression of Bcl-2 and Bax in rats. Brain Res Bull. 2016;125:134–143. doi:10.1016/j.brainresbull.2016.06.014

261. Sublette ME, Vaquero C, Baca-Garcia E, Pachano G, Huang YY, Oquendo MA, Mann JJ. Lack of association of SNPs from the FADS1-FADS2 gene cluster with major depression or suicidal behavior. Psychiatr Genet. 2016;26(2):81–86. doi:10.1097/YPG.0000000000000111

262. Wang L, Shi C, Zhang K, Xu Q. The gender-specific association of EHD3 polymorphisms with major depressive disorder. Neurosci Lett. 2014;567:11–14. doi:10.1016/j.neulet.2014.02.055

263. Ozeki Y, Pickard BS, Kano S, Malloy MP, Zeledon M, Sun DQ, Fujii K, Wakui K, Shirayama Y, Fukushima Y, et al. A novel balanced chromosomal translocation found in subjects with schizophrenia and schizotypal personality disorder: altered l-serine level associated with disruption of PSAT1 gene expression. Neurosci Res. 2011;69(2):154–160. doi:10.1016/j.neures.2010.10.003

264. Lin XM, Luo W, Wang H, Li RZ, Huang YS, Chen LK, Wu XP. The Role of Prostaglandin-Endoperoxide Synthase-2 in Chemoresistance of Non-Small Cell Lung Cancer. Front Pharmacol. 2019;10:836. doi:10.3389/fphar.2019.00836

265. Zhang L, Martin G, Mohankumar K, Hampton JT, Liu WR, Safe S. Resveratrol binds nuclear receptor 4a1 (nr4a1) and acts as an nr4a1 antagonist in lung cancer cells. Mol Pharmacol. 2022. doi:10.1124/molpharm.121.000481

266. Chien MH, Lee WJ, Yang YC, Li YL, Chen BR, Cheng TY, Yang PW, Wang MY, Jan YH, Lin YK, et al. KSRP suppresses cell invasion and metastasis through miR-23a-mediated EGR3 mRNA degradation in non-small cell lung cancer. Biochim Biophys Acta Gene Regul Mech. 2017;1860(10):1013–1024. doi:10.1016/j.bbagrm.2017.08.005

267. Zhang Y, Chen Z, Jiang A, Gao G. KLRK1 as a prognostic biomarker for lung adenocarcinoma cancer. Sci Rep. 2022;12(1):1976. doi:10.1038/s41598-022-05997-z

268. Ke W, Zhang L, Dai Y. The role of IL-6 in immunotherapy of non-small cell lung cancer (NSCLC) with immune-related adverse events (irAEs). Thorac Cancer. 2020;11(4):835–839. doi:10.1111/1759-7714.13341

269. Gu L, Yao Y, Chen Z. An inter-correlation among chemokine (C-X-C motif) ligand (CXCL) 1, CXCL2 and CXCL8, and their diversified potential as biomarkers for tumor features and survival profiles in non-small cell lung cancer patients. Transl Cancer Res. 2021;10(2):748–758. doi:10.21037/tcr-20-2539

270. Kiziltunc Ozmen H, Simsek M. Serum IL-23, E-selectin and sICAM levels in non-small cell lung cancer patients before and after radiotherapy. J Int Med Res. 2020;48(5):300060520923493. doi:10.1177/0300060520923493

271. Zhu X, Liu D, Wang Y, Dong M. Salidroside suppresses nonsmall cell lung cancer cells proliferation and migration via microRNA-103-3p/Mzb1. Anticancer Drugs. 2020;31(7):663–671. doi:10.1097/CAD.0000000000000926

272. Hu ZW, Sun W, Wen YH, Ma RQ, Chen L, Chen WQ, Lei WB, Wen WP. CD69 and SBK1 as potential predictors of responses to PD-1/PD-L1 blockade cancer immunotherapy in lung cancer and melanoma. Front Immunol. 2022;13:952059. doi:10.3389/fimmu.2022.952059

273. Zhang T, Qiu L, Cao J, Li Q, Zhang L, An G, Ni J, Jia H, Li S, Li K. ZFP36 loss-mediated BARX1 stabilization promotes malignant phenotypes by transactivating master oncogenes in NSCLC. Cell Death Dis. 2023;14(8):527. doi:10.1038/s41419-023-06044-z

274. Yu S, Yi M, Xu L, Qin S, Li A, Wu K. CXCL1 as an Unfavorable Prognosis Factor Negatively Regulated by DACH1 in Non-small Cell Lung Cancer. Front Oncol. 2020;9:1515. doi:10.3389/fonc.2019.01515

275. Abuduwaili K, Zhu X, Shen Y, Lu S, Liu C. circ_0008797 attenuates non-small cell lung cancer proliferation, metastasis, and aerobic glycolysis by sponging miR-301a-3p/SOCS2. Environ Toxicol. 2022;37(7):1697–1710. doi:10.1002/tox.23518

276. Son B, Jeon J, Lee S, Kim H, Kang H, Youn H, Jo S, Youn B. Radiotherapy in combination with hyperthermia suppresses lung cancer progression via increased NR4A3 and KLF11 expression. Int J Radiat Biol. 2019;95(12):1696–1707. doi:10.1080/09553002.2019.1665213

277. Wang L, Liu X, Ren Y, Zhang J, Chen J, Zhou W, Guo W, Wang X, Chen H, Li M, et al. Cisplatin-enriching cancer stem cells confer multidrug resistance in non-small cell lung cancer via enhancing TRIB1/HDAC activity. Cell Death Dis. 2017;8(4):e2746. doi:10.1038/cddis.2016.409

278. Sun Y, Gao Y, Dong M, Li J, Li X, He N, Song H, Zhang M, Ji K, Wang J, et al. Kremen2 drives the progression of non-small cell lung cancer by preventing SOCS3-mediated degradation of EGFR. J Exp Clin Cancer Res. 2023;42(1):140. doi:10.1186/s13046-023-02692-3

279. Song Q, Shang J, Zhang C, Chen J, Zhang L, Wu X. Transcription factor RUNX3 promotes CD8+ T cell recruitment by CCL3 and CCL20 in lung adenocarcinoma immune microenvironment. J Cell Biochem. 2020;121(5-6):3208–3220. doi:10.1002/jcb.29587

280. Melese ES, Franks E, Cederberg RA, Harbourne BT, Shi R, Wadsworth BJ, Collier JL, Halvorsen EC, Johnson F, Luu J, et al. CCL5 production in lung cancer cells leads to an altered immune microenvironment and promotes tumor development. Oncoimmunology. 2021;11(1):2010905.doi:10.1080/2162402X.2021.2010905

281. Zhou Z, Zhou Q, Wu X, Xu S, Hu X, Tao X, Li B, Peng J, Li D, Shen L, et al. VCAM-1 secreted from cancer-associated fibroblasts enhances the growth and invasion of lung cancer cells through AKT and MAPK signaling. Cancer Lett. 2020;473:62–73. doi:10.1016/j.canlet.2019.12.039

282. Li H, Harrison EB, Li H, Hirabayashi K, Chen J, Li QX, Gunn J, Weiss J, Savoldo B, Parker JS, et al. Targeting brain lesions of non-small cell lung cancer by enhancing CCL2-mediated CAR-T cell migration. Nat Commun. 2022;13(1):2154. doi:10.1038/s41467-022-29647-0

283. Yang W, Wei C, Cheng J, Ding R, Li Y, Wang Y, Yang Y, Wang J. BTG2 and SerpinB5, a novel gene pair to evaluate the prognosis of lung adenocarcinoma. Front Immunol. 2023;14:1098700. doi:10.3389/fimmu.2023.1098700

284. Ito T, Ozaki S, Chanasong R, Mizutani Y, Oyama T, Sakurai H, Matsumoto I, Takemura H, Kawahara E. Activation of ERK/IER3/PP2A-B56γ-positive feedback loop in lung adenocarcinoma by allelic deletion of B56γ gene. Oncol Rep. 2016;35(5):2635–2642. doi:10.3892/or.2016.4677

285. Zhang X, Lu Y, Huang K, Pan Q, Jia Y, Cui B, Yin P, Li J, Ju J, Fan X, et al. The synergized diagnostic value of VTQ with chemokine CXCL13 in lung tumors. Front Oncol. 2023;13:1115485. doi:10.3389/fonc.2023.1115485

286. Gan X, Liu Z, Tong B, Zhou J. Epigenetic downregulated ITGBL1 promotes non-small cell lung cancer cell invasion through Wnt/PCP signaling. Tumour Biol. 2016;37(2):1663–1669. doi:10.1007/s13277-015-3919-8

287. Majores M, Schindler A, Fuchs A, Stein J, Heukamp L, Altevogt P, Kristiansen G. Membranous CD24 expression as detected by the monoclonal antibody SWA11 is a prognostic marker in non-small cell lung cancer patients. BMC Clin Pathol. 2015;15:19. doi:10.1186/s12907-015-0019-z

288. Simoncello F, Piperno GM, Caronni N, Amadio R, Cappelletto A, Canarutto G, Piazza S, Bicciato S, Benvenuti F. CXCL5-mediated accumulation of mature neutrophils in lung cancer tissues impairs the differentiation program of anticancer CD8 T cells and limits the efficacy of checkpoint inhibitors. Oncoimmunology. 2022;11(1):2059876. doi:10.1080/2162402X.2022.2059876

289. Wang Z, Sun J, Feng Y, Tian X, Wang B, Zhou Y. Oncogenic roles and drug target of CXCR4/CXCL12 axis in lung cancer and cancer stem cell. Tumour Biol. 2016;37(7):8515–8528. doi:10.1007/s13277-016-5016-z

290. Wilson IM, Vucic EA, Enfield KS, Thu KL, Zhang YA, Chari R, Lockwood WW, Radulovich N, Starczynowski DT, Banáth JP, et al. EYA4 is inactivated biallelically at a high frequency in sporadic lung cancer and is associated with familial lung cancer risk. Oncogene. 2014;33(36):4464–4473. doi:10.1038/onc.2013.396

291. Zhang W, Gao Z, Zeng G, Xie H, Liu J, Liu N, Wang G. Clinical significance of urinary plasminogen and fibrinogen gamma chain as novel potential diagnostic markers for non-small-cell lung cancer. Clin Chim Acta. 2020;502:55–65. doi:10.1016/j.cca.2019.11.022

292. Zhou Y, Yan J, Chen H, Zhou W, Xiao G, Zou H, Yang J.MiR-4652-5p Targets RND1 to Regulate Cell Adhesion and Promote Lung Squamous Cell Carcinoma Progression. Appl Biochem Biotechnol. 2022;194(7):3031–3043. doi:10.1007/s12010-022-03897-6

293. Ning Y, Zheng H, Yang Y, Zang H, Wang W, Zhan Y, Wang H, Luo J, Wen Q, Peng J, et al. YAP1 synergize with YY1 transcriptional co-repress DUSP1 to induce osimertinib resistant by activating the EGFR/MAPK pathway and abrogating autophagy in non-small cell lung cancer. Int J Biol Sci. 2023;19(8):2458–2474. doi:10.7150/ijbs.79965

294. Liu Q, Qiao M, Lohinai Z, Mao S, Pan Y, Wang Y, Yang S, Zhou F, Jiang T, Yi X, et al. CCL19 associates with lymph node metastasis and inferior prognosis in patients with small cell lung cancer. Lung Cancer. 2021;162:194–202. doi:10.1016/j.lungcan.2021.11.003

295. Lv Y, Liu Z, Xiong K, Duan H, Yang J, Liao P. GADD45B predicts lung squamous cell carcinoma survival and impacts immune infiltration, and T cell exhaustion. Autoimmunity. 2023;56(1):2209706. doi:10.1080/08916934.2023.2209706

296. Lu J, Shi Q, Zhang L, Wu J, Lou Y, Qian J, Zhang B, Wang S, Wang H, Zhao X, et al. Integrated Transcriptome Analysis Reveals KLK5 and L1CAM Predict Response to Anlotinib in NSCLC at 3rd Line. Front Oncol. 2019;9:886. doi:10.3389/fonc.2019.00886

297. Prado-Garcia H, Aguilar-Cazares D, Meneses-Flores M, Morales-Fuentes J, Lopez-Gonzalez JS. Lung carcinomas do not induce T-cell apoptosis via the Fas/Fas ligand pathway but down-regulate CD3 epsilon expression. Cancer Immunol Immunother. 2008;57(3):325–336. doi:10.1007/s00262-007-0372-6

298. Seder CW, Hartojo W, Lin L, Silvers AL, Wang Z, Thomas DG, Giordano TJ, Chen G, Chang AC, Orringer MB, et al. Upregulated INHBA expression may promote cell proliferation and is associated with poor survival in lung adenocarcinoma. Neoplasia. 2009;11(4):388–396. doi:10.1593/neo.81582

299. Li J, Tang Z, Wang H, Wu W, Zhou F, Ke H, Lu W, Zhang S, Zhang Y, Yang S, et al. CXCL6 promotes non-small cell lung cancer cell survival and metastasis via down-regulation of miR-515-5p. Biomed Pharmacother. 2018;97:1182–1188. doi:10.1016/j.biopha.2017.11.004

300. Yin J, Wang C, Vogel U, Ma Y, Zhang Y, Wang H, Sun Z, Du S. Common variants of pro-inflammatory gene IL1B and interactions with PPP1R13L and POLR1G in relation to lung cancer among Northeast Chinese. Sci Rep. 2023;13(1):7352. doi:10.1038/s41598-023-34069-z

301. Taniguchi H, Takeuchi S, Fukuda K, Nakagawa T, Arai S, Nanjo S, Yamada T, Yamaguchi H, Mukae H, Yano S. Amphiregulin triggered epidermal growth factor receptor activation confers in vivo crizotinib-resistance of EML4-ALK lung cancer and circumvention by epidermal growth factor receptor inhibitors. Cancer Sci. 2017;108(1):53–60. doi:10.1111/cas.13111

302. Wang F, Gu T, Chen Y, Chen Y, Xiong D, Zhu Y. Long non-coding RNA SOX21-AS1 modulates lung cancer progress upon microRNA miR-24-3p/PIM2 axis. Bioengineered. 2021;12(1):6724–6737. doi:10.1080/21655979.2021.1955578

303. Ou J, Liao Q, Du Y, Xi W, Meng Q, Li K, Cai Q, Pang CLK. SERPINE1 and SERPINB7 as potential biomarkers for intravenous vitamin C treatment in non-small-cell lung cancer. Free Radic Biol Med. 2023;209(Pt 1):96–107. doi:10.1016/j.freeradbiomed.2023.10.391

304. Pan CM, Wang ML, Chiou SH, Chen HY, Wu CW. Oncostatin M suppresses metastasis of lung adenocarcinoma by inhibiting SLUG expression through coordination of STATs and PIASs signalings. Oncotarget. 2016;7(37):60395–60406. doi:10.18632/oncotarget.10939

305. Chang PM, Yeh YC, Chen TC, Wu YC, Lu PJ, Cheng HC, Lu HJ, Chen MH, Chou TY, Huang CY. High expression of CHRNA1 is associated with reduced survival in early stage lung adenocarcinoma after complete resection. Ann Surg Oncol. 2013;20(11):3648–3654. doi:10.1245/s10434-013-3034-2

306. Lin S, Zhang X, Huang G, Cheng L, Lv J, Zheng D, Lin S, Wang S, Wu Q, Long Y, et al. Myeloid-derived suppressor cells promote lung cancer metastasis by CCL11 to activate ERK and AKT signaling and induce epithelial-mesenchymal transition in tumor cells. Oncogene. 2021;40(8):1476–1489. doi:10.1038/s41388-020-01605-4

307. Feng YY, Liu CH, Xue Y, Chen YY, Wang YL, Wu XZ. MicroRNA-147b promotes lung adenocarcinoma cell aggressiveness through negatively regulating microfibril-associated glycoprotein 4 (MFAP4) and affects prognosis of lung adenocarcinoma patients. Gene. 2020;730:144316. doi:10.1016/j.gene.2019.144316

308. Li X, Min S, Wang H, Shen Y, Li W, Chen Y, Wang X. MLF1 protein is a potential therapy target for lung adenocarcinoma. Int J Clin Exp Pathol. 2018;11(7):3533–3541.

309. Zhang J, Xu L, Gao J, Li J, Zhao X, Yang P, Ge Y, Guo D, Liu Z, Wang X, et al. Leukocyte CH25H is a potential diagnostic and prognostic marker for lung adenocarcinoma. Sci Rep. 2022;12(1):22201. doi:10.1038/s41598-022-24183-9

310. Ye Q, Singh S, Qian PR, Guo NL. Immune-Omics Networks of CD27, PD1, and PDL1 in Non-Small Cell Lung Cancer. Cancers (Basel). 2021;13(17):4296. doi:10.3390/cancers13174296

311. Kuang M, Zhou Z, Lu Z, Shen W, Ge H, Tao X, Zhao Y, Zhuge L, Sun Y, Ji D, et al. Prognostic prediction of lung adenocarcinoma by integrative analysis of RHOH expression and methylation. Clin Respir J. 2023;17(3):148–156. doi:10.1111/crj.13574

312. Moreno-Manuel A, Jantus-Lewintre E, Simões I, Aranda F, Calabuig-Fariñas S, Carreras E, Zúñiga S, Saenger Y, Rosell R, Camps C, et al. CD5 and CD6 as immunoregulatory biomarkers in non-small cell lung cancer. Transl Lung Cancer Res. 2020;9(4):1074–1083. doi:10.21037/tlcr-19-445

313. Shen Q, Wang H, Zhang L. TP63 Functions as a Tumor Suppressor Regulated by GAS5/miR-221-3p Signaling Axis in Human Non-Small Cell Lung Cancer Cells. Cancer Manag Res. 2023;15:217–231. doi:10.2147/CMAR.S387781

314. Roncato F, Regev O, Feigelson SW, Yadav SK, Kaczmarczyk L, Levi N, Drago-Garcia D, Ovadia S, Kizner M, Addadi Y, et al. Reduced Lamin A/C Does Not Facilitate Cancer Cell Transendothelial Migration but Compromises Lung Metastasis. Cancers (Basel). 2021;13(10):2383. doi:10.3390/cancers13102383

315. Ma K, Qiao Y, Wang H, Wang S. Comparative expression analysis of PD-1, PD-L1, and CD8A in lung adenocarcinoma. Ann Transl Med. 2020;8(22):1478. doi:10.21037/atm-20-6486

316. Xie J, Zhang Y, Hu X, Lv R, Xiao D, Jiang L, Bao Q. Norcantharidin inhibits Wnt signal pathway via promoter demethylation of WIF-1 in human non-small cell lung cancer. Med Oncol. 2015;32(5):145. doi:10.1007/s12032-015-0592-0

317. Zhou H, Guan Q, Hou X, Liu L, Zhou L, Li W, Liu H. Epithelial-mesenchymal reprogramming by KLF4-regulated Rictor expression contributes to metastasis of non-small cell lung cancer cells. Int J Biol Sci. 2022;18(13):4869–4883.. doi:10.7150/ijbs.73548

318. Yigit M, Değirmencioğlu S, Ugurlu E, Yaren A. Effect of serum interleukin-1 receptor antagonist level on survival of patients with non-small cell lung cancer. Mol Clin Oncol. 2017;6(5):708–712. doi:10.3892/mco.2017.1195

319. Song M, Ping Y, Zhang K, Yang L, Li F, Zhang C, Cheng S, Yue D, Maimela NR, Qu J, et al. Low-Dose IFNγ Induces Tumor Cell Stemness in Tumor Microenvironment of Non-Small Cell Lung Cancer. Cancer Res. 2019;79(14):3737–3748. doi:10.1158/0008-5472.CAN-19-0596

320. Wright CM, Savarimuthu Francis SM, Tan ME, Martins MU, Winterford C, Davidson MR, Duhig EE, Clarke BE, Hayward NK, et al. MS4A1 dysregulation in asbestos-related lung squamous cell carcinoma is due to CD20 stromal lymphocyte expression. PLoS One. 2012;7(4):e34943. doi:10.1371/journal.pone.0034943

321. Pan B, Wan L, Li Y, Yang J, Chen Z, Tong X, Zhao J, Li C. Comprehensive pan-cancer analysis identifies FHL2 associated with poor prognosis in lung adenocarcinoma. Transl Cancer Res. 2023;12(6):1516–1534. doi:10.21037/tcr-22-2786

322. Liu Y, Ma H, Dong T, Yan Y, Sun L, Wang W. Clinical significance of expression level of CX3CL1-CX3CR1 axis in bone metastasis of lung cancer. Clin Transl Oncol. 2021;23(2):378–388. doi:10.1007/s12094-020-02431-6

323. Wu CK, Wei MT, Wu HC, Wu CL, Wu CJ, Liaw H, Su WP. BMP2 promotes lung adenocarcinoma metastasis through BMP receptor 2-mediated SMAD1/5 activation. Sci Rep. 2022;12(1):16310. doi:10.1038/s41598-022-20788-2

324. Yu L, Chen D, Song J. LncRNA SNHG16 promotes non-small cell lung cancer development through regulating EphA2 expression by sponging miR-520a-3p. Thorac Cancer. 2020;11(3):603–611. doi:10.1111/1759-7714.13304

325. Jeong SB, Das R, Kim DH, Lee S, Oh HI, Jo S, Lee Y, Kim J, Park S, Choi DK, et al. Anticancer effect of verteporfin on non-small cell lung cancer via downregulation of ANO1. Biomed Pharmacother. 2022;153:113373. doi:10.1016/j.biopha.2022.113373

326. Yang C, Gagnon C, Hou X, Hardy P. Low density lipoprotein receptor mediates anti-VEGF effect of lymphocyte T-derived microparticles in Lewis lung carcinoma cells. Cancer Biol Ther. 2010;10(5):448–456. doi:10.4161/cbt.10.5.12533

327. Bi G, Liang J, Zhao M, Zhang H, Jin X, Lu T, Zheng Y, Bian Y, Chen Z, Huang Y, et al. miR-6077 promotes cisplatin/pemetrexed resistance in lung adenocarcinoma via CDKN1A/cell cycle arrest and KEAP1/ferroptosis pathways. Mol Ther Nucleic Acids. 2022;28:366–386. doi:10.1016/j.omtn.2022.03.020

328. Yue Z, Ningning D, Lin Y, Jianming Y, Hongtu Z, Ligong Y, Feng L, Shuaibo W, Yousheng M. Correlation between CXCR4, CXCR5 and CCR7 expression and survival outcomes in patients with clinical T1N0M0 non-small cell lung cancer. Thorac Cancer. 2020;11(10):2955–2965. doi:10.1111/1759-7714.13645

329. Xuan X, Wang Z, Wang Y. Circ_0058608 contributes to the progression and taxol resistance of non-small cell lung cancer by sponging miR-1299 to upregulate GBP1. Anticancer Drugs. 2023;34(1):103–114. doi:10.1097/CAD.0000000000001346

330. Deng T, Lin D, Zhang M, Zhao Q, Li W, Zhong B, Deng Y, Fu X. Differential expression of bone morphogenetic protein 5 in human lung squamous cell carcinoma and adenocarcinoma. Acta Biochim Biophys Sin (Shanghai). 2015;47(7):557–563. doi:10.1093/abbs/gmv037

331. Bach DH, Luu TT, Kim D, An YJ, Park S, Park HJ, Lee SK. BMP4 Upregulation Is Associated with Acquired Drug Resistance and Fatty Acid Metabolism in EGFR-Mutant Non-Small-Cell Lung Cancer Cells. Mol Ther Nucleic Acids. 2018;12:817–828. doi:10.1016/j.omtn.2018.07.016

332. Kahm YJ, Jung U, Kim RK. Regulation of Cancer Stem Cells and Epithelial-Mesenchymal Transition by CTNNAL1 in Lung Cancer and Glioblastoma. Biomedicines. 2023;11(5):1462. doi:10.3390/biomedicines11051462

333. Zhao J, Feng M, Liu D, Liu H, Shi M, Zhang J, Qu J.Antagonism between HTRA3 and TGFβ1 Contributes to Metastasis in Non-Small Cell Lung Cancer. Cancer Res. 2019;79(11):2853–2864. doi:10.1158/0008-5472.CAN-18-2507

334. Bai X, Guo ZQ, Zhang YP, Fan ZZ, Liu LJ, Liu L, Long LL, Ma SC, Wang J, Fang Y, et al. CDK4/6 inhibition triggers ICAM1-driven immune response and sensitizes LKB1 mutant lung cancer to immunotherapy. Nat Commun. 2023;14(1):1247. doi:10.1038/s41467-023-36892-4

335. Zhang X, Wu Z, Ma K. SNCA correlates with immune infiltration and serves as a prognostic biomarker in lung adenocarcinoma. BMC Cancer. 2022;22(1):406. doi:10.1186/s12885-022-09289-7

336. Li S, Yang S, Qiu C, Sun D. LncRNA MSC-AS1 facilitates lung adenocarcinoma through sponging miR-33b-5p to up-regulate GPAM. Biochem Cell Biol. 2021;99(2):241–248. doi:10.1139/bcb-2020-0239

337. Wu JE, Wu YY, Tung CH, Tsai YT, Chen HY, Chen YL, Hong TM. DNA methylation maintains the CLDN1-EPHB6-SLUG axis to enhance chemotherapeutic efficacy and inhibit lung cancer progression. Theranostics. 2020;10(19):8903–8923. doi:10.7150/thno.45785

338. Aida R, Hagiwara K, Okano K, Nakata K, Obata Y, Yamashita T, Yoshida K, Hagiwara H. miR-34a-5p might have an important role for inducing apoptosis by down-regulation of SNAI1 in apigenin-treated lung cancer cells. Mol Biol Rep. 2021;48(3):2291–2297. doi:10.1007/s11033-021-06255-7

339. Bing Z, Jian-ru Y, Yao-quan J, Shi-feng C. Evaluation of angiogenesis in non-small cell lung carcinoma by CD34 immunohistochemistry. Cell Biochem Biophys. 2014;70(1):327–331. doi:10.1007/s12013-014-9916-5

340. Cui Z, Li D, Zhao J, Chen K. Falnidamol and cisplatin combinational treatment inhibits non-small cell lung cancer (NSCLC) by targeting DUSP26-mediated signal pathways. Free Radic Biol Med. 2022;183:106–124. doi:10.1016/j.freeradbiomed.2022.03.003

341. Yang K, Tian C, Zhang C, Xiang M. The Controversial Role of IL-33 in Lung Cancer. Front Immunol. 2022;13:897356. doi:10.3389/fimmu.2022.897356

342. Chen LM, Niu YD, Xiao M, Li XJ, Lin H. LncRNA NEAT1 regulated cell proliferation, invasion, migration and apoptosis by targeting has-miR-376b-3p/SULF1 axis in non-small cell lung cancer. Eur Rev Med Pharmacol Sci. 2020;24(9):4810–4821. doi:10.26355/eurrev_202005_21170

343. Sun P, Sun L, Cui J, Liu L, He Q. Long noncoding RNA HAS2-AS1 accelerates non-small cell lung cancer chemotherapy resistance by targeting LSD1/EphB3 pathway. Am J Transl Res. 2020;12(3):950–958.

344. Anantharajan J, Zhou H, Zhang L, Hotz T, Vincent MY, Blevins MA, Jansson AE, Kuan JWL, Ng EY, Yeo YK, et al. Structural and Functional Analyses of an Allosteric EYA2 Phosphatase Inhibitor That Has On-Target Effects in Human Lung Cancer Cells. Mol Cancer Ther. 2019;18(9):1484–1496. doi:10.1158/1535-7163.MCT-18-1239

345. Bonnet-Magnaval F, Diallo LH, Brunchault V, Laugero N, Morfoisse F, David F, Roussel E, Nougue M, Zamora A, Marchaud E, et al. High Level of Staufen1 Expression Confers Longer Recurrence Free Survival to Non-Small Cell Lung Cancer Patients by Promoting THBS1 mRNA Degradation. Int J Mol Sci. 2021;23(1):215. doi:10.3390/ijms23010215

346. Shimizu M, Kondo M, Ito Y, Kume H, Suzuki R, Yamaki K. Soluble Fas and Fas ligand provide new information on metastasis and response to chemotherapy in SCLC patients. Cancer Detect Prev. 2005;29(2):175–180. doi:10.1016/j.cdp.2004.09.001

347. Kuo IY, Liu D, Lai WW, Wang YC, Loh YP. Carboxypeptidase E mRNA: Overexpression predicts recurrence and death in lung adenocarcinoma cancer patients. Cancer Biomark. 2022;33(3):369–377. doi:10.3233/CBM-210206

348. Guo Q, Wang SH, Ji YM, Tong S, Li D, Ding XC, Wu CY. The Roles and Mechanisms of TRAT1 in the Progression of Non-Small Cell Lung Cancer. Curr Med Sci. 2022;42(6):1186–1200. doi:10.1007/s11596-022-2625-1

349. Lesbon JCC, Garnica TK, Xavier PLP, Rochetti AL, Reis RM, Müller S, Fukumasu H. A Screening of Epigenetic Therapeutic Targets for Non-Small Cell Lung Cancer Reveals PADI4 and KDM6B as Promising Candidates. Int J Mol Sci. 2022;23(19):11911. doi:10.3390/ijms231911911

350. Latina A, Viticchiè G, Lena AM, Piro MC, Annicchiarico-Petruzzelli M, Melino G, Candi E. ΔNp63 targets cytoglobin to inhibit oxidative stress-induced apoptosis in keratinocytes and lung cancer. Oncogene. 2016;35(12):1493–1503. doi:10.1038/onc.2015.222

351. Tang Y, Hu C, Yang H, Cao L, Li Y, Deng P, Huang L. Rnd3 regulates lung cancer cell proliferation through notch signaling. PLoS One. 2014;9(11):e111897. doi:10.1371/journal.pone.0111897

352. Lin L, Lin G, Lin H, Chen L, Chen X, Lin Q, Xu Y, Zeng Y. Integrated profiling of endoplasmic reticulum stress-related DERL3 in the prognostic and immune features of lung adenocarcinoma. Front Immunol. 2022;13:906420. doi:10.3389/fimmu.2022.906420

353. Gong K, Guo G, Beckley N, Zhang Y, Yang X, Sharma M, Habib AA. Tumor necrosis factor in lung cancer: Complex roles in biology and resistance to treatment. Neoplasia. 2021;23(2):189–196. doi:10.1016/j.neo.2020.12.006

354. Duan L, Pang HL, Chen WJ, Shen WW, Cao PP, Wang SM, Liu LL, Zhang HL. The role of GDF15 in bone metastasis of lung adenocarcinoma cells. Oncol Rep. 2019;41(4):2379–2388. doi:10.3892/or.2019.7024

355. Liang H, Peng J. LncRNA HOTAIR promotes proliferation, invasion and migration in NSCLC cells via the CCL22 signaling pathway. PLoS One. 2022;17(2):e0263997. doi:10.1371/journal.pone.0263997

356. Liu ZD, Mou ZX, Che XH, Wang K, Li HX, Chen XY, Guo XM. ARHGAP15 regulates lung cancer cell proliferation and metastasis via the STAT3 pathway. Eur Rev Med Pharmacol Sci. 2019;23(13):5840–5850. doi:10.26355/eurrev_201907_18326

357. Hu J, Wang L, Guan C. MiR-532-5p Suppresses Migration and Invasion of Lung Cancer Cells Through Inhibiting CCR4. Cancer Biother Radiopharm. 2020;35(9):673–681. doi:10.1089/cbr.2019.3258

358. Chaszczewska-Markowska M, Kosacka M, Chryplewicz A, Dyła T, Brzecka A, Bogunia-Kubik K. ECCR1 and NFKB2 Polymorphisms as Potential Biomarkers of Non-small Cell Lung Cancer in a Polish Population. Anticancer Res. 2019;39(6):3269–3272. doi:10.21873/anticanres.13469

359. Yang R, Yang M, Wu Z, Liu B, Zheng M, Lu L, Wu S. Tespa1 deficiency reduces the antitumour immune response by decreasing CD8+T cell activity in a mouse Lewis lung cancer model. Int Immunopharmacol. 2023;124(Pt A):110865. doi:10.1016/j.intimp.2023.110865

360. Mu X, Li H, Zhou L, Xu W. TRIM52 regulates the proliferation and invasiveness of lung cancer cells via the Wnt/βLcatenin pathway. Oncol Rep. 2019;41(6):3325–3334. doi:10.3892/or.2019.7110

361. Al Abdulmonem W, Rasheed Z, Aljohani ASM, Omran OM, Rasheed N, Alkhamiss A, A M Al Salloom A, Alhumaydhi F, Alblihed MA, Al Ssadh H, et al. Absence of CD74 Isoform at 41kDa Prevents the Heterotypic Associations between CD74 and CD44 in Human Lung Adenocarcinoma-derived Cells. Immunol Invest. 2021;50(8):891–905. doi:10.1080/08820139.2020.1790594

362. Moreno-Manuel A, Jantus-Lewintre E, Simões I, Aranda F, Calabuig-Fariñas S, Carreras E, Zúñiga S, Saenger Y, Rosell R, Camps C, et al. CD5 and CD6 as immunoregulatory biomarkers in non-small cell lung cancer. Transl Lung Cancer Res. 2020;9(4):1074–1083. doi:10.21037/tlcr-19-445

363. Miyata T, Oyama T, Yoshimatsu T, Higa H, Kawano D, Sekimura A, Yamashita N, So T, Gotoh A. The Clinical Significance of Cancer Stem Cell Markers ALDH1A1 and CD133 in Lung Adenocarcinoma. Anticancer Res. 2017;37(5):2541–2547. doi:10.21873/anticanres.11597

364. Horton BL, D’Souza AD, Zagorulya M, McCreery CV, Abhiraman GC, Picton L, Sheen A, Agarwal Y, Momin N, Wittrup KD, et al. Overcoming lung cancer immunotherapy resistance by combining nontoxic variants of IL-12 and IL-2. JCI Insight. 2023;8(19):e172728. doi:10.1172/jci.insight.172728

365. Karaki S, Blanc C, Tran T, Galy-Fauroux I, Mougel A, Dransart E, Anson M, Tanchot C, Paolini L, Gruel N, et al. CXCR6 deficiency impairs cancer vaccine efficacy and CD8+ resident memory T-cell recruitment in head and neck and lung tumors. J Immunother Cancer. 2021;9(3):e001948. doi:10.1136/jitc-2020-001948

366. Nozaki K, Kagamu H, Shoji S, Igarashi N, Ohtsubo A, Okajima M, Miura S, Watanabe S, Yoshizawa H, Narita I. DDX3X induces primary EGFR-TKI resistance based on intratumor heterogeneity in lung cancer cells harboring EGFR-activating mutations. PLoS One. 2014;9(10):e111019. doi:10.1371/journal.pone.0111019

367. Zhou H, Chang J, Zhang J, Zheng H, Miao X, Mo H, Sun J, Jia Q, Qi G. PRMT5 activates KLF5 by methylation to facilitate lung cancer. J Cell Mol Med. 2023. doi:10.1111/jcmm.17856

368. Wang J, Xu Y, Rao X, Zhang R, Tang J, Zhang D, Jie X, Zhu K, Wang X, Xu Y, et al. BRD4-IRF1 axis regulates chemoradiotherapy-induced PD-L1 expression and immune evasion in non-small cell lung cancer. Clin Transl Med. 2022;12(1):e718. doi:10.1002/ctm2.718

369. Sun Z, Zeng L, Zhang M, Zhang Y, Yang N. PIM1 inhibitor synergizes the anti-tumor effect of osimertinib via STAT3 dephosphorylation in EGFR-mutant non-small cell lung cancer. Ann Transl Med. 2020;8(6):366. doi:10.21037/atm.2020.02.43

370. Wiest J, Clark AM, Dai W. Intron/exon organization and polymorphisms of the PLK3/PRK gene in human lung carcinoma cell lines. Genes Chromosomes Cancer. 2001;32(4):384–389. doi:10.1002/gcc.1204

371. Li X, Zhai S, Zhang J, Zhang D, Wang S, Wang L, Yu J.Interferon Regulatory Factor 4 Correlated With Immune Cells Infiltration Could Predict Prognosis for Patients With Lung Adenocarcinoma. Front Oncol. 2021;11:698465. doi:10.3389/fonc.2021.698465

372. Jin Y, Pan Y, Zheng S, Liu Y, Xu J, Peng Y, Zhang Z, Wang Y, Xiong Y, Xu L, et al. Inactivation of EGLN3 hydroxylase facilitates Erk3 degradation via autophagy and impedes lung cancer growth. Oncogene. 2022;41(12):1752–1766. doi:10.1038/s41388-022-02203-2

373. Kanaji N, Tadokoro A, Susaki K, Yokokura S, Ohmichi K, Haba R, Watanabe N, Bandoh S, Ishii T, Dobashi H, et al. Higher susceptibility of NOD/LtSz-scid Il2rg (-/-) NSG mice to xenotransplanted lung cancer cell lines. Cancer Manag Res. 2014;6:431–436. doi:10.2147/CMAR.S71185

374. Yasuda Y, Ozasa H, Kim YH, Yamazoe M, Ajimizu H, Yamamoto Funazo T, Nomizo T, Tsuji T, Yoshida H, Sakamori Y, et al. MCL1 inhibition is effective against a subset of small-cell lung cancer with high MCL1 and low BCL-XL expression. Cell Death Dis. 2020;11(3):177. doi:10.1038/s41419-020-2379-2

375. Tang SC, Wu CH, Lai CH, Sung WW, Yang WJ, Tang LC, Hsu CP, Ko JL. Glutathione S-transferase mu2 suppresses cancer cell metastasis in non-small cell lung cancer. Mol Cancer Res. 2013;11(5):518–529. doi:10.1158/1541-7786.MCR-12-0488

376. Walker T, Nolte A, Steger V, Makowiecki C, Mustafi M, Friedel G, Schlensak C, Wendel HP. Small interfering RNA-mediated suppression of serum response factor, E2-promotor binding factor and survivin in non-small cell lung cancer cell lines by non-viral transfection. Eur J Cardiothorac Surg. 2013;43(3):628–634. doi:10.1093/ejcts/ezs337

377. Zhang L, Zeng D, Chen Y, Li N, Lv Y, Li Y, Xu X, Xu G. miR-937 contributes to the lung cancer cell proliferation by targeting INPP4B. Life Sci. 2016;155:110–115. doi:10.1016/j.lfs.2016.05.014

378. Takai D, Yagi Y, Wakazono K, Ohishi N, Morita Y, Sugimura T, Ushijima T. Silencing of HTR1B and reduced expression of EDN1 in human lung cancers, revealed by methylation-sensitive representational difference analysis. Oncogene. 2001;20(51):7505–7513. doi:10.1038/sj.onc.120494

379. Román M, López I, Guruceaga E, Baraibar I, Ecay M, Collantes M, Nadal E, Vallejo A, Cadenas S, Miguel ME, et al. Inhibitor of Differentiation-1 Sustains Mutant KRAS-Driven Progression, Maintenance, and Metastasis of Lung Adenocarcinoma via Regulation of a FOSL1 Network. Cancer Res. 2019;79(3):625–638. doi:10.1158/0008-5472.CAN-18-1479

380. Li Z, Yu DP, Wang N, Tao T, Luo W, Chen H. SIRT5 promotes non-small cell lung cancer progression by reducing FABP4 acetylation level. Neoplasma. 2022;69(4):909–917. doi:10.4149/neo_2022_220107N28

381. Tan T, Han G, Cheng Z, Jiang J, Zhang L, Xia Z, Wang X, Xia Q. Genetic Polymorphisms in CYP2C19 Cause Changes in Plasma Levels and Adverse Reactions to Anlotinib in Chinese Patients With Lung Cancer. Front Pharmacol. 2022;13:918219. doi:10.3389/fphar.2022.918219

382. Wald O. CXCR4 Based Therapeutics for Non-Small Cell Lung Cancer (NSCLC). J Clin Med. 2018;7(10):303. doi:10.3390/jcm7100303

383. Luo J, Jin Y, Li M, Dong L. Tumor suppressor miRL613 induces cisplatin sensitivity in nonLsmall cell lung cancer cells by targeting GJA1. Mol Med Rep. 2021;23(5):385. doi:10.3892/mmr.2021.12024

384. Duplaquet L, Li Y, Booker MA, Xie Y, Olsen SN, Patel RA, Hong D, Hatton C, Denize T, Walton E, et al. KDM6A epigenetically regulates subtype plasticity in small cell lung cancer. Nat Cell Biol. 2023;25(9):1346–1358. doi:10.1038/s41556-023-01210-z

385. Zhu L, Zeng Q, Wang J, Deng F, Jin S. Cathepsin V drives lung cancer progression by shaping the immunosuppressive environment and adhesion molecules cleavage. Aging (Albany NY). 2023;15(23):13961–13979. doi:10.18632/aging.205278

386. Sun CC, Zhu W, Li SJ, Hu W, Zhang J, Zhuo Y, Zhang H, Wang J, Zhang Y, Huang SX, et al. FOXC1-mediated LINC00301 facilitates tumor progression and triggers an immune-suppressing microenvironment in non-small cell lung cancer by regulating the HIF1α pathway. Genome Med. 2020;12(1):77. doi:10.1186/s13073-020-00773-y

387. Cao Y, Guo C, Yin Y, Li X, Zhou L. LysineLspecific demethylase 2 contributes to the proliferation of small cell lung cancer by regulating the expression of TFPIL2. Mol Med Rep. 2018;18(1):733–740. doi:10.3892/mmr.2018.9047

388. Yang X, Zheng Y, Han Z, Zhang X. Functions and clinical significance of KLRG1 in the development of lung adenocarcinoma and immunotherapy. BMC Cancer. 2021;21(1):752. doi:10.1186/s12885-021-08510-3

389. Miura K, Koyanagi-Aoi M, Maniwa Y, Aoi T. Chorioallantoic membrane assay revealed the role of TIPARP (2,3,7,8-tetrachlorodibenzo-p-dioxin-inducible poly (ADP-ribose) polymerase) in lung adenocarcinoma-induced angiogenesis. Cancer Cell Int. 2023;23(1):34. doi:10.1186/s12935-023-02870-5

390. Zhang P, Zhang Y, Yang H, Li W, Chen X, Long F. Association between EPHX1 rs1051740 and lung cancer susceptibility: a meta-analysis. Int J Clin Exp Med. 2015;8(10):17941–17949.

391. Tong Q, Li D, Yin Y, Cheng L, Ouyang S. GBP5 Expression Predicted Prognosis of Immune Checkpoint Inhibitors in Small Cell Lung Cancer and Correlated with Tumor Immune Microenvironment. J Inflamm Res. 2023;16:4153–4164. doi:10.2147/JIR.S401430

392. Wei X, Png CW, Weerasooriya M, Li H, Zhu C, Chen G, Xu C, Zhang Y, Xu X. Tumor Promoting Function of DUSP10 in Non-Small Cell Lung Cancer Is Associated With Tumor-Promoting Cytokines. Immune Netw. 2023;23(4):e34. doi:10.4110/in.2023.23.e34

393. Yoshimura K, Suzuki Y, Inoue Y, Tsuchiya K, Karayama M, Iwashita Y, Kahyo T, Kawase A, Tanahashi M, Ogawa H, et al. CD200 and CD200R1 are differentially expressed and have differential prognostic roles in non-small cell lung cancer. Oncoimmunology. 2020;9(1):1746554. doi:10.1080/2162402X.2020.1746554

394. Song W, Wu X, Cheng C, Li D, Chen J, Zhang W. ARHGAP9 knockdown promotes lung adenocarcinoma metastasis by activating Wnt/β-catenin signaling pathway via suppressing DKK2. Genomics. 2023;115(5):110684. doi:10.1016/j.ygeno.2023.110684

395. Peng J, Wang Q, Liu H, Ye M, Wu X, Guo L. EPHA3 regulates the multidrug resistance of small cell lung cancer via the PI3K/BMX/STAT3 signaling pathway. Tumour Biol. 2016;37(9):11959–11971. doi:10.1007/s13277-016-5048-4

396. Chen P, Quan Z, Song X, Gao Z, Yuan K. MDFI is a novel biomarker for poor prognosis in LUAD. Front Oncol. 2022;12:1005962. doi:10.3389/fonc.2022.1005962

397. Sarode P, Zheng X, Giotopoulou GA, Weigert A, Kuenne C, Günther S, Friedrich A, Gattenlöhner S, Stiewe T, Brüne B, et al. Reprogramming of tumor-associated macrophages by targeting β-catenin/FOSL2/ARID5A signaling: A potential treatment of lung cancer. Sci Adv. 2020;6(23):eaaz6105. doi:10.1126/sciadv.aaz6105

398. Wang L, Wen J, Sun Y, Yang X, Ma Y, Tian X. Knockdown of NUPR1 inhibits angiogenesis in lung cancer through IRE1/XBP1 and PERK/eIF2α/ATF4 signaling pathways. Open Med (Wars). 2023;18(1):20230796. Published 2023 Oct 16. doi:10.1515/med-2023-0796

399. Tang Q, Wang X, Zhou Q, Li Q, Yang X, Xu M, Wang R, Chen J, Wu W, Wang S. Fuzheng Kang-Ai inhibits NSCLC cell proliferation via regulating hsa_circ_0048091/hsa-miR-378g/ARRDC3 pathway. Phytomedicine. 2023;114:154819. doi:10.1016/j.phymed.2023.154819

400. Honma R, Kinoshita I, Miyoshi E, Tomaru U, Matsuno Y, Shimizu Y, Takeuchi S, Kobayashi Y, Kaga K, Taniguchi N, et al. Expression of fucosyltransferase 8 is associated with an unfavorable clinical outcome in non-small cell lung cancers. Oncology. 2015;88(5):298–308. doi:10.1159/000369495

401. Chen P, Min J, Wu H, Zhang H, Wang C, Tan G, Zhang F. Annexin A1 is a potential biomarker of bone metastasis in small cell lung cancer. Oncol Lett. 2021;21(2):141. doi:10.3892/ol.2020.12402

402. Niu N, Ma X, Liu H, Zhao J, Lu C, Yang F, Qi W. DLC1 inhibits lung adenocarcinoma cell proliferation, migration and invasion via regulating MAPK signaling pathway. Exp Lung Res. 2021;47(4):173–182. doi:10.1080/01902148.2021.1885524

403. Zhu H, Liu Q, Yang X, Ding C, Wang Q, Xiong Y. LncRNA LINC00649 recruits TAF15 and enhances MAPK6 expression to promote the development of lung squamous cell carcinoma via activating MAPK signaling pathway. Cancer Gene Ther. 2022;29(8-9):1285–1295. doi:10.1038/s41417-021-00410-9

404. Gao Y, Ma HY, Xu QY, Li Y, Zhang SL, Qu YY, Zhang Y, Yin H. Mechanism of decorin protein inhibiting invasion and metastasis of non-small cell lung cancer. Eur Rev Med Pharmacol Sci. 2019;23(4):1520–1527. doi:10.26355/eurrev_201902_17110

405. Jusue-Torres I, Tiv R, Ricarte-Filho JC, Mallisetty A, Contreras-Vargas L, Godoy-Calderon MJ, Khaddour K, Kennedy K, Valyi-Nagy K, David O, et al. Myo1e overexpression in lung adenocarcinoma is associated with increased risk of mortality. Sci Rep. 2023;13(1):4107. doi:10.1038/s41598-023-30765-y

406. Xu Z, Zuo Y, Wang J, Yu Z, Peng F, Chen Y, Dong Y, Hu X, Zhou Q, Ma H, et al. Overexpression of the regulator of G-protein signaling 5 reduces the survival rate and enhances the radiation response of human lung cancer cells. Oncol Rep. 2015;33(6):2899–2907. doi:10.3892/or.2015.3917

407. Ingram K, Samson SC, Zewdu R, Zitnay RG, Snyder EL, Mendoza MC. NKX2-1 controls lung cancer progression by inducing DUSP6 to dampen ERK activity. Oncogene. 2022;41(2):293–300. doi:10.1038/s41388-021-02076-x

408. Kim YJ, Li W, Zhelev DV, Mellors JW, Dimitrov DS, Baek DS. Chimeric antigen receptor-T cells are effective against CEACAM5 expressing non-small cell lung cancer cells resistant to antibody-drug conjugates. Front Oncol. 2023;13:1124039. doi:10.3389/fonc.2023.1124039

409. Liu M, Xiao Y. Trefoil factor 3 silencing can inhibit the proliferation and apoptosis of lung cancer cells. J BUON. 2021;26(5):1842–1849.

410. Tian B, Han X, Li G, Jiang H, Qi J, Li J, Tian Y, Wang C. A Long Intergenic Non-coding RNA, LINC01426, Promotes Cancer Progression via AZGP1 and Predicts Poor Prognosis in Patients with LUAD. Mol Ther Methods Clin Dev. 2020;18:765–780. doi:10.1016/j.omtm.2020.08.001

411. Hsiao KC, Chu PY, Chang GC, Liu KJ. Elevated Expression of Lumican in Lung Cancer Cells Promotes Bone Metastasis through an Autocrine Regulatory Mechanism. Cancers (Basel). 2020;12(1):233. doi:10.3390/cancers12010233

412. Pang Y, Zhang Y, Zhang HY, Wang WH, Jin G, Liu JW, Zhu ZJ. MUC13 promotes lung cancer development and progression by activating ERK signaling. Oncol Lett. 2022;23(1):37. doi:10.3892/ol.2021.13155

413. Yu G, Herazo-Maya JD, Nukui T, Romkes M, Parwani A, Juan-Guardela BM, Robertson J, Gauldie J, Siegfried JM, Kaminski N, et al. Matrix metalloproteinase-19 promotes metastatic behavior in vitro and is associated with increased mortality in non-small cell lung cancer. Am J Respir Crit Care Med. 2014;190(7):780–790. doi:10.1164/rccm.201310-1903OC

414. Zhang H, Chen K, Zhou Y, Cao Z, Xu C, Zhou L, Wu G, Peng C, Lai S, Wu X. PLA2G2D fosters angiogenesis in non-small cell lung cancer through aerobic glycolysis. Growth Factors. 2024. doi:10.1080/08977194.2023.2297702

415. Liang Y, Xia W, Zhang T, Chen B, Wang H, Song X, Zhang Z, Xu L, Dong G, Jiang F. Upregulated Collagen COL10A1 Remodels the Extracellular Matrix and Promotes Malignant Progression in Lung Adenocarcinoma. Front Oncol. 2020;10:573534. doi:10.3389/fonc.2020.573534

416. Wu F, Wu Q. Corin-mediated processing of pro-atrial natriuretic peptide in human small cell lung cancer cells. Cancer Res. 2003;63(23):8318–8322.

417. Liu B, Wang Z, Gu M, Zhao C, Ma T, Wang J. GEO Data Mining Identifies OLR1 as a Potential Biomarker in NSCLC Immunotherapy. Front Oncol. 2021;11:629333. doi:10.3389/fonc.2021.629333

418. Liu Y, Fan J, Xu T, Ahmadinejad N, Hess K, Lin SH, Zhang J, Liu X, Liu L, Ning B, et al. Extracellular vesicle tetraspanin-8 level predicts distant metastasis in non-small cell lung cancer after concurrent chemoradiation. Sci Adv. 2020;6(11):eaaz6162. doi:10.1126/sciadv.aaz6162

419. Bi Y, Cao K, Wang Y, Yang W, Ma N, Lei X, Chen Y. Radiosensitivity in Non-Small-Cell Lung Cancer by MMP10 through the DNA Damage Repair Pathway. J Oncol. 2023;2023:5636852. doi:10.1155/2023/5636852

420. Zhan XK, Liu XK, Zhang S, Chen H. Expression and prognosis of inducible T-cell co-stimulator and its ligand in Chinese stage I-III lung adenocarcinoma patients. Animal Model Exp Med. 2023;6(5):464–473. doi:10.1002/ame2.12355

421. Lu Y, Xu W, Gu Y, Chang X, Wei G, Rong Z, Qin L, Chen X, Zhou F. Non-small Cell Lung Cancer Cells Modulate the Development of Human CD1c+ Conventional Dendritic Cell Subsets Mediated by CD103 and CD205. Front Immunol. 2019;10:2829. doi:10.3389/fimmu.2019.02829

422. Ma Y, Nenkov M, Schröder DC, Abubrig M, Gassler N, Chen Y. Fibulin 2 Is Hypermethylated and Suppresses Tumor Cell Proliferation through Inhibition of Cell Adhesion and Extracellular Matrix Genes in Non-Small Cell Lung Cancer. Int J Mol Sci. 2021;22(21):11834. doi:10.3390/ijms222111834

423. Wei S, Liu W, Xu M, Qin H, Liu C, Zhang R, Zhou S, Li E, Liu Z, Wang Q. Cathepsin F and Fibulin-1 as novel diagnostic biomarkers for brain metastasis of non-small cell lung cancer. Br J Cancer. 2022;126(12):1795–1805. doi:10.1038/s41416-022-01744-3

424. Kang JU, Koo SH, Kwon KC, Park JW, Kim JM. Identification of novel candidate target genes, including EPHB3, MASP1 and SST at 3q26.2-q29 in squamous cell carcinoma of the lung. BMC Cancer. 2009;9:237. doi:10.1186/1471-2407-9-237

425. Castellón Rubio VE, Segura PP, Muñoz A, Farré AL, Ruiz LC, Lorente JA. High plasma levels of soluble P-Selectin and Factor VIII predict venous thromboembolism in non-small cell lung cancer patients: The Thrombo-Nsclc risk score. Thromb Res. 2020;196:349–354. doi:10.1016/j.thromres.2020.09.021

426. Bae SY, Park HJ, Hong JY, Lee HJ, Lee SK. Down-regulation of SerpinB2 is associated with gefitinib resistance in non-small cell lung cancer and enhances invadopodia-like structure protrusions. Sci Rep. 2016;6:32258. doi:10.1038/srep32258

427. Zhang Q, Feng J, Liu K, Yang X, Huang Y, Tang B. STK11 mutation impacts CD1E expression to regulate the differentiation of macrophages in lung adenocarcinoma. Immun Inflamm Dis. 2023;11(7):e958. doi:10.1002/iid3.958

428. Park EJ, Jun HW, Na IH, Lee HK, Yun J, Kim HS, Kim Y, Hong JT, Han SB. CD48-expressing non-small-cell lung cancer cells are susceptible to natural killer cell-mediated cytotoxicity. Arch Pharm Res. 2022;45(1):1–10. doi:10.1007/s12272-021-01365-z

429. Qi X, Qi C, Wu T, Hu Y. CSF1R and HCST: Novel Candidate Biomarkers Predicting the Response to Immunotherapy in Non-Small Cell Lung Cancer. Technol Cancer Res Treat. 2020;19:1533033820970663. doi:10.1177/1533033820970663

430. Zhang H, Liu Q, Lei Y, Zhou J, Jiang W, Cui Y, He Q, Zhu J, Zhu Z, Sun Y, et al. Direct interaction between CD155 and CD96 promotes immunosuppression in lung adenocarcinoma. Cell Mol Immunol. 2021;18(6):1575–1577. doi:10.1038/s41423-020-00538-y

431. Peng X, Yang R, Peng W, Zhao Z, Tu G, He B, Cai Q, Shi S, Yin W, Yu F, et al. Overexpression of LINC00551 promotes autophagy-dependent ferroptosis of lung adenocarcinoma via upregulating DDIT4 by sponging miR-4328. PeerJ. 2022;10:e14180. doi:10.7717/peerj.14180

432. Lai IL, Chang YS, Chan WL, Lee YT, Yen JC, Yang CA, Hung SY, Chang JG. Male-Specific Long Noncoding RNA TTTY15 Inhibits Non-Small Cell Lung Cancer Proliferation and Metastasis via TBX4. Int J Mol Sci. 2019;20(14):3473. doi:10.3390/ijms20143473

433. Sui Q, Hu Z, Liang J, Lu T, Bian Y, Jin X, Li M, Huang Y, Yang H, Wang Q, et al. Targeting TAM-secreted S100A9 effectively enhances the tumor-suppressive effect of metformin in treating lung adenocarcinoma. Cancer Lett. 2024;581:216497. doi:10.1016/j.canlet.2023.216497

434. Wang C, Zhang C, Xu J, Li Y, Wang J, Liu H, Liu Y, Chen Z, Lin H. Association between IL-1R2 polymorphisms and lung cancer risk in the Chinese Han population: A case-control study. Mol Genet Genomic Med. 2019;7(5):e644. doi:10.1002/mgg3.644

435. Chen B, Zhang L, Zhou H, Ye W, Luo C, Yang L, Fang N, Tang A. HMOX1 promotes lung adenocarcinoma metastasis by affecting macrophages and mitochondrion complexes. Front Oncol. 2022;12:978006. doi:10.3389/fonc.2022.978006

436. Xu W, Jiang GJ, Shi GZ, Chen MZ, Ma TL, Tan YF. Metallothionein 1M (MT1M) inhibits lung adenocarcinoma cell viability, migration, and expression of cell mobility-related proteins through MDM2/p53/MT1M signaling. Transl Cancer Res. 2020;9(4):2710–2720. doi:10.21037/tcr.2020.02.61

437. Wang Y, Ha M, Li M, Zhang L, Chen Y. Histone deacetylase 6-mediated downregulation of TMEM100 expedites the development and progression of non-small cell lung cancer. Hum Cell. 2022;35(1):271–285. doi:10.1007/s13577-021-00635-8

438. Liu L, Liu H, Luo S, Patz EF Jr, Glass C, Su L, Lin L, Christiani DC, Wei Q. Genetic Variants of CLEC4E and BIRC3 in Damage-Associated Molecular Patterns-Related Pathway Genes Predict Non-Small Cell Lung Cancer Survival. Front Oncol. 2021;11:717109. doi:10.3389/fonc.2021.717109

439. Bhardwaj M, Schöttker B, Holleczek B, Brenner H. Enhanced selection of people for lung cancer screening using AHRR (cg05575921) or F2RL3 (cg03636183) methylation as biological markers of smoking exposure. Cancer Commun (Lond). 2023;43(8):956–959. doi:10.1002/cac2.12450

440. Yuan C, Zheng Y, Zhang B, Shao L, Liu Y, Tian T, Gu X, Li X, Fan K. Thymosin α1 promotes the activation of myeloid-derived suppressor cells in a Lewis lung cancer model by upregulating Arginase 1. Biochem Biophys Res Commun. 2015;464(1):249–255. doi:10.1016/j.bbrc.2015.06.132

441. Zhang H, Wang SQ, Zhu JB, Wang LN, Lin H, Li LF, Cheng YD, Duan CJ, Zhang CF. LncRNA CALML3-AS1 modulated by m6A modification induces BTNL9 methylation to drive non-small-cell lung cancer progression. Cancer Gene Ther. 2023;30(12):1649–1662. doi:10.1038/s41417-023-00670-7

442. Shi Q, Zheng X, Hu Y, Zhou Z, Fang M, Huang X. Methylation of hypoxia-inducible factor 3 subunit alpha contributes to poor prognosis in lung adenocarcinoma. J Appl Genet. 2023;64(4):769–777. doi:10.1007/s13353-023-00784-6

443. Bellezza G, Vannucci J, Bianconi F, Metro G, Del Sordo R, Andolfi M, Ferri I, Siccu P, Ludovini V, Puma F, et al. Prognostic implication of aquaporin 1 overexpression in resected lung adenocarcinoma. Interact Cardiovasc Thorac Surg. 2017;25(6):856–861. doi:10.1093/icvts/ivx202

444. Zhou W, Ma J, Meng L, Liu D, Chen J. Deletion of TRIB3 disrupts the tumor progression induced by integrin αvβ3 in lung cancer. BMC Cancer. 2022;22(1):459. doi:10.1186/s12885-022-09593-2

445. Chanda M, Anuntasomboon P, Ruangritchankul K, Cheepsunthorn P, Cheepsunthorn CL. Inhibition of non-small cell lung cancer (NSCLC) proliferation through targeting G6PD. PeerJ. 2023;11:e16503. doi:10.7717/peerj.16503

446. Huang B, Ding J, Guo H, Wang H, Xu J, Zheng Q, Zhou L. SIRT3 Regulates the ROS-FPR1/HIF-1α Axis under Hypoxic Conditions to Influence Lung Cancer Progression. Cell Biochem Biophys. 2023;81(4):813–821. doi:10.1007/s12013-023-01180-x

447. Xia D, Chen Z, Liu Q. Circ-PGC increases the expression of FOXR2 by targeting miR-532-3p to promote the development of non-small cell lung cancer. Cell Cycle. 2021;20(21):2195–2209. doi:10.1080/15384101.2021.1974788

448. Gu W, Zhang M, Gao F, Niu Y, Sun L, Xia H, Li W, Zhang Y, Guo Z, Du G. Berberine regulates PADI4-related macrophage function to prevent lung cancer. Int Immunopharmacol. 2022;110:108965. doi:10.1016/j.intimp.2022.108965

449. Ren C, Cui L, Li R, Song X, Li J, Xi Q, Zhang Z, Zhao L. Hsa_circ_0080608 Attenuates Lung Cancer Progression by Functioning as a Competitive Endogenous RNA to Regulate the miR-661/ADRA1A Pathway. Horm Metab Res. 2023;55(12):876–884. doi:10.1055/a-2179-0283

450. La Fleur L, Botling J, He F, Pelicano C, Zhou C, He C, Palano G, Mezheyeuski A, Micke P, Ravetch JV, et al. Targeting MARCO and IL37R on Immunosuppressive Macrophages in Lung Cancer Blocks Regulatory T Cells and Supports Cytotoxic Lymphocyte Function. Cancer Res. 2021;81(4):956–967. doi:10.1158/0008-5472.CAN-20-1885

451. Jang H, Jun Y, Kim S, Kim E, Jung Y, Park BJ, Lee J, Kim J, Lee S, Kim J. FCN3 functions as a tumor suppressor of lung adenocarcinoma through induction of endoplasmic reticulum stress. Cell Death Dis. 2021;12(4):407. doi:10.1038/s41419-021-03675-y

452. Liu X, Yu X, Xie J, Zhan M, Yu Z, Xie L, Zeng H, Zhang F, Chen G, Yi X, et al. ANGPTL2/LILRB2 signaling promotes the propagation of lung cancer cells. Oncotarget. 2015;6(25):21004–21015. doi:10.18632/oncotarget.4217

453. Xiong W, Hu XHW. AKR1C3 and β-catenin expression in non-small cell lung cancer and relationship with radiation resistance. J BUON. 2021;26(3):802–811.

454. Liao Y, Guo S, Chen Y, Cao D, Xu H, Yang C, Fei L, Ni B, Ruan Z. VSIG4 expression on macrophages facilitates lung cancer development. Lab Invest. 2014;94(7):706–715. doi:10.1038/labinvest.2014.73

455. Ercan H, Mauracher LM, Grilz E, Hell L, Hellinger R, Schmid JA, Moik F, Ay C, Pabinger I, Zellner M. Alterations of the Platelet Proteome in Lung Cancer: Accelerated F13A1 and ER Processing as New Actors in Hypercoagulability. Cancers (Basel). 2021;13(9):2260. doi:10.3390/cancers13092260

456. Li SQ, Feng J, Yang M, Ai XP, He M, Liu F. Sauchinone: a prospective therapeutic agent-mediated EIF4EBP1 down-regulation suppresses proliferation, invasion and migration of lung adenocarcinoma cells. J Nat Med. 2020;74(4):777–787. doi:10.1007/s11418-020-01435-4

457. Xiang L, Wang Y, Lan J, Na F, Wu S, Gong Y, Du H, Shao B, Xie G. HIF-1-dependent heme synthesis promotes gemcitabine resistance in human non-small cell lung cancers via enhanced ABCB6 expression. Cell Mol Life Sci. 2022;79(6):343. doi:10.1007/s00018-022-04360-9

458. Jin Y, Zhang Y, Huang A, Chen Y, Wang J, Liu N, Wang X, Gong Y, Wang W, Pan J. Overexpression of SERPINA3 suppresses tumor progression by modulating SPOP/NFLκB in lung cancer. Int J Oncol. 2023;63(2):96. doi:10.3892/ijo.2023.5544

459. Luo L, Zheng Y, Lin Z, Li X, Li X, Li M, Cui L, Luo H. Identification of SHMT2 as a Potential Prognostic Biomarker and Correlating with Immune Infiltrates in Lung Adenocarcinoma. J Immunol Res. 2021;2021:6647122. doi:10.1155/2021/6647122

460. Jing X, Yun Y, Ji X, Yang E, Li P. Pyroptosis and Inflammasome-Related Genes-NLRP3, NLRC4 and NLRP7 Polymorphisms Were Associated with Risk of Lung Cancer. Pharmgenomics Pers Med. 2023;16:795–804. doi:10.2147/PGPM.S424326

461. Zhang S, Wang Y, Dai SD, Wang EH. Down-regulation of NKD1 increases the invasive potential of non-small-cell lung cancer and correlates with a poor prognosis. BMC Cancer. 2011;11:186. doi:10.1186/1471-2407-11-186

462. Xing S, Zeng T, Xue N, He Y, Lai YZ, Li HL, Huang Q, Chen SL, Liu WL. Development and Validation of Tumor-educated Blood Platelets Integrin Alpha 2b (ITGA2B) RNA for Diagnosis and Prognosis of Non-small-cell Lung Cancer through RNA-seq. Int J Biol Sci. 2019;15(9):1977–1992. doi:10.7150/ijbs.36284

463. Chen J, Ye L, Zhang L, Jiang WG. Placenta growth factor, PLGF, influences the motility of lung cancer cells, the role of Rho associated kinase, Rock1. J Cell Biochem. 2008;105(1):313–320. doi:10.1002/jcb.21831

464. Siddiqui MA, Gollavilli PN, Ramesh V, Parma B, Schwab A, Vazakidou ME, Natesan R, Saatci O, Rapa I, Bironzo P, et al. Thymidylate synthase drives the phenotypes of epithelial-to-mesenchymal transition in non-small cell lung cancer. Br J Cancer. 2021;124(1):281–289. doi:10.1038/s41416-020-01095-x

465. Lian YX, Chen R, Xu YH, Peng CL, Hu HC. Effect of protein-tyrosine phosphatase 4A3 by small interfering RNA on the proliferation of lung cancer. Gene. 2012;511(2):169–176. doi:10.1016/j.gene.2012.09.079

466. Peng F, Li Q, Niu SQ, Shen GP, Luo Y, Chen M, Bao Y. ZWINT is the next potential target for lung cancer therapy. J Cancer Res Clin Oncol. 2019;145(3):661–673. doi:10.1007/s00432-018-2823-1

467. Fan J, Xia X, Fan Z. Hsa_circ_0129047 regulates the miR-375/ACVRL1 axis to attenuate the progression of lung adenocarcinoma. J Clin Lab Anal. 2022;36(9):e24591. doi:10.1002/jcla.24591

468. Dai N, Ma H, Feng Y. Silencing of long non-coding RNA SDCBP2-AS1/microRNA-656-3p/CRIM1 axis promotes ferroptosis of lung cancer cells. Cell Mol Biol (Noisy-le-grand). 2023;69(9):189–194. doi:10.14715/cmb/2023.69.9.29

469. Wang P, Zhang Y, Lv X, Zhou J, Cang S, Song Y. LncRNA ADAMTS9-AS1 inhibits the stemness of lung adenocarcinoma cells by regulating miR-5009-3p/NPNT axis. Genomics. 2023;115(3):110596. doi:10.1016/j.ygeno.2023.110596

470. Li ZH, Liu C, Liu QH, Wang J, Wang Y, Wang YF, Deng SJ, Li DB. Cytoplasmic expression of G protein-coupled estrogen receptor 1 correlates with poor postoperative prognosis in non-small cell lung cancer. J Thorac Dis. 2022;14(5):1466–1477. doi:10.21037/jtd-22-29

471. Wu J, Zhang L, Li W, Wang L, Jia Q, Shi F, Li K, Liao L, Shi Y, Wu S. The role of TOP2A in immunotherapy and vasculogenic mimicry in non-small cell lung cancer and its potential mechanism. Sci Rep. 2023;13(1):10906. doi:10.1038/s41598-023-38117-6

472. Ali A, Levantini E, Teo JT, Goggi J, Clohessy JG, Wu CS, Chen L, Yang H, Krishnan I, Kocher O, et al. Fatty acid synthase mediates EGFR palmitoylation in EGFR mutated non-small cell lung cancer. EMBO Mol Med. 2018;10(3):e8313. doi:10.15252/emmm.201708313

473. Venuto S, Coda ARD, González-Pérez R, Laselva O, Tolomeo D, Storlazzi CT, Liso A, Conese M. IGFBP-6 Network in Chronic Inflammatory Airway Diseases and Lung Tumor Progression. Int J Mol Sci. 2023;24(5):4804. doi:10.3390/ijms24054804

474. Li D, Zhang X, Ding Z, Ai R, Shi L, Wang Z, He Q, Dong Y, Zhu Y, Ouyang W, et al. Identification and Exploration of Serine Peptidase Inhibitor Kazal Type I (SPINK1) as a Potential Biomarker Correlated with the Progression of Non-Small Cell Lung Cancer. Cell Biochem Biophys. 2022;80(4):807–818. doi:10.1007/s12013-022-01098-w

475. Yin J, Zhang Y, Zhang Y, Peng F, Lu Y. Reporting on Two Novel Fusions, DYSF-ALK and ITGAV-ALK, Coexisting in One Patient with Adenocarcinoma of Lung, Sensitive to Crizotinib. J Thorac Oncol. 2018;13(3):e43–e45. doi:10.1016/j.jtho.2017.10.025

476. Zhu X, Chen X, Shen X, Liu Y, Fu W, Wang B, Zhao L, Yang F, Mo N, Zhong G, et al. PP4R1 accelerates the malignant progression of NSCLC via up-regulating HSPA6 expression and HSPA6-mediated ER stress. Biochim Biophys Acta Mol Cell Res. 2024;1871(1):119588. doi:10.1016/j.bbamcr.2023.119588

477. Wang L, Wen J, Sun Y, Yang X, Ma Y, Tian X. Knockdown of NUPR1 inhibits angiogenesis in lung cancer through IRE1/XBP1 and PERK/eIF2α/ATF4 signaling pathways. Open Med (Wars). 2023;18(1):20230796. doi:10.1515/med-2023-0796

478. Zhang J, Gu Y, Liu X, Rao X, Huang G, Ouyang Y. Clinicopathological and prognostic value of S100A4 expression in non-small cell lung cancer: a meta-analysis. Biosci Rep. 2020;40(7):BSR20201710. doi:10.1042/BSR20201710

479. Olingy C, Alimadadi A, Araujo DJ, Barry D, Gutierrez NA, Werbin MH, Arriola E, Patel SP, OttenCD33 Expression on Peripheral Blood Monocytes Predicts Efficacy of Anti-PD-1 Immunotherapy Against Non-Small Cell Lung Cancer. Front Immunol. 2022;13:842653. doi:10.3389/fimmu.2022.

480. Wang W, Xiong Y, Ding X, Wang L, Zhao Y, Fei Y, Zhu Y, Shen X, Tan C, Liang Z. Cathepsin L activated by mutant p53 and Egr-1 promotes ionizing radiation-induced EMT in human NSCLC. J Exp Clin Cancer Res. 2019;38(1):61. doi:10.1186/s13046-019-1054-x

481. Wang YL, Wu ZZ, Zhang HR, Chen DS, Zhao X. Coexistence of a novel RGS18 downstream intergenic region ALK fusion and a THUMPD2-ALK fusion in a lung adenocarcinoma patient and response to crizotinib. Lung Cancer. 2021;154:216–218. doi:10.1016/j.lungcan.2021.02.008

482. Zeng B, Ge C, Li R, Zhang Z, Fu Q, Li Z, Lin Z, Liu L, Xue Y, Xu Y, et al. Knockdown of microsomal glutathione S-transferase 1 inhibits lung adenocarcinoma cell proliferation and induces apoptosis. Biomed Pharmacother. 2020;121:109562. doi:10.1016/j.biopha.2019.109562

483. Yu S, Li Y, Ren H, Zhou H, Ning Q, Chen X, Hu T, Yang L. PDK4 promotes tumorigenesis and cisplatin resistance in lung adenocarcinoma via transcriptional regulation of EPAS1. Cancer Chemother Pharmacol. 2021;87(2):207–215. doi:10.1007/s00280-020-04188-9

484. Goda H, Nakashiro KI, Sano Y, Adachi T, Tokuzen N, Kuribayashi N, Hino S, Uchida D. KRT13 and UPK1B for differential diagnosis between metastatic lung carcinoma from oral squamous cell carcinoma and lung squamous cell carcinoma. Sci Rep. 2023;13(1):22626. doi:10.1038/s41598-023-49545-9

485. Chang LL, Lu PH, Yang W, Hu Y, Zheng L, Zhao Q, Lin NM, Zhang WZ. AKR1C1 promotes non-small cell lung cancer proliferation via crosstalk between HIF-1α and metabolic reprogramming. Transl Oncol. 2022;20:101421. doi:10.1016/j.tranon.2022.101421

486. Kahm YJ, Kim RK. BIRC5: A novel therapeutic target for lung cancer stem cells and glioma stem cells. Biochem Biophys Res Commun. 2023;682:141–147. doi:10.1016/j.bbrc.2023.10.008

487. Wang T, Chen X, Qiao W, Kong L, Sun D, Li Z. Transcription factor E2F1 promotes EMT by regulating ZEB2 in small cell lung cancer. BMC Cancer. 2017;17(1):719. doi:10.1186/s12885-017-3701-y

488. Wang ZH, Ye LL, Xiang X, Wei XS, Niu YR, Peng WB, Zhang SY, Zhang P, Xue QQ, Wang HL, et al. Circular RNA circFBXO7 attenuates non-small cell lung cancer tumorigenesis by sponging miR-296-3p to facilitate KLF15-mediated transcriptional activation of CDKN1A. Transl Oncol. 2023;30:101635. doi:10.1016/j.tranon.2023.101635

489. Li HJ, Ke FY, Lin CC, Lu MY, Kuo YH, Wang YP, Liang KH, Lin SC, Chang YH, Chen HY, et al. ENO1 Promotes Lung Cancer Metastasis via HGFR and WNT Signaling-Driven Epithelial-to-Mesenchymal Transition. Cancer Res. 2021;81(15):4094–4109. doi:10.1158/0008-5472.CAN-20-3543

490. Li Y, Song X, Niu J, Ren M, Tang G, Sun Z, Kong F. Pentraxin 3 acts as a functional effector of Akt/NF-κB signaling to modulate the progression and cisplatin-resistance in non-small cell lung cancer. Arch Biochem Biophys. 2021;701:108818. doi:10.1016/j.abb.2021.108818

491. Rodriguez EF, De Marchi F, Lokhandwala PM, Belchis D, Xian R, Gocke CD, Eshleman JR, Illei P, Li MT. IDH1 and IDH2 mutations in lung adenocarcinomas: Evidences of subclonal evolution. Cancer Med. 2020;9(12):4386–4394. doi:10.1002/cam4.3058

492. Zhang Y, Song D, Peng Z, Wang R, Li K, Ren H, Sun X, Du N, Tang SC. LINC00891 regulated by miR-128-3p/GATA2 axis impedes lung cancer cell proliferation, invasion and EMT by inhibiting RhoA pathway. Acta Biochim Biophys Sin (Shanghai). 2022;54(3):378–387. doi:10.3724/abbs.202200

493. Kong X, Bu J, Chen J, Ni B, Fu B, Zhou F, Pang S, Zhang J, Xu S, He C. PIGF and Flt-1 on the surface of macrophages induces the production of TGF-β1 by polarized tumor-associated macrophages to promote lung cancer angiogenesis. Eur J Pharmacol. 2021;912:174550. doi:10.1016/j.ejphar.2021.174550

494. Gong F, Li R, Zheng X, Chen W, Zheng Y, Yang Z, Chen Y, Qu H, Mao E, Chen E. OLFM4 Regulates Lung Epithelial Cell Function in Sepsis-Associated ARDS/ALI via LDHA-Mediated NF-κB Signaling. J Inflamm Res. 2021;14:7035–7051. doi:10.2147/JIR.S335915

495. Song M, Gao L, Zang J, Xing X. ABCA3, a tumor suppressor gene, inhibits the proliferation, migration and invasion of lung adenocarcinoma by regulating the epithelialLmesenchymal transition process. Oncol Lett. 2023;26(4):420. doi:10.3892/ol.2023.14006

496. Potiron VA, Sharma G, Nasarre P, Clarhaut JA, Augustin HG, Gemmill RM, Roche J, Drabkin HA. Semaphorin SEMA3F affects multiple signaling pathways in lung cancer cells. Cancer Res. 2007;67(18):8708–8715. doi:10.1158/0008-5472.CAN-06-3612

497. Deng P, Zhou R, Zhang J, Cao L. Increased Expression of KNSTRN in Lung Adenocarcinoma Predicts Poor Prognosis: A Bioinformatics Analysis Based on TCGA Data. J Cancer. 2021;12(11):3239–3248. doi:10.7150/jca.51591

498. Wang D, Lin Y, Gao B, Yan S, Wu H, Li Y, Wu Q, Wei Y. Reduced Expression of FADS1 Predicts Worse Prognosis in Non-Small-Cell Lung Cancer. J Cancer. 2016;7(10):1226–1232. doi:10.7150/jca.15403

499. Little AC, Sham D, Hristova M, Danyal K, Heppner DE, Bauer RA, Sipsey LM, Habibovic A, van der Vliet A.DUOX1 silencing in lung cancer promotes EMT, cancer stem cell characteristics and invasive properties. Oncogenesis. 2016;5(10):e261. doi:10.1038/oncsis.2016.61

500. Zhu D, Nie Y, Zhao Y, Chen X, Yang Z, Yang Y. RNF152 Suppresses Fatty Acid Oxidation and Metastasis of Lung Adenocarcinoma by Inhibiting IRAK1-Mediated AKR1B10 Expression. Am J Pathol. 2023;193(10):1603–1617. doi:10.1016/j.ajpath.2023.06.014

501. Ochiai R, Hayashi K, Yamamoto H, Fujii R, Saichi N, Shinchi H, Ishida T, Honda T, Shimizu T, Matsutani N, et al. Plasma exosomal DOK3 reflects immunological states in lung tumor and predicts prognosis of gefitinib treatment. Cancer Sci. 2022;113(11):3960–3971. doi:10.1111/cas.15512

502. Wang Y, Chen L, Huang G, He D, He J, Xu W, Zou C, Zong F, Li Y, Chen B, et al. Klotho sensitizes human lung cancer cell line to cisplatin via PI3k/Akt pathway. PLoS One. 2013;8(2):e57391. doi:10.1371/journal.pone.0057391

503. Li Z, Chen S, He X, Gong S, Sun L, Weng L. SLC3A2 promotes tumor-associated macrophage polarization through metabolic reprogramming in lung cancer. Cancer Sci. 2023;114(6):2306–2317. doi:10.1111/cas.15760

504. Wu DM, Liu T, Deng SH, Han R, Xu Y. SLC39A4 expression is associated with enhanced cell migration, cisplatin resistance, and poor survival in non-small cell lung cancer. Sci Rep. 2017;7(1):7211. doi:10.1038/s41598-017-07830-4

505. Wu Y, Feng J, Hu W, Zhang Y. T-box 3 overexpression is associated with poor prognosis of non-small cell lung cancer. Oncol Lett. 2017;13(5):3335–3341. doi:10.3892/ol.2017.5855

506. Zhang C, Liu T, Wang G, Wang H, Che X, Gao X, Liu H. Rac3 Regulates Cell Invasion, Migration and EMT in Lung Adenocarcinoma through p38 MAPK Pathway. J Cancer. 2017;8(13):2511–2522. doi:10.7150/jca.18161

507. Tang M, Sun J, Cai Z. PCK2 inhibits lung adenocarcinoma tumor cell immune escape through oxidative stress-induced senescence as a potential therapeutic target. J Thorac Dis. 2023;15(5):2601–2615. doi:10.21037/jtd-23-542

508. Reda M, Ngamcherdtrakul W, Nelson MA, Siriwon N, Wang R, Zaidan HY, Bejan DS, Reda S, Hoang NH, Crumrine NA, et al. Development of a nanoparticle-based immunotherapy targeting PD-L1 and PLK1 for lung cancer treatment. Nat Commun. 2022;13(1):4261. doi:10.1038/s41467-022-31926-9

509. Han Y, Dong Q, Hao J, Fu L, Han X, Zheng X, Wang E. RASSF4 is downregulated in nonsmall cell lung cancer and inhibits cancer cell proliferation and invasion. Tumour Biol. 2016;37(4):4865–4871. doi:10.1007/s13277-015-4343-9

510. Jiang B, Shi W, Li P, Wu Y, Li Y, Bao C. The mechanism of and the association between interleukin-27 and chemotherapeutic drug sensitivity in lung cancer. Oncol Lett. 2021;21(1):14. doi:10.3892/ol.2020.12275

511. Liu YN, Tsai MF, Wu SG, Chang TH, Tsai TH, Gow CH, Chang YL, Shih JY. Acquired resistance to EGFR tyrosine kinase inhibitors is mediated by the reactivation of STC2/JUN/AXL signaling in lung cancer. Int J Cancer. 2019;145(6):1609–1624. doi:10.1002/ijc.32487

512. Gürz S, Çelik B, Menteşe A, Us Altay D. Diagnostic value of signal peptide-Complement C1r/C1s, Uegf, and Bmp1-epidermal growth factor domain-containing protein 1 on serum and tissue samples in non-small cell lung cancer. Turk Gogus Kalp Damar Cerrahisi Derg. 2018;26(2):246–253. doi:10.5606/tgkdc.dergisi.2018.14600

513. Whitt AG, Neely AM, Sarkar OS, Meng S, Arumugam S, Yaddanapudi K, Li C. Paraoxonase 2 (PON2) plays a limited role in murine lung tumorigenesis. Sci Rep. 2023;13(1):9929. doi:10.1038/s41598-023-37146-5

514. Xu M, Zheng J, Wang J, Huang H, Hu G, He H. MCF2L-AS1/miR-874-3p/STAT3 feedback loop contributes to lung adenocarcinoma cell growth and cisplatin resistance. Heliyon. 2023;9(11):e21342. doi:10.1016/j.heliyon.2023.e21342

515. Zhang L, Zhao X, Wang E, Yang Y, Hu L, Xu H, Zhang B. PYCR1 promotes the malignant progression of lung cancer through the JAK-STAT3 signaling pathway via PRODH-dependent glutamine synthesize. Transl Oncol. 2023;32:101667. doi:10.1016/j.tranon.2023.101667

516. Tokunaga K, Nakamura Y, Sakata K, Fujimori K, Ohkubo M, Sawada K, Sakiyama S. Enhanced expression of a glyceraldehyde-3-phosphate dehydrogenase gene in human lung cancers. Cancer Res. 1987;47(21):5616–5619.

517. Yang B, Zhang B, Qi Q, Wang C. CircRNA has_circ_0017109 promotes lung tumor progression via activation of Wnt/β-catenin signaling due to modulating miR-671-5p/FZD4 axis. BMC Pulm Med. 2022;22(1):443. doi:10.1186/s12890-022-02209-2

518. Liu Y, Fan X, Jiang C, Xu S. SPOCK2 and SPRED1 function downstream of EZH2 to impede the malignant progression of lung adenocarcinoma in vitro and in vivo. Hum Cell. 2023;36(2):812–821. doi:10.1007/s13577-023-00855-0

519. Chakrabarti G. Mutant KRAS associated malic enzyme 1 expression is a predictive marker for radiation therapy response in non-small cell lung cancer. Radiat Oncol. 2015;10:145. doi:10.1186/s13014-015-0457-x

520. Liu HY, Zhao H, Li WX. Integrated Analysis of Transcriptome and Prognosis Data Identifies FGF22 as a Prognostic Marker of Lung Adenocarcinoma. Technol Cancer Res Treat. 2019;18:1533033819827317. doi:10.1177/1533033819827317

521. Cosic-Mujkanovic N, Valadez-Cosmes P, Maitz K, Lueger A, Mihalic ZN, Runtsch MC, Kienzl M, Davies MJ, Chuang CY, Heinemann A, et al. Myeloperoxidase Alters Lung Cancer Cell Function to Benefit Their Survival. Antioxidants (Basel). 2023;12(8):1587. doi:10.3390/antiox12081587

522. Yang XG, Li YY, Zhao DX, Cui W, Li H, Li XY, Li YX, Wang D. Repurposing of a monoamine oxidase A inhibitorLheptamethine carbocyanine dye conjugate for paclitaxelLresistant nonLsmall cell lung cancer. Oncol Rep. 2021;45(3):1306–1314. doi:10.3892/or.2021.7950

523. Nashed M, Chisholm JW, Igal RA. Stearoyl-CoA desaturase activity modulates the activation of epidermal growth factor receptor in human lung cancer cells. Exp Biol Med (Maywood). 2012;237(9):1007–1017. doi:10.1258/ebm.2012.012126

524. Podgornik H, Sok M, Kern I, Marc J, Cerne D. Lipoprotein lipase in non-small cell lung cancer tissue is highly expressed in a subpopulation of tumor-associated macrophages. Pathol Res Pract. 2013;209(8):516–520. doi:10.1016/j.prp.2013.06.004

525. Jovanović M, Stanković T, Stojković Burić S, Banković J, Dinić J, Ljujić M, Pešić M, Dragoj M. Decreased TSPAN14 Expression Contributes to NSCLC Progression. Life (Basel). 2022;12(9):1291. doi:10.3390/life12091291

526. Park SL, Murphy SE, Wilkens LR, Stram DO, Hecht SS, Le Marchand L. Association of CYP2A6 activity with lung cancer incidence in smokers: The multiethnic cohort study. PLoS One. 2017;12(5):e0178435. doi:10.1371/journal.pone.0178435

527. Tang D, Zhao YC, Liu H, Luo S, Clarke JM, Glass C, Su L, Shen S, Christiani DC, Gao W, et al. Potentially functional genetic variants in PLIN2, SULT2A1 and UGT1A9 genes of the ketone pathway and survival of nonsmall cell lung cancer. Int J Cancer. 2020;147(6):1559–1570. doi:10.1002/ijc.32932

528. Zhang W, Liu K, Pei Y, Tan J, Ma J, Zhao J. Long Noncoding RNA HIF1A-AS2 Promotes Non-Small Cell Lung Cancer Progression by the miR-153-5p/S100A14 Axis. Onco Targets Ther. 2020;13:8715–8722. doi:10.2147/OTT.S262293

529. Chen M, Zhang J, Berger AH, Diolombi MS, Ng C, Fung J, Bronson RT, Castillo-Martin M, Thin TH, Cordon-Cardo C, et al. Compound haploinsufficiency of Dok2 and Dusp4 promotes lung tumorigenesis. J Clin Invest. 2019;129(1):215–222. doi:10.1172/JCI99699

530. Shen M, Wang YJ, Liu ZH, Chen YW, Liang QK, Li Y, Ming HX. Inhibitory Effect of Astragalus Polysaccharide on Premetastatic Niche of Lung Cancer through the S1PR1-STAT3 Signaling Pathway. Evid Based Complement Alternat Med. 2023;2023:4010797. doi:10.1155/2023/4010797

531. Liu F, Wu H. Prognostic Value of Gastrokine-2 (GKN2) and Its Correlation with Tumor-Infiltrating Immune Cells in Lung Cancer and Gastric Cancers. J Inflamm Res. 2020;13:933–944. doi:10.2147/JIR.S277353

532. Zhou J, Jiang G, Xu E, Zhou J, Liu L, Yang Q. Identification of SRXN1 and KRT6A as Key Genes in Smoking-Related Non-Small-Cell Lung Cancer Through Bioinformatics and Functional Analyses. Front Oncol. 2022;11:810301. doi:10.3389/fonc.2021.810301

533. Wang L, Zhao X, Zheng H, Zhu C, Liu Y. AIF-1, a potential biomarker of aggressive tumor behavior in patients with non-small cell lung cancer. PLoS One. 2022;17(12):e0279211. doi:10.1371/journal.pone.0279211

534. Xia F, Xie M, He J, Cheng D. Circ_0004140 promotes lung adenocarcinoma progression by upregulating NOVA2 via sponging miR-330-5p. Thorac Cancer. 2023;14(35):3483–3494. doi:10.1111/1759-7714.15141

535. Fan C, Lin X, Wang E. Clinicopathological significance of cathepsin D expression in non-small cell lung cancer is conditional on apoptosis-associated protein phenotype: an immunohistochemistry study. Tumour Biol. 2012;33(4):1045–1052. doi:10.1007/s13277-012-0338-y

536. Patel V, Bardoliwala D, Lalani R, Patil S, Ghosh S, Javia A, Misra A. Development of a dry powder for inhalation of nanoparticles codelivering cisplatin and ABCC3 siRNA in lung cancer. Ther Deliv. 2021;12(9):651–670. doi:10.4155/tde-2020-0117

537. Zhou J, Xu Y, Wang G, Mei T, Yang H, Liu Y. The TLR7/8 agonist R848 optimizes host and tumor immunity to improve therapeutic efficacy in murine lung cancer. Int J Oncol. 2022;61(1):81. doi:10.3892/ijo.2022.5371

538. Xu T, Li D, Wang H, Zheng T, Wang G, Xin Y. MUC1 downregulation inhibits non-small cell lung cancer progression in human cell lines. Exp Ther Med. 2017;14(5):4443–4447. doi:10.3892/etm.2017.5062

539. Li Z, Jiang D, Liu F, Li Y. Involvement of ZDHHC9 in lung adenocarcinoma: regulation of PD-L1 stability via palmitoylation. In Vitro Cell Dev Biol Anim. 2023;59(3):193–203. doi:10.1007/s11626-023-00755-5

540. Yang S, Tang D, Zhao YC, Liu H, Luo S, Stinchcombe TE, Glass C, Su L, Shen S, Christiani DC, et al. Potentially functional variants of ERAP1, PSMF1 and NCF2 in the MHC-I-related pathway predict non-small cell lung cancer survival. Cancer Immunol Immunother. 2021;70(10):2819–2833. doi:10.1007/s00262-021-02877-9

541. Wu CY, Chan CH, Dubey NK, Wei HJ, Lu JH, Chang CC, Cheng HC, Ou KL, Deng WP. Highly Expressed FOXF1 Inhibit Non-Small-Cell Lung Cancer Growth via Inducing Tumor Suppressor and G1-Phase Cell-Cycle Arrest. Int J Mol Sci. 2020;21(9):3227. doi:10.3390/ijms21093227

542. Gao W, Liu Y, Qin R, Liu D, Feng Q. Silence of fibronectin 1 increases cisplatin sensitivity of non-small cell lung cancer cell line. Biochem Biophys Res Commun. 2016;476(1):35–41. doi:10.1016/j.bbrc.2016.05.081

543. Alam S, Mohammad T, Padder RA, Hassan MI, Husain M. Thymoquinone and quercetin induce enhanced apoptosis in non-small cell lung cancer in combination through the Bax/Bcl2 cascade. J Cell Biochem. 2022;123(2):259–274. doi:10.1002/jcb.30162

544. Chen F, Takenaka K, Ogawa E, Yanagihara K, Otake Y, Wada H, Tanaka F. Flt-4-positive endothelial cell density and its clinical significance in non-small cell lung cancer. Clin Cancer Res. 2004;10(24):8548–8553. doi:10.1158/1078-0432.CCR-04-0950

545. Zhang B, Jin Z, Zhang H. LINC01207 promotes the progression of non-small cell lung cancer via regulating ARHGAP11A by sponging miR-525-5p. Cancer Biomark. 2022;33(3):401–414. doi:10.3233/CBM-203197

546. Chang W, Zhu J, Yang D, Shang A, Sun Z, Quan W, Li D. Plasma versican and plasma exosomal versican as potential diagnostic markers for non-small cell lung cancer. Respir Res. 2023;24(1):140. doi:10.1186/s12931-023-02423-4

547. Xu B, Lou Y, Xu X, Li X, Tian X, Yu Z, Chen X. Carbonic Anhydrase 4 Serves as A Novel Prognostic Biomarker and Therapeutic Target for Non-Small Cell Lung Cancer: A Study Based on TCGA Samples. Comb Chem High Throughput Screen. 2023;26(14):2527–2540. doi:10.2174/1386207326666230321091943

548. Li R, Mu C, Cao Y, Fan Y. METTL7B serves as a prognostic biomarker and promotes metastasis of lung adenocarcinoma cells. Ann Transl Med. 2022;10(16):895. doi:10.21037/atm-22-3849

549. Niu C, Qiu W, Li X, Li H, Zhou J, Zhu H. Transketolase Serves as a Biomarker for Poor Prognosis in Human Lung Adenocarcinoma. J Cancer. 2022;13(8):2584–2593. doi:10.7150/jca.69583

550. Wei S, Shao J, Wang J, Zhao T, Liu J, Shen X, Wang Y, Chen H, Wang G. EHD2 inhibits the invasive ability of lung adenocarcinoma and improves the prognosis of patients. J Thorac Dis. 2022;14(7):2652–2664. doi:10.21037/jtd-22-842

551. Li M, Li J, Guo X, Pan H, Zhou Q. Absence of HTATIP2 Expression in A549 Lung Adenocarcinoma Cells Promotes Tumor Plasticity in Response to Hypoxic Stress. Cancers (Basel). 2020;12(6):1538. doi:10.3390/cancers12061538

552. Lee YS, Oh JH, Yoon S, Kwon MS, Song CW, Kim KH, Cho MJ, Mollah ML, Je YJ, Kim YD, et al. ifferential gene expression profiles of radioresistant non-small-cell lung cancer cell lines established by fractionated irradiation: tumor protein p53-inducible protein 3 confers sensitivity to ionizing radiation. Int J Radiat Oncol Biol Phys. 2010;77(3):858–866. doi:10.1016/j.ijrobp.2009.12.076

553. Li H, Wu C, Chang W, Zhong L, Gao W, Zeng M, Wen Z, Mai S, Chen Y. Overexpression of PSAT1 is Correlated with Poor Prognosis and Immune Infiltration in Non-Small Cell Lung Cancer. Front Biosci (Landmark Ed). 2023;28(10):243. doi:10.31083/j.fbl2810243

554. Zhang S, You X, Zheng Y, Shen Y, Xiong X, Sun Y. The UBE2C/CDH1/DEPTOR axis is an oncogene and tumor suppressor cascade in lung cancer cells. J Clin Invest. 2023;133(4):e162434. doi:10.1172/JCI162434

555. Zhou F, Yuan Z, Gong Y, Li L, Wang Y, Wang X, Ma C, Yang L, Liu Z, Wang L, et al. Pharmacological targeting of MTHFD2 suppresses NSCLC via the regulation of ILK signaling pathway. Biomed Pharmacother. 2023;161:114412. doi:10.1016/j.biopha.2023.114412

556. Zhang Z, Rui W, Wang ZC, Liu DX, Du L. Anti-proliferation and anti-metastasis effect of barbaloin in non-small cell lung cancer via inactivating p38MAPK/Cdc25B/Hsp27 pathway. Oncol Rep. 2017;38(2):1172–1180. doi:10.3892/or.2017.5760

557. Malvi P, Janostiak R, Nagarajan A, Cai G, Wajapeyee N. Loss of thymidine kinase 1 inhibits lung cancer growth and metastatic attributes by reducing GDF15 expression. PLoS Genet. 2019;15(10):e1008439. doi:10.1371/journal.pgen.1008439

558. Zhang Z, Bi X, Lian X, Niu Z. BDH1 promotes lung cancer cell proliferation and metastases by PARP1-mediated autophagy. J Cell Mol Med. 2023;27(7):939–949. doi:10.1111/jcmm.17700

559. Jiang Y, Hu X, Pang M, Huang Y, Ren B, He L, Jiang L. RRM2Lmediated Wnt/βLcatenin signaling pathway activation in lung adenocarcinoma: A potential prognostic biomarker. Oncol Lett. 2023;26(4):417. doi:10.3892/ol.2023.14003

560. Dong JK, Lei HM, Liang Q, Tang YB, Zhou Y, Wang Y, Zhang S, Li WB, Tong Y, Zhuang G, et al. Overcoming erlotinib resistance in EGFR mutation-positive lung adenocarcinomas through repression of phosphoglycerate dehydrogenase. Theranostics. 2018;8(7):1808–1823. doi:10.7150/thno.23177

561. Peng W, Wen L, Jiang R, Deng J, Chen M. CHAC2 promotes lung adenocarcinoma by regulating ROS-mediated MAPK pathway activation. J Cancer. 2023;14(8):1309–1320. doi:10.7150/jca.84036

562. Guijo M, Ceballos-Chávez M, Gómez-Marín E, Basurto-Cayuela L, Reyes JC. Expression of TDRD9 in a subset of lung carcinomas by CpG island hypomethylation protects from DNA damage. Oncotarget. 2017;9(11):9618–9631. doi:10.18632/oncotarget.22709

563. Yim JH, Yun HS, Lee SJ, Baek JH, Lee CW, Song JY, Um HD, Park JK, Kim JS, Park IC, et al. Radiosensitizing effect of PSMC5, a 19S proteasome ATPase, in H460 lung cancer cells. Biochem Biophys Res Commun. 2016;469(1):94–100. doi:10.1016/j.bbrc.2015.11.077

564. Xing Y, Zhao S, Wei Q, Gong S, Zhao X, Zhou F, Ai-Lamki R, Ortmann D, Du M, Pedersen R, et al. A novel piperidine identified by stem cell-based screening attenuates pulmonary arterial hypertension by regulating BMP2 and PTGS2 levels. Eur Respir J. 2018;51(4):1702229. doi:10.1183/13993003.02229-2017

565. Xu WJ, Wu Q, He WN, Wang S, Zhao YL, Huang JX, Yan XS, Jiang R. Interleukin-6 and pulmonary hypertension: from physiopathology to therapy. Front Immunol. 2023;14:1181987. doi:10.3389/fimmu.2023.1181987

566. Li Z, Jiang J, Gao S. Potential of C-X-C motif chemokine ligand 1/8/10/12 as diagnostic and prognostic biomarkers in idiopathic pulmonary arterial hypertension. Clin Respir J. 2021;15(12):1302–1309. doi:10.1111/crj.13421

567. Ma Y, Chen SS, Jiang F, Ma RY, Wang HL. Bioinformatic analysis and validation of microRNA-508-3p as a protective predictor by targeting NR4A3/MEK axis in pulmonary arterial hypertension. J Cell Mol Med. 2021;25(11):5202–5219. doi:10.1111/jcmm.16523

568. Benincasa G, Maron BA, Affinito O, D’Alto M, Franzese M, Argiento P, Schiano C, Romeo E, Bontempo P, Golino P, et al. Association Between Circulating CD4+ T Cell Methylation Signatures of Network-Oriented SOCS3 Gene and Hemodynamics in Patients Suffering Pulmonary Arterial Hypertension. J Cardiovasc Transl Res. 2023;16(1):17–30. doi:10.1007/s12265-022-10294-1

569. Nie X, Tan J, Dai Y, Liu Y, Zou J, Sun J, Ye S, Shen C, Fan L, Chen J, et al. CCL5 deficiency rescues pulmonary vascular dysfunction, and reverses pulmonary hypertension via caveolin-1-dependent BMPR2 activation. J Mol Cell Cardiol. 2018;116:41–56. doi:10.1016/j.yjmcc.2018.01.016

570. Amsellem V, Abid S, Poupel L, Parpaleix A, Rodero M, Gary-Bobo G, Latiri M, Dubois-Rande JL, Lipskaia L, Combadiere C, et al. Roles for the CX3CL1/CX3CR1 and CCL2/CCR2 Chemokine Systems in Hypoxic Pulmonary Hypertension. Am J Respir Cell Mol Biol. 2017;56(5):597–608. doi:10.1165/rcmb.2016-0201OC

571. Olsson KM, Olle S, Fuge J, Welte T, Hoeper MM, Lerch C, Maegel L, Haller H, Jonigk D, Schiffer L. CXCL13 in idiopathic pulmonary arterial hypertension and chronic thromboembolic pulmonary hypertension. Respir Res. 2016;17:21. doi:10.1186/s12931-016-0336-5

572. Ferguson BS, Wennersten SA, Demos-Davies KM, Rubino M, Robinson EL, Cavasin MA, Stratton MS, Kidger AM, Hu T, Keyse SM, et al. DUSP5-mediated inhibition of smooth muscle cell proliferation suppresses pulmonary hypertension and right ventricular hypertrophy. Am J Physiol Heart Circ Physiol. 2021;321(2):H382–H389. doi:10.1152/ajpheart.00115.2021

573. DeVallance ER, Dustin CM, de Jesus DS, Ghouleh IA, Sembrat JC, Cifuentes-Pagano E, Pagano PJ. Specificity Protein 1-Mediated Promotion of CXCL12 Advances Endothelial Cell Metabolism and Proliferation in Pulmonary Hypertension. Antioxidants (Basel). 2022;12(1):71. doi:10.3390/antiox12010071

574. Powell PP, Klagsbrun M, Abraham JA, Jones RC. Eosinophils expressing heparin-binding EGF-like growth factor mRNA localize around lung microvessels in pulmonary hypertension. Am J Pathol. 1993;143(3):784–793.

575. Florentin J, Zhao J, Tai YY, Sun W, Ohayon LL, O’Neil SP, Arunkumar A, Zhang X, Zhu J, Al Aaraj Y, et al. Loss of Amphiregulin drives inflammation and endothelial apoptosis in pulmonary hypertension. Life Sci Alliance. 2022;5(11):e202101264. doi:10.26508/lsa.202101264

576. Kim J, Kang Y, Kojima Y, Lighthouse JK, Hu X, Aldred MA, McLean DL, Park H, Comhair SA, Greif DM, et al. An endothelial apelin-FGF link mediated by miR-424 and miR-503 is disrupted in pulmonary arterial hypertension. Nat Med. 2013;19(1):74–82. doi:10.1038/nm.3040

577. Kania K, Ahmed A, Ahmed S, Rådegran G. Elevated plasma WIF-1 levels are associated with worse prognosis in heart failure with pulmonary hypertension. ESC Heart Fail. 2022;9(6):4139–4149. doi:10.1002/ehf2.14148

578. Liu Y, Luo Y, Shi X, Lu Y, Li H, Fu G, Li X, Shan L. Role of KLF4/NDRG1/DRP1 axis in hypoxia-induced pulmonary hypertension. Biochim Biophys Acta Mol Basis Dis. 2023;1869(7):166794. doi:10.1016/j.bbadis.2023.166794

579. Swain SD, Siemsen DW, Pullen RR, Han S. CD4+ T cells and IFN-γ are required for the development of Pneumocystis-associated pulmonary hypertension. Am J Pathol. 2014;184(2):483–493. doi:10.1016/j.ajpath.2013.10.027

580. Amsellem V, Abid S, Poupel L, Parpaleix A, Rodero M, Gary-Bobo G, Latiri M, Dubois-Rande JL, Lipskaia L, Combadiere C, et al. Roles for the CX3CL1/CX3CR1 and CCL2/CCR2 Chemokine Systems in Hypoxic Pulmonary Hypertension. Am J Respir Cell Mol Biol. 2017;56(5):597–608. doi:10.1165/rcmb.2016-0201OC

581. Anderson L, Lowery JW, Frank DB, Novitskaya T, Jones M, Mortlock DP, Chandler RL, de Caestecker MP. Bmp2 and Bmp4 exert opposing effects in hypoxic pulmonary hypertension. Am J Physiol Regul Integr Comp Physiol. 2010;298(3):R833–R842. doi:10.1152/ajpregu.00534.2009

582. Wang B, Li C, Huai R, Qu Z. Overexpression of ANO1/TMEM16A, an arterial Ca2+-activated Cl-channel, contributes to spontaneous hypertension. J Mol Cell Cardiol. 2015;82:22–32. doi:10.1016/j.yjmcc.2015.02.020

583. Calvier L, Herz J, Hansmann G. Interplay of Low-Density Lipoprotein Receptors, LRPs, and Lipoproteins in Pulmonary Hypertension. JACC Basic Transl Sci. 2022;7(2):164–180. doi:10.1016/j.jacbts.2021.09.011

584. Shinya Y, Hiraide T, Momoi M, Goto S, Suzuki H, Katsumata Y, Kurebayashi Y, Endo J, Sano M, Fukuda K, et al. TNFRSF13B c.226G>A (p.Gly76Ser) as a Novel Causative Mutation for Pulmonary Arterial Hypertension. J Am Heart Assoc. 2021;10(5):e019245. doi:10.1161/JAHA.120.019245

585. Cai M, Li X, Dong H, Wang Y, Huang X. CCR7 and its related molecules may be potential biomarkers of pulmonary arterial hypertension. FEBS Open Bio. 2021;11(6):1565–1578. doi:10.1002/2211-5463.13130

586. Koudstaal T, van Hulst JAC, Das T, Neys SFH, Merkus D, Bergen IM, de Raaf MA, Bogaard HJ, Boon L, van Loo G, et al. DNGR1-Cre-mediated Deletion of Tnfaip3/A20 in Conventional Dendritic Cells Induces Pulmonary Hypertension in Mice. Am J Respir Cell Mol Biol. 2020;63(5):665–680. doi:10.1165/rcmb.2019-0443OC

587. Jia D, Bai P, Wan N, Liu J, Zhu Q, He Y, Chen G, Wang J, Chen H, Wang C, et al. Niacin Attenuates Pulmonary Hypertension Through H-PGDS in Macrophages. Circ Res. 2020;127(10):1323–1336. doi:10.1161/CIRCRESAHA.120.316784

588. Wang S, Wang Y, Liu C, Xu G, Gao W, Hao J, Zhang M, Wu G, Yang Y, Huang J, et al. EPAS1 (Endothelial PAS Domain Protein 1) Orchestrates Transactivation of Endothelial ICAM1 (Intercellular Adhesion Molecule 1) by Small Nucleolar RNA Host Gene 5 (SNHG5) to Promote Hypoxic Pulmonary Hypertension. Hypertension. 2021;78(4):1080–1091. doi:10.1161/HYPERTENSIONAHA.121.16949

589. Fadini GP, Schiavon M, Cantini M, Avogaro A, Agostini C. Circulating CD34+ cells, pulmonary hypertension, and myelofibrosis. Blood. 2006;108(5):1776–1777. doi:10.1182/blood-2006-02-005892

590. Liu J, Wang W, Wang L, Chen S, Tian B, Huang K, Corrigan CJ, Ying S, Wang W, Wang C. IL-33 Initiates Vascular Remodelling in Hypoxic Pulmonary Hypertension by up-Regulating HIF-1α and VEGF Expression in Vascular Endothelial Cells. EBioMedicine. 2018;33:196–210. doi:10.1016/j.ebiom.2018.06.003

591. Kentera D, Susić D, Zdravković M. Separate and combined use of verapamil, aspirin and captopril in experimental chronic pulmonary hypertension. Basic Res Cardiol. 1984;79(3):375–378. doi:10.1007/BF01908038

592. Tseng V, Collum SD, Allawzi A, Crotty K, Yeligar S, Trammell A, Ryan Smith M, Kang BY, Sutliff RL, Ingram JL, et al. 3’UTR shortening of HAS2 promotes hyaluronan hyper-synthesis and bioenergetic dysfunction in pulmonary hypertension. Matrix Biol. 2022;111:53–75. doi:10.1016/j.matbio.2022.06.001

593. Santulli G, Cipolletta E, Sorriento D, Del Giudice C, Anastasio A, Monaco S, Maione AS, Condorelli G, Puca A, Trimarco B, et al. CaMK4 Gene Deletion Induces Hypertension. J Am Heart Assoc. 2012;1(4):e001081. doi:10.1161/JAHA.112.001081

594. Kumar R, Mickael C, Kassa B, Sanders L, Hernandez-Saavedra D, Koyanagi DE, Kumar S, Pugliese SC, Thomas S, McClendon J, et al. Interstitial macrophage-derived thrombospondin-1 contributes to hypoxia-induced pulmonary hypertension. Cardiovasc Res. 2020;116(12):2021–2030. doi:10.1093/cvr/cvz304

595. Ding XH, Chai X, Zheng J, Chang H, Zheng W, Bian SZ, Ye P. Baseline Ratio of Soluble Fas/FasL Predicts Onset of Pulmonary Hypertension in Elder Patients Undergoing Maintenance Hemodialysis: A Prospective Cohort Study. Front Physiol. 2022;13:847172. doi:10.3389/fphys.2022.847172

596. Pan Z, Wu X, Zhang X, Hu K. Phosphodiesterase 4B activation exacerbates pulmonary hypertension induced by intermittent hypoxia by regulating mitochondrial injury and cAMP/PKA/p-CREB/PGC-1α signaling. Biomed Pharmacother. 2023;158:114095. doi:10.1016/j.biopha.2022.114095

597. Zhou C, Chen Y, Kang W, Lv H, Fang Z, Yan F, Li L, Zhang W, Shi J. Mir-455-3p-1 represses FGF7 expression to inhibit pulmonary arterial hypertension through inhibiting the RAS/ERK signaling pathway. J Mol Cell Cardiol. 2019;130:23–35. doi:10.1016/j.yjmcc.2019.03.002

598. Bell RD, White RJ, Garcia-Hernandez ML, Wu E, Rahimi H, Marangoni RG, Slattery P, Duemmel S, Nuzzo M, Huertas N, et al. Tumor Necrosis Factor Induces Obliterative Pulmonary Vascular Disease in a Novel Model of Connective Tissue Disease-Associated Pulmonary Arterial Hypertension. Arthritis Rheumatol. 2020;72(10):1759–1770. doi:10.1002/art.41309

599. Le Hiress M, Tu L, Ricard N, Phan C, Thuillet R, Fadel E, Dorfmüller P, Montani D, de Man F, Humbert M, et al. Proinflammatory Signature of the Dysfunctional Endothelium in Pulmonary Hypertension. Role of the Macrophage Migration Inhibitory Factor/CD74 Complex. Am J Respir Crit Care Med. 2015;192(8):983–997. doi:10.1164/rccm.201402-0322OC

600. Liu Y, Tian XY, Mao G, Fang X, Fung ML, Shyy JY, Huang Y, Wang N. Response to overexpression of 5-hydroxytryptamine 2B receptor gene in pulmonary hypertension: still a long way to understand its transcriptional regulation. Hypertension. 2013;61(4):e30. doi:10.1161/hypertensionaha.111.00714

601. Wang Q, Chai L, Zhang Q, Wang J, Liu J, Chen H, Wang Y, Chen Y, Shen N, Xie X, et al. Induction of GLI1 by miR-27b-3p/FBXW7/KLF5 pathway contributes to pulmonary arterial hypertension. J Mol Cell Cardiol. 2022;171:16–29. doi:10.1016/j.yjmcc.2022.06.01

602. Pu A, Ramani G, Chen YJ, Perry JA, Hong CC. Identification of novel genetic variants, including PIM1 and LINC01491, with ICD-10 based diagnosis of pulmonary arterial hypertension in the UK Biobank cohort. Front Drug Discov (Lausanne). 2023;3:1127736. doi:10.3389/fddsv.2023.1127736

603. Ding X, Zhou S, Li M, Cao C, Wu P, Sun L, Fei G, Wang R. Upregulation of SRF Is Associated With Hypoxic Pulmonary Hypertension by Promoting Viability of Smooth Muscle Cells via Increasing Expression of Bcl-2. J Cell Biochem. 2017;118(9):2731–2738. doi:10.1002/jcb.25922

604. Jiao YR, Wang W, Lei PC, Jia HP, Dong J, Gou YQ, Chen CL, Cao J, Wang YF, Zhu YK. 5-HTT, BMPR2, EDN1, ENG, KCNA5 gene polymorphisms and susceptibility to pulmonary arterial hypertension: A meta-analysis. Gene. 2019;680:34–42. doi:10.1016/j.gene.2018.09.020

605. Yi D, Liu B, Wang T, Liao Q, Zhu MM, Zhao YY, Dai Z. Endothelial Autocrine Signaling through CXCL12/CXCR4/FoxM1 Axis Contributes to Severe Pulmonary Arterial Hypertension. Int J Mol Sci. 2021;22(6):3182. doi:10.3390/ijms22063182

606. Salguero MV, Chan K, Greeley SAW, Dyamenahalli U, Waggoner D, Del Gaudio D, Rajiyah T, Lemelman M. Novel KDM6A Kabuki Syndrome Mutation With Hyperinsulinemic Hypoglycemia and Pulmonary Hypertension Requiring ECMO. J Endocr Soc. 2022;6(4):bvac015. doi:10.1210/jendso/bvac015

607. Yang L, Liang H, Shen L, Guan Z, Meng X. LncRNA Tug1 involves in the pulmonary vascular remodeling in mice with hypoxic pulmonary hypertension via the microRNA-374c-mediated Foxc1. Life Sci. 2019;237:116769. doi:10.1016/j.lfs.2019.116769

608. Novoyatleva T, Kojonazarov B, Owczarek A, Veeroju S, Rai N, Henneke I, Böhm M, Grimminger F, Ghofrani HA, Seeger W, et al. Evidence for the Fucoidan/P-Selectin Axis as a Therapeutic Target in Hypoxia-induced Pulmonary Hypertension. Am J Respir Crit Care Med. 2019;199(11):1407–1420. doi:10.1164/rccm.201806-1170OC

609. Zhang W, Lin W, Zeng X, Zhang M, Chen Q, Tang Y, Sun J, Liang B, Zha L, Yu Z. FUT8-Mediated Core Fucosylation Promotes the Pulmonary Vascular Remodeling in Pulmonary Arterial Hypertension. Aging Dis. 2023;14(5):1927–1944. doi:10.14336/AD.2023.0218

610. Arvidsson M, Ahmed A, Säleby J, Ahmed S, Hesselstrand R, Rådegran G. Plasma TRAIL and ANXA1 in diagnosis and prognostication of pulmonary arterial hypertension. Pulm Circ. 2023;13(3):e12269. doi:10.1002/pul2.12269

611. Lu G, Du R, Liu Y, Zhang S, Li J, Pei J. RGS5 as a Biomarker of Pericytes, Involvement in Vascular Remodeling and Pulmonary Arterial Hypertension. Vasc Health Risk Manag. 2023;19:673–688. doi:10.2147/VHRM.S429535

612. Bai Z, Xu L, Dai Y, Yuan Q, Zhou Z. ECM2 and GLT8D2 in human pulmonary artery hypertension: fruits from weighted gene co-expression network analysis. J Thorac Dis. 2021;13(4):2242–2254. doi:10.21037/jtd-20-3069

613. Yamada Y, Maruyama J, Zhang E, Okada A, Yokochi A, Sawada H, Mitani Y, Hayashi T, Suzuki K, Maruyama K. Effect of thrombomodulin on the development of monocrotaline-induced pulmonary hypertension. J Anesth. 2014;28(1):26–33. doi:10.1007/s00540-013-1663-z

614. Chi PL, Cheng CC, Hung CC, Wang MT, Liu HY, Ke MW, Shen MC, Lin KC, Kuo SH, Hsieh PP, et al. MMP-10 from M1 macrophages promotes pulmonary vascular remodeling and pulmonary arterial hypertension. Int J Biol Sci. 2022;18(1):331–348. doi:10.7150/ijbs.66472

615. Bellan M, Murano F, Ceruti F, Piccinino C, Tonello S, Minisini R, Giubertoni A, Sola D, Pedrazzoli R, Maglione V, et al. Increased Levels of ICOS and ICOSL Are Associated to Pulmonary Arterial Hypertension in Patients Affected by Connective Tissue Diseases. Diagnostics (Basel). 2022;12(3):704. doi:10.3390/diagnostics12030704

616. Zhu N, Swietlik EM, Welch CL, Pauciulo MW, Hagen JJ, Zhou X, Guo Y, Karten J, Pandya D, Tilly T, et al. Correction to: Rare variant analysis of 4241 pulmonary arterial hypertension cases from an international consortium implicates FBLN2, PDGFD, and rare de novo variants in PAH. Genome Med. 2021;13(1):106. doi:10.1186/s13073-021-00915-w

617. Myllylahti L, Ropponen J, Lax M, Lassila R, Nykänen AI. Upregulation of Coagulation Factor VIII and Fibrinogen After Pulmonary Endarterectomy in Patients with Chronic Thromboembolic Pulmonary Hypertension. Clin Appl Thromb Hemost. 2023;29:10760296231158369. doi:10.1177/10760296231158369

618. Song J, Chen Y, Chen Y, Wang S, Dong Z, Liu X, Li X, Zhang Z, Sun L, Zhong J. Ferrostatin-1 Blunts Right Ventricular Hypertrophy and Dysfunction in Pulmonary Arterial Hypertension by Suppressing the HMOX1/GSH Signaling. J Cardiovasc Transl Res. 2023. doi:10.1007/s12265-023-10423-4

619. Xu Y, Hu T, Ding H, Yuan Y, Chen R. miR-485-5p alleviates obstructive sleep apnea syndrome with hypertension by inhibiting PI3K/AKT signaling pathway via downregulating HIF3A expression. Sleep Breath. 2023;27(1):109–119. doi:10.1007/s11325-022-02580-8

620. Liang KW, Chang SK, Chen YW, Tsai WJ, Wang KY. A cluster of heritable pulmonary arterial hypertension cases in a family with all three siblings carrying the same novel AQP1 c.273C>G variant-a case report. Pulm Circ. 2023;13(2):e12211. doi:10.1002/pul2.12211

621. Cao X, Fang X, Guo M, Li X, He Y, Xie M, Xu Y, Liu X. TRB3 mediates vascular remodeling by activating the MAPK signaling pathway in hypoxic pulmonary hypertension. Respir Res. 2021;22(1):312. doi:10.1186/s12931-021-01908-4

622. Jacob C, Kitagawa A, Signoretti C, Dzieciatkowska M, D’Alessandro A, Gupte A, Hossain S, D’Addario CA, Gupte R, Gupte SA. Mediterranean G6PD variant mitigates expression of DNA methyltransferases and right heart pressure in experimental model of pulmonary hypertension. J Biol Chem. 2022;298(12):102691. doi:10.1016/j.jbc.2022.102691

623. Lago-Docampo M, Tenorio J, Hernández-González I, Pérez-Olivares C, Escribano-Subías P, Pousada G, Baloira A, Arenas M, Lapunzina P, Valverde D. Characterization of rare ABCC8 variants identified in Spanish pulmonary arterial hypertension patients. Sci Rep. 2020;10(1):15135. doi:10.1038/s41598-020-72089-1

624. Li X, Zhang Y, Su L, Cai L, Zhang C, Zhang J, Sun J, Chai M, Cai M, Wu Q, et al. FGF21 alleviates pulmonary hypertension by inhibiting mTORC1/EIF4EBP1 pathway via H19. J Cell Mol Med. 2022;26(10):3005–3021. doi:10.1111/jcmm.17318

625. Wang Q, Tian J, Li X, Liu X, Zheng T, Zhao Y, Li X, Zhong H, Liu D, Zhang W, et al. Upregulation of Endothelial DKK1 (Dickkopf 1) Promotes the Development of Pulmonary Hypertension Through the Sp1 (Specificity Protein 1)/SHMT2 (Serine Hydroxymethyltransferase 2) Pathway. Hypertension. 2022;79(5):960–973. doi:10.1161/HYPERTENSIONAHA.121.18672

626. Patel N, Moenkemeyer F, Germano S, Cheung MM. Plasma vascular endothelial growth factor A and placental growth factor: novel biomarkers of pulmonary hypertension in congenital diaphragmatic hernia. Am J Physiol Lung Cell Mol Physiol. 2015;308(4):L378–L383. doi:10.1152/ajplung.00261.2014

627. Zhang X, Zhang C, Li Q, Piao C, Zhang H, Gu H. Clinical characteristics and prognosis analysis of idiopathic and hereditary pulmonary hypertension patients with ACVRL1 gene mutations. Pulm Circ. 2021;11(4):20458940211044577. doi:10.1177/20458940211044577

628. Singh N, Manhas A, Kaur G, Jagavelu K, Hanif K. Inhibition of fatty acid synthase is protective in pulmonary hypertension. Br J Pharmacol. 2016;173(12):2030–2045. doi:10.1111/bph.13495

629. Laggner M, Hacker P, Oberndorfer F, Bauer J, Raunegger T, Gerges C, Szerafin T, Thanner J, Lang I, Skoro-Sajer N, et al. The Roles of S100A4 and the EGF/EGFR Signaling Axis in Pulmonary Hypertension with Right Ventricular Hypertrophy. Biology (Basel). 2022;11(1):118. doi:10.3390/biology11010118

630. Shen M, Zheng C, Chen L, Li M, Huang X, He M, Liu C, Lin H, Liao W, Bin J, et al. LCZ696 (sacubitril/valsartan) inhibits pulmonary hypertension induced right ventricular remodeling by targeting pyruvate dehydrogenase kinase 4. Biomed Pharmacother. 2023;162:114569. doi:10.1016/j.biopha.2023.114569

631. Jiang J, Xia Y, Liang Y, Yang M, Zeng W, Zeng X. miR-190a-5p participates in the regulation of hypoxia-induced pulmonary hypertension by targeting KLF15 and can serve as a biomarker of diagnosis and prognosis in chronic obstructive pulmonary disease complicated with pulmonary hypertension. Int J Chron Obstruct Pulmon Dis. 2018;13:3777–3790. doi:10.2147/COPD.S182504

632. Shi Y, Liu J, Zhang R, Zhang M, Cui H, Wang L, Cui Y, Wang W, Sun Y, Wang C. Targeting Endothelial ENO1 (Alpha-Enolase) -PI3K-Akt-mTOR Axis Alleviates Hypoxic Pulmonary Hypertension. Hypertension. 2023;80(5):1035–1047. doi:10.1161/HYPERTENSIONAHA.122.19857

633. Zheng Q, Zhang B, Lu N, Li X, Jin B, Jin P. Diagnostic values of serum BNP, PTX3, and VEGF in acute pulmonary embolism complicated by pulmonary artery hypertension and their correlations with severity of pulmonary artery hypertension. Immun Inflamm Dis. 2023;11(9):e986. doi:10.1002/iid3.986

634. Sanges S, Prévotat A, Fertin M, Terriou L, Lefèvre G, Quesnel B, Hatron PY, Hachulla É, Copin MC, Launay D. Haemodynamically proven pulmonary hypertension in a patient with GATA2 deficiency-associated pulmonary alveolar proteinosis and fibrosis. Eur Respir J. 2017;49(5):1700178. doi:10.1183/13993003.00178-2017

635. Mata-Greenwood E, Meyrick B, Soifer SJ, Fineman JR, Black SM. Expression of VEGF and its receptors Flt-1 and Flk-1/KDR is altered in lambs with increased pulmonary blood flow and pulmonary hypertension. Am J Physiol Lung Cell Mol Physiol. 2003;285(1):L222–L231. doi:10.1152/ajplung.00388.2002

636. Ota C, Kimura M, Kure S. ABCA3 mutations led to pulmonary fibrosis and emphysema with pulmonary hypertension in an 8-year-old girl. Pediatr Pulmonol. 2016;51(6):E21–E23. doi:10.1002/ppul.23379

637. Batlahally S, Franklin A, Damianos A, Huang J, Chen P, Sharma M, Duara J, Keerthy D, Zambrano R, Shehadeh LA, et al. Soluble Klotho, a biomarker and therapeutic strategy to reduce bronchopulmonary dysplasia and pulmonary hypertension in preterm infants. Sci Rep. 2020;10(1):12368. doi:10.1038/s41598-020-69296-1

638. Chen R, Wang H, Zheng C, Zhang X, Li L, Wang S, Chen H, Duan J, Zhou X, Peng H, et al. Polo-like kinase 1 promotes pulmonary hypertension. Respir Res. 2023;24(1):204. doi:10.1186/s12931-023-02498-z

639. Tan R, Li J, Peng X, Zhu L, Cai L, Wang T, Su Y, Irani K, Hu Q. APDH is critical for superior efficacy of female bone marrow-derived mesenchymal stem cells on pulmonary hypertension. Cardiovasc Res. 2013;100(1):19–27. doi:10.1093/cvr/cvt165

640. Klinke A, Berghausen E, Friedrichs K, Molz S, Lau D, Remane L, Berlin M, Kaltwasser C, Adam M, Mehrkens D, et al. Myeloperoxidase aggravates pulmonary arterial hypertension by activation of vascular Rho-kinase. JCI Insight. 2018;3(11):e97530. doi:10.1172/jci.insight.97530

641. Ahmed S, Ahmed A, Rådegran G. Plasma tumour and metabolism related biomarkers AMBP, LPL and Glyoxalase I differentiate heart failure with preserved ejection fraction with pulmonary hypertension from pulmonary arterial hypertension. Int J Cardiol. 2021;345:68–76. doi:10.1016/j.ijcard.2021.10.136

642. Yeh FC, Chen CN, Xie CY, Baxan N, Zhao L, Ashek A, Sabrin F, Lawrie A, Wilkins M, Zhao L. TLR7/8 activation induces autoimmune vasculopathy and causes severe pulmonary arterial hypertension. Eur Respir J. 2023;62(1):2300204. doi:10.1183/13993003.00204-2023

643. Cicco S, Leone P, Racanelli V, Vacca A. Mucine-1 Is Related to Cell-Mediated Immunoexpression and Blood Pressure in Pulmonary Artery in Pulmonary Arterial Hypertension (PAH): Preliminary Results. Adv Exp Med Biol. 2018;1072:275–280. doi:10.1007/978-3-319-91287-5_44

644. Isobe S, Nair RV, Kang HY, Wang L, Moonen JR, Shinohara T, Cao A, Taylor S, Otsuki S, Marciano DP, et al. Reduced FOXF1 links unrepaired DNA damage to pulmonary arterial hypertension. Nat Commun. 2023;14(1):7578. doi:10.1038/s41467-023-43039-y

645. Chang YT, Chan CK, Eriksson I, Johnson PY, Cao X, Westöö C, Norvik C, Andersson-Sjöland A, Westergren-Thorsson G, Johansson S, et al. Versican accumulates in vascular lesions in pulmonary arterial hypertension. Pulm Circ. 2016;6(3):347–359. doi:10.1086/686994

646. Jin T, Lu J, Lv Q, Gong Y, Feng Z, Ying H, Wang M, Fu G, Jiang D. Farnesyl diphosphate synthase regulated endothelial proliferation and autophagy during rat pulmonary arterial hypertension induced by monocrotaline. Mol Med. 2022;28(1):94. doi:10.1186/s10020-022-00511-7

647. Martín-Vázquez E, Cobo-Vuilleumier N, López-Noriega L, Lorenzo PI, Gauthier BR. The PTGS2/COX2-PGE2 signaling cascade in inflammation: Pro or anti? A case study with type 1 diabetes mellitus. Int J Biol Sci. 2023;19(13):4157–4165. doi:10.7150/ijbs.86492

648. Li Q, Li Y, Huang W, Wang X, Liu Z, Chen J, Fan Y, Peng T, Sadayappan S, Wang Y, et al. Loss of Lipocalin 10 Exacerbates Diabetes-Induced Cardiomyopathy via Disruption of Nr4a1-Mediated Anti-Inflammatory Response in Macrophages. Front Immunol. 2022;13:930397. doi:10.3389/fimmu.2022.930397

649. Rehman K, Akash MSH, Liaqat A, Kamal S, Qadir MI, Rasul A. Role of Interleukin-6 in Development of Insulin Resistance and Type 2 Diabetes Mellitus. Crit Rev Eukaryot Gene Expr. 2017;27(3):229–236. doi:10.1615/CritRevEukaryotGeneExpr.2017019712

650. Silva BRD, Cirelli T, Nepomuceno R, Theodoro LH, Orrico SRP, Cirelli JA, Barros SP, Scarel-Caminaga RM. Functional haplotype in the Interleukin8 (CXCL8) gene is associated with type 2 Diabetes Mellitus and Periodontitis in Brazilian population. Diabetes Metab Syndr. 2020;14(6):1665–1672. doi:10.1016/j.dsx.2020.08.036

651. Lei L, Cui L, Mao Y, Zhang X, Jiang Q, Dong S, Wang Y. ugmented CD25 and CD69 expression on circulating CD8+ T cells in type 2 diabetes mellitus with albuminuria. Diabetes Metab. 2017;43(4):382–384. doi:10.1016/j.diabet.2016.10.002

652. Takahashi K, Ohara M, Sasai T, Homma H, Nagasawa K, Takahashi T, Yamashina M, Ishii M, Fujiwara F, Kajiwara T, et al. Serum CXCL1 concentrations are elevated in type 1 diabetes mellitus, possibly reflecting activity of anti-islet autoimmune activity. Diabetes Metab Res Rev. 2011;27(8):830–833. doi:10.1002/dmrr.1257

653. Kato H, Nomura K, Osabe D, Shinohara S, Mizumori O, Katashima R, Iwasaki S, Nishimura K, Yoshino M, Kobori M, et al. Association of single-nucleotide polymorphisms in the suppressor of cytokine signaling 2 (SOCS2) gene with type 2 diabetes in the Japanese. Genomics. 2006;87(4):446–458. doi:10.1016/j.ygeno.2005.11.009

654. Zhang Y, Lin C, Chen R, Luo L, Huang J, Liu H, Chen W, Xu J, Yu H, Ding Y. Association analysis of SOCS3, JAK2 and STAT3 gene polymorphisms and genetic susceptibility to type 2 diabetes mellitus in Chinese population. Diabetol Metab Syndr. 2022;14(1):4. doi:10.1186/s13098-021-00774-w

655. Zhong Y, Du G, Liu J, Li S, Lin J, Deng G, Wei J, Huang J. RUNX1 and CCL3 in Diabetes Mellitus-Related Coronary Artery Disease: A Bioinformatics Analysis. Int J Gen Med. 2022;15:955–963. doi:10.2147/IJGM.S350732

656. Alshammary AF, Alshammari AM, Alsobaie SF, Alageel AA, Ali Khan I. Evidence from genetic studies among rs2107538 variant in the CCL5 gene and Saudi patients diagnosed with type 2 diabetes mellitus. Saudi J Biol Sci. 2023;30(6):103658. doi:10.1016/j.sjbs.2023.103658

657. Fadel MM, Abdel Ghaffar FR, Zwain SK, Ibrahim HM, Badr EA. Serum netrin and VCAM-1 as biomarker for Egyptian patients with type IΙ diabetes mellitus. Biochem Biophys Rep. 2021;27:101045. doi:10.1016/j.bbrep.2021.101045

658. Yahya MJ, Ismail PB, Nordin NB, Akim ABM, Yusuf WSBM, Adam NLB, Yusoff MJ. Association of CCL2, CCR5, ELMO1, and IL8 Polymorphism with Diabetic Nephropathy in Malaysian Type 2 Diabetic Patients. Int J Chronic Dis. 2019;2019:2053015. doi:10.1155/2019/2053015

659. Chen C, Lin LY, Chen JW, Chang TT. CXCL5 suppression recovers neovascularization and accelerates wound healing in diabetes mellitus. Cardiovasc Diabetol. 2023;22(1):172. doi:10.1186/s12933-023-01900-w

660. Xu Z, Tong Q, Zhang Z, Wang S, Zheng Y, Liu Q, Qian LB, Chen SY, Sun J, Cai L. Inhibition of HDAC3 prevents diabetic cardiomyopathy in OVE26 mice via epigenetic regulation of DUSP5-ERK1/2 pathway. Clin Sci (Lond). 2017;131(15):1841–1857. doi:10.1042/CS20170064

661. Alagpulinsa DA, Cao JJL, Sobell D, Poznansky MC. Harnessing CXCL12 signaling to protect and preserve functional β-cell mass and for cell replacement in type 1 diabetes. Pharmacol Ther. 2019;193:63–74. doi:10.1016/j.pharmthera.2018.08.011

662. Gutierrez-Aguilar R, Benmezroua Y, Balkau B, Marre M, Helbecque N, Charpentier G, Polychronakos C, Sladek R, Froguel P, Neve B. Minor contribution of SMAD7 and KLF10 variants to genetic susceptibility of type 2 diabetes. Diabetes Metab. 2007;33(5):372–378. doi:10.1016/j.diabet.2007.06.002

663. Nargis T, Kumar K, Ghosh AR, Sharma A, Rudra D, Sen D, Chakrabarti S, Mukhopadhyay S, Ganguly D, Chakrabarti P. KLK5 induces shedding of DPP4 from circulatory Th17 cells in type 2 diabetes. Mol Metab. 2017;6(11):1529–1539. doi:10.1016/j.molmet.2017.09.004

664. Wong S, Moore S, Orisio S, Millward A, Demaine AG. Susceptibility to type I diabetes in women is associated with the CD3 epsilon locus on chromosome 11. Clin Exp Immunol. 1991;83(1):69–73. doi:10.1111/j.1365-2249.1991.tb05590.x

665. Aparicio JM, Wakisaka A, Takada A, Matsuura N, Yoshiki T. Non-HLA genetic factors and insulin dependent diabetes mellitus in the Japanese: TCRA, TCRB and TCRG, INS, THY1, CD3D and ETS1. Dis Markers. 1990;8(5):283–294.

666. Jiao J, Wang Z, Guo Y, Liu J, Huang X, Ni X, Gao D, Sun L, Zhu X, Zhou Q, et al. Association between IL-1B (-511)/IL-1RN (VNTR) polymorphisms and type 2 diabetes: a systematic review and meta-analysis. PeerJ. 2021;9:e12384. doi:10.7717/peerj.12384

667. Raugh A, Jing Y, Bettini ML, Bettini M. The amphiregulin/EGFR axis has limited contribution in controlling autoimmune diabetes. Sci Rep. 2023;13(1):18653. doi:10.1038/s41598-023-45738-4

668. Fan Q, Li H, Qin Y, Li L, Chen L, Zhang L, Lv Y, Liang D, Liang Y, Long T, et al. Association of SERPINE1 rs6092 with type 2 diabetes and related metabolic traits in a Chinese population. Gene. 2018;661:176–181. doi:10.1016/j.gene.2018.04.011

669. Ikeda S, Sato K, Takeda M, Miki K, Aizawa K, Takada T, Fukuda K, Shiba N. Oncostatin M is a novel biomarker for coronary artery disease - A possibility as a screening tool of silent myocardial ischemia for diabetes mellitus. Int J Cardiol Heart Vasc. 2021;35:100829. doi:10.1016/j.ijcha.2021.100829

670. Juhas U, Ryba-Stanisławowska M, Ławrynowicz U, Myśliwiec M, Myśliwska J. Putative loss of CD83 immunosuppressive activity in long-standing complication-free juvenile diabetic patients during disease progression. Immunol Res. 2019;67(1):70–76. doi:10.1007/s12026-019-09074-y

671. Blindbæk SL, Schlosser A, Green A, Holmskov U, Sorensen GL, Grauslund J. Association between microfibrillar-associated protein 4 (MFAP4) and micro-and macrovascular complications in long-term type 1 diabetes mellitus. Acta Diabetol. 2017;54(4):367–372. doi:10.1007/s00592-016-0953-y

672. Elsehmawy AAEW, El-Toukhy SE, Seliem NMA, Moustafa RS, Mohammed DS. Apelin and chemerin as promising adipokines in children with type 1 diabetes mellitus. Diabetes Metab Syndr Obes. 2019;12:383–389. doi:10.2147/DMSO.S189264

673. Zhao K, Ding W, Zhang Y, Ma K, Wang D, Hu C, Liu J, Zhang X. Variants in the RARRES2 gene are associated with serum chemerin and increase the risk of diabetic kidney disease in type 2 diabetes. Int J Biol Macromol. 2020;165(Pt A):1574–1580. doi:10.1016/j.ijbiomac.2020.10.030

674. Russo L, Muir L, Geletka L, Delproposto J, Baker N, Flesher C, O’Rourke R, Lumeng CN. Cholesterol 25-hydroxylase (CH25H) as a promoter of adipose tissue inflammation in obesity and diabetes. Mol Metab. 2020;39:100983. doi:10.1016/j.molmet.2020.100983

675. Li Y, Yang Y, Wang J, Cai P, Li M, Tang X, Tan Y, Wang Y, Zhang F, Wen X, et al. Bacteroides ovatus-mediated CD27-MAIT cell activation is associated with obesity-related T2D progression. Cell Mol Immunol. 2022;19(7):791–804. doi:10.1038/s41423-022-00871-4

676. Miranda M, Chacón MR, Gutiérrez C, Vilarrasa N, Gómez JM, Caubet E, Megía A, Vendrell J. LMNA mRNA expression is altered in human obesity and type 2 diabetes. Obesity (Silver Spring). 2008;16(8):1742–1748. doi:10.1038/oby.2008.276

677. Wang Y, Zhang R, Shen H, Kong J, Lv X. Pioglitazone protects blood vessels through inhibition of the apelin signaling pathway by promoting KLF4 expression in rat models of T2DM. Biosci Rep. 2019;39(12):BSR20190317. doi:10.1042/BSR20190317

678. Jiao J, Wang Z, Guo Y, Liu J, Huang X, Ni X, Gao D, Sun L, Zhu X, Zhou Q, et al. Association between IL-1B (-511)/IL-1RN (VNTR) polymorphisms and type 2 diabetes: a systematic review and meta-analysis. PeerJ. 2021;9:e12384. doi:10.7717/peerj.12384

679. Stalenhoef JE, Alisjahbana B, Nelwan EJ, van der Ven-Jongekrijg J, Ottenhoff TH, van der Meer JW, Nelwan RH, Netea MG, van Crevel R. The role of interferon-gamma in the increased tuberculosis risk in type 2 diabetes mellitus. Eur J Clin Microbiol Infect Dis. 2008;27(2):97–103. doi:10.1007/s10096-007-0395-0

680. Habibe JJ, Clemente-Olivo MP, Scheithauer TPM, Rampanelli E, Herrema H, Vos M, Mieremet A, Nieuwdorp M, van Raalte DH, Eringa EC, et al. Glucose-mediated insulin secretion is improved in FHL2-deficient mice and elevated FHL2 expression in humans is associated with type 2 diabetes. Diabetologia. 2022;65(10):1721–1733. doi:10.1007/s00125-022-05750-1

681. Sindhu S, Akhter N, Arefanian H, Al-Roub AA, Ali S, Wilson A, Al-Hubail A, Al-Beloushi S, Al-Zanki S, Ahmad R. Increased circulatory levels of fractalkine (CX3CL1) are associated with inflammatory chemokines and cytokines in individuals with type-2 diabetes. J Diabetes Metab Disord. 2017;16:15. doi:10.1186/s40200-017-0297-3

682. Mollah ZUA, Quah HS, Graham KL, Jhala G, Krishnamurthy B, Dharma JFM, Chee J, Trivedi PM, Pappas EG, Mackin L, et al. Granzyme A Deficiency Breaks Immune Tolerance and Promotes Autoimmune Diabetes Through a Type I Interferon-Dependent Pathway. Diabetes. 2017;66(12):3041–3050. doi:10.2337/db17-0517

683. Zhang JM, Yu RQ, Wu FZ, Qiao L, Wu XR, Fu YJ, Liang YF, Pang Y, Xie CY. BMP-2 alleviates heart failure with type 2 diabetes mellitus and doxorubicin-induced AC16 cell injury by inhibiting NLRP3 inflammasome-mediated pyroptosis. Exp Ther Med. 2021;22(2):897. doi:10.3892/etm.2021.10329

684. Willecke F, Yuan C, Oka K, Chan L, Hu Y, Barnhart S, Bornfeldt KE, Goldberg IJ, Fisher EA. Effects of High Fat Feeding and Diabetes on Regression of Atherosclerosis Induced by Low-Density Lipoprotein Receptor Gene Therapy in LDL Receptor-Deficient Mice. PLoS One. 2015;10(6):e0128996. doi:10.1371/journal.pone.0128996

685. Muhammad SA, Qousain Naqvi ST, Nguyen T, Wu X, Munir F, Jamshed MB, Zhang Q. Cisplatin’s potential for type 2 diabetes repositioning by inhibiting CDKN1A, FAS, and SESN1. Comput Biol Med. 2021;135:104640. doi:10.1016/j.compbiomed.2021.104640

686. Villasenor A, Wang ZV, Rivera LB, Ocal O, Asterholm IW, Scherer PE, Brekken RA, Cleaver O, Wilkie TM. Rgs16 and Rgs8 in embryonic endocrine pancreas and mouse models of diabetes. Dis Model Mech. 2010;3(9-10):567–580. doi:10.1242/dmm.003210

687. Cao C, Fu X, Wang X. Case Report: A novel mutation in TNFAIP3 in a patient with type 1 diabetes mellitus and haploinsufficiency of A20. Front Endocrinol (Lausanne). 2023;14:1131437. doi:10.3389/fendo.2023.1131437

688. Liang C, Sun R, Xu Y, Geng W, Li J. Effect of the Abnormal Expression of BMP-4 in the Blood of Diabetic Patients on the Osteogenic Differentiation Potential of Alveolar BMSCs and the Rescue Effect of Metformin: A Bioinformatics-Based Study. Biomed Res Int. 2020;2020:7626215. doi:10.1155/2020/7626215

689. Yao Y, Du J, Li R, Zhao L, Luo N, Zhai JY, Long L. Association between ICAM-1 level and diabetic retinopathy: a review and meta-analysis. Postgrad Med J. 2019;95(1121):162–168. doi:10.1136/postgradmedj-2018-136102

690. Melnik BC. Synergistic Effects of Milk-Derived Exosomes and Galactose on α-Synuclein Pathology in Parkinson’s Disease and Type 2 Diabetes Mellitus. Int J Mol Sci. 2021;22(3):1059. doi:10.3390/ijms22031059

691. Pala C, Altun I, Koker Y, Kurnaz F, Sivgin S, Koçyiğit I, Tanrıverdi F, Kaynar L, Elmali F, Cetin M, et al. The effect of diabetes mellitus and end-stage renal disease on the number of CD34+ cells in the blood. Ann Hematol. 2013;92(9):1189–1194. doi:10.1007/s00277-013-1760-y

692. Zhang K, Yang J, Ao N, Jin S, Qi R, Shan F, Du J. Methionine enkephalin (MENK) regulates the immune pathogenesis of type 2 diabetes mellitus via the IL-33/ST2 pathway. Int Immunopharmacol. 2019;73:23–40. doi:10.1016/j.intimp.2019.04.054

693. Pearl-Yafe M, Yolcu ES, Yaniv I, Stein J, Shirwan H, Askenasy N. The dual role of Fas-ligand as an injury effector and defense strategy in diabetes and islet transplantation. Bioessays. 2006;28(2):211–222. doi:10.1002/bies.20356

694. Alsters SI, Goldstone AP, Buxton JL, Zekavati A, Sosinsky A, Yiorkas AM, Holder S, Klaber RE, Bridges N, van Haelst MM, et al. Truncating Homozygous Mutation of Carboxypeptidase E (CPE) in a Morbidly Obese Female with Type 2 Diabetes Mellitus, Intellectual Disability and Hypogonadotrophic Hypogonadism. PLoS One. 2015;10(6):e0131417. doi:10.1371/journal.pone.0131417

695. Raza W, Guo J, Qadir MI, Bai B, Muhammad SA. qPCR Analysis Reveals Association of Differential Expression of SRR, NFKB1, and PDE4B Genes With Type 2 Diabetes Mellitus. Front Endocrinol (Lausanne). 2022;12:774696. doi:10.3389/fendo.2021.774696

696. Alwahsh SM, Qutachi O, Starkey Lewis PJ, Bond A, Noble J, Burgoyne P, Morton N, Carter R, Mann J, Ferreira-Gonzalez S, et al. Fibroblast growth factor 7 releasing particles enhance islet engraftment and improve metabolic control following islet transplantation in mice with diabetes. Am J Transplant. 2021;21(9):2950–2963. doi:10.1111/ajt.16488

697. Kawabata Y, Nishida N, Awata T, Kawasaki E, Imagawa A, Shimada A, Osawa H, Tanaka S, Takahashi K, Nagata M, et al. Genome-Wide Association Study Confirming a Strong Effect of HLA and Identifying Variants in CSAD/lnc-ITGB7-1 on Chromosome 12q13.13 Associated With Susceptibility to Fulminant Type 1 Diabetes. Diabetes. 2019;68(3):665–675. doi:10.2337/db18-0314

698. Zhang Y, Cao Y, Zheng R, Xiong Z, Zhu Z, Gao F, Man W, Duan Y, Lin J, Zhang X, et al. Fibroblast-specific activation of Rnd3 protects against cardiac remodeling in diabetic cardiomyopathy via suppression of Notch and TGF-β signaling. Theranostics. 2022;12(17):7250–7266. doi:10.7150/thno.77043

699. Akash MSH, Rehman K, Liaqat A. Tumor Necrosis Factor-Alpha: Role in Development of Insulin Resistance and Pathogenesis of Type 2 Diabetes Mellitus. J Cell Biochem. 2018;119(1):105–110. doi:10.1002/jcb.26174

700. Cheng YS, Dai DZ, Ji H, Zhang Q, Dai Y. Sildenafil and FDP-Sr attenuate diabetic cardiomyopathy by suppressing abnormal expression of myocardial CASQ2, FKBP12.6, and SERCA2a in rats. Acta Pharmacol Sin. 2011;32(4):441–448. doi:10.1038/aps.2010.226

701. Al-Kuraishy HM, Al-Gareeb AI, Alexiou A, Papadakis M, Nadwa EH, Albogami SM, Alorabi M, Saad HM, Batiha GE. Metformin and growth differentiation factor 15 (GDF15) in type 2 diabetes mellitus: A hidden treasure. J Diabetes. 2022;14(12):806–814. doi:10.1111/1753-0407.13334

702. Guimarães JB, Rodrigues VF, Pereira ÍS, Manso GMDC, Elias-Oliveira J, Leite JA, Waldetario MCGM, de Oliveira S, Gomes ABDSP, Faria AMC, et al. Inulin prebiotic ameliorates type 1 diabetes dictating regulatory T cell homing via CCR4 to pancreatic islets and butyrogenic gut microbiota in murine model. J Leukoc Biol. 2023. doi:10.1093/jleuko/qiad132

703. Chen L, Yin Z, Qin X, Zhu X, Chen X, Ding G, Sun D, Wu NN, Fei J, Bi Y, et al. CD74 ablation rescues type 2 diabetes mellitus-induced cardiac remodeling and contractile dysfunction through pyroptosis-evoked regulation of ferroptosis. Pharmacol Res. 2022;176:106086. doi:10.1016/j.phrs.2022.106086

704. Long SA, Buckner JH, Greenbaum CJ. IL-2 therapy in type 1 diabetes: “Trials” and tribulations. Clin Immunol. 2013;149(3):324–331. doi:10.1016/j.clim.2013.02.005

705. Khan R, Kadamkode V, Kesharwani D, Purkayastha S, Banerjee G, Datta M. Circulatory miR-98-5p levels are deregulated during diabetes and it inhibits proliferation and promotes apoptosis by targeting PPP1R15B in keratinocytes. RNA Biol. 2020;17(2):188–201. doi:10.1080/15476286.2019.1673117

706. Fallahi P, Corrado A, Di Domenicantonio A, Frenzilli G, Antonelli A, Ferrari SM. CXCR3, CXCR5, CXCR6, and CXCR7 in Diabetes. Curr Drug Targets. 2016;17(5):515–519. doi:10.2174/1389450115666141229153949

707. Choi WG, Choi W, Oh TJ, Cha HN, Hwang I, Lee YK, Lee SY, Shin H, Lim A, Ryu D, et al. nhibiting serotonin signaling through HTR2B in visceral adipose tissue improves obesity-related insulin resistance. J Clin Invest. 2021;131(23):e145331. doi:10.1172/JCI145331

708. You S, Xu J, Yin Z, Wu B, Wang P, Hao M, Cheng C, Liu M, Zhao Y, Jia P, et al. Down-regulation of WWP2 aggravates Type 2 diabetes mellitus-induced vascular endothelial injury through modulating ubiquitination and degradation of DDX3X. Cardiovasc Diabetol. 2023;22(1):107. doi:10.1186/s12933-023-01818-3

709. Kyriazis ID, Hoffman M, Gaignebet L, Lucchese AM, Markopoulou E, Palioura D, Wang C, Bannister TD, Christofidou-Solomidou M, Oka SI, et al. KLF5 Is Induced by FOXO1 and Causes Oxidative Stress and Diabetic Cardiomyopathy. Circ Res. 2021;128(3):335–357. doi:10.1161/CIRCRESAHA.120.316738

710. Colli ML, Hill JLE, Marroquí L, Chaffey J, Dos Santos RS, Leete P, Coomans de Brachène A, Paula FMM, Op de Beeck A, Castela A, et al. PDL1 is expressed in the islets of people with type 1 diabetes and is up-regulated by interferons-α and-γ via IRF1 induction. EBioMedicine. 2018;36:367–375. doi:10.1016/j.ebiom.2018.09.040

711. Xia X, Liang Y, Zheng W, Lin D, Sun S. miR-410-5p promotes the development of diabetic cardiomyopathy by suppressing PIM1-induced anti-apoptosis. Mol Cell Probes. 2020;52:101558. doi:10.1016/j.mcp.2020.101558

712. Shim JY, Chung JO, Jung D, Kang PS, Park SY, Kendi AT, Lowe VJ, Lee S. Parkin-mediated mitophagy is negatively regulated by FOXO3A, which inhibits Plk3-mediated mitochondrial ROS generation in STZ diabetic stress-treated pancreatic β cells. PLoS One. 2023;18(5):e0281496. doi:10.1371/journal.pone.0281496

713. Akazawa S, Kobayashi M, Kuriya G, Horie I, Yu L, Yamasaki H, Okita M, Nagayama Y, Matsuyama T, Akbari M, et al. Haploinsufficiency of interferon regulatory factor 4 strongly protects against autoimmune diabetes in NOD mice. Diabetologia. 2015;58(11):2606–2614. doi:10.1007/s00125-015-3724-3

714. Zhu Y, Yang X, Zhou J, Chen L, Zuo P, Chen L, Jiang L, Li T, Wang D, Xu Y, et al. miR-340-5p Mediates Cardiomyocyte Oxidative Stress in Diabetes-Induced Cardiac Dysfunction by Targeting Mcl-1. Oxid Med Cell Longev. 2022;2022:3182931. doi:10.1155/2022/3182931

715. Liu L, Sun K, Luo Y, Wang B, Yang Y, Chen L, Zheng S, Wu T, Xiao P. Myocardin-related transcription factor A, regulated by serum response factor, contributes to diabetic cardiomyopathy in mice. Life Sci. 2023;317:121470. doi:10.1016/j.lfs.2023.121470

716. Shimizu I, Yoshida Y, Moriya J, Nojima A, Uemura A, Kobayashi Y, Minamino T. Semaphorin3E-induced inflammation contributes to insulin resistance in dietary obesity. Cell Metab. 2013;18(4):491–504. doi:10.1016/j.cmet.2013.09.001

717. Biswas S, Feng B, Thomas A, Chen S, Aref-Eshghi E, Sadikovic B, Chakrabarti S. Endothelin-1 regulation is entangled in a complex web of epigenetic mechanisms in diabetes. Physiol Res. 2018;67(Suppl 1):S115–S125. doi:10.33549/physiolres.933836

718. Zhou C, Wang F, Ma H, Xing N, Hou L, Du Y, Ding H. Silencing of FOS-like antigen 1 represses restenosis via the ERK/AP-1 pathway in type 2 diabetic mice. Diab Vasc Dis Res. 2021;18(6):14791641211058855. doi:10.1177/14791641211058855

719. He YL, Chen MT, Wang T, Zhang MM, Li YX, Wang HY, Ding N. Development of FABP4/5 inhibitors with potential therapeutic effect on type 2 Diabetes Mellitus. Eur J Med Chem. 2021;224:113720. doi:10.1016/j.ejmech.2021.113720

720. Li JY, Tao F, Wu XX, Tan YZ, He L, Lu H. Common RASGRP1 Gene Variants That Confer Risk of Type 2 Diabetes. Genet Test Mol Biomarkers. 2015;19(8):439–443. doi:10.1089/gtmb.2015.0005

721. Austin-Zimmerman I, Wronska M, Wang B, Irizar H, Thygesen JH, Bhat A, Denaxas S, Fatemifar G, Finan C, Harju-Seppänen J, et al. The Influence of CYP2D6 and CYP2C19 Genetic Variation on Diabetes Mellitus Risk in People Taking Antidepressants and Antipsychotics. Genes (Basel). 2021;12(11):1758. doi:10.3390/genes12111758

722. Mayorga ME, Kiedrowski M, McCallinhart P, Forudi F, Ockunzzi J, Weber K, Chilian W, Penn MS, Dong F. Role of SDF-1:CXCR4 in Impaired Post-Myocardial Infarction Cardiac Repair in Diabetes. Stem Cells Transl Med. 2018;7(1):115–124. doi:10.1002/sctm.17-0172

723. Guo M, Xia Z, Hong Y, Ji H, Li F, Liu W, Li S, Xin H, Tan K, Lian Z. The TFPI2-PPARγ axis induces M2 polarization and inhibits fibroblast activation to promote recovery from post-myocardial infarction in diabetic mice. J Inflamm (Lond). 2023;20(1):35. doi:10.1186/s12950-023-00357-8

724. Gautheron J, Morisseau C, Chung WK, Zammouri J, Auclair M, Baujat G, Capel E, Moulin C, Wang Y, Yang J, et al. EPHX1 mutations cause a lipoatrophic diabetes syndrome due to impaired epoxide hydrolysis and increased cellular senescence. Elife. 2021;10:e68445. doi:10.7554/eLife.68445

725. Alzahrani FM, Alhassan JA, Alshehri AM, Farooqi FA, Aldossary MA, Abdelghany MK, Ibrahim H, El-Masry OS. The impact of SELP gene Thr715Pro polymorphism on sP-selectin level and association with cardiovascular disease in Saudi diabetic patients: A cross-sectional case-control study. Saudi J Biol Sci. 2023;30(3):103579. doi:10.1016/j.sjbs.2023.103579

726. Ibarra-Tapia IY, Juárez-Sandoval A, Pérez IT, Cano-Martínez LJ, Sánchez-García S, Ruiz-Batalla JM, Aroche-Reyes IA, García S, Canto P, Mejía DR, et al. Association of polymorphisms rs2303729, rs10880, and rs1131620 of LTBP4 with sarcopenia in elderly patients with type 2 diabetes mellitus. Ann Hum Biol. 2022;49(7-8):311–316. doi:10.1080/03014460.2022.2152489

727. Gu HF. Genetic variation screening and association studies of the adenylate cyclase activating polypeptide 1 (ADCYAP1) gene in patients with type 2 diabetes. Hum Mutat. 2002;19(5):572–573. doi:10.1002/humu.9034

728. Tang D, Liu L, Ajiakber D, Ye J, Xu J, Xin X, Aisa HA. Anti-diabetic Effect of Punica granatum Flower Polyphenols Extract in Type 2 Diabetic Rats: Activation of Akt/GSK-3β and Inhibition of IRE1α-XBP1 Pathways. Front Endocrinol (Lausanne). 2018;9:586. doi:10.3389/fendo.2018.00586

729. Purvis GSD, Collino M, Loiola RA, Baragetti A, Chiazza F, Brovelli M, Sheikh MH, Collotta D, Cento A, Mastrocola R, et al. Identification of AnnexinA1 as an Endogenous Regulator of RhoA, and Its Role in the Pathophysiology and Experimental Therapy of Type-2 Diabetes. Front Immunol. 2019;10:571. doi:10.3389/fimmu.2019.00571

730. Chen F, Lai J, Zhu Y, He M, Hou H, Wang J, Chen C, Wang DW, Tang J. Cardioprotective Effect of Decorin in Type 2 Diabetes. Front Endocrinol (Lausanne). 2020;11:479258. doi:10.3389/fendo.2020.479258

731. Deng W, Wang X, Xiao J, Chen K, Zhou H, Shen D, Li H, Tang Q. Loss of regulator of G protein signaling 5 exacerbates obesity, hepatic steatosis, inflammation and insulin resistance. PLoS One. 2012;7(1):e30256. doi:10.1371/journal.pone.0030256

732. Smith MJ, Pastor L, Newman JRB, Concannon P. Genetic Control of Splicing at SIRPG Modulates Risk of Type 1 Diabetes. Diabetes. 2022;71(2):350–358. doi:10.2337/db21-0194

733. Rochmah N, Arief F, Faizi M, Basuki S. The association between PTPN22 C1858T gene polymorphism and type 1 diabetes mellitus: an Indonesian study. Ann Med. 2023;55(1):1211–1215. doi:10.1080/07853890.2023.2190162

734. Strieder-Barboza C, Flesher CG, Geletka LM, Eichler T, Akinleye O, Ky A, Ehlers AP, Lumeng CN, O’Rourke RW. Lumican modulates adipocyte function in obesity-associated type 2 diabetes. Adipocyte. 2022;11(1):665–675. doi:10.1080/21623945.2022.2154112

735. Zhong W, Dong YJ, Hong C, Li YH, Xiao CX, Liu XH, Chang J. ASH2L upregulation contributes to diabetic endothelial dysfunction in mice through STEAP4-mediated copper uptake. Acta Pharmacol Sin. 2023. doi:10.1038/s41401-023-01174-8

736. Pang A, Hu Y, Zhou P, Long G, Tian X, Men L, Shen Y, Liu Y, Cui Y. Corin is down-regulated and exerts cardioprotective action via activating pro-atrial natriuretic peptide pathway in diabetic cardiomyopathy. Cardiovasc Diabetol. 2015;14:134. doi:10.1186/s12933-015-0298-9

737. Alharbi KK, Ali Khan I, Syed R, Alharbi FK, Mohammed AK, Vinodson B, Al-Daghri NM. Association of JAZF1 and TSPAN8/LGR5 variants in relation to type 2 diabetes mellitus in a Saudi population. Diabetol Metab Syndr. 2015;7:92. doi:10.1186/s13098-015-0091-7

738. Abasheva D, Dolcet-Negre MM, Fernández-Seara MA, Mora-Gutiérrez JM, Orbe J, Escalada FJ, Garcia-Fernandez N. Association between Circulating Levels of 25-Hydroxyvitamin D3 and Matrix Metalloproteinase-10 (MMP-10) in Patients with Type 2 Diabetes. Nutrients. 2022;14(17):3484. doi:10.3390/nu14173484

739. Zhang HY, Ruan LB, Li Y, Yang TR, Liu WJ, Jiang YX, Li TR, Quan J, Xuan W. ICOS/ICOSL upregulation mediates inflammatory response and endothelial dysfunction in type 2 diabetes mellitus. Eur Rev Med Pharmacol Sci. 2018;22(24):8898–8908. doi:10.26355/eurrev_201812_16659

740. Krogh SS, Holt CB, Steffensen R, Funck KL, Høyem P, Laugesen E, Poulsen PL, Thiel S, Hansen TK. Plasma levels of MASP-1, MASP-3 and MAp44 in patients with type 2 diabetes: influence of glycaemic control, body composition and polymorphisms in the MASP1 gene. Clin Exp Immunol. 2017;189(1):103–112. doi:10.1111/cei.12963

741. Babić N, Dervisević A, Huskić J, Musić M. Coagulation factor VIII activity in diabetic patients. Med Glas (Zenica). 2011;8(1):134–139.

742. Ramos-Lopez E, Ghebru S, Van Autreve J, Aminkeng F, Herwig J, Seifried E, Seidl C, Van der Auwera B, Badenhoop K. Neither an intronic CA repeat within the CD48 gene nor the HERV-K18 polymorphisms are associated with type 1 diabetes. Tissue Antigens. 2006;68(2):147–152. doi:10.1111/j.1399-0039.2006.00637.x

743. Santin I, Castellanos-Rubio A, Aransay AM, Gutierrez G, Gaztambide S, Rica I, Vicario JL, Noble JA, Castaño L, Bilbao JR. Exploring the diabetogenicity of the HLA-B18-DR3 CEH: independent association with T1D genetic risk close to HLA-DOA. Genes Immun. 2009;10(6):596–600. doi:10.1038/gene.2009.41

744. Pan X, Liu C, Wang X, Zhao M, Zhang Z, Zhang X, Wang C, Song G. Resveratrol improves palmitic acidLinduced insulin resistance via the DDIT4/mTOR pathway in C2C12 cells. Mol Med Rep. 2023;28(4):181. doi:10.3892/mmr.2023.13068

745. Kawakami R, Katsuki S, Travers R, Romero DC, Becker-Greene D, Passos LSA, Higashi H, Blaser MC, Sukhova GK, Buttigieg J, et al. S100A9-RAGE Axis Accelerates Formation of Macrophage-Mediated Extracellular Vesicle Microcalcification in Diabetes Mellitus. Arterioscler Thromb Vasc Biol. 2020;40(8):1838–1853. doi:10.1161/ATVBAHA.118.314087

746. Dance A, Fernandes J, Toussaint B, Vaillant E, Boutry R, Baron M, Loiselle H, Balkau B, Charpentier G, Franc S, et al. xploring the role of purinergic receptor P2RY1 in type 2 diabetes risk and pathophysiology: Insights from human functional genomics. Mol Metab. 2023. doi:10.1016/j.molmet.2023.101867

747. Sidibeh CO, Pereira MJ, Abalo XM, J Boersma G, Skrtic S, Lundkvist P, Katsogiannos P, Hausch F, Castillejo-López C, Eriksson JW. FKBP5 expression in human adipose tissue: potential role in glucose and lipid metabolism, adipogenesis and type 2 diabetes. Endocrine. 2018;62(1):116–128. doi:10.1007/s12020-018-1674-5

748. Meng Z, Liang H, Zhao J, Gao J, Liu C, Ma X, Liu J, Liang B, Jiao X, Cao J, et al. HMOX1 upregulation promotes ferroptosis in diabetic atherosclerosis. Life Sci. 2021;284:119935. doi:10.1016/j.lfs.2021.119935

749. Liang C, Liu X, Liu C, Xu Y, Geng W, Li J. Integrin α10 regulates adhesion, migration, and osteogenic differentiation of alveolar bone marrow mesenchymal stem cells in type 2 diabetic patients who underwent dental implant surgery. Bioengineered. 2022;13(5):13252–13268. doi:10.1080/21655979.2022.2079254

750. Fawad Ali Shah S, Iqbal T, Naveed N, Akram S, Arshad Rafiq M, Hussain S. ARG1 single nucleotide polymorphisms rs2781666 and rs2781665 confer risk of Type 2 diabetes mellitus. EXCLI J. 2018;17:847–855. doi:10.17179/excli2018-1178

751. Guo Y, Zou J, Xu X, Zhou H, Sun X, Wu L, Zhang S, Zhong X, Xiong Z, Lin Y, et al. Short-chain fatty acids combined with intronic DNA methylation of HIF3A: Potential predictors for diabetic cardiomyopathy. Clin Nutr. 2021;40(6):3708–3717. doi:10.1016/j.clnu.2021.04.007

752. Madonna R, De Caterina R. Aquaporin-1 and sodium-hydrogen exchangers as pharmacological targets in diabetic atherosclerosis. Curr Drug Targets. 2015;16(4):361–365. doi:10.2174/1389450116666141219115720

753. Lu G, Li J, Gao T, Liu Q, Chen O, Zhang X, Xiao M, Guo Y, Wang J, Tang Y, et al. Integration of dietary nutrition and TRIB3 action into diabetes mellitus. Nutr Rev. 2023. doi:10.1093/nutrit/nuad056

754. Karadsheh NS, Quttaineh NA, Karadsheh SN, El-Khateeb M. Effect of combined G6PD deficiency and diabetes on protein oxidation and lipid peroxidation. BMC Endocr Disord. 2021;21(1):246. doi:10.1186/s12902-021-00911-6

755. Li M, Han X, Ji L. Clinical and Genetic Characteristics of ABCC8 Nonneonatal Diabetes Mellitus: A Systematic Review. J Diabetes Res. 2021;2021:9479268. doi:10.1155/2021/9479268

756. Kavian Z, Sargazi S, Majidpour M, Sarhadi M, Saravani R, Shahraki M, Mirinejad S, Heidari Nia M, Piri M. Association of SLC11A1 polymorphisms with anthropometric and biochemical parameters describing Type 2 Diabetes Mellitus. Sci Rep. 2023;13(1):6195. doi:10.1038/s41598-023-33239-3

757. Saulnier PJ, Roussel R, Halimi JM, Lebrec J, Dardari D, Maimaitiming S, Guilloteau G, Prugnard X, Marechaud R, Ragot S, et al. Impact of natriuretic peptide clearance receptor (NPR3) gene variants on blood pressure in type 2 diabetes. Diabetes Care. 2011;34(5):1199–1204. doi:10.2337/dc10-2057

758. Xu L, Sun X, Xia Y, Luo S, Lin J, Xiao Y, Liu Y, Wang Y, Huang G, Li X, et al. Polymorphisms of the NLRC4 Gene are Associated with the Onset Age, Positive Rate of GADA and 2-h Postprandial C-Peptide in Patients with Type 1 Diabetes. Diabetes Metab Syndr Obes. 2020;13:811–818. doi:10.2147/DMSO.S244882

759. Wang X, Ding Y, Zhang X, Rao J, Yu H, Pan H. The association between HHEX single-nucleotide polymorphism rs5015480 and gestational diabetes mellitus: A meta-analysis. Medicine (Baltimore). 2020;99(12):e19478. doi:10.1097/MD.0000000000019478

760. Simonis-Bik AM, Nijpels G, van Haeften TW, Houwing-Duistermaat JJ, Boomsma DI, Reiling E, van Hove EC, Diamant M, Kramer MH, Heine RJ, et al. Gene variants in the novel type 2 diabetes loci CDC123/CAMK1D, THADA, ADAMTS9, BCL11A, and MTNR1B affect different aspects of pancreatic beta-cell function. Diabetes. 2010;59(1):293–301. doi:10.2337/db09-1048

761. Menendez JA, Vazquez-Martin A, Ortega FJ, Fernandez-Real JM. Fatty acid synthase: association with insulin resistance, type 2 diabetes, and cancer. Clin Chem. 2009;55(3):425–438. doi:10.1373/clinchem.2008.115352

762. Snabboon T, Plengpanich W, Sridama V, Sunthornyothin S, Suwanwalaikorn S, Khovidhunkit W. A SPINK1 gene mutation in a Thai patient with fibrocalculous pancreatic diabetes. Southeast Asian J Trop Med Public Health. 2006;37(3):559–562.

763. Arner P, Petrus P, Esteve D, Boulomié A, Näslund E, Thorell A, Gao H, Dahlman I, Rydén M. Screening of potential adipokines identifies S100A4 as a marker of pernicious adipose tissue and insulin resistance. Int J Obes (Lond). 2018;42(12):2047–2056. doi:10.1038/s41366-018-0018-0

764. Hassan M, Raslan HM, Eldin HG, Mahmoud E, Elwajed HAA. CD33+ HLA-DR-Myeloid-Derived Suppressor Cells Are Increased in Frequency in the Peripheral Blood of Type1 Diabetes Patients with Predominance of CD14+ Subset. Open Access Maced J Med Sci. 2018;6(2):303–309. doi:10.3889/oamjms.2018.080

765. Brings S, Fleming T, Herzig S, Nawroth PP, Kopf S. Urinary cathepsin L is predictive of changes in albuminuria and correlates with glucosepane in patients with type 2 diabetes in a closed-cohort study. J Diabetes Complications. 2020;34(9):107648. doi:10.1016/j.jdiacomp.2020.107648

766. Khattab A, Torkamani A. Nidogen-1 could play a role in diabetic kidney disease development in type 2 diabetes: a genome-wide association meta-analysis. Hum Genomics. 2022;16(1):47. doi:10.1186/s40246-022-00422-y

767. Dlamini Z, Ntlabati P, Mbita Z, Shoba-Zikhali L. Pyruvate dehydrogenase kinase 4 (PDK4) could be involved in a regulatory role in apoptosis and a link between apoptosis and insulin resistance. Exp Mol Pathol. 2015;98(3):574–584. doi:10.1016/j.yexmp.2015.03.022

768. Ali Beg MM, Verma AK, Saleem M, Saud Alreshidi F, Alenazi F, Ahmad H, Joshi PC. Role and Significance of Circulating Biomarkers: miRNA and E2F1 mRNA Expression and Their Association with Type-2 Diabetic Complications. Int J Endocrinol. 2020;2020:6279168. doi:10.1155/2020/6279168

769. Tian Y, Wang Z, Zheng X, Song W, Cai L, Rane M, Zhao Y. KLF15 negatively regulates cardiac fibrosis by which SDF-1β attenuates cardiac fibrosis in type 2 diabetic mice. Toxicol Appl Pharmacol. 2021;427:115654. doi:10.1016/j.taap.2021.115654

770. Mutlu M, Yuksel N, Takmaz T, Dincel AS, Bilgihan A, Altınkaynak H. Aqueous humor pentraxin-3 levels in patients with diabetes mellitus. Eye (Lond). 2017;31(10):1463–1467. doi:10.1038/eye.2017.87

771. Nandy D, Mukhopadhyay D, Basu A. Both vascular endothelial growth factor and soluble Flt-1 are increased in type 2 diabetes but not in impaired fasting glucose. J Investig Med. 2010;58(6):804–806. doi:10.231/JIM.0b013e3181e96203

772. Liu W, Aerbajinai W, Li H, Liu Y, Gavrilova O, Jain S, Rodgers GP. Olfactomedin 4 Deletion Improves Male Mouse Glucose Intolerance and Insulin Resistance Induced by a High-Fat Diet. Endocrinology. 2018;159(9):3235–3244. doi:10.1210/en.2018-00451

773. Li SW, Wang J, Yang Y, Liu ZJ, Cheng L, Liu HY, Ma P, Luo W, Liu SM. Polymorphisms in FADS1 and FADS2 alter plasma fatty acids and desaturase levels in type 2 diabetic patients with coronary artery disease. J Transl Med. 2016;14:79. doi:10.1186/s12967-016-0834-8

774. Degirmenci I, Ozbayer C, Kebapci MN, Kurt H, Colak E, Gunes HV. Common variants of genes encoding TLR4 and TLR4 pathway members TIRAP and IRAK1 are effective on MCP1, IL6, IL1β, and TNFα levels in type 2 diabetes and insulin resistance. Inflamm Res. 2019;68(9):801–814. doi:10.1007/s00011-019-01263-7

775. Gu H, Jiang W, You N, Huang X, Li Y, Peng X, Dong R, Wang Z, Zhu Y, Wu K, et al. Soluble Klotho Improves Hepatic Glucose and Lipid Homeostasis in Type 2 Diabetes. Mol Ther Methods Clin Dev. 2020;18:811–823. doi:10.1016/j.omtm.2020.08.002

776. Liu D, Liu L, Hu Z, Song Z, Wang Y, Chen Z. Evaluation of the oxidative stress-related genes ALOX5, ALOX5AP, GPX1, GPX3 and MPO for contribution to the risk of type 2 diabetes mellitus in the Han Chinese population. Diab Vasc Dis Res. 2018;15(4):336–339. doi:10.1177/1479164118755044

777. Bobbala D, Mayhue M, Menendez A, Ilangumaran S, Ramanathan S. Trans-presentation of interleukin-15 by interleukin-15 receptor alpha is dispensable for the pathogenesis of autoimmune type 1 diabetes. Cell Mol Immunol. 2017;14(7):590–596. doi:10.1038/cmi.2015.102

778. Güneş S, Wu J, Özyılmaz B, Deveci Sevim R, Ünüvar T, Anık A. Cooccurring Type 1 Diabetes Mellitus and Autoimmune Thyroiditis in a Girl with Craniofrontonasal Syndrome: Are EFNB1 Variants Associated with Autoimmunity?. Pharmaceuticals (Basel). 2022;15(12):1535. doi:10.3390/ph15121535

779. Beale EG, Harvey BJ, Forest C. PCK1 and PCK2 as candidate diabetes and obesity genes. Cell Biochem Biophys. 2007;48(2-3):89–95. doi:10.1007/s12013-007-0025-6

780. Hao JS, Zhu CJ, Yan BY, Yan CY, Ling R. Stimulation of KLF14/PLK1 pathway by thrombin signaling potentiates endothelial dysfunction in Type 2 diabetes mellitus. Biomed Pharmacother. 2018;99:859–866. doi:10.1016/j.biopha.2018.01.151

781. Sasso GRDS, Cerri PS, Sasso-Cerri E, Simões MJ, Gil CD, Florencio-Silva R. Possible role of annexin A1/FPR2 pathway in COX2/NLRP3 inflammasome regulation in alveolar bone cells of estrogen-deficient female rats with diabetes mellitus. J Periodontol. 2023. doi:10.1002/JPER.23-0530

782. Łukawska-Tatarczuk M, Franek E, Czupryniak L, Joniec-Maciejak I, Pawlak A, Wojnar E, Zieliński J, Mirowska-Guzel D, Mrozikiewicz-Rakowska B. Sirtuin 1, Visfatin and IL-27 Serum Levels of Type 1 Diabetic Females in Relation to Cardiovascular Parameters and Autoimmune Thyroid Disease. Biomolecules. 2021;11(8):1110. doi:10.3390/biom11081110

783. Ren H, Tan SL, Liu MZ, Banh HL, Luo JQ. Association of PON2 Gene Polymorphisms (Ser311Cys and Ala148Gly) With the Risk of Developing Type 2 Diabetes Mellitus in the Chinese Population. Front Endocrinol (Lausanne). 2018;9:495. doi:10.3389/fendo.2018.00495

784. Liu J, Wang Y, Gong L, Sun C. Oxidation of glyceraldehyde-3-phosphate dehydrogenase decreases sperm motility in diabetes mellitus. Biochem Biophys Res Commun. 2015;465(2):245–248. doi:10.1016/j.bbrc.2015.08.006

785. Wang J, Xia Y, Li J, Wang W. miR-129-5p in exosomes inhibits diabetes-associated osteogenesis in the jaw via targeting FZD4. Biochem Biophys Res Commun. 2021;566:87–93. doi:10.1016/j.bbrc.2021.05.072

786. Müssig K, Staiger H, Machicao F, Machann J, Schick F, Schäfer SA, Claussen CD, Holst JJ, Gallwitz B, Stefan N, et al. Preliminary report: genetic variation within the GPBAR1 gene is not associated with metabolic traits in white subjects at an increased risk for type 2 diabetes mellitus. Metabolism. 2009;58(12):1809–1811. doi:10.1016/j.metabol.2009.06.012

787. Jelić-Knezović N, Galijašević S, Lovrić M, Vasilj M, Selak S, Mikulić I. Levels of Nitric Oxide Metabolites and Myeloperoxidase in Subjects with Type 2 Diabetes Mellitus on Metformin TherapyL. Exp Clin Endocrinol Diabetes. 2019;127(1):56–61. doi:10.1055/a-0577-7776

788. Moura Alves Seixas G, de Souza Freitas R, Ferreira Fratelli C, de Souza Silva CM, Ramos de Lima L, Morato Stival M, Schwerz Funghetto S, Rodrigues da Silva IC. MAOA uVNTR Polymorphism Influence on Older Adults Diagnosed with Diabetes Mellitus/Systemic Arterial Hypertension. J Aging Res. 2023;2023:8538027. doi:10.1155/2023/8538027

789. Flowers JB, Rabaglia ME, Schueler KL, Flowers MT, Lan H, Keller MP, Ntambi JM, Attie AD. Loss of stearoyl-CoA desaturase-1 improves insulin sensitivity in lean mice but worsens diabetes in leptin-deficient obese mice. Diabetes. 2007;56(5):1228–1239. doi:10.2337/db06-1142

790. Cho YS, Go MJ, Han HR, Cha SH, Kim HT, Min H, Shin HD, Park C, Han BG, Cho NH, et al. Association of lipoprotein lipase (LPL) single nucleotide polymorphisms with type 2 diabetes mellitus. Exp Mol Med. 2008;40(5):523–532. doi:10.3858/emm.2008.40.5.523

791. Haghvirdizadeh P, Mohamed Z, Abdullah NA, Haghvirdizadeh P, Haerian MS, Haerian BS. KCNJ11: Genetic Polymorphisms and Risk of Diabetes Mellitus. J Diabetes Res. 2015;2015:908152. doi:10.1155/2015/908152

792. Liu T, Chen WQ, David SP, Tyndale RF, Wang H, Chen YM, Yu XQ, Chen W, Zhou Q, Ling WH. Interaction between heavy smoking and CYP2A6 genotypes on type 2 diabetes and its possible pathways. Eur J Endocrinol. 2011;165(6):961–967. doi:10.1530/EJE-11-0596

793. Shukla N, Kumari S, Verma P, Kushwah AS, Banarjee M, Sankhwar SN, Srivastava A, Ansari MS, Gautam NK. Genotypic Analysis of COL4A1 Gene in Diabetic Nephropathy and Type 2 Diabetes Mellitus Patients: A Comparative Genetic Study. DNA Cell Biol. 2023;42(9):541–547. doi:10.1089/dna.2023.0125

794. Zhao YY, Huang XY, Chen ZW. Daintain/AIF-1 (Allograft Inflammatory Factor-1) accelerates type 1 diabetes in NOD mice. Biochem Biophys Res Commun. 2012;427(3):513–517. doi:10.1016/j.bbrc.2012.09.087

795. Yang Y, Qiu W, Meng Q, Liu M, Lin W, Yang H, Wang R, Dong J, Yuan N, Zhou Z, et al. GRB10 rs1800504 Polymorphism Is Associated With the Risk of Coronary Heart Disease in Patients With Type 2 Diabetes Mellitus. Front Cardiovasc Med. 2021;8:728976. doi:10.3389/fcvm.2021.728976

796. Mannerås-Holm L, Kirchner H, Björnholm M, Chibalin AV, Zierath JR. mRNA expression of diacylglycerol kinase isoforms in insulin-sensitive tissues: effects of obesity and insulin resistance. Physiol Rep. 2015;3(4):e12372. doi:10.14814/phy2.12372

797. Limonte CP, Valo E, Drel V, Natarajan L, Darshi M, Forsblom C, Henderson CM, Hoofnagle AN, Ju W, Kretzler M, et al. Urinary Proteomics Identifies Cathepsin D as a Biomarker of Rapid eGFR Decline in Type 1 Diabetes. Diabetes Care. 2022;45(6):1416–1427. doi:10.2337/dc21-2204

798. Ahmad R, Kochumon S, Thomas R, Atizado V, Sindhu S. Increased adipose tissue expression of TLR8 in obese individuals with or without type-2 diabetes: significance in metabolic inflammation. J Inflamm (Lond). 2016;13:38. doi:10.1186/s12950-016-0147-y

799. Zarei R, Nikpour P, Rashidi B, Eskandari N, Aboutorabi R. Evaluation of Muc1 Gene Expression at The Time of Implantation in Diabetic Rat Models Treated with Insulin, Metformin and Pioglitazone in The Normal Cycle and Ovulation Induction Cycle. Int J Fertil Steril. 2020;14(3):218–222. doi:10.22074/ijfs.2020.44409

800. Mohamed AA, Khater SI, Hamed Arisha A, Metwally MMM, Mostafa-Hedeab G, El-Shetry ES. Chitosan-stabilized selenium nanoparticles alleviate cardio-hepatic damage in type 2 diabetes mellitus model via regulation of caspase, Bax/Bcl-2, and Fas/FasL-pathway. Gene. 2021;768:145288. doi:10.1016/j.gene.2020.145288

801. Ha X, Cai X, Cao H, Li J, Yang B, Jiang R, Li X, Li B, Xin Y. Docking protein 1 and free fatty acids are associated with insulin resistance in patients with type 2 diabetes mellitus. J Int Med Res. 2021;49(11):3000605211048293. doi:10.1177/03000605211048293

802. Zhao Q, Du X, Liu F, Zhang Y, Qin W, Zhang Q. ECHDC3 Variant Regulates the Right Hippocampal Microstructural Integrity and Verbal Memory in Type 2 Diabetes Mellitus. Neuroscience. 2023. doi:10.1016/j.neuroscience.2023.12.003

803. Li S, Li N, Li L, Zhan J. Sex Difference in the Association Between Serum Versican and Albuminuria in Patients with Type 2 Diabetes Mellitus. Diabetes Metab Syndr Obes. 2023;16:3631–3639. doi:10.2147/DMSO.S434287

804. Basu J, Mukherjee R, Sahu P, Datta C, Chowdhury S, Mandal D, Ghosh A. Association of common variants of TCF7L2 and PCSK2 with gestational diabetes mellitus in West Bengal, India. Nucleosides Nucleotides Nucleic Acids. 2023. doi:10.1080/15257770.2023.2248201

805. Shetty SS, Kumari NS. Fatty acid desaturase 2 (FADS 2) rs174575 (C/G) polymorphism, circulating lipid levels and susceptibility to type-2 diabetes mellitus. Sci Rep. 2021;11(1):13151. doi:10.1038/s41598-021-92572-7

806. Coy JF, Dressler D, Wilde J, Schubert P. Mutations in the transketolase-like gene TKTL1: clinical implications for neurodegenerative diseases, diabetes and cancer. Clin Lab. 2005;51(5-6):257–273.

807. Azarova YE, Klyosova EY, Ivakin VE, Churilin MI, Kolomoets II, Sunyaykina OA, Ragulina VA, Polonikov AV. Polymorphisms of the NCF4 Gene Increase the Risk of Chronic Heart Failure in Patients with Type 2 Diabetes Mellitus. Bull Exp Biol Med. 2023;176(1):77–81. doi:10.1007/s10517-023-05974-0

808. Cardinale CJ, Chang X, Wei Z, Qu HQ, Bradfield JP, Polychronakos C, Hakonarson H. Genome-wide association study of the age of onset of type 1 diabetes reveals HTATIP2 as a novel T cell regulator. Front Immunol. 2023;14:1101488. doi:10.3389/fimmu.2023.1101488

809. Chappell JE, Stewart JK. Soluble and particulate phenylethanolamine N-methyltransferase in hypothalamus of diabetic rats. Am J Physiol. 1992;263(2 Pt 1):E335–E339. doi:10.1152/ajpendo.1992.263.2.E335

810. Kolodka T, Charles ML, Raghavan A, Radichev IA, Amatya C, Ellefson J, Savinov AY, Nag A, Williams MS, Robbins MS. Preclinical characterization of recombinant human tissue kallikrein-1 as a novel treatment for type 2 diabetes mellitus. PLoS One. 2014;9(8):e103981. doi:10.1371/journal.pone.0103981

811. Mychaleckyj JC, Valo E, Ichimura T, Ahluwalia TS, Dina C, Miller RG, Shabalin IG, Gyorgy B, Cao J, Onengut-Gumuscu S, et al. Association of Coding Variants in Hydroxysteroid 17-beta Dehydrogenase 14 (HSD17B14) with Reduced Progression to End Stage Kidney Disease in Type 1 Diabetes. J Am Soc Nephrol. 2021;32(10):2634–2651. doi:10.1681/ASN.2020101457

812. Yang S, Chen C, Wang H, Rao X, Wang F, Duan Q, Chen F, Long G, Gong W, Zou MH, et al. Protective effects of Acyl-coA thioesterase 1 on diabetic heart via PPARα/PGC1α signaling. PLoS One. 2012;7(11):e50376. doi:10.1371/journal.pone.0050376

813. Wang Z, Zhang W, Huo B, Dong L, Zhang J. Relationship between thymidine kinase 1 before radiotherapy and prognosis in breast cancer patients with diabetes. Biosci Rep. 2020;40(4):BSR20192813. doi:10.1042/BSR20192813

814. Lourenço C, Kelly D, Cantillon J, Cauchi M, Yon MA, Bentley L, Cox RD, Turner C. Monitoring type 2 diabetes from volatile faecal metabolome in Cushing’s syndrome and single Afmid mouse models via a longitudinal study. Sci Rep. 2019;9(1):18779. doi:10.1038/s41598-019-55339-9

815. Cui P, Wu S, Xu X, Ye H, Hou J, Liu X, Wang H, Fang X. Deficiency of the Transcription Factor NR4A1 Enhances Bacterial Clearance and Prevents Lung Injury During Escherichia Coli Pneumonia. Shock. 2019;51(6):787–794. doi:10.1097/SHK.0000000000001184

816. de Brito RC, Lucena-Silva N, Torres LC, Luna CF, Correia JB, da Silva GA. The balance between the serum levels of IL-6 and IL-10 cytokines discriminates mild and severe acute pneumonia. BMC Pulm Med. 2016;16(1):170. doi:10.1186/s12890-016-0324-z

817. Chen Z, Chen X, Cheng HT, Yeh SC, Yu HY, Cheng JW, Li F. A novel CXCL8-IP10 hybrid protein is effective in blocking pulmonary pathology in a mouse model of Klebsiella pneumoniae infection. Int Immunopharmacol. 2018;62:40–45. doi:10.1016/j.intimp.2018.06.040

818. Nishikawa K, Morii T, Ako H, Hamada K, Saito S, Narita N. In vivo expression of CD69 on lung eosinophils in eosinophilic pneumonia: CD69 as a possible activation marker for eosinophils. J Allergy Clin Immunol. 1992;90(2):169–174. doi:10.1016/0091-6749(92)90068-d

819. Martins FRB, de Oliveira MD, Souza JAM, Queiroz-Junior CM, Lobo FP, Teixeira MM, Malacco NL, Soriani FM. Chronic ethanol exposure impairs alveolar leukocyte infiltration during pneumococcal pneumonia, leading to an increased bacterial burden despite increased CXCL1 and nitric oxide levels. Front Immunol. 2023;14:1175275. doi:10.3389/fimmu.2023.1175275

820. Chi X, Ding B, Zhang L, Zhang J, Wang J, Zhang W. lncRNA GAS5 promotes M1 macrophage polarization via miR-455-5p/SOCS3 pathway in childhood pneumonia. J Cell Physiol. 2019;234(8):13242–13251. doi:10.1002/jcp.27996

821. Zeng X, Moore TA, Newstead MW, Hernandez-Alcoceba R, Tsai WC, Standiford TJ. Intrapulmonary expression of macrophage inflammatory protein 1alpha (CCL3) induces neutrophil and NK cell accumulation and stimulates innate immunity in murine bacterial pneumonia. Infect Immun. 2003;71(3):1306–1315. doi:10.1128/IAI.71.3.1306-1315.2003

822. Zhou Z, Zhu Y, Gao G, Zhang Y. Long noncoding RNA SNHG16 targets miR-146a-5p/CCL5 to regulate LPS-induced WI-38 cell apoptosis and inflammation in acute pneumonia. Life Sci. 2019;228:189–197. doi:10.1016/j.lfs.2019.05.008

823. Parra ER, Silvério da Costa LR, Ab’Saber A, Ribeiro de Carvalho CR, Kairalla RA, Fernezlian SM, Teixeira LR, Capelozzi VL. Nonhomogeneous density of CD34 and VCAM-1 alveolar capillaries in major types of idiopathic interstitial pneumonia. Lung. 2005;183(5):363–373. doi:10.1007/s00408-005-2548-1

824. Cao M, Liu H, Dong Y, Liu W, Yu Z, Wang Q, Wang Q, Liang Z, Li Y, Ren H. Mesenchymal stem cells alleviate idiopathic pneumonia syndrome by modulating T cell function through CCR2-CCL2 axis. Stem Cell Res Ther. 2021;12(1):378. doi:10.1186/s13287-021-02459-7

825. Xue M, Guo Z, Cai C, Sun B, Wang H. Evaluation of the Diagnostic Efficacies of Serological Markers KL-6, SP-A, SP-D, CCL2, and CXCL13 in Idiopathic Interstitial Pneumonia. Respiration. 2019;98(6):534–545. doi:10.1159/000503689

826. Song H, Xi J, Li GG, Xu S, Wang C, Cheng T, Li H, Zhang Y, Liu X, Bai J. Upregulation of CD19LCD24(hi)CD38(hi) regulatory B cells is associated with a reduced risk of acute lung injury in elderly pneumonia patients. Intern Emerg Med. 2016;11(3):415–423. doi:10.1007/s11739-015-1377-3

827. Traber KE, Hilliard KL, Allen E, Wasserman GA, Yamamoto K, Jones MR, Mizgerd JP, Quinton LJ. Induction of STAT3-Dependent CXCL5 Expression and Neutrophil Recruitment by Oncostatin-M during Pneumonia. Am J Respir Cell Mol Biol. 2015;53(4):479–488. doi:10.1165/rcmb.2014-0342OC

828. De Leo F, Rossi A, De Marchis F, Cigana C, Melessike M, Quilici G, De Fino I, Mantonico MV, Fabris C, Bragonzi A, et al. Pamoic acid is an inhibitor of HMGB1·CXCL12 elicited chemotaxis and reduces inflammation in murine models of Pseudomonas aeruginosa pneumonia. Mol Med. 2022;28(1):108. doi:10.1186/s10020-022-00535-z

829. Traber KE, Dimbo EL, Shenoy AT, Symer EM, Allen E, Mizgerd JP, Quinton LJ. Neutrophil-Derived Oncostatin M Triggers Diverse Signaling Pathways during Pneumonia. Infect Immun. 2021;89(4):e00655–20. doi:10.1128/IAI.00655-20

830. Cho SJ, Pronko A, Yang J, Pagan K, Stout-Delgado H. Role of Cholesterol 25-Hydroxylase (Ch25h) in Mediating Innate Immune Responses to Streptococcus pneumoniae Infection. Cells. 2023;12(4):570. doi:10.3390/cells12040570

831. Franco-Leyva T, Torres OH, Saez Prieto ME, Boera-Carnicero G, Santos Á, Clotet S, Albert-Jares D, El-Ebiary Y, Agustí-Martí M, Casademont J, et al. Early differentiated CD28+ CD27+ T lymphocytes as a biomarker for short and long-term outcomes in older patients with pneumonia. J Leukoc Biol. 2022;112(5):1183–1190. doi:10.1002/JLB.5MA0422-370R

832. Herta T, Bhattacharyya A, Rosolowski M, Conrad C, Gurtner C, Gruber AD, Ahnert P, Gutbier B, Frey D, Suttorp N, et al. Krueppel-Like Factor 4 Expression in Phagocytes Regulates Early Inflammatory Response and Disease Severity in Pneumococcal Pneumonia. Front Immunol. 2021;12:726135. doi:10.3389/fimmu.2021.726135

833. Nguyen LS, Ait Hamou Z, Gastli N, Chapuis N, Pène F. Potential role for interferon gamma in the treatment of recurrent ventilator-acquired pneumonia in patients with COVID-19: a hypothesis. Intensive Care Med. 2021;47(5):619–621. doi:10.1007/s00134-021-06377-3

834. Baranek T, Morello E, Valayer A, Aimar RF, Bréa D, Henry C, Besnard AG, Dalloneau E, Guillon A, Dequin PF, et al. HL2 Regulates Natural Killer Cell Development and Activation during Streptococcus pneumoniae Infection. Front Immunol. 2017;8:123. doi:10.3389/fimmu.2017.00123

835. Kikuchi T, Andarini S, Xin H, Gomi K, Tokue Y, Saijo Y, Honjo T, Watanabe A, Nukiwa T. Involvement of fractalkine/CX3CL1 expression by dendritic cells in the enhancement of host immunity against Legionella pneumophila. Infect Immun. 2005;73(9):5350–5357. doi:10.1128/IAI.73.9.5350-5357.2005

836. van den Boogaard FE, van Gisbergen KP, Vernooy JH, Medema JP, Roelofs JJ, van Zoelen MA, Endeman H, Biesma DH, Boon L, Van’t Veer C, et al. Granzyme A impairs host defense during Streptococcus pneumoniae pneumonia. Am J Physiol Lung Cell Mol Physiol. 2016;311(2):L507–L516. doi:10.1152/ajplung.00116.2016

837. Nureki S, Miyazaki E, Ishi T, et al. Elevated concentrations of CCR7 ligands in patients with eosinophilic pneumonia. Allergy. 2013;68(11):1387–1395. doi:10.1111/all.12243

838. Chang PY, Tsao SM, Chang JH, Chien MH, Hung WY, Huang YW, Yang SF. Plasma levels of soluble intercellular adhesion molecule-1 as a biomarker for disease severity of patients with community-acquired pneumonia. Clin Chim Acta. 2016;463:174–180. doi:10.1016/j.cca.2016.10.030

839. Akaba T, Kondo M, Hara K, Mizobuchi R, Abe K, Miyoshi A, Yagi O, Tagaya E. Tryptase and IL-33 in Bronchoalveolar Lavage Fluid May Predict the Types of Eosinophilic Pneumonia and Disease Recurrence. Int Arch Allergy Immunol. 2022;183(4):415–423. doi:10.1159/000520180

840. Lopez AD, Avasarala S, Grewal S, Murali AK, London L. Differential role of the Fas/Fas ligand apoptotic pathway in inflammation and lung fibrosis associated with reovirus 1/L-induced bronchiolitis obliterans organizing pneumonia and acute respiratory distress syndrome. J Immunol. 2009;183(12):8244–8257. doi:10.4049/jimmunol.0901958

841. Mancuso P, O Brien E, Prano J, Goel D, Aronoff DM. No Impairment in host defense against Streptococcus pneumoniae in obese CPEfat/fat mice. PLoS One. 2014;9(9):e106420. doi:10.1371/journal.pone.0106420

842. Connor MG, Camarasa TMN, Patey E, Rasid O, Barrio L, Weight CM, Miller DP, Heyderman RS, Lamont RJ, Enninga J, et al. The histone demethylase KDM6B fine-tunes the host response to Streptococcus pneumoniae. Nat Microbiol. 2021;6(2):257–269. doi:10.1038/s41564-020-00805-8

843. Pandey P, Karupiah G. Targeting tumour necrosis factor to ameliorate viral pneumonia. FEBS J. 2022;289(4):883–900. doi:10.1111/febs.15782

844. Sekiguchi N, Komatsu M, Ichiyama T, Kobayashi A, Gomi D, Fukushima T, Kobayashi T, Noguchi T, Nakazawa H, Asano N, et al. CCR4-positive peripheral T-cell lymphoma presenting as eosinophilic pneumonia and developing from prolonged pustular psoriasis. J Int Med Res. 2021;49(2):300060521996165. doi:10.1177/0300060521996165

845. Wang TJ, Wu LF, Chen J, Zhu W, Wang H, Liu XL, Teng YQ. X-linked hyper-IgM syndrome complicated with interstitial pneumonia and liver injury: a new mutation locus in the CD40LG gene. Immunol Res. 2019;67(4-5):454–459. doi:10.1007/s12026-019-09098-4

846. Shi H, Wang W, Yin J, Ouyang Y, Pang L, Feng Y, Qiao L, Guo X, Shi H, Jin R, et al. The inhibition of IL-2/IL-2R gives rise to CD8+ T cell and lymphocyte decrease through JAK1-STAT5 in critical patients with COVID-19 pneumonia. Cell Death Dis. 2020;11(6):429. doi:10.1038/s41419-020-2636-4

847. Lin YC, Lu MC, Lin C, Chiang MK, Jan MS, Tang HL, Liu HC, Lin WL, Huang CY, et al. Activation of IFN-γ/STAT/IRF-1 in hepatic responses to Klebsiella pneumoniae infection. PLoS One. 2013;8(11):e79961. doi:10.1371/journal.pone.0079961

848. St Jean SC, Ricart Arbona RJ, Mishkin N, Monette S, Wipf JRK, Henderson KS, Cheleuitte-Nieves C, Lipman NS, Carrasco SE. Chlamydia muridarum infection causes bronchointerstitial pneumonia in NOD.Cg-PrkdcscidIl2rgtm1Wjl/SzJ (NSG) mice. Vet Pathol. 2024;61(1):145–156. doi:10.1177/03009858231183907

849. Liang X, Gupta K, Quintero JR, Cernadas M, Kobzik L, Christou H, Pier GB, Owen CA, Çataltepe S. Macrophage FABP4 is required for neutrophil recruitment and bacterial clearance in Pseudomonas aeruginosa pneumonia. FASEB J. 2019;33(3):3562–3574. doi:10.1096/fj.201802002R

850. Shimizu Y, Dobashi K, Endou K, Ono A, Yanagitani N, Utsugi M, Sano T, Ishizuka T, Shimizu K, Tanaka S, et al. Decreased interstitial FOXP3(+) lymphocytes in usual interstitial pneumonia with discrepancy of CXCL12/CXCR4 axis. Int J Immunopathol Pharmacol. 2010;23(2):449–461. doi:10.1177/039463201002300207

851. de Stoppelaar SF, Van’t Veer C, Roelofs JJ, Claushuis TA, de Boer OJ, Tanck MW, Hoogendijk AJ, van der Poll T. Platelet and endothelial cell P-selectin are required for host defense against Klebsiella pneumoniae-induced pneumosepsis. J Thromb Haemost. 2015;13(6):1128–1138. doi:10.1111/jth.12893

852. Fernández JJ, Mancebo C, Garcinuño S, March G, Alvarez Y, Alonso S, Inglada L, Blanco J, Orduña A, Montero O, et al. Innate IRE1α-XBP1 activation by viral single-stranded RNA and its influence on lung cytokine production during SARS-CoV-2 pneumonia. Genes Immun. 2023. doi:10.1038/s41435-023-00243-6

853. Gu M, Han X, Liu X, Sui F, Zhang Q, Pan S. Predictive Value of Annenxin A1 for Disease Severity and Prognosis in Patients with Community-Acquired Pneumonia. Diagnostics (Basel). 2023;13(3):396. doi:10.3390/diagnostics13030396

854. Xu L, Song Q, Ouyang Z, Zhang X, Zhang C. let7fL5p attenuates inflammatory injury in in vitro pneumonia models by targeting MAPK6. Mol Med Rep. 2021;23(2):95. doi:10.3892/mmr.2020.11734

855. Watanabe E, Akamatsu T, Ohmori M, Kato M, Takeuchi N, Ishiwada N, Nishimura R, Hishiki H, Fujimura L, Ito C, et al. Recombinant thrombomodulin attenuates hyper-inflammation and glycocalyx damage in a murine model of Streptococcus pneumoniae-induced sepsis. Cytokine. 2022;149:155723. doi:10.1016/j.cyto.2021.155723

856. Sakthivel P, Gereke M, Breithaupt A, Fuchs D, Gigliotti L, Gruber AD, Dianzani U, Bruder D. Attenuation of immune-mediated influenza pneumonia by targeting the inducible co-stimulator (ICOS) molecule on T cells. PLoS One. 2014;9(7):e100970. doi:10.1371/journal.pone.0100970

857. De Cristofaro R, Liuzzo G, Sacco M, Lancellotti S, Pedicino D, Andreotti F. Marked von Willebrand factor and factor VIII elevations in severe acute respiratory syndrome coronavirus-2-positive, but not severe acute respiratory syndrome coronavirus-2-negative, pneumonia: a case-control study. Blood Coagul Fibrinolysis. 2021;32(4):285–289. doi:10.1097/MBC.0000000000000998

858. Song X, Liu B, Zhao G, Pu X, Liu B, Ding M, Xue Y. Streptococcus pneumoniae promotes migration and invasion of A549 cells in vitro by activating mTORC2/AKT through up-regulation of DDIT4 expression. Front Microbiol. 2022;13:1046226. doi:10.3389/fmicb.2022.1046226

859. Haydar D, Gonzalez R, Garvy BA, Garneau-Tsodikova S, Thamban Chandrika N, Bocklage TJ, Feola DJ. Myeloid arginase-1 controls excessive inflammation and modulates T cell responses in Pseudomonas aeruginosa pneumonia. Immunobiology. 2021;226(1):152034. doi:10.1016/j.imbio.2020.152034

860. Domon H, Terao Y. The Role of Neutrophils and Neutrophil Elastase in Pneumococcal Pneumonia. Front Cell Infect Microbiol. 2021;11:615959. doi:10.3389/fcimb.2021.615959

861. Yu R, Chen CR, Evans D, Qing X, Gotesman M, Chandramohan G, Kallay T, Lin HJ, Pedigo TP. Glucose-6-phosphate dehydrogenase deficiency presenting with rhabdomyolysis in a patient with coronavirus disease 2019 pneumonia: a case report. J Med Case Rep. 2022;16(1):106. doi:10.1186/s13256-022-03322-w

862. Wu M, Gibbons JG, DeLoid GM, Bedugnis AS, Thimmulappa RK, Biswal S, Kobzik L. Immunomodulators targeting MARCO expression improve resistance to postinfluenza bacterial pneumonia. Am J Physiol Lung Cell Mol Physiol. 2017;313(1):L138–L153. doi:10.1152/ajplung.00075.2017

863. Paudel S, Ghimire L, Jin L, Baral P, Cai S, Jeyaseelan S. NLRC4 suppresses IL-17A-mediated neutrophil-dependent host defense through upregulation of IL-18 and induction of necroptosis during Gram-positive pneumonia. Mucosal Immunol. 2019;12(1):247–257. doi:10.1038/s41385-018-0088-2

864. Steiner P, Otth M, Casaulta C, Aebi C. Autoantibodies against bactericidal/permeability-increasing protein (BPI) in children with acute pneumonia. FEMS Immunol Med Microbiol. 2009;57(2):125–128. doi:10.1111/j.1574-695X.2009.00593.x

865. Cheng SL, Wang HC, Cheng SJ, Yu CJ. Elevated placenta growth factor predicts pneumonia in patients with chronic obstructive pulmonary disease under inhaled corticosteroids therapy. BMC Pulm Med. 2011;11:46. doi:10.1186/1471-2466-11-46

866. Kagimoto A, Tsutani Y, Kushitani K, Kambara T, Mimae T, Miyata Y, Takeshima Y, Okada M. Usefulness of serum S100A4 and positron-emission tomography on lung cancer accompanied by interstitial pneumonia. Thorac Cancer. 2023;14(4):381–388. doi:10.1111/1759-7714.14757

867. Shi GQ, Yang L, Shan LY, Yin LZ, Jiang W, Tian HT, Yang DD. Investigation of the clinical significance of detecting PTX3 for community-acquired pneumonia. Eur Rev Med Pharmacol Sci. 2020;24(16):8477–8482. doi:10.26355/eurrev_202008_22645

868. González-Lara MF, Wisniowski-Yáñez A, Pérez-Patrigeon S, Hsu AP, Holland SM, Cuellar-Rodríguez JM. Pneumocystis jiroveci pneumonia and GATA2 deficiency: Expanding the spectrum of the disease. J Infect. 2017;74(4):425–427. doi:10.1016/j.jinf.2017.01.005

869. Young LR, Nogee LM, Barnett B, Panos RJ, Colby TV, Deutsch GH. Usual interstitial pneumonia in an adolescent with ABCA3 mutations. Chest. 2008;134(1):192–195. doi:10.1378/chest.07-2652

870. Machado MG, Tavares LP, Souza GVS, Queiroz-Junior CM, Ascenção FR, Lopes ME, Garcia CC, Menezes GB, Perretti M, Russo RC, et al. The Annexin A1/FPR2 pathway controls the inflammatory response and bacterial dissemination in experimental pneumococcal pneumonia. FASEB J. 2020;34(2):2749–2764. doi:10.1096/fj.201902172R

871. Xu Z, Wang XM, Cao P, Zhang C, Feng CM, Zheng L, Xu DX, Fu L, Zhao H. Serum IL-27 predicts the severity and prognosis in patients with community-acquired pneumonia: a prospective cohort study. Int J Med Sci. 2022;19(1):74–81. doi:10.7150/ijms.67028

872. Salcines-Cuevas D, Terán-Navarro H, Calderón-Gonzalez R, Torres-Rodriguez P, Tobes R, Fresno M, Calvo-Montes J, Molino-Bernal ICPD, Yañez-Diaz S, Alvarez-Dominguez C. Glyceraldehyde-3-phosphate Dehydrogenase Common Peptides of Listeria monocytogenes, Mycobacterium marinum and Streptococcus pneumoniae as Universal Vaccines. Vaccines (Basel). 2021;9(3):269. doi:10.3390/vaccines9030269

873. Bando M, Homma S, Harigai M. MPO-ANCA positive interstitial pneumonia: Current knowledge and future perspectives. Sarcoidosis Vasc Diffuse Lung Dis. 2022;38(4):e2021045. doi:10.36141/svdld.v38i4.11808

874. Coon DR, Roberts DJ, Loscertales M, Kradin R. Differential epithelial expression of SHH and FOXF1 in usual and nonspecific interstitial pneumonia. Exp Mol Pathol. 2006;80(2):119–123. doi:10.1016/j.yexmp.2005.12.003

875. Sand JMB, Tanino Y, Karsdal MA, Nikaido T, Misa K, Sato Y, Togawa R, Wang X, Leeming DJ, Munakata M. A Serological Biomarker of Versican Degradation is Associated with Mortality Following Acute Exacerbations of Idiopathic Interstitial Pneumonia. Respir Res. 2018;19(1):82. doi:10.1186/s12931-018-0779-y

876. Higgins MA, Boraston AB. Structure of the fucose mutarotase from Streptococcus pneumoniae in complex with L-fucose. Acta Crystallogr Sect F Struct Biol Cryst Commun. 2011;67(Pt 12):1524–1530. doi:10.1107/S1744309111046343

877. Egarnes B, Blanchet MR, Gosselin J. Treatment with the NR4A1 agonist cytosporone B controls influenza virus infection and improves pulmonary function in infected mice. PLoS One. 2017;12(10):e0186639. doi:10.1371/journal.pone.0186639

878. Yang ML, Wang CT, Yang SJ, Leu CH, Chen SH, Wu CL, Shiau AL. IL-6 ameliorates acute lung injury in influenza virus infection. Sci Rep. 2017;7:43829. doi:10.1038/srep43829

879. McGrath JJC, Vanderstocken G, Dvorkin-Gheva A, Cass SP, Afkhami S, Fantauzzi MF, Thayaparan D, Reihani A, Wang P, Beaulieu A, et al. Cigarette smoke augments CSF3 expression in neutrophils to compromise alveolar-capillary barrier function during influenza infection. Eur Respir J. 2022;60(2):2102049. doi:10.1183/13993003.02049-2021

880. Huipao N, Borwornpinyo S, Wiboon-Ut S, Campbell CR, Lee IH, Hiranyachattada S, Sukasem C, Thitithanyanont A, Pholpramool C, Cook DI, et al. P2Y6 receptors are involved in mediating the effect of inactivated avian influenza virus H5N1 on IL-6 & CXCL8 mRNA expression in respiratory epithelium. PLoS One. 2017;12(5):e0176974. doi:10.1371/journal.pone.0176974

881. Du N, Kwon H, Li P, West EE, Oh J, Liao W, Yu Z, Ren M, Leonard WJ.EGR2 is critical for peripheral naïve T-cell differentiation and the T-cell response to influenza. Proc Natl Acad Sci U S A. 2014;111(46):16484–16489. doi:10.1073/pnas.1417215111

882. Li S, Liu S, Chen RA, Huang M, Fung TS, Liu DX. Activation of the MKK3-p38-MK2-ZFP36 Axis by Coronavirus Infection Restricts the Upregulation of AU-Rich Element-Containing Transcripts in Proinflammatory Responses. J Virol. 2022;96(5):e0208621. doi:10.1128/jvi.02086-21

883. Antoniak S, Tatsumi K, Schmedes CM, Egnatz GJ, Auriemma AC, Bharathi V, Stokol T, Beck MA, Griffin JH, Palumbo JS, et al. PAR1 regulation of CXCL1 expression and neutrophil recruitment to the lung in mice infected with influenza A virus. J Thromb Haemost. 2021;19(4):1103–1111. doi:10.1111/jth.15221

884. Pothlichet J, Chignard M, Si-Tahar M. Cutting edge: innate immune response triggered by influenza A virus is negatively regulated by SOCS1 and SOCS3 through a RIG-I/IFNAR1-dependent pathway. J Immunol. 2008;180(4):2034–2038. doi:10.4049/jimmunol.180.4.2034

885. Tregoning JS, Pribul PK, Pennycook AM, Hussell T, Wang B, Lukacs N, Schwarze J, Culley FJ, Openshaw PJ. The chemokine MIP1alpha/CCL3 determines pathology in primary RSV infection by regulating the balance of T cell populations in the murine lung. PLoS One. 2010;5(2):e9381. doi:10.1371/journal.pone.0009381

886. Silva T, Temerozo JR, do Vale G, Ferreira AC, Soares VC, Dias SSG, Sardella G, Bou-Habib DC, Siqueira M, Souza TML, et al. The Chemokine CCL5 Inhibits the Replication of Influenza A Virus Through SAMHD1 Modulation. Front Cell Infect Microbiol. 2021;11:549020. doi:10.3389/fcimb.2021.549020

887. Alturaiki W, McFarlane AJ, Rose K, Corkhill R, McNamara PS, Schwarze J, Flanagan BF. Expression of the B cell differentiation factor BAFF and chemokine CXCL13 in a murine model of Respiratory Syncytial Virus infection. Cytokine. 2018;110:267–271. doi:10.1016/j.cyto.2018.01.014

888. Liu W, Zhao Y, Fan J, Shen J, Tang H, Tang W, Wu D, Huang W, Ding Y, Qiao P, et al. Smoke and Spike: Benzo[a]pyrene Enhances SARS-CoV-2 Infection by Boosting NR4A2-Induced ACE2 and TMPRSS2 Expression. Adv Sci (Weinh). 2023;10(26):e2300834. doi:10.1002/advs.202300834

889. Foo CX, Bartlett S, Chew KY, Ngo MD, Bielefeldt-Ohmann H, Arachchige BJ, Matthews B, Reed S, Wang R, Smith C, et al. GPR183 antagonism reduces macrophage infiltration in influenza and SARS-CoV-2 infection. Eur Respir J. 2023;61(3):2201306. doi:10.1183/13993003.01306-2022

890. Nguyen Thanh D, Thanh Giang NT, Le TV, Truong NM, Ngo TV, Lam TN, Nguyen DT, Tran QH, Nguyen MN. Predicting the severity of COVID-19 patients using the CD24-CSF1R index in whole blood samples. Heliyon. 2023;9(3):e13945. doi:10.1016/j.heliyon.2023.e13945

891. Guo L, Li N, Yang Z, Li H, Zheng H, Yang J, Chen Y, Zhao X, Mei J, Shi H, et al. Role of CXCL5 in Regulating Chemotaxis of Innate and Adaptive Leukocytes in Infected Lungs Upon Pulmonary Influenza Infection. Front Immunol. 2021;12:785457. doi:10.3389/fimmu.2021.785457

892. Goel S, Saheb Sharif-Askari F, Saheb Sharif Askari N, Madkhana B, Alwaa AM, Mahboub B, Zakeri AM, Ratemi E, Hamoudi R, Hamid Q, et al. SARS-CoV-2 Switches ‘on’ MAPK and NFκB Signaling via the Reduction of Nuclear DUSP1 and DUSP5 Expression. Front Pharmacol. 2021;12:631879. doi:10.3389/fphar.2021.631879

893. Fleming-Canepa X, Brusnyk C, Aldridge JR, Ross KL, Moon D, Wang D, Xia J, Barber MR, Webster RG, Magor KE. Expression of duck CCL19 and CCL21 and CCR7 receptor in lymphoid and influenza-infected tissues. Mol Immunol. 2011;48(15-16):1950–1957. doi:10.1016/j.molimm.2011.05.025

894. Touzelet O, Broadbent L, Armstrong SD, Aljabr W, Cloutman-Green E, Power UF, Hiscox JA. The Secretome Profiling of a Pediatric Airway Epithelium Infected with hRSV Identified Aberrant Apical/Basolateral Trafficking and Novel Immune Modulating (CXCL6, CXCL16, CSF3) and Antiviral (CEACAM1) Proteins. Mol Cell Proteomics. 2020;19(5):793–807. doi:10.1074/mcp.RA119.001546

895. García-Ramírez RA, Ramírez-Venegas A, Quintana-Carrillo R, Camarena ÁE, Falfán-Valencia R, Mejía-Aranguré JM. TNF, IL6, and IL1B Polymorphisms Are Associated with Severe Influenza A (H1N1) Virus Infection in the Mexican Population. PLoS One. 2015;10(12):e0144832. doi:10.1371/journal.pone.0144832

896. Fricke-Galindo I, Buendia-Roldan I, Chavez-Galan L, Pérez-Rubio G, Hernández-Zenteno RJ, Ramos-Martinez E, Zazueta-Márquez A, Reyes-Melendres F, Alarcón-Dionet A, Guzmán-Vargas J, et al. SERPINE1 rs6092 Variant Is Related to Plasma Coagulation Proteins in Patients with Severe COVID-19 from a Tertiary Care Hospital. Biology (Basel). 2022;11(4):595. doi:10.3390/biology11040595

897. Tian T, Zi X, Peng Y, Wang Z, Hong H, Yan Y, Guan W, Tan KS, Liu J, Ong HH, et al. H3N2 influenza virus infection enhances oncostatin M expression in human nasal epithelium. Exp Cell Res. 2018;371(2):322–329. doi:10.1016/j.yexcr.2018.08.022

898. Suryadevara M, Bonville CA, Rosenberg HF, Domachowske JB. Local production of CCL3, CCL11, and IFN-γ correlates with disease severity in murine parainfluenza virus infection. Virol J. 2013;10:357. doi:10.1186/1743-422X-10-357

899. Akauliya M, Gautam A, Maharjan S, Park BK, Kim J, Kwon HJ. CD83 expression regulates antibody production in response to influenza A virus infection. Virol J. 2020;17(1):194. doi:10.1186/s12985-020-01465-0

900. Park J, Park MY, Kim Y, Jun Y, Lee U, Oh CM. Apelin as a new therapeutic target for COVID-19 treatment. QJM. 2023;116(3):197–204. doi:10.1093/qjmed/hcac229

901. Shirato K, Ujike M, Kawase M, Matsuyama S. Identification of CCL2, RARRES2 and EFNB2 as host cell factors that influence the multistep replication of respiratory syncytial virus. Virus Res. 2015;210:213–226. doi:10.1016/j.virusres.2015.08.006

902. Song Z, Zhang Q, Liu X, Bai J, Zhao Y, Wang X, Jiang P. Cholesterol 25-hydroxylase is an interferon-inducible factor that protects against porcine reproductive and respiratory syndrome virus infection. Vet Microbiol. 2017;210:153–161. doi:10.1016/j.vetmic.2017.09.011

903. Rokni M, Sarhadi M, Heidari Nia M, Mohamed Khosroshahi L, Asghari S, Sargazi S, Mirinejad S, Saravani R. Single nucleotide polymorphisms located in TNFA, IL1RN, IL6R, and IL6 genes are associated with COVID-19 risk and severity in an Iranian population. Cell Biol Int. 2022;46(7):1109–1127. doi:10.1002/cbin.11807

904. Masemann D, Leite Dantas R, Sitnik S, Schied T, Nordhoff C, Ludwig S, Wixler V. The Four-and-a-Half LIM Domain Protein 2 Supports Influenza A Virus-Induced Lung Inflammation by Restricting the Host Adaptive Immune Response. Am J Pathol. 2018;188(5):1236–1245. doi:10.1016/j.ajpath.2018.02.004

905. Loxham M, Smart DE, Bedke NJ, Smithers NP, Filippi I, Blume C, Swindle EJ, Tariq K, Howarth PH, Holgate ST, et al. Allergenic proteases cleave the chemokine CX3CL1 directly from the surface of airway epithelium and augment the effect of rhinovirus. Mucosal Immunol. 2018;11(2):404–414. doi:10.1038/mi.2017.63

906. Shin JM, Han MS, Park JH, Lee SH, Kim TH, Lee SH. The EphA1 and EphA2 Signaling Modulates the Epithelial Permeability in Human Sinonasal Epithelial Cells and the Rhinovirus Infection Induces Epithelial Barrier Dysfunction via EphA2 Receptor Signaling. Int J Mol Sci. 2023;24(4):3629. doi:10.3390/ijms24043629

907. Pearson H, Todd EJAA, Ahrends M, Hover SE, Whitehouse A, Stacey M, Lippiat JD, Wilkens L, Fieguth HG, Danov O, et al. TMEM16A/ANO1 calcium-activated chloride channel as a novel target for the treatment of human respiratory syncytial virus infection. Thorax. 2021;76(1):64–72. doi:10.1136/thoraxjnl-2020-215171

908. Uppal S, Postnikova O, Villasmil R, Rogozin IB, Bocharov AV, Eggerman TL, Poliakov E, Redmond TM. Low-Density Lipoprotein Receptor (LDLR) Is Involved in Internalization of Lentiviral Particles Pseudotyped with SARS-CoV-2 Spike Protein in Ocular Cells. Int J Mol Sci. 2023;24(14):11860. doi:10.3390/ijms241411860

909. Inchley CS, Osterholt HC, Sonerud T, Fjærli HO, Nakstad B. Downregulation of IL7R, CCR7, and TLR4 in the cord blood of children with respiratory syncytial virus disease. J Infect Dis. 2013;208(9):1431–1435. doi:10.1093/infdis/jit336

910. Semiz S. SIT1 transporter as a potential novel target in treatment of COVID-19. Biomol Concepts. 2021;12(1):156–163. doi:10.1515/bmc-2021-0017

911. Maelfait J, Roose K, Bogaert P, Sze M, Saelens X, Pasparakis M, Carpentier I, van Loo G, Beyaert R. A20 (Tnfaip3) deficiency in myeloid cells protects against influenza A virus infection. PLoS Pathog. 2012;8(3):e1002570. doi:10.1371/journal.ppat.1002570

912. Ma P, Gu K, Wen R, Li C, Zhou C, Zhao Y, Li H, Lei C, Yang X, Wang H. Guanylate-binding protein 1 restricts avian coronavirus infectious bronchitis virus-infected HD11 cells. Poult Sci. 2023;102(3):102398. doi:10.1016/j.psj.2022.102398

913. Zhou B, Wang L, Yang S, Liang Y, Zhang Y, Pan X, Li J. Rosmarinic acid treatment protects against lethal H1N1 virus-mediated inflammation and lung injury by promoting activation of the h-PGDS-PGD2-HO-1 signal axis. Chin Med. 2023;18(1):139. doi:10.1186/s13020-023-00847-0

914. Othumpangat S, Noti JD, McMillen CM, Beezhold DH. ICAM-1 regulates the survival of influenza virus in lung epithelial cells during the early stages of infection. Virology. 2016;487:85–94. doi:10.1016/j.virol.2015.10.005

915. Yumine N, Matsumoto Y, Ohta K, Fukasawa M, Nishio M. Claudin-1 inhibits human parainfluenza virus type 2 dissemination. Virology. 2019;531:93–99. doi:10.1016/j.virol.2019.01.031

916. Murdaca G, Paladin F, Tonacci A, Borro M, Greco M, Gerosa A, Isola S, Allegra A, Gangemi S. Involvement of Il-33 in the Pathogenesis and Prognosis of Major Respiratory Viral Infections: Future Perspectives for Personalized Therapy. Biomedicines. 2022;10(3):715. doi:10.3390/biomedicines10030715

917. Hamldar S, Kiani SJ, Khoshmirsafa M, Nahand JS, Mirzaei H, Khatami A, Kahyesh-Esfandiary R, Khanaliha K, Tavakoli A, Babakhaniyan K, et al. Expression profiling of inflammation-related genes including IFI-16, NOTCH2, CXCL8, THBS1 in COVID-19 patients. Biologicals. 2022;80:27–34. doi:10.1016/j.biologicals.2022.09.001

918. Olson MR, Varga SM. Fas ligand is required for the development of respiratory syncytial virus vaccine-enhanced disease. J Immunol. 2009;182(5):3024–3031. doi:10.4049/jimmunol.0803585

919. Giuzio F, Bonomo MG, Catalano A, Infantino V, Salzano G, Monné M, Geronikaki A, Petrou A, Aquaro S, Sinicropi MS, et al. Potential PDE4B inhibitors as promising candidates against SARS-CoV-2 infection. Biomol Concepts. 2023;14(1):10.1515/bmc-2022-0033. doi:10.1515/bmc-2022-0033

920. Wu Q, Jiang D, Schaefer NR, Harmacek L, O’Connor BP, Eling TE, Eickelberg O, Chu HW. Overproduction of growth differentiation factor 15 promotes human rhinovirus infection and virus-induced inflammation in the lung. Am J Physiol Lung Cell Mol Physiol. 2018;314(3):L514–L527. doi:10.1152/ajplung.00324.2017

921. Ibañez LI, Martinez VP, Iglesias AA, Bellomo CM, Alonso DO, Coelho RM, Martinez Peralta L, Periolo N. Decreased expression of surfactant Protein-C and CD74 in alveolar epithelial cells during influenza virus A(H1N1)pdm09 and H3N2 infection. Microb Pathog. 2023;176:106017. doi:10.1016/j.micpath.2023.106017

922. Ashhurst AS, Flórido M, Lin LCW, Quan D, Armitage E, Stifter SA, Stambas J, Britton WJ. CXCR6-Deficiency Improves the Control of Pulmonary Mycobacterium tuberculosis and Influenza Infection Independent of T-Lymphocyte Recruitment to the Lungs. Front Immunol. 2019;10:339 doi:10.3389/fimmu.2019.00339

923. Kesavardhana S, Samir P, Zheng M, Malireddi RKS, Karki R, Sharma BR, Place DE, Briard B, Vogel P, Kanneganti TD. DDX3X coordinates host defense against influenza virus by activating the NLRP3 inflammasome and type I interferon response. J Biol Chem. 2021;296:100579. doi:10.1016/j.jbc.2021.100579

924. Kuriakose T, Zheng M, Neale G, Kanneganti TD. IRF1 Is a Transcriptional Regulator of ZBP1 Promoting NLRP3 Inflammasome Activation and Cell Death during Influenza Virus Infection. J Immunol. 2018;200(4):1489–1495. doi:10.4049/jimmunol.1701538

925. Ismail MMF, El-Awady RR, Farrag AM, Mahmoud SH, Abo Shama NM, Mostafa A, Ali MA, Rashed MH, Ibrahim IH. Potential role of PIM1 inhibition in the treatment of SARS-CoV-2 infection. J Genet Eng Biotechnol. 2023;21(1):65. doi:10.1186/s43141-023-00520-x

926. Basavaraju S, Mishra S, Jindal R, Kesavardhana S. Emerging Role of ZBP1 in Z-RNA Sensing, Influenza Virus-Induced Cell Death, and Pulmonary Inflammation. mBio. 2022;13(3):e0040122. doi:10.1128/mbio.00401-22

927. Ren C, Chen T, Zhang S, Gao Q, Zou J, Li P, Wang B, Zhao Y, OuYang A, Suolang S, et al. PLK3 facilitates replication of swine influenza virus by phosphorylating viral NP protein. Emerg Microbes Infect. 2023;12(2):2275606. doi:10.1080/22221751.2023.2275606

928. Ainsua-Enrich E, Hatipoglu I, Kadel S, Turner S, Paul J, Singh S, Bagavant H, Kovats S. IRF4-dependent dendritic cells regulate CD8+ T-cell differentiation and memory responses in influenza infection. Mucosal Immunol. 2019;12(4):1025–1037. doi:10.1038/s41385-019-0173-1

929. Pan P, Ge W, Lei Z, Luo W, Liu Y, Guan Z, Chen L, Yu Z, Shen M, Hu D, et al. SARS-CoV-2 N protein enhances the anti-apoptotic activity of MCL-1 to promote viral replication. Signal Transduct Target Ther. 2023;8(1):194. doi:10.1038/s41392-023-01459-8

930. Chun JY, Kim KJ, Hwang IT, Kim YJ, Lee DH, Lee IK, Kim JK. Dual priming oligonucleotide system for the multiplex detection of respiratory viruses and SNP genotyping of CYP2C19 gene. Nucleic Acids Res. 2007;35(6):e40. doi:10.1093/nar/gkm051

931. Chen S, Tang W, Yu G, Tang Z, Liu E. CXCL12/CXCR4 Axis is Involved in the Recruitment of NK Cells by HMGB1 Contributing to Persistent Airway Inflammation and AHR During the Late Stage of RSV Infection. J Microbiol. 2023;61(4):461–469. doi:10.1007/s12275-023-00018-8

932. Li Z, Qu X, Liu X, Huan C, Wang H, Zhao Z, Yang X, Hua S, Zhang W. GBP5 Is an Interferon-Induced Inhibitor of Respiratory Syncytial Virus. J Virol. 2020;94(21):e01407–20. doi:10.1128/JVI.01407-20

933. Karsli E, Sabirli R, Altintas E, Canacik O, Sabirli GT, Kaymaz B, Kurt Ö, Koseler A. Soluble P-selectin as a potential diagnostic and prognostic biomarker for COVID-19 disease: A case-control study. Life Sci. 2021;277:119634. doi:10.1016/j.lfs.2021.119634

934. Manley GCA, Stokes CA, Marsh EK, Sabroe I, Parker LC. DUSP10 Negatively Regulates the Inflammatory Response to Rhinovirus through Interleukin-1β Signaling. J Virol. 2019;93(2):e01659–18. doi:10.1128/JVI.01659-18

935. Cui J, Morgan D, Cheng DH, Foo SL, Yap GLR, Ampomah PB, Arora S, Sachaphibulkij K, Periaswamy B, Fairhurst AM, et al. RNA-Sequencing-Based Transcriptomic Analysis Reveals a Role for Annexin-A1 in Classical and Influenza A Virus-Induced Autophagy. Cells. 2020;9(6):1399. doi:10.3390/cells9061399

936. Wang H, Liu D, Sun Y, Meng C, Tan L, Song C, Qiu X, Liu W, Ding C, Ying L. Upregulation of DUSP6 impairs infectious bronchitis virus replication by negatively regulating ERK pathway and promoting apoptosis. Vet Res. 2021;52(1):7. doi:10.1186/s13567-020-00866-x

937. Wu X, Qi H, Yang Y, Yin Y, Ma D, Li H, Qu Y. Downregulation of matrix metalloproteinaseL19 induced by respiratory syncytial viral infection affects the interaction between epithelial cells and fibroblasts. Mol Med Rep. 2016;13(1):167–173. doi:10.3892/mmr.2015.4518

938. Kawakami E, Saiki N, Yoneyama Y, Moriya C, Maezawa M, Kawamura S, Kinebuchi A, Kono T, Funata M, Sakoda A, et al. Complement factor D targeting protects endotheliopathy in organoid and monkey models of COVID-19. Cell Stem Cell. 2023;30(10):1315–1330.e10. doi:10.1016/j.stem.2023.09.001

939. Malla R, Kamal MA. Tetraspanin-enriched Microdomain Containing CD151, CD9, and TSPAN 8 -Potential Mediators of Entry and Exit Mechanisms in Respiratory Viruses Including SARS-CoV-2. Curr Pharm Des. 2022;28(46):3649–3657. doi:10.2174/1381612828666220907105543

940. Hirakawa S, Kojima T, Obata K, Okabayashi T, Yokota S, Nomura K, Obonai T, Fuchimoto J, Himi T, Tsutsumi H, et al. Marked induction of matrix metalloproteinase-10 by respiratory syncytial virus infection in human nasal epithelial cells. J Med Virol. 2013;85(12):2141–2150. doi:10.1002/jmv.23718

941. Castelli EC, de Castro MV, Naslavsky MS, Scliar MO, Silva NSB, Pereira RN, Ciriaco VAO, Castro CFB, Mendes-Junior CT, Silveira ES, et al. MUC22, HLA-A, and HLA-DOB variants and COVID-19 in resilient super-agers from Brazil. Front Immunol. 2022;13:975918. doi:10.3389/fimmu.2022.975918

942. Lévy Y, Wiedemann A, Hejblum BP, Durand M, Lefebvre C, Surénaud M, Lacabaratz C, Perreau M, Foucat E, Déchenaud M, et al. CD177, a specific marker of neutrophil activation, is associated with coronavirus disease 2019 severity and death. iScience. 2021;24(7):102711. doi:10.1016/j.isci.2021.102711

943. Song Z, Bai J, Liu X, Nauwynck H, Wu J, Liu X, Jiang P. S100A9 regulates porcine reproductive and respiratory syndrome virus replication by interacting with the viral nucleocapsid protein. Vet Microbiol. 2019;239:108498. doi:10.1016/j.vetmic.2019.108498

944. Hao W, Wang L, Li S. FKBP5 Regulates RIG-I-Mediated NF-κB Activation and Influenza A Virus Infection. Viruses. 2020;12(6):672. doi:10.3390/v12060672

945. Zhang S, Wang J, Wang L, Aliyari S, Cheng G. SARS-CoV-2 virus NSP14 Impairs NRF2/HMOX1 activation by targeting Sirtuin 1. Cell Mol Immunol. 2022;19(8):872–882. doi:10.1038/s41423-022-00887-w

946. Derakhshani A, Hemmat N, Asadzadeh Z, Ghaseminia M, Shadbad MA, Jadideslam G, Silvestris N, Racanelli V, Baradaran B. Arginase 1 (Arg1) as an Up-Regulated Gene in COVID-19 Patients: A Promising Marker in COVID-19 Immunopathy. J Clin Med. 2021;10(5):1051. doi:10.3390/jcm10051051

947. Zhang L, Li M, Wang Z, Sun P, Wei S, Zhang C, Wu H, Bai H. Cardiovascular Risk After SARS-CoV-2 Infection Is Mediated by IL18/IL18R1/HIF-1 Signaling Pathway Axis. Front Immunol. 2022;12:780804. doi:10.3389/fimmu.2021.780804

948. de Moraes D, Paiva BVB, Cury SS, Ludwig RG, Junior JPA, Mori MADS, Carvalho RF. Prediction of SARS-CoV Interaction with Host Proteins during Lung Aging Reveals a Potential Role for TRIB3 in COVID-19. Aging Dis. 2021;12(1):42–49. doi:10.14336/AD.2020.1112

949. Vargas-Alarcón G, Posadas-Sánchez R, Ramírez-Bello J. Variability in genes related to SARS-CoV-2 entry into host cells (ACE2, TMPRSS2, TMPRSS11A, ELANE, and CTSL) and its potential use in association studies. Life Sci. 2020;260:118313. doi:10.1016/j.lfs.2020.118313

950. Zhang X, Chen Y, Li S, Wang J, He Z, Yan J, Liu X, Guo C. MARCO Inhibits Porcine Reproductive and Respiratory Syndrome Virus Infection through Intensifying Viral GP5-Induced Apoptosis. Microbiol Spectr. 2023;11(3):e0475322. doi:10.1128/spectrum.04753-22

951. Petersen BC, Dolgachev V, Rasky A, Lukacs NW. IL-17E (IL-25) and IL-17RB promote respiratory syncytial virus-induced pulmonary disease. J Leukoc Biol. 2014;95(5):809–815. doi:10.1189/jlb.0913482

952. Ercan H, Schrottmaier WC, Pirabe A, Schmuckenschlager A, Pereyra D, Santol J, Pawelka E, Traugott MT, Schörgenhofer C, Seitz T, et al. Platelet Phenotype Analysis of COVID-19 Patients Reveals Progressive Changes in the Activation of Integrin αIIbβ3, F13A1, the SARS-CoV-2 Target EIF4A1 and Annexin A5. Front Cardiovasc Med. 2021;8:779073. doi:10.3389/fcvm.2021.779073

953. Hornick EE, Dagvadorj J, Zacharias ZR, Miller AM, Langlois RA, Chen P, Legge KL, Bishop GA, Sutterwala FS, Cassel SL. Dendritic cell NLRC4 regulates influenza A virus-specific CD4 T cell responses through FasL expression. J Clin Invest. 2019;129(7):2888–2897. doi:10.1172/JCI124937

954. Pinkenburg O, Meyer T, Bannert N, Norley S, Bolte K, Czudai-Matwich V, Herold S, Gessner A, Schnare M. The Human Antimicrobial Protein Bactericidal/Permeability-Increasing Protein (BPI) Inhibits the Infectivity of Influenza A Virus. PLoS One. 2016;11(6):e0156929. doi:10.1371/journal.pone.0156929

955. Costa AJ, Lemes RMR, Bartolomeo CS, Nunes TA, Pereira GC, Oliveira RB, Gomes AL, Smaili SS, Maciel RMB, Newson L, et al. Overexpression of estrogen receptor GPER1 and G1 treatment reduces SARS-CoV-2 infection in BEAS-2B bronchial cells. Mol Cell Endocrinol. 2022;558:111775. doi:10.1016/j.mce.2022.111775

956. Ohol YM, Wang Z, Kemble G, Duke G. Direct Inhibition of Cellular Fatty Acid Synthase Impairs Replication of Respiratory Syncytial Virus and Other Respiratory Viruses. PLoS One. 2015;10(12):e0144648. doi:10.1371/journal.pone.0144648

957. Chi L, Shan Y, Cui Z. N-Acetyl-L-Cysteine Protects Airway Epithelial Cells during Respiratory Syncytial Virus Infection against Mucin Synthesis, Oxidative Stress, and Inflammatory Response and Inhibits HSPA6 Expression. Anal Cell Pathol (Amst). 2022;2022:4846336. doi:10.1155/2022/4846336

958. Guzmán-Beltrán S, Herrera MT, Torres M, Gonzalez Y. CD33 is downregulated by influenza virus H1N1pdm09 and induces ROS and the TNF-α, IL-1β, and IL-6 cytokines in human mononuclear cells. Braz J Microbiol. 2022;53(1):89–97. doi:10.1007/s42770-021-00663-4

959. Zhao MM, Zhu Y, Zhang L, Zhong G, Tai L, Liu S, Yin G, Lu J, He Q, Li MJ, et al. Novel cleavage sites identified in SARS-CoV-2 spike protein reveal mechanism for cathepsin L-facilitated viral infection and treatment strategies. Cell Discov. 2022;8(1):53. doi:10.1038/s41421-022-00419-w

960. Reading PC, Bozza S, Gilbertson B, Tate M, Moretti S, Job ER, Crouch EC, Brooks AG, Brown LE, Bottazzi B, et al. Antiviral activity of the long chain pentraxin PTX3 against influenza viruses. J Immunol. 2008;180(5):3391–3398. doi:10.4049/jimmunol.180.5.3391

961. Brand HK, Ahout IM, de Ridder D, van Diepen A, Li Y, Zaalberg M, Andeweg A, Roeleveld N, de Groot R, Warris A, et al. Olfactomedin 4 Serves as a Marker for Disease Severity in Pediatric Respiratory Syncytial Virus (RSV) Infection. PLoS One. 2015;10(7):e0131927. doi:10.1371/journal.pone.0131927

962. Kaltenborn E, Kern S, Frixel S, Fragnet L, Conzelmann KK, Zarbock R, Griese M. Respiratory syncytial virus potentiates ABCA3 mutation-induced loss of lung epithelial cell differentiation. Hum Mol Genet. 2012;21(12):2793–2806. doi:10.1093/hmg/dds107

963. Wang H, Du L, Liu F, Wei Z, Gao L, Feng WH. Highly Pathogenic Porcine Reproductive and Respiratory Syndrome Virus Induces Interleukin-17 Production via Activation of the IRAK1-PI3K-p38MAPK-C/EBPβ/CREB Pathways. J Virol. 2019;93(21):e01100–19. doi:10.1128/JVI.01100-19

964. Loh JT, Teo JKH, Lam KP. Dok3 restrains neutrophil production of calprotectin during TLR4 sensing of SARS-CoV-2 spike protein. Front Immunol. 2022;13:996637. doi:10.3389/fimmu.2022.996637

965. Sun D, Luthra P, Li Z, He B. PLK1 down-regulates parainfluenza virus 5 gene expression. PLoS Pathog. 2009;5(7):e1000525. doi:10.1371/journal.ppat.1000525

966. Schloer S, Hübel N, Masemann D, Pajonczyk D, Brunotte L, Ehrhardt C, Brandenburg LO, Ludwig S, Gerke V, Rescher U. The annexin A1/FPR2 signaling axis expands alveolar macrophages, limits viral replication, and attenuates pathogenesis in the murine influenza A virus infection model. FASEB J. 2019;33(11):12188–12199. doi:10.1096/fj.201901265

967. Zeng R, Zhang H, Hai Y, Cui Y, Wei L, Li N, Liu J, Li C, Liu Y. Interleukin-27 inhibits vaccine-enhanced pulmonary disease following respiratory syncytial virus infection by regulating cellular memory responses. J Virol. 2012;86(8):4505–4517. doi:10.1128/JVI.07091-11

968. Karabulut Uzunçakmak S, Naldan ME, Dirican E, Kerget F, Halıcı Z. Preliminary investigation of gene expression levels of PAPP-A, STC-2, and HIF-1α in SARS-Cov-2 infected patients. Mol Biol Rep. 2022;49(9):8693–8699. doi:10.1007/s11033-022-07710-9

969. Liu X, Liu X, Bai J, Gao Y, Song Z, Nauwynck H, Wang X, Yang Y, Jiang P. Glyceraldehyde-3-Phosphate Dehydrogenase Restricted in Cytoplasmic Location by Viral GP5 Facilitates Porcine Reproductive and Respiratory Syndrome Virus Replication via Its Glycolytic Activity. J Virol. 2021;95(18):e0021021. doi:10.1128/JVI.00210-21

970. Siekacz K, Kumor-Kisielewska A, Miłkowska-Dymanowska J, Pietrusińska M, Bartczak K, Majewski S, Stańczyk A, Piotrowski WJ, Białas AJ. Soluble ITGaM and ITGb2 Integrin Subunits Are Involved in Long-Term Pulmonary Complications after COVID-19 Infection. J Clin Med. 2023;12(1):342. doi:10.3390/jcm12010342

971. Sugamata R, Dobashi H, Nagao T, Yamamoto K, Nakajima N, Sato Y, Aratani Y, Oshima M, Sata T, Kobayashi K, et al. Contribution of neutrophil-derived myeloperoxidase in the early phase of fulminant acute respiratory distress syndrome induced by influenza virus infection. Microbiol Immunol. 2012;56(3):171–182. doi:10.1111/j.1348-0421.2011.00424.x

972. Menendez D, Snipe J, Marzec J, Innes CL, Polack FP, Caballero MT, Schurman SH, Kleeberger SR, Resnick MA. p53-responsive TLR8 SNP enhances human innate immune response to respiratory syncytial virus. J Clin Invest. 2019;129(11):4875–4884. doi:10.1172/JCI128626

973. McAuley JL, Corcilius L, Tan HX, Payne RJ, McGuckin MA, Brown LE. The cell surface mucin MUC1 limits the severity of influenza A virus infection. Mucosal Immunol. 2017;10(6):1581–1593. doi:10.1038/mi.2017.16

974. Pashkov E, Korchevaya E, Faizuloev E, Rtishchev A, Cherepovich B, Bystritskaya E, Sidorov A, Poddubikov A, Bykov A, Dronina Y, et al. Knockdown of FLT4, Nup98, and Nup205 Cellular Genes Effectively Suppresses the Reproduction of Influenza Virus Strain A/WSN/1933 (H1N1) In vitro. Infect Disord Drug Targets. 2022;22(5):e250322202629. doi:10.2174/1871526522666220325121403

975. Kellar GG, Barrow KA, Rich LM, Debley JS, Wight TN, Ziegler SF, Reeves SR. Loss of versican and production of hyaluronan in lung epithelial cells are associated with airway inflammation during RSV infection. J Biol Chem. 2021;296:100076. doi:10.1074/jbc.RA120.016196

976. Cui L, Mahesutihan M, Zheng W, Meng L, Fan W, Li J, Ye X, Liu W, Sun L. CDC25B promotes influenza A virus replication by regulating the phosphorylation of nucleoprotein. Virology. 2018;525:40–47. doi:10.1016/j.virol.2018.09.005

977. Zhang Y, Federation AJ, Kim S, O’Keefe JP, Lun M, Xiang D, Brown JD, Steinhauser ML. Targeting nuclear receptor NR4A1-dependent adipocyte progenitor quiescence promotes metabolic adaptation to obesity. J Clin Invest. 2018;128(11):4898–4911. doi:10.1172/JCI98353

978. Sindhu S, Thomas R, Shihab P, Sriraman D, Behbehani K, Ahmad R. Obesity Is a Positive Modulator of IL-6R and IL-6 Expression in the Subcutaneous Adipose Tissue: Significance for Metabolic Inflammation. PLoS One. 2015;10(7):e0133494. doi:10.1371/journal.pone.0133494

979. Lima RS, Mattos RT, Medeiros NI, Kattah FM, Nascimento JRS, Menezes CA, Rios-Santos F, Dutra WO, Gomes JAS, Moreira PR. CXCL8 expression and methylation are correlated with anthropometric and metabolic parameters in childhood obesity. Cytokine. 2021;143:155538. doi:10.1016/j.cyto.2021.155538

980. Yamagata AS, Rizzo LB, Cerqueira RO, Scott J, Cordeiro Q, McIntyre RS, Mansur RB, Brietzke E. Differential Impact of Obesity on CD69 Expression in Individuals with Bipolar Disorder and Healthy Controls. Mol Neuropsychiatry. 2018;3(4):192–196. doi:10.1159/000486396

981. Caracciolo V, Young J, Gonzales D, Ni Y, Flowers SJ, Summer R, Waldman SA, Kim JK, Jung DY, Noh HL, et al. Myeloid-specific deletion of Zfp36 protects against insulin resistance and fatty liver in diet-induced obese mice. Am J Physiol Endocrinol Metab. 2018;315(4):E676–E693. doi:10.1152/ajpendo.00224.2017

982. Nunemaker CS, Chung HG, Verrilli GM, Corbin KL, Upadhye A, Sharma PR. Increased serum CXCL1 and CXCL5 are linked to obesity, hyperglycemia, and impaired islet function. J Endocrinol. 2014;222(2):267–276. doi:10.1530/JOE-14-0126

983. Yang HL, Feng M, Tan X, Yan GY, Sun C. The role of SOCS2 in recombinant human growth hormone (rhGH) regulating lipid metabolism in high-fat-diet-induced obesity mice. Mol Biol Rep. 2013;40(3):2319–2326. doi:10.1007/s11033-012-2313-5

984. Terán-Cabanillas E, Hernández J. Role of Leptin and SOCS3 in Inhibiting the Type I Interferon Response During Obesity. Inflammation. 2017;40(1):58–67. doi:10.1007/s10753-016-0452-x

985. Chan PC, Hung LM, Huang JP, Day YJ, Yu CL, Kuo FC, Lu CH, Tian YF, Hsieh PS. Augmented CCL5/CCR5 signaling in brown adipose tissue inhibits adaptive thermogenesis and worsens insulin resistance in obesity. Clin Sci (Lond). 2022;136(1):121–137. doi:10.1042/CS20210959

986. Widjaja NA, Caesar LA, Nova S, Ardianah E. Beyond the Scale: Investigating Adiponectin, ICAM-1, and VCAM-1 as Metabolic Markers in Obese Adolescents with Metabolic Syndrome. J Obes. 2023;2023:4574042. doi:10.1155/2023/4574042

987. Dommel S, Blüher M. Does C-C Motif Chemokine Ligand 2 (CCL2) Link Obesity to a Pro-Inflammatory State?. Int J Mol Sci. 2021;22(3):1500. doi:10.3390/ijms22031500

988. Gan M, Shen L, Wang S, Guo Z, Zheng T, Tan Y, Fan Y, Liu L, Chen L, Jiang A, et al. Genistein inhibits high fat diet-induced obesity through miR-222 by targeting BTG2 and adipor1. Food Funct. 2020;11(3):2418–2426. doi:10.1039/c9fo00861f

989. Shapira S, Kazanov D, Dankner R, Fishman S, Stern N, Arber N. High Expression Level of PPARγ in CD24 Knockout Mice and Gender-Specific Metabolic Changes: A Model of Insulin-Sensitive Obesity. J Pers Med. 2021;11(1):50. doi:10.3390/jpm11010050

990. Wara AK, Wang S, Wu C, Fang F, Haemmig S, Weber BN, Aydogan CO, Tesmenitsky Y, Aliakbarian H, Hawse JR, et al. KLF10 Deficiency in CD4+ T Cells Triggers Obesity, Insulin Resistance, and Fatty Liver. Cell Rep. 2020;33(13):108550. doi:10.1016/j.celrep.2020.108550

991. Kupcinskiene K, Murnikovaite M, Varkalaite G, Juzenas S, Trepenaitis D, Petereit R, Maleckas A, Kupcinskas J, Macas A. Thrombosis Related ABO, F5, MTHFR, and FGG Gene Polymorphisms in Morbidly Obese Patients. Dis Markers. 2016;2016:7853424. doi:10.1155/2016/7853424

992. Lancaster GI, Kraakman MJ, Kammoun HL, Langley KG, Estevez E, Banerjee A, Grumont RJ, Febbraio MA, Gerondakis S. The dual-specificity phosphatase 2 (DUSP2) does not regulate obesity-associated inflammation or insulin resistance in mice. PLoS One. 2014;9(11):e111524. doi:10.1371/journal.pone.0111524

993. Khadir A, Tiss A, Abubaker J, Abu-Farha M, Al-Khairi I, Cherian P, John J, Kavalakatt S, Warsame S, Al-Madhoun A, et al. MAP kinase phosphatase DUSP1 is overexpressed in obese humans and modulated by physical exercise. Am J Physiol Endocrinol Metab. 2015;308(1):E71–E83. doi:10.1152/ajpendo.00577.2013

994. Sano T, Nagayasu S, Suzuki S, Iwashita M, Yamashita A, Shinjo T, Sanui T, Kushiyama A, Kanematsu T, Asano T, et al. Epicatechin downregulates adipose tissue CCL19 expression and thereby ameliorates diet-induced obesity and insulin resistance. Nutr Metab Cardiovasc Dis. 2017;27(3):249–259. doi:10.1016/j.numecd.2016.11.008

995. De Los Santos S, Reyes-Castro LA, Coral-Vázquez RM, Mendez JP, Zambrano E, Canto P. (-)-Epicatechin increases apelin/APLNR expression and modifies proteins involved in lipid metabolism of offspring descendants of maternal obesity. J Nutr Biochem. 2023;117:109350. doi:10.1016/j.jnutbio.2023.109350

996. Maculewicz E, Antkowiak B, Antkowiak O, Borecka A, Mastalerz A, Leońska-Duniec A, Humińska-Lisowska K, Michałowska-Sawczyn M, Garbacz A, Lorenz K, et al. The interactions between interleukin-1 family genes: IL1A, IL1B, IL1RN, and obesity parameters. BMC Genomics. 2022;23(1):112. doi:10.1186/s12864-021-08258-x

997. Lopez-Legarrea P, Mansego ML, Zulet MA, Martinez JA. SERPINE1, PAI-1 protein coding gene, methylation levels and epigenetic relationships with adiposity changes in obese subjects with metabolic syndrome features under dietary restriction. J Clin Biochem Nutr. 2013;53(3):139–144. doi:10.3164/jcbn.13-54

998. Piquer-Garcia I, Campderros L, Taxerås SD, Gavaldà-Navarro A, Pardo R, Vila M, Pellitero S, Martínez E, Tarascó J, Moreno P, et al. A Role for Oncostatin M in the Impairment of Glucose Homeostasis in Obesity. J Clin Endocrinol Metab. 2020;105(3):e337–e348. doi:10.1210/clinem/dgz090

999. Suriyaprom K, Pheungruang B, Tungtrongchitr R, Sroijit OY. Relationships of apelin concentration and APLN T-1860C polymorphism with obesity in Thai children. BMC Pediatr. 2020;20(1):455. doi:10.1186/s12887-020-02350-z

1000. Gustafson B, Smith U. The WNT inhibitor Dickkopf 1 and bone morphogenetic protein 4 rescue adipogenesis in hypertrophic obesity in humans. Diabetes. 2012;61(5):1217–1224. doi:10.2337/db11-1419

1001. Deng Y, Qiu T, Zhang M, Wu J, Zhang X, Wang J, Chen K, Feng J, Ha X, Xie J, et al. High Level of Palmitic Acid Induced Over-Expressed Methyltransferase Inhibits Anti-Inflammation Factor KLF4 Expression in Obese Status. Inflammation. 2020;43(3):821–832. doi:10.1007/s10753-019-01168-x

1002. Bradley D, Smith AJ, Blaszczak A, Shantaram D, Bergin SM, Jalilvand A, Wright V, Wyne KL, Dewal RS, Baer LA, et al. Interferon gamma mediates the reduction of adipose tissue regulatory T cells in human obesity. Nat Commun. 2022;13(1):5606. doi:10.1038/s41467-022-33067-5

1003. Sommer J, Ehnis H, Seitz T, Schneider J, Wild AB, Moceri S, Buechler C, Bozec A, Weber GF, Merkel S, et al. Four-and-a-Half LIM-Domain Protein 2 (FHL2) Induces Neuropeptide Y (NPY) in Macrophages in Visceral Adipose Tissue and Promotes Diet-Induced Obesity. Int J Mol Sci. 2023;24(19):14943. doi:10.3390/ijms241914943

1004. Banerjee J, Dorfman MD, Fasnacht R, Douglass JD, Wyse-Jackson AC, Barria A, Thaler JP. CX3CL1 Action on Microglia Protects from Diet-Induced Obesity by Restoring POMC Neuronal Excitability and Melanocortin System Activity Impaired by High-Fat Diet Feeding. Int J Mol Sci. 2022;23(12):6380. doi:10.3390/ijms23126380

1005. Ribeiro SMTL, Lopes LR, Paula Costa G, Figueiredo VP, Shrestha D, Batista AP, Nicolato RLC, Oliveira FLP, Gomes JAS, Talvani A. et al. CXCL-16, IL-17, and bone morphogenetic protein 2 (BMP-2) are associated with overweight and obesity conditions in middle-aged and elderly women. Immun Ageing. 2017;14:6. doi:10.1186/s12979-017-0089-0

1006. Du Y, Li S, Cui CJ, Zhang Y, Yang SH, Li JJ. Leptin decreases the expression of low-density lipoprotein receptor via PCSK9 pathway: linking dyslipidemia with obesity. J Transl Med. 2016;14(1):276. doi:10.1186/s12967-016-1032-4

1007. Sano T, Sanada T, Sotomaru Y, Shinjo T, Iwashita M, Yamashita A, Fukuda T, Sanui T, Asano T, Kanematsu T, et al. Ccr7 null mice are protected against diet-induced obesity via Ucp1 upregulation and enhanced energy expenditure. Nutr Metab (Lond). 2019;16:43. doi:10.1186/s12986-019-0372-5

1008. Hoffmann JM, Grünberg JR, Church C, Elias I, Palsdottir V, Jansson JO, Bosch F, Hammarstedt A, Hedjazifar S, Smith U. BMP4 Gene Therapy in Mature Mice Reduces BAT Activation but Protects from Obesity by Browning Subcutaneous Adipose Tissue. Cell Rep. 2017;20(5):1038–1049. doi:10.1016/j.celrep.2017.07.020

1009. Ugi S, Maeda S, Kawamura Y, Kobayashi MA, Imamura M, Yoshizaki T, Morino K, Sekine O, Yamamoto H, Tani T, et al. CCDC3 is specifically upregulated in omental adipose tissue in subjects with abdominal obesity. Obesity (Silver Spring). 2014;22(4):1070–1077. doi:10.1002/oby.20645

1010. Carpagnano GE, Spanevello A, Sabato R, Depalo A, Palladino GP, Bergantino L, Foschino Barbaro MP. Systemic and airway inflammation in sleep apnea and obesity: the role of ICAM-1 and IL-8. Transl Res. 2010;155(1):35–43. doi:10.1016/j.trsl.2009.09.004

1011. Kinik ST, Ozbek N, Yücel M, Haberal A, Cetintas S. Correlations among serum leptin levels, complete blood count parameters and peripheral CD34(+) cell count in prepubertal obese children. Ann Hematol. 2005;84(9):605–608. doi:10.1007/s00277-005-1064-y

1012. Tang H, Liu N, Feng X, Yang Y, Fang Y, Zhuang S, Dai Y, Liu M, Tang L. Circulating levels of IL-33 are elevated by obesity and positively correlated with metabolic disorders in Chinese adults. J Transl Med. 2021;19(1):52. doi:10.1186/s12967-021-02711-x

1013. Li M, Liu L, Kang Y, Huang S, Xiao Y. Circulating THBS1: A Risk Factor for Nonalcoholic Fatty Liver Disease in Obese Children. Ann Nutr Metab. 2023;79(1):16–28. doi:10.1159/000527780

1014. Blüher M, Klöting N, Wueest S, Schoenle EJ, Schön MR, Dietrich A, Fasshauer M, Stumvoll M, Konrad D. Fas and FasL expression in human adipose tissue is related to obesity, insulin resistance, and type 2 diabetes. J Clin Endocrinol Metab. 2014;99(1):E36–E44. doi:10.1210/jc.2013-2488

1015. Clapcote SJ. Phosphodiesterase-4B as a Therapeutic Target for Cognitive Impairment and Obesity-Related Metabolic Diseases. Adv Neurobiol. 2017;17:103–131. doi:10.1007/978-3-319-58811-7_5

1016. Olszanecka-Glinianowicz M, Zahorska-Markiewicz B, Janowska J, Zurakowski A. Serum concentrations of nitric oxide, tumor necrosis factor (TNF)-alpha and TNF soluble receptors in women with overweight and obesity. Metabolism. 2004;53(10):1268–1273. doi:10.1016/j.metabol.2004.07.001

1017. Siddiqui JA, Pothuraju R, Khan P, Sharma G, Muniyan S, Seshacharyulu P, Jain M, Nasser MW, Batra SK. Pathophysiological role of growth differentiation factor 15 (GDF15) in obesity, cancer, and cachexia. Cytokine Growth Factor Rev. 2022;64:71–83. doi:10.1016/j.cytogfr.2021.11.002

1018. Hueso L, Marques P, Morant B, Gonzalez-Navarro H, Ortega J, Real JT, Sanz MJ, Piqueras L. CCL17 and CCL22 chemokines are upregulated in human obesity and play a role in vascular dysfunction. Front Endocrinol (Lausanne). 2023;14:1154158. doi:10.3389/fendo.2023.1154158

1019. Chan PC, Wu TN, Chen YC, Lu CH, Wabitsch M, Tian YF, Hsieh PS. Targetted inhibition of CD74 attenuates adipose COX-2-MIF-mediated M1 macrophage polarization and retards obesity-related adipose tissue inflammation and insulin resistance. Clin Sci (Lond). 2018;132(14):1581–1596. doi:10.1042/CS20180041

1020. Haenisch M, Nguyen T, Fihn CA, Goldstein AS, Amory JK, Treuting P, Brabb T, Paik J. Investigation of an ALDH1A1-specific inhibitor for suppression of weight gain in a diet-induced mouse model of obesity. Int J Obes (Lond). 2021;45(7):1542–1552. doi:10.1038/s41366-021-00818-1

1021. Kochumon S, Al Madhoun A, Al-Rashed F, Thomas R, Sindhu S, Al-Ozairi E, Al-Mulla F, Ahmad R. Elevated adipose tissue associated IL-2 expression in obesity correlates with metabolic inflammation and insulin resistance. Sci Rep. 2020;10(1):16364. doi:10.1038/s41598-020-73347-y

1022. Liong S, Barker G, Lappas M. Placental Pim-1 expression is increased in obesity and regulates cytokine-and toll-like receptor-mediated inflammation. Placenta. 2017;53:101–112. doi:10.1016/j.placenta.2017.04.010

1023. Duggan BM, Singh AM, Chan DY, Schertzer JD. Postbiotics engage IRF4 in adipocytes to promote sex-dependent changes in blood glucose during obesity. Physiol Rep. 2022;10(16):e15439. doi:10.14814/phy2.15439

1024. Gruber S, Straub BK, Ackermann PJ, Wunderlich CM, Mauer J, Seeger JM, Büning H, Heukamp L, Kashkar H, Schirmacher P, et al. Obesity promotes liver carcinogenesis via Mcl-1 stabilization independent of IL-6Rα signaling. Cell Rep. 2013;4(4):669–680. doi:10.1016/j.celrep.2013.07.023

1025. Osorio-Conles Ó, Ibarzabal A, Balibrea JM, Vidal J, Ortega E, de Hollanda A. FABP4 Expression in Subcutaneous Adipose Tissue Is Independently Associated with Circulating Triglycerides in Obesity. J Clin Med. 2023;12(3):1013. doi:10.3390/jcm12031013

1026. Moriyama B, Jarosinski PF, Figg WD, Henning SA, Danner RL, Penzak SR, Wayne AS, Walsh TJ. Pharmacokinetics of intravenous voriconazole in obese patients: implications of CYP2C19 homozygous poor metabolizer genotype. Pharmacotherapy. 2013;33(3):e19–e22. doi:10.1002/phar.1192

1027. Quirós PM, Ramsay AJ, Sala D, Fernández-Vizarra E, Rodríguez F, Peinado JR, Fernández-García MS, Vega JA, Enríquez JA, Zorzano A, et al. Loss of mitochondrial protease OMA1 alters processing of the GTPase OPA1 and causes obesity and defective thermogenesis in mice. EMBO J. 2012;31(9):2117–2133. doi:10.1038/emboj.2012.70

1028. Saha A, Ahn S, Blando J, Su F, Kolonin MG, DiGiovanni J. Proinflammatory CXCL12-CXCR4/CXCR7 Signaling Axis Drives Myc-Induced Prostate Cancer in Obese Mice. Cancer Res. 2017;77(18):5158–5168. doi:10.1158/0008-5472.CAN-17-0284

1029. Chen J, Xu X, Li Y, Li F, Zhang J, Xu Q, Chen W, Wei Y, Wang X. Kdm6a suppresses the alternative activation of macrophages and impairs energy expenditure in obesity. Cell Death Differ. 2021;28(5):1688–1704. doi:10.1038/s41418-020-00694-8

1030. Sato C, Shikata K, Hirota D, Sasaki M, Nishishita S, Miyamoto S, Kodera R, Ogawa D, Tone A, Kataoka HU, et al. P-selectin glycoprotein ligand-1 deficiency is protective against obesity-related insulin resistance. Diabetes. 2011;60(1):189–199. doi:10.2337/db09-1894

1031. Zhang J, Chen Y, Yan L, Zhang X, Zheng X, Qi J, Yang F, Li J. EphA3 deficiency in the hypothalamus promotes high-fat diet-induced obesity in mice. J Biomed Res. 2022;37(3):179–193. doi:10.7555/JBR.36.20220168

1032. Sha H, He Y, Yang L, Qi L. Stressed out about obesity: IRE1α-XBP1 in metabolic disorders. Trends Endocrinol Metab. 2011;22(9):374–381. doi:10.1016/j.tem.2011.05.002

1033. Pietrani NT, Ferreira CN, Rodrigues KF, Perucci LO, Carneiro FS, Bosco AA, Oliveira MC, Pereira SS, Teixeira AL, Alvarez-Leite JI, et al. Proresolving protein Annexin A1: The role in type 2 diabetes mellitus and obesity. Biomed Pharmacother. 2018;103:482–489. doi:10.1016/j.biopha.2018.04.024

1034. Liu R, Peters M, Urban N, Knowlton J, Napierala T, Gabrysiak J. Mice lacking DUSP6/8 have enhanced ERK1/2 activity and resistance to diet-induced obesity. Biochem Biophys Res Commun. 2020;533(1):17–22. doi:10.1016/j.bbrc.2020.08.106

1035. Shayo SC, Ogiso K, Kawade S, Hashiguchi H, Deguchi T, Nishio Y. Dietary obesity and glycemic excursions cause a parallel increase in STEAP4 and pro-inflammatory gene expression in murine PBMCs. Diabetol Int. 2021;13(2):358–371. doi:10.1007/s13340-021-00542-1

1036. Pendás AM, Folgueras AR, Llano E, Caterina J, Frerard F, Rodríguez F, Astudillo A, Noël A, Birkedal-Hansen H, López-Otín C. Diet-induced obesity and reduced skin cancer susceptibility in matrix metalloproteinase 19-deficient mice. Mol Cell Biol. 2004;24(12):5304–5313. doi:10.1128/MCB.24.12.5304-5313.2004

1037. Catalán V, Domench P, Gómez-Ambrosi J, Ramírez B, Becerril S, Mentxaka A, Rodríguez A, Valentí V, Moncada R, Baixauli J, et al. Dermatopontin Influences the Development of Obesity-Associated Colon Cancer by Changes in the Expression of Extracellular Matrix Proteins. Int J Mol Sci. 2022;23(16):9222. doi:10.3390/ijms23169222

1038. Mathews JA, Wurmbrand AP, Ribeiro L, Neto FL, Shore SA. Induction of IL-17A Precedes Development of Airway Hyperresponsiveness during Diet-Induced Obesity and Correlates with Complement Factor D. Front Immunol. 2014;5:440. doi:10.3389/fimmu.2014.00440

1039. Peng H, Zhang Q, Shen H, Liu Y, Chao X, Tian H, Cai X, Jin J. Association between serum soluble corin and obesity in Chinese adults: a cross-sectional study. Obesity (Silver Spring). 2015;23(4):856–861. doi:10.1002/oby.21016

1040. Khaidakov M, Mitra S, Kang BY, Wang X, Kadlubar S, Novelli G, Raj V, Winters M, Carter WC, Mehta JL. Oxidized LDL receptor 1 (OLR1) as a possible link between obesity, dyslipidemia and cancer. PLoS One. 2011;6(5):e20277. doi:10.1371/journal.pone.0020277

1041. Lijnen HR, Van Hoef B, Rodriguez JA, Paramo JA. Stromelysin-2 (MMP-10) deficiency does not affect adipose tissue formation in a mouse model of nutritionally induced obesity. Biochem Biophys Res Commun. 2009;389(2):378–381. doi:10.1016/j.bbrc.2009.08.170

1042. Nishimura S, Nagasaki M, Okudaira S, Aoki J, Ohmori T, Ohkawa R, Nakamura K, Igarashi K, Yamashita H, Eto K, et al. ENPP2 contributes to adipose tissue expansion and insulin resistance in diet-induced obesity. Diabetes. 2014;63(12):4154–4164. doi:10.2337/db13-1694

1043. Parikh D, Riascos-Bernal DF, Egaña-Gorroño L, Jayakumar S, Almonte V, Chinnasamy P, Sibinga NES. Allograft inflammatory factor-1-like is not essential for age dependent weight gain or HFD-induced obesity and glucose insensitivity. Sci Rep. 2020;10(1):3594. doi:10.1038/s41598-020-60433-4

1044. Yamaguchi S, Zhang D, Katayama A, Kurooka N, Sugawara R, Albuayjan HHH, Nakatsuka A, Eguchi J, Wada J. Adipocyte-Specific Inhibition of Mir221/222 Ameliorates Diet-Induced Obesity Through Targeting Ddit4. Front Endocrinol (Lausanne). 2022;12:750261. doi:10.3389/fendo.2021.750261

1045. Pan X, Yang L, Wang S, Liu Y, Yue L, Chen S. Semaglutide ameliorates obesity-induced cardiac inflammation and oxidative stress mediated via reduction of neutrophil Cxcl2, S100a8, and S100a9 expression. Mol Cell Biochem. 2023. doi:10.1007/s11010-023-04784-2

1046. Willmer T, Oosthuizen A, Dias S, Mendham AE, Goedecke JH, Pheiffer C. A pilot investigation of genetic and epigenetic variation of FKBP5 and response to exercise intervention in African women with obesity. Sci Rep. 2022;12(1):11771. doi:10.1038/s41598-022-15678-6

1047. Jiménez-Osorio AS, González-Reyes S, García-Niño WR, Moreno-Macías H, Rodríguez-Arellano ME, Vargas-Alarcón G, Zúñiga J, Barquera R, Pedraza-Chaverri J. Association of Nuclear Factor-Erythroid 2-Related Factor 2, Thioredoxin Interacting Protein, and Heme Oxygenase-1 Gene Polymorphisms with Diabetes and Obesity in Mexican Patients. Oxid Med Cell Longev. 2016;2016:7367641. doi:10.1155/2016/7367641

1048. Bhatta A, Yao L, Xu Z, Toque HA, Chen J, Atawia RT, Fouda AY, Bagi Z, Lucas R, Caldwell RB, et al. Obesity-induced vascular dysfunction and arterial stiffening requires endothelial cell arginase 1. Cardiovasc Res. 2017;113(13):1664–1676. doi:10.1093/cvr/cvx164

1049. Shen J, Song R, Ye Y, Wu X, Chow WH, Zhao H. HIF3A DNA methylation, obesity and weight gain, and breast cancer risk among Mexican American women. Obes Res Clin Pract. 2020;14(6):548–553. doi:10.1016/j.orcp.2020.10.001

1050. Mirea AM, Stienstra R, Kanneganti TD, Tack CJ, Chavakis T, Toonen EJM, Joosten LAB. Mice Deficient in the IL-1β Activation Genes Prtn3, Elane, and Casp1 Are Protected Against the Development of Obesity-Induced NAFLD. Inflammation. 2020;43(3):1054–1064. doi:10.1007/s10753-020-01190-4

1051. Lee SK, Park CY, Kim J, Kim D, Choe H, Kim JH, Hong JP, Lee YJ, Heo Y, Park HS, et al. TRIB3 Is Highly Expressed in the Adipose Tissue of Obese Patients and Is Associated With Insulin Resistance. J Clin Endocrinol Metab. 2022;107(3):e1057–e1073. doi:10.1210/clinem/dgab780

1052. Ushakumari CJ, Zhou QL, Wang YH, Na S, Rigor MC, Zhou CY, Kroll MK, Lin BD, Jiang ZY. Neutrophil Elastase Increases Vascular Permeability and Leukocyte Transmigration in Cultured Endothelial Cells and Obese Mice. Cells. 2022;11(15):2288. doi:10.3390/cells11152288

1053. Park YJ, Choe SS, Sohn JH, Kim JB. The role of glucose-6-phosphate dehydrogenase in adipose tissue inflammation in obesity. Adipocyte. 2017;6(2):147–153. doi:10.1080/21623945.2017.1288321

1054. Stefanski A, Majkowska L, Ciechanowicz A, Frankow M, Safranow K, Parczewski M, Pilarska K. The common C49620T polymorphism in the sulfonylurea receptor gene (ABCC8), pancreatic beta cell function and long-term diabetic complications in obese patients with long-lasting type 2 diabetes mellitus. Exp Clin Endocrinol Diabetes. 2007;115(5):317–321. doi:10.1055/s-2007-967086

1055. Kaartinen MT, Arora M, Heinonen S, Rissanen A, Kaprio J, Pietiläinen KH. Transglutaminases and Obesity in Humans: Association of F13A1 to Adipocyte Hypertrophy and Adipose Tissue Immune Response. Int J Mol Sci. 2020;21(21):8289. doi:10.3390/ijms21218289

1056. Guarino BD, Dado CD, Kumar A, Braza J, Harrington EO, Klinger JR. Deletion of the Npr3 gene increases severity of acute lung injury in obese mice. Pulm Circ. 2023;13(3):e12270. doi:10.1002/pul2.12270

1057. Kolb R, Phan L, Borcherding N, Liu Y, Yuan F, Janowski AM, Xie Q, Markan KR, Li W, Potthoff MJ, et al. Obesity-associated NLRC4 inflammasome activation drives breast cancer progression. Nat Commun. 2016;7:13007. doi:10.1038/ncomms13007

1058. Pervanidou P, Chouliaras G, Akalestos A, Bastaki D, Apostolakou F, Papassotiriou I, Chrousos GP. Increased placental growth factor (PlGF) concentrations in children and adolescents with obesity and the metabolic syndrome. Hormones (Athens). 2014;13(3):369–374. doi:10.14310/horm.2002.1491

1059. Nuncia-Cantarero M, Martinez-Canales S, Andrés-Pretel F, Santpere G, Ocaña A, Galan-Moya EM. Functional transcriptomic annotation and protein-protein interaction network analysis identify NEK2, BIRC5, and TOP2A as potential targets in obese patients with luminal A breast cancer. Breast Cancer Res Treat. 2018;168(3):613–623. doi:10.1007/s10549-017-4652-3

1060. Sun D, Zhao T, Zhang Q, Wu M, Zhang Z. Fat mass and obesity-associated protein regulates lipogenesis via m6 A modification in fatty acid synthase mRNA. Cell Biol Int. 2021;45(2):334–344. doi:10.1002/cbin.11490

1061. Abass MK, Al Shamsi A, Jan I, Masalawala MSY, Deeb A. Combined SPINK1 mutations induce early-onset severe chronic pancreatitis in a child with severe obesity. Endocrinol Diabetes Metab Case Rep. 2022. doi:10.1530/EDM-22-0273

1062. Tan L, Armstrong AR, Rosas S, Patel CM, Vander Wiele SS, Willey JS, Carlson CS, Yammani RR. Nuclear protein-1 is the common link for pathways activated by aging and obesity in chondrocytes: A potential therapeutic target for osteoarthritis. FASEB J. 2023;37(9):e23133. doi:10.1096/fj.202201700RR

1063. Xi P, Zhu W, Zhang Y, Wang M, Liang H, Wang H, Tian D. Upregulation of hypothalamic TRPV4 via S100a4/AMPKα signaling pathway promotes the development of diet-induced obesity. Biochim Biophys Acta Mol Basis Dis. 2024;1870(1):166883. doi:10.1016/j.bbadis.2023.166883

1064. Zhou Q, Zhu Y, Li C, Li Z, Tang Z, Yuan B, Wang X, Zhang S, Wu X. Elevated CTSL Gene Expression Correlated with Proinflammatory Cytokines in Omental Adipose Tissue of Patients with Obesity. Diabetes Metab Syndr Obes. 2022;15:2277–2285. doi:10.2147/DMSO.S373203

1065. Pérez-Díaz S, Koumaiha Z, Borok MJ, Aurade F, Pini M, Periou B, Rouault C, Baba-Amer Y, Clément K, Derumeaux G, et al. Obesity impairs skeletal muscle repair through NID-1 mediated extracellular matrix remodeling by mesenchymal progenitors. Matrix Biol. 2022;112:90–115. doi:10.1016/j.matbio.2022.08.006

1066. Yiew NKH, Chatterjee TK, Tang YL, Pellenberg R, Stansfield BK, Bagi Z, Fulton DJ, Stepp DW, Chen W, Patel V, et al. A novel role for the Wnt inhibitor APCDD1 in adipocyte differentiation: Implications for diet-induced obesity. J Biol Chem. 2017;292(15):6312–6324. doi:10.1074/jbc.M116.758078

1067. Thoudam T, Ha CM, Leem J, Chanda D, Park JS, Kim HJ, Jeon JH, Choi YK, Liangpunsakul S, Huh YH, et al. PDK4 Augments ER-Mitochondria Contact to Dampen Skeletal Muscle Insulin Signaling During Obesity. Diabetes. 2019;68(3):571–586. doi:10.2337/db18-0363

1068. Maixner N, Haim Y, Blüher M, Chalifa-Caspi V, Veksler-Lublinsky I, Makarenkov N, Yoel U, Bashan N, Liberty IF, Kukeev I, et al. Visceral Adipose Tissue E2F1-miRNA206/210 Pathway Associates with Type 2 Diabetes in Humans with Extreme Obesity. Cells. 2022;11(19):3046. doi:10.3390/cells11193046

1069. Hu Y, Xu J, Gao R, Xu Y, Huangfu B, Asakiya C, Huang X, Zhang F, Huang K, He X, et al. Diallyl Trisulfide Prevents Adipogenesis and Lipogenesis by Regulating the Transcriptional Activation Function of KLF15 on PPARγ to Ameliorate Obesity. Mol Nutr Food Res. 2022;66(22):e2200173. doi:10.1002/mnfr.202200173

1070. Bonacina F, Moregola A, Porte R, Baragetti A, Bonavita E, Salatin A, Grigore L, Pellegatta F, Molgora M, Sironi M, et al. Pentraxin 3 deficiency protects from the metabolic inflammation associated to diet-induced obesity. Cardiovasc Res. 2019;115(13):1861–1872. doi:10.1093/cvr/cvz068

1071. Menghini R, Marchetti V, Cardellini M, Hribal ML, Mauriello A, Lauro D, Sbraccia P, Lauro R, Federici M. Phosphorylation of GATA2 by Akt increases adipose tissue differentiation and reduces adipose tissue-related inflammation: a novel pathway linking obesity to atherosclerosis. Circulation. 2005;111(15):1946–1953. doi:10.1161/01.CIR.0000161814.02942.B2

1072. Albuquerque D, Nóbrega C, Rodríguez-López R, Manco L. Association study of common polymorphisms in MSRA, TFAP2B, MC4R, NRXN3, PPARGC1A, TMEM18, SEC16B, HOXB5 and OLFM4 genes with obesity-related traits among Portuguese children. J Hum Genet. 2014;59(6):307–313. doi:10.1038/jhg.2014.23

1073. Wang C, Murphy J, Delaney KZ, Khor N, Morais JA, Tsoukas MA, Lowry DE, Mutch DM, Santosa S. ssociation between rs174537 FADS1 polymorphism and immune cell profiles in abdominal and femoral subcutaneous adipose tissue: an exploratory study in adults with obesity. Adipocyte. 2021;10(1):124–130. doi:10.1080/21623945.2021.1888470

1074. Habibovic A, Hristova M, Morris CR, Lin MJ, Cruz LC, Ather JL, Geiszt M, Anathy V, Janssen-Heininger YMW, Poynter ME, et al. Diet-induced obesity worsens allergen-induced type 2/type 17 inflammation in airways by enhancing DUOX1 activation. Am J Physiol Lung Cell Mol Physiol. 2023;324(2):L228–L242. doi:10.1152/ajplung.00331.2022

1075. Ahmad R, Shihab PK, Thomas R, Alghanim M, Hasan A, Sindhu S, Behbehani K. Increased expression of the interleukin-1 receptor-associated kinase (IRAK)-1 is associated with adipose tissue inflammatory state in obesity. Diabetol Metab Syndr. 2015;7:71. doi:10.1186/s13098-015-0067-7

1076. Smith DC, Karahan H, Wijeratne HRS, Al-Amin M, McCord B, Moon Y, Kim J. Deletion of the Alzheimer’s disease risk gene Abi3 locus results in obesity and systemic metabolic disruption in mice. Front Aging Neurosci. 2022;14:1035572. doi:10.3389/fnagi.2022.1035572

1077. Elias I, Ferré T, Vilà L, Muñoz S, Casellas A, Garcia M, Molas M, Agudo J, Roca C, Ruberte J, et al. ALOX5AP Overexpression in Adipose Tissue Leads to LXA4 Production and Protection Against Diet-Induced Obesity and Insulin Resistance. Diabetes. 2016;65(8):2139–2150. doi:10.2337/db16-0040

1078. Guizar-Heredia R, Tovar AR, Granados-Portillo O, Pichardo-Ontiveros E, Flores-López A, González-Salazar LE, Arteaga-Sanchez L, Medina-Vera I, Orozco-Ruiz X, Torres N, et al. Serum amino acid concentrations are modified by age, insulin resistance, and BCAT2 rs11548193 and BCKDH rs45500792 polymorphisms in subjects with obesity. Clin Nutr. 2021;40(6):4209–4215. doi:10.1016/j.clnu.2021.01.037

1079. Accattatis FM, Caruso A, Carleo A, Del Console P, Gelsomino L, Bonofiglio D, Giordano C, Barone I, Andò S, Bianchi L, et al. CEBP-β and PLK1 as Potential Mediators of the Breast Cancer/Obesity Crosstalk: In Vitro and In Silico Analyses. Nutrients. 2023;15(13):2839. doi:10.3390/nu15132839

1080. Yang Y, Liu H, Liu D. Preventing high-fat diet-induced obesity and related metabolic disorders by hydrodynamic transfer of Il-27 gene. Int J Obes (Lond). 2023;47(5):413–421. doi:10.1038/s41366-023-01293-6

1081. Shih DM, Meng Y, Sallam T, Vergnes L, Shu ML, Reue K, Tontonoz P, Fogelman AM, Lusis AJ, Reddy ST, et al. PON2 Deficiency Leads to Increased Susceptibility to Diet-Induced Obesity. Antioxidants (Basel). 2019;8(1):19. doi:10.3390/antiox8010019

1082. Simmen FA, Pabona JMP, Al-Dwairi A, Alhallak I, Montales MTE, Simmen RCM. Malic Enzyme 1 (ME1) Promotes Adiposity and Hepatic Steatosis and Induces Circulating Insulin and Leptin in Obese Female Mice. Int J Mol Sci. 2023;24(7):6613. doi:10.3390/ijms24076613

1083. Vassileva G, Hu W, Hoos L, Tetzloff G, Yang S, Liu L, Kang L, Davis HR, Hedrick JA, Lan H, et al. Gender-dependent effect of Gpbar1 genetic deletion on the metabolic profiles of diet-induced obese mice. J Endocrinol. 2010;205(3):225–232. doi:10.1677/JOE-10-0009

1084. Piek A, Koonen DPY, Schouten EM, Lindtstedt EL, Michaëlsson E, de Boer RA, Silljé HHW. Pharmacological myeloperoxidase (MPO) inhibition in an obese/hypertensive mouse model attenuates obesity and liver damage, but not cardiac remodeling. Sci Rep. 2019;9(1):18765. doi:10.1038/s41598-019-55263-y

1085. Manco L, Machado-Rodrigues AM, Padez C. Association study of common functional genetic polymorphisms in SLC6A4 (5-HTT) and MAOA genes with obesity in portuguese children. Arch Physiol Biochem. 2022;128(6):1510–1515. doi:10.1080/13813455.2020.1779312

1086. Wang H, Astarita G, Taussig MD, Bharadwaj KG, DiPatrizio NV, Nave KA, Piomelli D, Goldberg IJ, Eckel RH. Deficiency of lipoprotein lipase in neurons modifies the regulation of energy balance and leads to obesity. Cell Metab. 2011;13(1):105–113. doi:10.1016/j.cmet.2010.12.006

1087. Katsouda A, Valakos D, Dionellis VS, Bibli SI, Akoumianakis I, Karaliota S, Zuhra K, Fleming I, Nagahara N, Havaki S, et al. MPST sulfurtransferase maintains mitochondrial protein import and cellular bioenergetics to attenuate obesity. J Exp Med. 2022;219(7):e20211894. doi:10.1084/jem.20211894

1088. Liu T, David SP, Tyndale RF, Wang H, Yu XQ, Chen W, Zhou Q, Chen WQ. Relationship between amounts of daily cigarette consumption and abdominal obesity moderated by CYP2A6 genotypes in Chinese male current smokers. Ann Behav Med. 2012;43(2):253–261. doi:10.1007/s12160-011-9318-5

1089. Pisano E, Pacifico L, Perla FM, Liuzzo G, Chiesa C, Lavorato M, Mingrone G, Fabrizi M, Fintini D, Severino A, et al. Upregulated monocyte expression of PLIN2 is associated with early arterial injury in children with overweight/obesity. Atherosclerosis. 2021;327:68–75. doi:10.1016/j.atherosclerosis.2021.04.016

1090. Gholami M, Zoughi M, Behboo R, Taslimi R, Kazemeini A, Bastami M, Hasani-Ranjbar S, Larijani B, Amoli MM. Association of miRNA targetome variants in LAMC1 and GNB3 genes with colorectal cancer and obesity. Cancer Med. 2022;11(21):3923–3938. doi:10.1002/cam4.4713

1091. Chinnasamy P, Casimiro I, Riascos-Bernal DF, Venkatesh S, Parikh D, Maira A, Srinivasan A, Zheng W, Tarabra E, Zong H, et al. Increased adipose catecholamine levels and protection from obesity with loss of Allograft Inflammatory Factor-1. Nat Commun. 2023;14(1):38. doi:10.1038/s41467-022-35683-7

1092. Ding L, Goossens GH, Oligschlaeger Y, Houben T, Blaak EE, Shiri-Sverdlov R. Plasma cathepsin D activity is negatively associated with hepatic insulin sensitivity in overweight and obese humans. Diabetologia. 2020;63(2):374–384. doi:10.1007/s00125-019-05025-2

1093. Hosooka T, Noguchi T, Kotani K, Nakamura T, Sakaue H, Inoue H, Ogawa W, Tobimatsu K, Takazawa K, Sakai M, et al. Dok1 mediates high-fat diet-induced adipocyte hypertrophy and obesity through modulation of PPAR-gamma phosphorylation. Nat Med. 2008;14(2):188–193. doi:10.1038/nm1706

1094. Backe MB, Andersen RC, Jensen M, Jin C, Hundahl C, Dmytriyeva O, Treebak JT, Hansen JB, Gerhart-Hines Z, Madsen KL, et al. PICK1-Deficient Mice Maintain Their Glucose Tolerance During Diet-Induced Obesity. J Endocr Soc. 2023;7(6):bvad057. doi:10.1210/jendso/bvad057

1095. Hao J, Liu Z, Ju W, He F, Liu K, Wu J. Role and mechanism of FLT4 in high-fat diet-induced obesity in mice. Biochem Biophys Res Commun. 2023;675:61–70. doi:10.1016/j.bbrc.2023.06.025

1096. Deveci Sevim R, Gök M, Çevik Ö, Erdoğan Ö, Güneş S, Ünüvar T, Anık A. Associations of Adipocyte-Derived Versican and Macrophage-Derived Biglycan with Body Adipose Tissue and Hepatosteatosis in Obese Children. J Clin Res Pediatr Endocrinol. 2024. doi:10.4274/jcrpe.galenos.2024.2023-9-18

1097. Maguolo A, Zusi C, Giontella A, Miraglia Del Giudice E, Tagetti A, Fava C, Morandi A, Maffeis C. Influence of genetic variants in FADS2 and ELOVL2 genes on BMI and PUFAs homeostasis in children and adolescents with obesity. Int J Obes (Lond). 2021;45(1):56–65. doi:10.1038/s41366-020-00662-9

1098. Tian N, Liu Q, Li Y, Tong L, Lu Y, Zhu Y, Zhang P, Chen H, Hu L, Meng J, et al. Transketolase Deficiency in Adipose Tissues Protects Mice From Diet-Induced Obesity by Promoting Lipolysis. Diabetes. 2020;69(7):1355–1367. doi:10.2337/db19-1087

1099. Peters WR, MacMurry JP, Walker J, Giese RJ Jr, Comings DE. Phenylethanolamine N-methyltransferase G-148A genetic variant and weight loss in obese women. Obes Res. 2003;11(3):415–419. doi:10.1038/oby.2003.56

1100. Hamano M, Esaki K, Moriyasu K, Yasuda T, Mohri S, Tashiro K, Hirabayashi Y, Furuya S. Hepatocyte-Specific Phgdh-Deficient Mice Culminate in Mild Obesity, Insulin Resistance, and Enhanced Vulnerability to Protein Starvation. Nutrients. 2021;13(10):3468. doi:10.3390/nu13103468

1101. Chirita-Emandi A, Serban CL, Paul C, Andreescu N, Velea I, Mihailescu A, Serafim V, Tiugan DA, Tutac P, Zimbru C, et al. CHDH-PNPLA3 Gene-Gene Interactions Predict Insulin Resistance in Children with Obesity. Diabetes Metab Syndr Obes. 2020;13:4483–4494. doi:10.2147/DMSO.S277268

1102. Sie MP, Sayed-Tabatabaei FA, Oei HH, Uitterlinden AG, Pols HA, Hofman A, van Duijn CM, Witteman JC. Interleukin 6 -174 g/c promoter polymorphism and risk of coronary heart disease: results from the rotterdam study and a meta-analysis. Arterioscler Thromb Vasc Biol. 2006;26(1):212–217. doi:10.1161/01.ATV.0000194099.65024.17

1103. Guo LY, Yang F, Peng LJ, Li YB, Wang AP. CXCL2, a new critical factor and therapeutic target for cardiovascular diseases. Clin Exp Hypertens. 2020;42(5):428–437. doi:10.1080/10641963.2019.1693585

1104. Zhang L, Wang YN, Ju JM, Shabanova A, Li Y, Fang RN, Sun JB, Guo YY, Jin TZ, Liu YY, et al. Mzb1 protects against myocardial infarction injury in mice via modulating mitochondrial function and alleviating inflammation. Acta Pharmacol Sin. 2021;42(5):691–700. doi:10.1038/s41401-020-0489-0

1105. Korbecki J, Maruszewska A, Bosiacki M, Chlubek D, Baranowska-Bosiacka I. The Potential Importance of CXCL1 in the Physiological State and in Noncancer Diseases of the Cardiovascular System, Respiratory System and Skin. Int J Mol Sci. 2022;24(1):205. doi:10.3390/ijms24010205

1106. Gaio P, Gualdrón-López M, Cramer A, Esper L, de Menezes Filho JER, Cruz JS, Teixeira MM, Machado FS. SOCS2 expression in hematopoietic and non-hematopoietic cells during Trypanosoma cruzi infection: Correlation with immune response and cardiac dysfunction. Clin Immunol. 2022;234:108913. doi:10.1016/j.clim.2021.108913

1107. Varbo A, Benn M, Tybjærg-Hansen A, Grande P, Nordestgaard BG. TRIB1 and GCKR polymorphisms, lipid levels, and risk of ischemic heart disease in the general population. Arterioscler Thromb Vasc Biol. 2011;31(2):451–457. doi:10.1161/ATVBAHA.110.216333

1108. Pedroso JAB, Silva IBD, Zampieri TT, Totola LT, Moreira TS, Taniguti APT, Diniz GP, Barreto-Chaves MLM, Donato J Jr. SOCS3 Ablation in Leptin Receptor-Expressing Cells Causes Autonomic and Cardiac Dysfunctions in Middle-Aged Mice despite Improving Energy and Glucose Metabolism. Int J Mol Sci. 2022;23(12):6484. doi:10.3390/ijms23126484

1109. Lin CS, Hsieh PS, Hwang LL, Lee YH, Tsai SH, Tu YC, Hung YW, Liu CC, Chuang YP, et al. The CCL5/CCR5 Axis Promotes Vascular Smooth Muscle Cell Proliferation and Atherogenic Phenotype Switching. Cell Physiol Biochem. 2018;47(2):707–720. doi:10.1159/000490024

1110. Singh V, Kaur R, Kumari P, Pasricha C, Singh R. ICAM-1 and VCAM-1: Gatekeepers in various inflammatory and cardiovascular disorders. Clin Chim Acta. 2023;548:117487. doi:10.1016/j.cca.2023.117487

1111. Coto E, Reguero JR, Avanzas P, Pascual I, Martín M, Hevia S, Morís C, Díaz-Molina B, Lambert JL, Alonso B, et al. Gene variants in the NF-KB pathway (NFKB1, NFKBIA, NFKBIZ) and risk for early-onset coronary artery disease. Immunol Lett. 2019;208:39–43. doi:10.1016/j.imlet.2019.02.007

1112. Crea F. New therapeutic targets to reduce inflammation-associated cardiovascular risk: the CCL2-CCR2 axis, LOX-1, and IRF5. Eur Heart J. 2022;43(19):1777–1781. doi:10.1093/eurheartj/ehac233

1113. Zhou Q, Hahn JK, Neupane B, Aidery P, Labeit S, Gawaz M, Gramlich M. Dysregulated IER3 Expression is Associated with Enhanced Apoptosis in Titin-Based Dilated Cardiomyopathy. Int J Mol Sci. 2017;18(4):723. doi:10.3390/ijms18040723

1114. Kardys I, van Tiel CM, de Vries CJ, Pannekoek H, Uitterlinden AG, Hofman A, Witteman JC, de Maat MP. Haplotypes of the NR4A2/NURR1 gene and cardiovascular disease: the Rotterdam Study. Hum Mutat. 2009;30(3):417–423. doi:10.1002/humu.20902

1115. Chen X, Li X, Wu X, Ding Y, Li Y, Zhou G, Wei Y, Chen S, Lu X, Xu J, et al. Integrin beta-like 1 mediates fibroblast-cardiomyocyte crosstalk to promote cardiac fibrosis and hypertrophy. Cardiovasc Res. 2023;119(10):1928–1941. doi:10.1093/cvr/cvad104

1116. Zhang S, Ding Y, Feng F, Gao Y. The role of blood CXCL12 level in prognosis of coronary artery disease: A meta-analysis. Front Cardiovasc Med. 2022;9:938540. doi:10.3389/fcvm.2022.938540

1117. Williams T, Hundertmark M, Nordbeck P, Voll S, Arias-Loza PA, Oppelt D, Mühlfelder M, Schraut S, Elsner I, Czolbe M, et al. Eya4 Induces Hypertrophy via Regulation of p27kip1. Circ Cardiovasc Genet. 2015;8(6):752–764. doi:10.1161/CIRCGENETICS.115.001134

1118. Hilfiker-Kleiner D, Hilfiker A, Castellazzi M, Wollert KC, Trautwein C, Schunkert H, Drexler H. JunD attenuates phenylephrine-mediated cardiomyocyte hypertrophy by negatively regulating AP-1 transcriptional activity. Cardiovasc Res. 2006;71(1):108–117. doi:10.1016/j.cardiores.2006.02.032

1119. Stegger JG, Schmidt EB, Tjønneland A, Kopp TI, Sørensen TI, Vogel U, Overvad K. Single nucleotide polymorphisms in IL1B and the risk of acute coronary syndrome: a Danish case-cohort study. PLoS One. 2012;7(6):e36829. doi:10.1371/journal.pone.0036829

1120. Ji M, Liu Y, Zuo Z, Xu C, Lin L, Li Y. Downregulation of amphiregulin improves cardiac hypertrophy via attenuating oxidative stress and apoptosis. Biol Direct. 2022;17(1):21. doi:10.1186/s13062-022-00334-w

1121. Rui H, Zhao F, Yuhua L, Hong J. Suppression of SMOC2 alleviates myocardial fibrosis via the ILK/p38 pathway. Front Cardiovasc Med. 2023;9:951704. doi:10.3389/fcvm.2022.951704

1122. Wang HB, Yang J, Shuai W, Yang J, Liu LB, Xu M, Tang QZ. Deletion of Microfibrillar-Associated Protein 4 Attenuates Left Ventricular Remodeling and Dysfunction in Heart Failure. J Am Heart Assoc. 2020;9(17):e015307. doi:10.1161/JAHA.119.015307

1123. Er LK, Hsu LA, Juang JJ, Chiang FT, Teng MS, Tzeng IS, Wu S, Lin JF, Ko YL. Circulating Chemerin Levels, but not the RARRES2 Polymorphisms, Predict the Long-Term Outcome of Angiographically Confirmed Coronary Artery Disease. Int J Mol Sci. 2019;20(5):1174. doi:10.3390/ijms20051174

1124. Liu ZY, Liu F, Cao Y, Peng SL, Pan HW, Hong XQ, Zheng PF. ACSL1, CH25H, GPCPD1, and PLA2G12A as the potential lipid-related diagnostic biomarkers of acute myocardial infarction. Aging (Albany NY). 2023;15(5):1394–1411. doi:10.18632/aging.204542

1125. Poloni G, Calore M, Rigato I, Marras E, Minervini G, Mazzotti E, Lorenzon A, Li Mura IEA, Telatin A, Zara I, et al. A targeted next-generation gene panel reveals a novel heterozygous nonsense variant in the TP63 gene in patients with arrhythmogenic cardiomyopathy. Heart Rhythm. 2019;16(5):773–780. doi:10.1016/j.hrthm.2018.11.015

1126. Toyohara T, Roudnicky F, Florido MHC, Nakano T, Yu H, Katsuki S, Lee M, Meissner T, Friesen M, Davidow LS, et al. Patient hiPSCs Identify Vascular Smooth Muscle Arylacetamide Deacetylase as Protective against Atherosclerosis. Cell Stem Cell. 2020;27(1):147–157.e7. doi:10.1016/j.stem.2020.04.018

1127. Crasto S, My I, Di Pasquale E. The Broad Spectrum of LMNA Cardiac Diseases: From Molecular Mechanisms to Clinical Phenotype. Front Physiol. 2020;11:761. doi:10.3389/fphys.2020.00761

1128. Zhao L, Zhang Q, Liang J, Li J, Tan X, Tang N. Astrocyte elevated gene-1 induces autophagy in diabetic cardiomyopathy through upregulation of KLF4. J Cell Biochem. 2019;120(6):9709–9715. doi:10.1002/jcb.28249

1129. Crossman DC, Morton AC, Gunn JP, Greenwood JP, Hall AS, Fox KA, Lucking AJ, Flather MD, Lees B, Foley CE. Investigation of the effect of Interleukin-1 receptor antagonist (IL-1ra) on markers of inflammation in non-ST elevation acute coronary syndromes (The MRC-ILA-HEART Study). Trials. 2008;9:8. doi:10.1186/1745-6215-9-8

1130. Meng XW, Zhang M, Hu JK, Chen XY, Long YQ, Liu H, Feng XM, Ji FH, Peng K. Activation of CCL21-GPR174/CCR7 on cardiac fibroblasts underlies myocardial ischemia/reperfusion injury. Front Genet. 2022;13:946524. doi:10.3389/fgene.2022.946524

1131. Nagano H, Mitchell RN, Taylor MK, Hasegawa S, Tilney NL, Libby P. Interferon-gamma deficiency prevents coronary arteriosclerosis but not myocardial rejection in transplanted mouse hearts. J Clin Invest. 1997;100(3):550–557. doi:10.1172/JCI119564

1132. Hojayev B, Rothermel BA, Gillette TG, Hill JA. FHL2 binds calcineurin and represses pathological cardiac growth. Mol Cell Biol. 2012;32(19):4025–4034. doi:10.1128/MCB.05948-11

1133. Funk SD, Yurdagul A Jr, Albert P, Traylor JG Jr, Jin L, Chen J, Orr AW. EphA2 activation promotes the endothelial cell inflammatory response: a potential role in atherosclerosis. Arterioscler Thromb Vasc Biol. 2012;32(3):686–695. doi:10.1161/ATVBAHA.111.242792

1134. Li B, Xie X. A20 (TNFAIP3) alleviates viral myocarditis through ADAR1/miR-1a-3p-dependent regulation. BMC Cardiovasc Disord. 2022;22(1):10. doi:10.1186/s12872-021-02438-z

1135. Vrtovec B, Poglajen G, Sever M, Lezaic L, Socan A, Haddad F, Wu JC. CD34+ stem cell therapy in nonischemic dilated cardiomyopathy patients. Clin Pharmacol Ther. 2013;94(4):452–458. doi:10.1038/clpt.2013.134

1136. Zhang Z, Wang L, Zhan Y, Xie C, Xiang Y, Chen D, Wu Y. Clinical value and expression of Homer 1, homocysteine, S-adenosyl-l-homocysteine, fibroblast growth factors 23 in coronary heart disease. BMC Cardiovasc Disord. 2022;22(1):215. doi:10.1186/s12872-022-02554-4

1137. Zhao J, Jiang X, Liu J, Ye P, Jiang L, Chen M, Xia J. Dual-Specificity Phosphatase 26 Protects Against Cardiac Hypertrophy Through TAK1. J Am Heart Assoc. 2021;10(4):e014311. doi:10.1161/JAHA.119.014311

1138. Thanikachalam PV, Ramamurthy S, Mallapu P, Varma SR, Narayanan J, Abourehab MA, Kesharwani P. Modulation of IL-33/ST2 signaling as a potential new therapeutic target for cardiovascular diseases. Cytokine Growth Factor Rev. 2023;71-72:94–104. doi:10.1016/j.cytogfr.2023.06.003

1139. Zhang K, Wu M, Qin X, Wen P, Wu Y, Zhuang J. Asporin is a Potential Promising Biomarker for Common Heart Failure. DNA Cell Biol. 2021;40(2):303–315. doi:10.1089/dna.2020.5995

1140. Setsuta K, Seino Y, Ogawa T, Ohtsuka T, Seimiya K, Takano T. Ongoing myocardial damage in chronic heart failure is related to activated tumor necrosis factor and Fas/Fas ligand system. Circ J. 2004;68(8):747–750. doi:10.1253/circj.68.747

1141. Jia EZ, Wang J, Yang ZJ, Zhu TB, Wang LS, Chen B, Cao KJ, Huang J, Ma WZ. Molecular scanning of the human carboxypeptidase E gene for mutations in Chinese subjects with coronary atherosclerosis. Mol Cell Biochem. 2008;307(1-2):31–39. doi:10.1007/s11010-007-9581-8

1142. Sinagra E, Perricone G, Romano C, Cottone M. Heart failure and anti tumor necrosis factor-alpha in systemic chronic inflammatory diseases. Eur J Intern Med. 2013;24(5):385–392. doi:10.1016/j.ejim.2012.12.015

1143. Xiao QA, He Q, Zeng J, Xia X. GDF-15, a future therapeutic target of glucolipid metabolic disorders and cardiovascular disease. Biomed Pharmacother. 2022;146:112582. doi:10.1016/j.biopha.2021.112582

1144. Noori F, Naeimi S, Zibaeenezhad MJ, Gharemirshamlu FR. CCL22 and CCR4 Gene Polymorphisms in Myocardial Infarction: Risk Assessment of rs4359426 and rs2228428 in Iranian Population. Clin Lab. 2018;64(6):907–913. doi:10.7754/Clin.Lab.2018.171106

1145. Xiao J, Yu K, Li M, Xiong C, Wei Y, Zeng Q. The IL-2/Anti-IL-2 Complex Attenuates Cardiac Ischaemia-Reperfusion Injury Through Expansion of Regulatory T Cells. Cell Physiol Biochem. 2017;44(5):1810–1827. doi:10.1159/000485818

1146. Lopez-Canoa JN, Baluja A, Couselo-Seijas M, Naveira AB, Gonzalez-Melchor L, Rozados A, Martínez-Sande L, García-Seara J, Fernandez-Lopez XA, Fernandez AL, et al. Plasma FABP4 levels are associated with left atrial fat volume in persistent atrial fibrillation and predict recurrence after catheter ablation. Int J Cardiol. 2019;292:131–135. doi:10.1016/j.ijcard.2019.04.031

1147. Sun Y, Lu Q, Tao X, Cheng B, Yang G. Cyp2C19*2 Polymorphism Related to Clopidogrel Resistance in Patients With Coronary Heart Disease, Especially in the Asian Population: A Systematic Review and Meta-Analysis. Front Genet. 2020;11:576046. doi:10.3389/fgene.2020.576046

1148. Acin-Perez R, Lechuga-Vieco AV, Del Mar Muñoz M, Nieto-Arellano R, Torroja C, Sánchez-Cabo F, Jiménez C, González-Guerra A, Carrascoso I, Benincá C, et al. Ablation of the stress protease OMA1 protects against heart failure in mice. Sci Transl Med. 2018;10(434):eaan4935. doi:10.1126/scitranslmed.aan4935

1149. Chen YL, Chen YC, Chang YT, Wang HT, Liu WH, Chong SZ, Lin PT, Hsu PY, Su MC, Lin MC. GJA1 Expression and Left Atrial Remodeling in the Incidence of Atrial Fibrillation in Patients with Obstructive Sleep Apnea Syndrome. Biomedicines. 2021;9(10):1463. doi:10.3390/biomedicines9101463

1150. Khalil A, Al-Haddad C, Hariri H, Shibbani K, Bitar F, Kurban M, Nemer G, Arabi M. A Novel Mutation in FOXC1 in a Lebanese Family with Congenital Heart Disease and Anterior Segment Dysgenesis: Potential Roles for NFATC1 and DPT in the Phenotypic Variations. Front Cardiovasc Med. 2017;4:58. doi:10.3389/fcvm.2017.00058

1151. Sterner-Kock A, Thorey IS, Koli K, Wempe F, Otte J, Bangsow T, Kuhlmeier K, Kirchner T, Jin S, Keski-Oja J, et al. Disruption of the gene encoding the latent transforming growth factor-beta binding protein 4 (LTBP-4) causes abnormal lung development, cardiomyopathy, and colorectal cancer. Genes Dev. 2002;16(17):2264–2273. doi:10.1101/gad.229102

1152. Gao J, Wang S, Liu S. The involvement of protein TNFSF18 in promoting p-STAT1 phosphorylation to induce coronary microcirculation disturbance in atherosclerotic mouse model. Drug Dev Res. 2021;82(1):115–122. doi:10.1002/ddr.21735

1153. Fu F, Doroudgar S. IRE1/XBP1 and endoplasmic reticulum signaling -from basic to translational research for cardiovascular disease. Curr Opin Physiol. 2022;28:100552. doi:10.1016/j.cophys.2022.100552

1154. Arvanitis M, Tampakakis E, Zhang Y, Wang W, Auton A; 23andMe Research Team; Dutta D, Glavaris S, Keramati A, Chatterjee N, Chi NC, et al. Genome-wide association and multi-omic analyses reveal ACTN2 as a gene linked to heart failure. Nat Commun. 2020;11(1):1122. doi:10.1038/s41467-020-14843-7

1155. Mozaffari MS. Therapeutic Potential of Annexin A1 Modulation in Kidney and Cardiovascular Disorders. Cells. 2021;10(12):3420. doi:10.3390/cells10123420

1156. Li Y, Yan H, Guo J, Han Y, Zhang C, Liu X, Du J, Tian XL. Down-regulated RGS5 by genetic variants impairs endothelial cell function and contributes to coronary artery disease. Cardiovasc Res. 2021;117(1):240–255. doi:10.1093/cvr/cvz268

1157. Zhou X, Zhang C, Wu X, Hu X, Zhang Y, Wang X, Zheng L, Gao P, Du J, Zheng W, et al. Dusp6 deficiency attenuates neutrophil-mediated cardiac damage in the acute inflammatory phase of myocardial infarction. Nat Commun. 2022;13(1):6672. doi:10.1038/s41467-022-33631-z

1158. Mahendra J, Srinivasan S, Kanakamedala A, D N, Namasivayam A, Mahendra L, Muralidharan J, Cherian SM, Ilango P. Expression of trefoil factor 2 and 3 and adrenomedullin in chronic periodontitis subjects with coronary heart disease. J Periodontol. 2023;94(5):694–703. doi:10.1002/JPER.22-0467

1159. Chan BT, Lim E, Chee KH, Abu Osman NA. Review on CFD simulation in heart with dilated cardiomyopathy and myocardial infarction. Comput Biol Med. 2013;43(4):377–385. doi:10.1016/j.compbiomed.2013.01.013

1160. Khosravi E, Sadeghian L, Mohamadynejad P, Dianatkhah M, Hajizadeh M, Gharipour M. Association study of polymorphism in Thrombomodulin gene [rs1042579] with cardiovascular disease. Acta Biomed. 2022;92(6):e2021282. doi:10.23750/abm.v92i6.9622

1161. García-Quintáns N, Sacristán S, Márquez-López C, Sánchez-Ramos C, Martinez-de-Benito F, Siniscalco D, González-Guerra A, Camafeita E, Roche-Molina M, Lytvyn M, et al. MYH10 activation rescues contractile defects in arrhythmogenic cardiomyopathy (ACM). Nat Commun. 2023;14(1):6461. doi:10.1038/s41467-023-41981-5

1162. Salehipour P, Rezagholizadeh F, Mahdiannasser M, Kazerani R, Modarressi MH. Association of OLR1 gene polymorphisms with the risk of coronary artery disease: A systematic review and meta-analysis. Heart Lung. 2021;50(2):334–343. doi:10.1016/j.hrtlng.2021.01.015

1163. Bern MM, Cassani MP, Horton J, Rand L, Davis G. Changes of fibrinolysis and factor VIII coagulant, antigen, and ristocetin cofactor in diabetes mellitus and atherosclerosis. Thromb Res. 1980;19(6):831–839. doi:10.1016/0049-3848(80)90011-0

1164. Grabowski K, Herlan L, Witten A, Qadri F, Eisenreich A, Lindner D, Schädlich M, Schulz A, Subrova J, Mhatre KN, et al. Cpxm2 as a novel candidate for cardiac hypertrophy and failure in hypertension. Hypertens Res. 2022;45(2):292–307. doi:10.1038/s41440-021-00826-8

1165. Zheng PF, Yin RX, Cao XL, Chen WX, Wu JZ, Huang F. Effect of SYTL3-SLC22A3 Variants, Their Haplotypes, and G × E Interactions on Serum Lipid Levels and the Risk of Coronary Artery Disease and Ischaemic Stroke. Front Cardiovasc Med. 2021;8:713068. doi:10.3389/fcvm.2021.713068

1166. Xia Z, Gu M, Jia X, Wang X, Wu C, Guo J, Zhang L, Du Y, Wang J. Integrated DNA methylation and gene expression analysis identifies SLAMF7 as a key regulator of atherosclerosis. Aging (Albany NY). 2018;10(6):1324–1337. doi:10.18632/aging.101470

1167. Maneerat Y, Prasongsukarn K, Benjathummarak S, Dechkhajorn W. PPBP and DEFA1/DEFA3 genes in hyperlipidaemia as feasible synergistic inflammatory biomarkers for coronary heart disease. Lipids Health Dis. 2017;16(1):80. doi:10.1186/s12944-017-0471-0

1168. Xie J, Liao W, Chen W, Lai D, Tang Q, Li Y. Circulating long non-coding RNA TTTY15 and HULC serve as potential novel biomarkers for predicting acute myocardial infarction. BMC Cardiovasc Disord. 2022;22(1):86. doi:10.1186/s12872-022-02529-5

1169. Chen Q, Li Z, Wang M, Li G. Over-expression of IL1R2 in PBMCs of Patients with Coronary Artery Disease and Its Clinical Significance. Anatol J Cardiol. 2022;26(9):710–716. doi:10.5152/AnatolJCardiol.2022.1241

1170. Timur AA, Murugesan G, Zhang L, Aung PP, Barnard J, Wang QK, Gaussem P, Silverstein RL, Bhatt DL, Kottke-Marchant K. P2RY1 and P2RY12 polymorphisms and on-aspirin platelet reactivity in patients with coronary artery disease. Int J Lab Hematol. 2012;34(5):473–483. doi:10.1111/j.1751-553X.2012.01420.x

1171. Delgobo M, Frantz S. When Sensing Goes Wrong: Role of Clec4e in Ischemic Heart Injury. JACC Basic Transl Sci. 2021;6(8):647–649. doi:10.1016/j.jacbts.2021.07.003

1172. Corbin LJ, White SJ, Taylor AE, Williams CM, Taylor K, van den Bosch MT, Teasdale JE, Jones M, Bond M, Harper MT, et al. Epigenetic Regulation of F2RL3 Associates With Myocardial Infarction and Platelet Function. Circ Res. 2022;130(3):384–400. doi:10.1161/CIRCRESAHA.121.318836

1173. Zhang R, Ji Z, Qu Y, Yang M, Su Y, Zuo W, Zhao Q, Ma G, Li Y. Clinical value of ARG1 in acute myocardial infarction patients: Bioinformatics-based approach. Biomed Pharmacother. 2020;121:109590. doi:10.1016/j.biopha.2019.109590

1174. Ogura Y, Tajiri K, Murakoshi N, Xu D, Yonebayashi S, Li S, Okabe Y, Feng D, Shimoda Y, Song Z, et al. Neutrophil Elastase Deficiency Ameliorates Myocardial Injury Post Myocardial Infarction in Mice. Int J Mol Sci. 2021;22(2):722. doi:10.3390/ijms22020722

1175. Hecker PA, Leopold JA, Gupte SA, Recchia FA, Stanley WC. Impact of glucose-6-phosphate dehydrogenase deficiency on the pathophysiology of cardiovascular disease. Am J Physiol Heart Circ Physiol. 2013;304(4):H491–H500. doi:10.1152/ajpheart.00721.2012

1176. Xie Z, Shen Y, Huang S, Shen W, Liu J. Abnormal ADAMTS2 and VSIG4 in Serum of HF Patients and their Relationship with CRP, UA, and HCY. Clin Lab. 2022;68(5):10.7754/Clin.Lab.2021.210811. doi:10.7754/Clin.Lab.2021.210811

1177. Carreras-Torres R, Athanasiadis G, Via M, Trenchs J, Gayà-Vidal M, Santamaria J, Esteban E, Moral P. Allele-allele interaction within the F13A1 gene: a risk factor for ischaemic heart disease in Spanish population. Thromb Res. 2010;126(3):e241–e245. doi:10.1016/j.thromres.2010.04.021

1178. Yu Y, Song G. Lipopolysaccharide-Binding Protein and Bactericidal/Permeability-Increasing Protein in Lipid Metabolism and Cardiovascular Diseases. Adv Exp Med Biol. 2020;1276:27–35. doi:10.1007/978-981-15-6082-8_3

1179. Wei M, Pan H, Guo K. Association Between Plasma ADAMTS-9 Levels and Severity of Coronary Artery Disease. Angiology. 2021;72(4):371–380. doi:10.1177/0003319720979238

1180. Wang C, Wong J, Fung G, Shi J, Deng H, Zhang J, Bernatchez P, Luo H. Dysferlin deficiency confers increased susceptibility to coxsackievirus-induced cardiomyopathy. Cell Microbiol. 2015;17(10):1423–1430. doi:10.1111/cmi.12473

1181. Mehra S, Kumar M, Manchanda M, Singh R, Thakur B, Rani N, Arava S, Narang R, Arya DS, Chauhan SS. Clinical significance of cathepsin L and cathepsin B in dilated cardiomyopathy. Mol Cell Biochem. 2017;428(1-2):139–147. doi:10.1007/s11010-016-2924-6

1182. Zhao Z, Liu G, Zhang H, Ruan P, Ge J, Liu Q. BIRC5, GAJ5, and lncRNA NPHP3-AS1 Are Correlated with the Development of Atrial Fibrillation-Valvular Heart Disease. Int Heart J. 2021;62(1):153–161. doi:10.1536/ihj.20-238

1183. Dervisoglu P, Elmas B. Pentraxin 3 as a Marker for Cardiovascular Disease Risk in Overweight and Obese Children. Acta Cardiol Sin. 2021;37(2):177–183. doi:10.6515/ACS.202103_37(2).20201006A

1184. Damp J, Givertz MM, Semigran M, Alharethi R, Ewald G, Felker GM, Bozkurt B, Boehmer J, Haythe J, Skopicki H, et al. Relaxin-2 and Soluble Flt1 Levels in Peripartum Cardiomyopathy: Results of the Multicenter IPAC Study. JACC Heart Fail. 2016;4(5):380–388. doi:10.1016/j.jchf.2016.01.004

1185. Fujimaki T, Kato K, Yokoi K, Oguri M, Yoshida T, Watanabe S, Metoki N, Yoshida H, Satoh K, Aoyagi Y, et al. Association of genetic variants in SEMA3F, CLEC16A, LAMA3, and PCSK2 with myocardial infarction in Japanese individuals. Atherosclerosis. 2010;210(2):468–473. doi:10.1016/j.atherosclerosis.2009.11.050

1186. Inagaki K, Koyanagi T, Berry NC, Sun L, Mochly-Rosen D. Pharmacological inhibition of epsilon-protein kinase C attenuates cardiac fibrosis and dysfunction in hypertension-induced heart failure. Hypertension. 2008;51(6):1565–1569. doi:10.1161/HYPERTENSIONAHA.107.109637

1187. Olejnik A, Franczak A, Krzywonos-Zawadzka A, Kałużna-Oleksy M, Bil-Lula I. The Biological Role of Klotho Protein in the Development of Cardiovascular Diseases. Biomed Res Int. 2018;2018:5171945. doi:10.1155/2018/5171945

1188. Huang H, Zeng Z, Li J, Zhang L, Chen Y. Variants of arachidonate 5-lipoxygenase-activating protein (ALOX5AP) gene and risk of coronary heart disease: A meta-analysis. Arch Med Res. 2010;41(8):634–641. doi:10.1016/j.arcmed.2010.11.001

1189. Alaei Z, Zamani N, Rabbani B, Mahdieh N. TCAP gene is not a common cause of cardiomyopathy in Iranian patients. Eur J Med Res. 2023;28(1):376. doi:10.1186/s40001-023-01019-4

1190. Fu F, Lai Q, Hu J, Zhang L, Zhu X, Kou J, Yu B, Li F.Ruscogenin Alleviates Myocardial Ischemia-Induced Ferroptosis through the Activation of BCAT1/BCAT2. Antioxidants (Basel). 2022;11(3):583. doi:10.3390/antiox11030583

1191. Heo SC, Kwon YW, Jang IH, Jeong GO, Lee TW, Yoon JW, Shin HJ, Jeong HC, Ahn Y, Ko TH, et al Formyl Peptide Receptor 2 Is Involved in Cardiac Repair After Myocardial Infarction Through Mobilization of Circulating Angiogenic Cells. Stem Cells. 2017;35(3):654–665. doi:10.1002/stem.253

1192. Jafarizade M, Kahe F, Sharfaei S, Momenzadeh K, Pitliya A, Zahedi Tajrishi F, Singh P, Chi G. The Role of Interleukin-27 in Atherosclerosis: A Contemporary Review. Cardiology. 2021;146(4):517–530. doi:10.1159/000515359

1193. Devarajan A, Bourquard N, Hama S, Navab M, Grijalva VR, Morvardi S, Clarke CF, Vergnes L, Reue K, Teiber JF, et al. Paraoxonase 2 deficiency alters mitochondrial function and exacerbates the development of atherosclerosis. Antioxid Redox Signal. 2011;14(3):341–351. doi:10.1089/ars.2010.3430

1194. Haarhaus M, Cianciolo G, Barbuto S, La Manna G, Gasperoni L, Tripepi G, Plebani M, Fusaro M, Magnusson P. Alkaline Phosphatase: An Old Friend as Treatment Target for Cardiovascular and Mineral Bone Disorders in Chronic Kidney Disease. Nutrients. 2022;14(10):2124. doi:10.3390/nu14102124

1195. Janus SE, Hajjari J, Chami T, Karnib M, Al-Kindi SG, Rashid I. Myeloperoxidase is Independently Associated with Incident Heart Failure in Patients with Coronary Artery Disease and Kidney Disease. Curr Probl Cardiol. 2022;47(11):101080. doi:10.1016/j.cpcardiol.2021.101080

1196. Machado-Vieira R, Mallinger AG. Abnormal function of monoamine oxidase-A in comorbid major depressive disorder and cardiovascular disease: pathophysiological and therapeutic implications (review). Mol Med Rep. 2012;6(5):915–922. doi:10.3892/mmr.2012.1062

1197. Geldenhuys WJ, Lin L, Darvesh AS, Sadana P. Emerging strategies of targeting lipoprotein lipase for metabolic and cardiovascular diseases. Drug Discov Today. 2017;22(2):352–365. doi:10.1016/j.drudis.2016.10.007

1198. Jeron A, Hengstenberg C, Holmer S, Wollnik B, Riegger GA, Schunkert H, Erdmann J. KCNJ11 polymorphisms and sudden cardiac death in patients with acute myocardial infarction. J Mol Cell Cardiol. 2004;36(2):287–293. doi:10.1016/j.yjmcc.2003.11.009

1199. Al-Eitan LN, Almasri AY, Alnaamneh AH, Aman HA, Alrabadi NN, Khasawneh RH, Alghamdi MA. Influence of CYP4F2, ApoE, and CYP2A6 gene polymorphisms on the variability of Warfarin dosage requirements and susceptibility to cardiovascular disease in Jordan. Int J Med Sci. 2021;18(3):826–834. doi:10.7150/ijms.51546

1200. Yari A, Saleh-Gohari N, Mirzaee M, Hashemi F, Saeidi K. A Study of Associations Between rs9349379 (PHACTR1), rs2891168 (CDKN2B-AS), rs11838776 (COL4A2) and rs4880 (SOD2) Polymorphic Variants and Coronary Artery Disease in Iranian Population. Biochem Genet. 2022;60(1):106–126. doi:10.1007/s10528-021-10089-0

1201. Yang W, Ng FL, Chan K, Pu X, Poston RN, Ren M, An W, Zhang R, Wu J, Yan S, et al. Coronary-Heart-Disease-Associated Genetic Variant at the COL4A1/COL4A2 Locus Affects COL4A1/COL4A2 Expression, Vascular Cell Survival, Atherosclerotic Plaque Stability and Risk of Myocardial Infarction. PLoS Genet. 2016;12(7):e1006127. doi:10.1371/journal.pgen.1006127

1202. Cannavo A, Rengo G, Liccardo D, Pagano G, Zincarelli C, De Angelis MC, Puglia R, Di Pietro E, Rabinowitz JE, Barone MV, et al. β1-adrenergic receptor and sphingosine-1-phosphate receptor 1 (S1PR1) reciprocal downregulation influences cardiac hypertrophic response and progression to heart failure: protective role of S1PR1 cardiac gene therapy. Circulation. 2013;128(15):1612–1622. doi:10.1161/CIRCULATIONAHA.113.002659

1203. Dai J, Zhang Q, Wan C, Liu J, Zhang Q, Yu Y, Wang J. Significances of viable synergistic autophagy-associated cathepsin B and cathepsin D (CTSB/CTSD) as potential biomarkers for sudden cardiac death. BMC Cardiovasc Disord. 2021;21(1):233. doi:10.1186/s12872-021-02040-3

1204. Yu S, Shao L, Kilbride H, Zwick DL. Haploinsufficiencies of FOXF1 and FOXC2 genes associated with lethal alveolar capillary dysplasia and congenital heart disease. Am J Med Genet A. 2010;152A(5):1257–1262. doi:10.1002/ajmg.a.33378

1205. Misao J, Hayakawa Y, Ohno M, Kato S, Fujiwara T, Fujiwara H. Expression of bcl-2 protein, an inhibitor of apoptosis, and Bax, an accelerator of apoptosis, in ventricular myocytes of human hearts with myocardial infarction. Circulation. 1996;94(7):1506–1512. doi:10.1161/01.cir.94.7.1506

1206. Barascuk N, Genovese F, Larsen L, Byrjalsen I, Zheng Q, Sun S, Hosbond S, Poulsen TS, Diederichsen A, Jensen JM, et al. A MMP derived versican neo-epitope is elevated in plasma from patients with atherosclerotic heart disease. Int J Clin Exp Med. 2013;6(3):174–184.

1207. Boyle L, Wamelink MMC, Salomons GS, Roos B, Pop A, Dauber A, Hwa V, Andrew M, Douglas J, Feingold M, et al. Mutations in TKT Are the Cause of a Syndrome Including Short Stature, Developmental Delay, and Congenital Heart Defects. Am J Hum Genet. 2016;98(6):1235–1242. doi:10.1016/j.ajhg.2016.03.03

1208. Zhang X, Zhang MC, Wang CT. Loss of LRRC25 accelerates pathological cardiac hypertrophy through promoting fibrosis and inflammation regulated by TGF-β1. Biochem Biophys Res Commun. 2018;506(1):137–144. doi:10.1016/j.bbrc.2018.09.065

1209. Dai SH, Li JF, Feng JB, Li RJ, Li CB, Li Z, Zhang Y, Li DQ. Association of serum levels of AngII, KLK1, and ACE/KLK1 polymorphisms with acute myocardial infarction induced by coronary artery stenosis. J Renin Angiotensin Aldosterone Syst. 2016;17(2):1470320316655037. doi:10.1177/1470320316655037

1210. Xie Z, Shen Y, Huang S, Shen W, Liu J. Abnormal ADAMTS2 and VSIG4 in Serum of HF Patients and their Relationship with CRP, UA, and HCY. Clin Lab. 2022;68(5):10.7754/Clin.Lab.2021.210811. doi:10.7754/Clin.Lab.2021.210811

1211. Uchihashi M, Hoshino A, Okawa Y, Ariyoshi M, Kaimoto S, Tateishi S, Ono K, Yamanaka R, Hato D, Fushimura Y, et al. Cardiac-Specific Bdh1 Overexpression Ameliorates Oxidative Stress and Cardiac Remodeling in Pressure Overload-Induced Heart Failure. Circ Heart Fail. 2017;10(12):e004417. doi:10.1161/CIRCHEARTFAILURE.117.004417

1212. Jevnikar Z, Östling J, Ax E, Calvén J, Thörn K, Israelsson E, Öberg L, Singhania A, Lau LCK, Wilson SJ, et al. Epithelial IL-6 trans-signaling defines a new asthma phenotype with increased airway inflammation. J Allergy Clin Immunol. 2019;143(2):577–590. doi:10.1016/j.jaci.2018.05.026

1213. Ouyang S, Liu C, Xiao J, Chen X, Lui AC, Li X. Targeting IL-17A/glucocorticoid synergy to CSF3 expression in neutrophilic airway diseases. JCI Insight. 2020;5(3):e132836. doi:10.1172/jci.insight.132836

1214. Adage T, Konya V, Weber C, Strutzmann E, Fuchs T, Zankl C, Gerlza T, Jeremic D, Heinemann A, Kungl AJ. Targeting glycosaminoglycans in the lung by an engineered CXCL8 as a novel therapeutic approach to lung inflammation. Eur J Pharmacol. 2015;748:83–92. doi:10.1016/j.ejphar.2014.12.019

1215. Liu S, Liu J, Yang X, Jiang M, Wang Q, Zhang L, Ma Y, Shen Z, Tian Z, Cao X. Cis-acting lnc-Cxcl2 restrains neutrophil-mediated lung inflammation by inhibiting epithelial cell CXCL2 expression in virus infection. Proc Natl Acad Sci U S A. 2021;118(41):e2108276118. doi:10.1073/pnas.2108276118

1216. Kimura MY, Hayashizaki K, Tokoyoda K, Takamura S, Motohashi S, Nakayama T. Crucial role for CD69 in allergic inflammatory responses: CD69-Myl9 system in the pathogenesis of airway inflammation. Immunol Rev. 2017;278(1):87–100. doi:10.1111/imr.12559

1217. Sharma S, Ursery LT, Bharathi V, Miles SD, Williams WA, Elzawam AZ, Schmedes CM, Egnatz GJ, Fernandez JA, Palumbo JS, et al. APC-PAR1-R46 signaling limits CXCL1 expression during poly IC-induced airway inflammation in mice. J Thromb Haemost. 2023;21(11):3279–3282. doi:10.1016/j.jtha.2023.08.018

1218. Lear TB, McKelvey AC, Evankovich JW, Rajbhandari S, Coon TA, Dunn SR, Londino JD, McVerry BJ, Zhang Y, Valenzi E, et al. KIAA0317 regulates pulmonary inflammation through SOCS2 degradation. JCI Insight. 2019;4(19):e129110. doi:10.1172/jci.insight.129110

1219. Draijer C, Speth JM, Penke LRK, Zaslona Z, Bazzill JD, Lugogo N, Huang YJ, Moon JJ, Peters-Golden M. Resident alveolar macrophage-derived vesicular SOCS3 dampens allergic airway inflammation. FASEB J. 2020;34(3):4718–4731. doi:10.1096/fj.201903089R

1220. John AE, Berlin AA, Lukacs NW. Respiratory syncytial virus-induced CCL5/RANTES contributes to exacerbation of allergic airway inflammation. Eur J Immunol. 2003;33(6):1677–1685. doi:10.1002/eji.200323930

1221. Perkins TN, Oczypok EA, Milutinovic PS, Dutz RE, Oury TD. RAGE-dependent VCAM-1 expression in the lung endothelium mediates IL-33-induced allergic airway inflammation. Allergy. 2019;74(1):89–99. doi:10.1111/all.13500

1222. Hong L, Wang Q, Chen M, Shi J, Guo Y, Liu S, Pan R, Yuan X, Jiang S. Mas receptor activation attenuates allergic airway inflammation via inhibiting JNK/CCL2-induced macrophage recruitment. Biomed Pharmacother. 2021;137:111365. doi:10.1016/j.biopha.2021.111365

1223. Liu Y, Huo SG, Xu L, Che YY, Jiang SY, Zhu L, Zhao M, Teng YC. MiR-135b Alleviates Airway Inflammation in Asthmatic Children and Experimental Mice with Asthma via Regulating CXCL12. Immunol Invest. 2022;51(3):496–510. doi:10.1080/08820139.2020.1841221

1224. Huang LT, Chang HW, Wu MJ, Lai YT, Wu WC, Yu WCY, Chang VHS. Klf10 deficiency in mice exacerbates pulmonary inflammation by increasing expression of the proinflammatory molecule NPRA. Int J Biochem Cell Biol. 2016;79:231–238. doi:10.1016/j.biocel.2016.08.027

1225. Nakano K, Whitehead GS, Lyons-Cohen MR, Grimm SA, Wilkinson CL, Izumi G, Livraghi-Butrico A, Cook DN, Nakano H. Chemokine CCL19 promotes type 2 T-cell differentiation and allergic airway inflammation. J Allergy Clin Immunol. 2023. doi:10.1016/j.jaci.2023.10.024

1226. Besnard AG, Struyf S, Guabiraba R, Fauconnier L, Rouxel N, Proost P, Uyttenhove C, Van Snick J, Couillin I, Ryffel B. CXCL6 antibody neutralization prevents lung inflammation and fibrosis in mice in the bleomycin model. J Leukoc Biol. 2013;94(6):1317–1323. doi:10.1189/jlb.0313140

1227. Yi G, Liang M, Li M, Fang X, Liu J, Lai Y, Chen J, Yao W, Feng X, Hu L, et al. A large lung gene expression study identifying IL1B as a novel player in airway inflammation in COPD airway epithelial cells. Inflamm Res. 2018;67(6):539–551. doi:10.1007/s00011-018-1145-8

1228. Wang J, Zhu M, Wang L, Chen C, Song Y. Amphiregulin potentiates airway inflammation and mucus hypersecretion induced by urban particulate matter via the EGFR-PI3Kα-AKT/ERK pathway. Cell Signal. 2019;53:122–131. doi:10.1016/j.cellsig.2018.10.002

1229. Headland SE, Dengler HS, Xu D, Teng G, Everett C, Ratsimandresy RA, Yan D, Kang J, Ganeshan K, Nazarova EV, et al. Oncostatin M expression induced by bacterial triggers drives airway inflammatory and mucus secretion in severe asthma. Sci Transl Med. 2022;14(627):eabf8188. doi:10.1126/scitranslmed.abf8188

1230. Shaw OM, Nyanhanda T, McGhie TK, Harper JL, Hurst RD. Blackcurrant anthocyanins modulate CCL11 secretion and suppress allergic airway inflammation. Mol Nutr Food Res. 2017;61(9):10.1002/mnfr.201600868. doi:10.1002/mnfr.201600868

1231. Ye C, Zhang N, Zhao Q, Xie X, Li X, Zhu HP, Peng C, Huang W, Han B. Evodiamine alleviates lipopolysaccharide-induced pulmonary inflammation and fibrosis by activating apelin pathway. Phytother Res. 2021;35(6):3406–3417. doi:10.1002/ptr.7062

1232. Madenspacher JH, Morrell ED, Gowdy KM, McDonald JG, Thompson BM, Muse G, Martinez J, Thomas S, Mikacenic C, Nick JA, et al. Cholesterol 25-hydroxylase promotes efferocytosis and resolution of lung inflammation. JCI Insight. 2020;5(11):e137189. doi:10.1172/jci.insight.137189

1233. Zhang Y, Yan L, Yang J, Li X. Silencing of FSTL1 Alleviated LPS-Induced Inflammatory Damage and Oxidative Damage in Human Bronchial Epithelial Cells via BMP4/KLF4 Axis. Int Arch Allergy Immunol. 2022;183(7):785–795. doi:10.1159/000521852

1234. Coyle AJ, Tsuyuki S, Bertrand C, Huang S, Aguet M, Alkan SS, Anderson GP. Mice lacking the IFN-gamma receptor have impaired ability to resolve a lung eosinophilic inflammatory response associated with a prolonged capacity of T cells to exhibit a Th2 cytokine profile. J Immunol. 1996;156(8):2680–2685.

1235. Kurakula K, Vos M, Logiantara A, Roelofs JJ, Nieuwenhuis MA, Koppelman GH, Postma DS, Brandsma CA, Sin DD, Bossé Y, et al. Deficiency of FHL2 attenuates airway inflammation in mice and genetic variation associates with human bronchial hyper-responsiveness. Allergy. 2015;70(12):1531–1544. doi:10.1111/all.12709

1236. Godwin MS, Jones M, Blackburn JP, Yu Z, Matalon S, Hastie AT, Meyers DA, Steele C. The chemokine CX3CL1/fractalkine regulates immunopathogenesis during fungal-associated allergic airway inflammation. Am J Physiol Lung Cell Mol Physiol. 2021;320(3):L393–L404. doi:10.1152/ajplung.00376.2020

1237. Xu Z, Ye Y, Huang G, Li Y, Guo X, Li L, Wu Y, Xu W, Nian S, Yuan Q. EphA2 recognizes Dermatophagoidespteronyssinus to mediate airway inflammation in asthma. Int Immunopharmacol. 2022;111:109106. doi:10.1016/j.intimp.2022.109106

1238. Knight D. Leukaemia inhibitory factor (LIF): a cytokine of emerging importance in chronic airway inflammation. Pulm Pharmacol Ther. 2001;14(3):169–176. doi:10.1006/pupt.2001.0282

1239. Shankar SP, Wilson MS, DiVietro JA, Mentink-Kane MM, Xie Z, Wynn TA, Druey KM. RGS16 attenuates pulmonary Th2/Th17 inflammatory responses. J Immunol. 2012;188(12):6347–6356. doi:10.4049/jimmunol.1103781

1240. Kawakami M, Narumoto O, Matsuo Y, Horiguchi K, Horiguchi S, Yamashita N, Sakaguchi M, Lipp M, Nagase T, Yamashita N. The role of CCR7 in allergic airway inflammation induced by house dust mite exposure. Cell Immunol. 2012;275(1-2):24–32. doi:10.1016/j.cellimm.2012.03.009

1241. Yokoyama Y, Tamachi T, Iwata A, Maezawa Y, Meguro K, Yokota M, Takatori H, Suto A, Suzuki K, Hirose K, et al. A20 (Tnfaip3) expressed in CD4+ T cells suppresses Th2 cell-mediated allergic airway inflammation in mice. Biochem Biophys Res Commun. 2022;629:47–53. doi:10.1016/j.bbrc.2022.08.097

1242. Li Z, Wang J, Wang Y, Jiang H, Xu X, Zhang C, Li D, Xu C, Zhang K, Qi Y, et al. Bone morphogenetic protein 4 inhibits liposaccharide-induced inflammation in the airway. Eur J Immunol. 2014;44(11):3283–3294. doi:10.1002/eji.201344287

1243. Nadeem A, Ahmad SF, Al-Harbi NO, Ibrahim KE, Siddiqui N, Al-Harbi MM, Attia SM, Bakheet SA. Inhibition of Bruton’s tyrosine kinase and IL-2 inducible T-cell kinase suppresses both neutrophilic and eosinophilic airway inflammation in a cockroach allergen extract-induced mixed granulocytic mouse model of asthma using preventative and therapeutic strategy. Pharmacol Res. 2019;148:104441. doi:10.1016/j.phrs.2019.104441

1244. Chong DLW, Rebeyrol C, José RJ, Williams AE, Brown JS, Scotton CJ, Porter JC. ICAM-1 and ICAM-2 Are Differentially Expressed and Up-Regulated on Inflamed Pulmonary Epithelium, but Neither ICAM-2 nor LFA-1: ICAM-1 Are Required for Neutrophil Migration Into the Airways In Vivo. Front Immunol. 2021;12:691957. doi:10.3389/fimmu.2021.691957

1245. Mao H, Wang ZL, Li FY, Liu CT, Lei S. Relationship between bone marrow-derived CD34 + cells expressing interleukin-5 receptor messenger RNA and asthmatic airway inflammation. Chin Med J (Engl). 2004;117(1):24–29.

1246. Hong H, Liao S, Chen F, Yang Q, Wang DY. Role of IL-25, IL-33, and TSLP in triggering united airway diseases toward type 2 inflammation. Allergy. 2020;75(11):2794–2804. doi:10.1111/all.14526

1247. Oshima K, Han X, Ouyang Y, El Masri R, Yang Y, Haeger SM, McMurtry SA, Lane TC, Davizon-Castillo P, Zhang F, et al. Loss of endothelial sulfatase-1 after experimental sepsis attenuates subsequent pulmonary inflammatory responses. Am J Physiol Lung Cell Mol Physiol. 2019;317(5):L667–L677. doi:10.1152/ajplung.00175.2019

1248. Lee SN, Yoon SA, Song JM, Kim HC, Cho HJ, Choi AMK, Yoon JH. Cell-Type-Specific Expression of Hyaluronan Synthases HAS2 and HAS3 Promotes Goblet Cell Hyperplasia in Allergic Airway Inflammation. Am J Respir Cell Mol Biol. 2022;67(3):360–374. doi:10.1165/rcmb.2021-0527OC

1249. Kearns MT, Barthel L, Bednarek JM, Yunt ZX, Henson PM, Janssen WJ. Fas ligand-expressing lymphocytes enhance alveolar macrophage apoptosis in the resolution of acute pulmonary inflammation. Am J Physiol Lung Cell Mol Physiol. 2014;307(1):L62–L70. doi:10.1152/ajplung.00273.2013

1250. Dahm PH, Richards JB, Karmouty-Quintana H, Cromar KR, Sur S, Price RE, Malik F, Spencer CY, Barreno RX, Hashmi SS, et al. Effect of antigen sensitization and challenge on oscillatory mechanics of the lung and pulmonary inflammation in obese carboxypeptidase E-deficient mice. Am J Physiol Regul Integr Comp Physiol. 2014;307(6):R621–R633. doi:10.1152/ajpregu.00205.2014

1251. Faustino L, Fonseca DM, Florsheim EB, Resende RR, Lepique AP, Faquim-Mauro E, Gomes E, Silva JS, Yagita H, Russo M. Tumor necrosis factor-related apoptosis-inducing ligand mediates the resolution of allergic airway inflammation induced by chronic allergen inhalation. Mucosal Immunol. 2014;7(5):1199–1208. doi:10.1038/mi.2014.9

1252. Verhamme FM, Seys LJM, De Smet EG, Provoost S, Janssens W, Elewaut D, Joos GF, Brusselle GG, Bracke KR. Elevated GDF-15 contributes to pulmonary inflammation upon cigarette smoke exposure. Mucosal Immunol. 2017;10(6):1400–1411. doi:10.1038/mi.2017.3

1253. Perros F, Hoogsteden HC, Coyle AJ, Lambrecht BN, Hammad H. Blockade of CCR4 in a humanized model of asthma reveals a critical role for DC-derived CCL17 and CCL22 in attracting Th2 cells and inducing airway inflammation. Allergy. 2009;64(7):995–1002. doi:10.1111/j.1398-9995.2009.02095.x

1254. Faustino L, da Fonseca DM, Takenaka MC, Mirotti L, Florsheim EB, Guereschi MG, Silva JS, Basso AS, Russo M. Regulatory T cells migrate to airways via CCR4 and attenuate the severity of airway allergic inflammation. J Immunol. 2013;190(6):2614–2621. doi:10.4049/jimmunol.1202354

1255. Zhu B, Wu G, Wang C, Xiao Y, Jin J, Wang K, Jiang Y, Sun Y, Ben D, Xia Z. Soluble cluster of differentiation 74 regulates lung inflammation through the nuclear factor-κB signaling pathway. Immunobiology. 2020;225(5):152007. doi:10.1016/j.imbio.2020.152007

1256. Meunier S, Chea S, Garrido D, Perchet T, Petit M, Cumano A, Golub R. Maintenance of Type 2 Response by CXCR6-Deficient ILC2 in Papain-Induced Lung Inflammation. Int J Mol Sci. 2019;20(21):5493. doi:10.3390/ijms20215493

1257. Yang T, Wang H, Li Y, Zeng Z, Shen Y, Wan C, Wu Y, Dong J, Chen L, Wen F. Serotonin receptors 5-HTR2A and 5-HTR2B are involved in cigarette smoke-induced airway inflammation, mucus hypersecretion and airway remodeling in mice. Int Immunopharmacol. 2020;81:106036. doi:10.1016/j.intimp.2019.106036

1258. Zhao Z, Qian Y, Wald D, Xia YF, Geng JG, Li X. IFN regulatory factor-1 is required for the up-regulation of the CD40-NF-kappa B activator 1 axis during airway inflammation. J Immunol. 2003;170(11):5674–5680. doi:10.4049/jimmunol.170.11.5674

1259. de Vries M, Heijink IH, Gras R, den Boef LE, Reinders-Luinge M, Pouwels SD, Hylkema MN, van der Toorn M, Brouwer U, van Oosterhout AJ, et al. Pim1 kinase protects airway epithelial cells from cigarette smoke-induced damage and airway inflammation. Am J Physiol Lung Cell Mol Physiol. 2014;307(3):L240–L251. doi:10.1152/ajplung.00156.2013

1260. Felton JM, Dorward DA, Cartwright JA, Potey PM, Robb CT, Gui J, Craig RW, Schwarze J, Haslett C, Duffin R, et al. Mcl-1 protects eosinophils from apoptosis and exacerbates allergic airway inflammation. Thorax. 2020;75(7):600–605. doi:10.1136/thoraxjnl-2019-213204

1261. Ge XN, Bastan I, Dileepan M, Greenberg Y, Ha SG, Steen KA, Bernlohr DA, Rao SP, Sriramarao P. FABP4 regulates eosinophil recruitment and activation in allergic airway inflammation. Am J Physiol Lung Cell Mol Physiol. 2018;315(2):L227–L240. doi:10.1152/ajplung.00429.2017

1262. Jiang T, Zhao D, Zheng Z, Li Z. Sigma-1 Receptor Alleviates Airway Inflammation and Airway Remodeling Through AMPK/CXCR4 Signal Pathway. Inflammation. 2022;45(3):1298–1312. doi:10.1007/s10753-022-01621-4

1263. Blanquart E, Mandonnet A, Mars M, Cenac C, Anesi N, Mercier P, Audouard C, Roga S, Serrano de Almeida G, Bevan CL, Girard JP, et al. Targeting androgen signaling in ILC2s protects from IL-33-driven lung inflammation, independently of KLRG1. J Allergy Clin Immunol. 2022;149(1):237–251.e12. doi:10.1016/j.jaci.2021.04.029

1264. Ribeiro CM, Lubamba BA. Role of IRE1α/XBP-1 in Cystic Fibrosis Airway Inflammation. Int J Mol Sci. 2017;18(1):118. doi:10.3390/ijms18010118

1265. Ferreira TPT, Guimarães FV, Sá YAPJ, da Silva Ribeiro NB, de Arantes ACS, de Frias Carvalho V, Sousa LP, Perretti M, Martins MA, et al. Annexin-A1-Derived Peptide Ac2-26 Suppresses Allergic Airway Inflammation and Remodelling in Mice. Cells. 2022;11(5):759. doi:10.3390/cells11050759

1266. Sharma N, Nagaraj C, Nagy BM, Marsh LM, Bordag N, Zabini D, Wygrecka M, Klepetko W, Gschwandtner E, Genové G, et al. RGS5 Determines Neutrophil Migration in the Acute Inflammatory Phase of Bleomycin-Induced Lung Injury. Int J Mol Sci. 2021;22(17):9342. doi:10.3390/ijms22179342

1267. Gueders MM, Hirst SJ, Quesada-Calvo F, Paulissen G, Hacha J, Gilles C, Gosset P, Louis R, Foidart JM, Lopez-Otin C, et al. Matrix metalloproteinase-19 deficiency promotes tenascin-C accumulation and allergen-induced airway inflammation. Am J Respir Cell Mol Biol. 2010;43(3):286–295. doi:10.1165/rcmb.2008-0426OC

1268. Weijer S, Wieland CW, Florquin S, van der Poll T. A thrombomodulin mutation that impairs activated protein C generation results in uncontrolled lung inflammation during murine tuberculosis. Blood. 2005;106(8):2761–2768. doi:10.1182/blood-2004-12-4623

1269. Wuttge DM, Andreasson A, Tufvesson E, Johansson ÅC, Scheja A, Hellmark T, Hesselstrand R, Truedsson L. CD81 and CD48 show different expression on blood eosinophils in systemic sclerosis: new markers for disease and pulmonary inflammation?. Scand J Rheumatol. 2016;45(2):107–113. doi:10.3109/03009742.2015.1054877

1270. Palmer LD, Maloney KN, Boyd KL, Goleniewska AK, Toki S, Maxwell CN, Chazin WJ, Peebles RS Jr, Newcomb DC, Skaar EP. The Innate Immune Protein S100A9 Protects from T-Helper Cell Type 2-mediated Allergic Airway Inflammation. Am J Respir Cell Mol Biol. 2019;61(4):459–468. doi:10.1165/rcmb.2018-0217OC

1271. Barron L, Smith AM, El Kasmi KC, Qualls JE, Huang X, Cheever A, Borthwick LA, Wilson MS, Murray PJ, Wynn TA. Role of arginase 1 from myeloid cells in th2-dominated lung inflammation. PLoS One. 2013;8(4):e61961. doi:10.1371/journal.pone.0061961

1272. Ijaz B, Shabbir A, Shahzad M, Mobashar A, Sharif M, Basheer MI, Tareen RB, Syed NI. Amelioration of airway inflammation and pulmonary edema by Teucrium stocksianum via attenuation of pro-inflammatory cytokines and up-regulation of AQP1 and AQP5. Respir Physiol Neurobiol. 2021;284:103569. doi:10.1016/j.resp.2020.103569

1273. Thakur SA, Beamer CA, Migliaccio CT, Holian A. Critical role of MARCO in crystalline silica-induced pulmonary inflammation. Toxicol Sci. 2009;108(2):462–471. doi:10.1093/toxsci/kfp011

1274. Plovsing RR, Berg RM, Munthe-Fog L, Konge L, Iversen M, Møller K, Garred P.Alveolar recruitment of ficolin-3 in response to acute pulmonary inflammation in humans. Immunobiology. 2016;221(5):690–697. doi:10.1016/j.imbio.2015.11.015

1275. Li H, Edin ML, Bradbury JA, Graves JP, DeGraff LM, Gruzdev A, Cheng J, Dackor RT, Wang PM, Bortner CD, et al. Cyclooxygenase-2 inhibits T helper cell type 9 differentiation during allergic lung inflammation via down-regulation of IL-17RB. Am J Respir Crit Care Med. 2013;187(8):812–822. doi:10.1164/rccm.201211-2073OC

1276. Schleimer RP, Schnaar RL, Bochner BS. Regulation of airway inflammation by Siglec-8 and Siglec-9 sialoglycan ligand expression. Curr Opin Allergy Clin Immunol. 2016;16(1):24–30. doi:10.1097/ACI.0000000000000234

1277. Wu Y, Zhang W, Gunst SJ. S100A4 is secreted by airway smooth muscle tissues and activates inflammatory signaling pathways via receptors for advanced glycation end products. Am J Physiol Lung Cell Mol Physiol. 2020;319(1):L185–L195. doi:10.1152/ajplung.00347.2019

1278. Han X, Liu L, Huang S, Xiao W, Gao Y, Zhou W, Zhang C, Zheng H, Yang L, Xie X, et al. RNA m6A methylation modulates airway inflammation in allergic asthma via PTX3-dependent macrophage homeostasis. Nat Commun. 2023;14(1):7328. doi:10.1038/s41467-023-43219-w

1279. Zhang S, Lu X, Fang X, Wang Z, Cheng S, Song J. Cigarette smoke extract combined with LPS reduces ABCA3 expression in chronic pulmonary inflammation may be related to PPARγ/ P38 MAPK signaling pathway. Ecotoxicol Environ Saf. 2022;244:114086. doi:10.1016/j.ecoenv.2022.114086

1280. Poole JA, Romberger DJ, Bauer C, Gleason AM, Sisson JH, Oldenburg PJ, West WW, Wyatt TA. Protein kinase C epsilon is important in modulating organic-dust-induced airway inflammation. Exp Lung Res. 2012;38(8):383–395. doi:10.3109/01902148.2012.714841

1281. Krick S, Grabner A, Baumlin N, Yanucil C, Helton S, Grosche A, Sailland J, Geraghty P, Viera L, Russell DW, et al. Fibroblast growth factor 23 and Klotho contribute to airway inflammation. Eur Respir J. 2018;52(1):1800236. doi:10.1183/13993003.00236-2018

1282. Tang W, Dong M, Teng F, Cui J, Zhu X, Wang W, Wuniqiemu T, Qin J, Yi L, Wang S, et al. TMT-based quantitative proteomics reveals suppression of SLC3A2 and ATP1A3 expression contributes to the inhibitory role of acupuncture on airway inflammation in an OVA-induced mouse asthma model. Biomed Pharmacother. 2021;134:111001. doi:10.1016/j.biopha.2020.111001

1283. Chen K, Liu M, Liu Y, Wang C, Yoshimura T, Gong W, Le Y, Tessarollo L, Wang JM. Signal relay by CC chemokine receptor 2 (CCR2) and formylpeptide receptor 2 (Fpr2) in the recruitment of monocyte-derived dendritic cells in allergic airway inflammation. J Biol Chem. 2013;288(23):16262–16273. doi:10.1074/jbc.M113.450635

1284. Nguyen QT, Jang E, Le HT, Kim S, Kim D, Dvorina N, Aronica MA, Baldwin WM 3rd, Asosingh K, Comhair S, et al. IL-27 targets Foxp3+ Tregs to mediate antiinflammatory functions during experimental allergic airway inflammation. JCI Insight. 2019;4(2):e123216. doi:10.1172/jci.insight.123216

1285. Dickerhof N, Huang J, Min E, Michaëlsson E, Lindstedt EL, Pearson JF, Kettle AJ, Day BJ. Myeloperoxidase inhibition decreases morbidity and oxidative stress in mice with cystic fibrosis-like lung inflammation. Free Radic Biol Med. 2020;152:91–99. doi:10.1016/j.freeradbiomed.2020.03.001

1286. Abonyo BO, Alexander MS, Heiman AS. Autoregulation of CCL26 synthesis and secretion in A549 cells: a possible mechanism by which alveolar epithelial cells modulate airway inflammation. Am J Physiol Lung Cell Mol Physiol. 2005;289(3):L478–L488. doi:10.1152/ajplung.00032.2005

1287. Zhou Y, Huang X, Yu H, Shi H, Chen M, Song J, Tang W, Teng F, Li C, Yi L, et al. TMT-based quantitative proteomics revealed protective efficacy of Icariside II against airway inflammation and remodeling via inhibiting LAMP2, CTSD and CTSS expression in OVA-induced chronic asthma mice. Phytomedicine. 2023;118:154941. doi:10.1016/j.phymed.2023.154941

1288. Liu L, Zhou L, Wang L, Mao Z, Zheng P, Zhang F, Zhang H, Liu H. MUC1 attenuates neutrophilic airway inflammation in asthma by reducing NLRP3 inflammasome-mediated pyroptosis through the inhibition of the TLR4/MyD88/NF-κB pathway. Respir Res. 2023;24(1):255. doi:10.1186/s12931-023-02550-y

1289. Kalin TV, Meliton L, Meliton AY, Zhu X, Whitsett JA, Kalinichenko VV. Pulmonary mastocytosis and enhanced lung inflammation in mice heterozygous null for the Foxf1 gene. Am J Respir Cell Mol Biol. 2008;39(4):390–399. doi:10.1165/rcmb.2008-0044OC

1290. Pilzner C, Bühling F, Reinheckel T, Chwieralski C, Rathinasamy A, Lauenstein HD, Wex T, Welte T, Braun A, Groneberg DA. Allergic airway inflammation in mice deficient for the antigen-processing protease cathepsin E. Int Arch Allergy Immunol. 2012;159(4):367–383. doi:10.1159/000338288

1291. Epstein Shochet G, Brook E, Bardenstein-Wald B, Shitrit D. TGF-β pathway activation by idiopathic pulmonary fibrosis (IPF) fibroblast derived soluble factors is mediated by IL-6 trans-signaling. Respir Res. 2020;21(1):56. doi:10.1186/s12931-020-1319-0

1292. Roels E, Krafft E, Farnir F, Holopainen S, Laurila HP, Rajamäki MM, Day MJ, Antoine N, Pirottin D, Clercx C. Assessment of CCL2 and CXCL8 chemokines in serum, bronchoalveolar lavage fluid and lung tissue samples from dogs affected with canine idiopathic pulmonary fibrosis. Vet J. 2015;206(1):75–82. doi:10.1016/j.tvjl.2015.06.001aaa

1293. Nie Y, Zhai X, Li J, Sun A, Che H, Christman JW, Chai G, Zhao P, Karpurapu M. NFATc3 Promotes Pulmonary Inflammation and Fibrosis by Regulating Production of CCL2 and CXCL2 in Macrophages. Aging Dis. 2023;14(4):1441–1457. doi:10.14336/AD.2022.1202

1294. Schiller HB, Mayr CH, Leuschner G, Strunz M, Staab-Weijnitz C, Preisendörfer S, Eckes B, Moinzadeh P, Krieg T, Schwartz DA, et al. Deep Proteome Profiling Reveals Common Prevalence of MZB1-Positive Plasma B Cells in Human Lung and Skin Fibrosis. Am J Respir Crit Care Med. 2017;196(10):1298–1310. doi:10.1164/rccm.201611-2263OC

1295. Yamauchi K, Kasuya Y, Kuroda F, Tanaka K, Tsuyusaki J, Ishizaki S, Matsunaga H, Iwamura C, Nakayama T, Tatsumi K. Attenuation of lung inflammation and fibrosis in CD69-deficient mice after intratracheal bleomycin. Respir Res. 2011;12(1):131. doi:10.1186/1465-9921-12-131

1296. Cao Y, Huang W, Wu F, Shang J, Ping F, Wang W, Li Y, Zhao X, Zhang X. ZFP36 protects lungs from intestinal I/R-induced injury and fibrosis through the CREBBP/p53/p21/Bax pathway. Cell Death Dis. 2021;12(7):685. doi:10.1038/s41419-021-03950-y

1297. Yang X, Walton W, Cook DN, Hua X, Tilley S, Haskell CA, Horuk R, Blackstock AW, Kirby SL. The chemokine, CCL3, and its receptor, CCR1, mediate thoracic radiation-induced pulmonary fibrosis. Am J Respir Cell Mol Biol. 2011;45(1):127–135. doi:10.1165/rcmb.2010-0265OC

1298. Emad A, Emad Y. Relationship between eosinophilia and levels of chemokines (CCL5 and CCL11) and IL-5 in bronchoalveolar lavage fluid of patients with mustard gas-induced pulmonary fibrosis. J Clin Immunol. 2008;28(4):298–305. doi:10.1007/s10875-007-9109-8

1299. Pulito-Cueto V, Remuzgo-Martínez S, Genre F, Atienza-Mateo B, Mora-Cuesta VM, Iturbe-Fernández D, Lera-Gómez L, Sebastián Mora-Gil M, Prieto-Peña D, Portilla V, et al. Elevated VCAM-1, MCP-1 and ADMA serum levels related to pulmonary fibrosis of interstitial lung disease associated with rheumatoid arthritis. Front Mol Biosci. 2022;9:1056121. doi:10.3389/fmolb.2022.1056121

1300. Gui X, Qiu X, Tian Y, Xie M, Li H, Gao Y, Zhuang Y, Cao M, Ding H, Ding J, et al. Prognostic value of IFN-γ, sCD163, CCL2 and CXCL10 involved in acute exacerbation of idiopathic pulmonary fibrosis. Int Immunopharmacol. 2019;70:208–215. doi:10.1016/j.intimp.2019.02.039

1301. Vuga LJ, Tedrow JR, Pandit KV, Tan J, Kass DJ, Xue J, Chandra D, Leader JK, Gibson KF, Kaminski N, et al. C-X-C motif chemokine 13 (CXCL13) is a prognostic biomarker of idiopathic pulmonary fibrosis. Am J Respir Crit Care Med. 2014;189(8):966–974. doi:10.1164/rccm.201309-1592OC

1302. Song X, Xu P, Meng C, Song C, Blackwell TS, Li R, Li H, Zhang J, Lv C. lncITPF Promotes Pulmonary Fibrosis by Targeting hnRNP-L Depending on Its Host Gene ITGBL1. Mol Ther. 2019;27(2):380–393. doi:10.1016/j.ymthe.2018.08.026

1303. Fukushima K, Akira S. Novel insights into the pathogenesis of lung fibrosis: the RBM7-NEAT1-CXCL12-SatM axis at fibrosis onset. Int Immunol. 2021;33(12):659–663. doi:10.1093/intimm/dxab034

1304. Bahudhanapati H, Tan J, Apel RM, Seeliger B, Schupp J, Li X, Sullivan DI, Sembrat J, Rojas M, Tabib T, et al. Increased Expression of CXCL6 in Secretory Cells Drives Fibroblast Collagen Synthesis and is Associated With Increased Mortality in Idiopathic Pulmonary Fibrosis. Eur Respir J. 2023. doi:10.1183/13993003.00088-2023

1305. Barlo NP, van Moorsel CH, Korthagen NM, Heron M, Rijkers GT, Ruven HJ, van den Bosch JM, Grutters JC. Genetic variability in the IL1RN gene and the balance between interleukin (IL)-1 receptor agonist and IL-1β in idiopathic pulmonary fibrosis. Clin Exp Immunol. 2011;166(3):346–351. doi:10.1111/j.1365-2249.2011.04468.x

1306. Ding L, Liu T, Wu Z, Hu B, Nakashima T, Ullenbruch M, Gonzalez De Los Santos F, Phan SH. Bone Marrow CD11c+ Cell-Derived Amphiregulin Promotes Pulmonary Fibrosis. J Immunol. 2016;197(1):303–312. doi:10.4049/jimmunol.1502479

1307. Jiang C, Liu G, Luckhardt T, Antony V, Zhou Y, Carter AB, Thannickal VJ, Liu RM. Serpine 1 induces alveolar type II cell senescence through activating p53-p21-Rb pathway in fibrotic lung disease. Aging Cell. 2017;16(5):1114–1124. doi:10.1111/acel.12643

1308. Lan YW, Theng SM, Huang TT, Choo KB, Chen CM, Kuo HP, Chong KY. Oncostatin M-Preconditioned Mesenchymal Stem Cells Alleviate Bleomycin-Induced Pulmonary Fibrosis Through Paracrine Effects of the Hepatocyte Growth Factor. Stem Cells Transl Med. 2017;6(3):1006–1017. doi:10.5966/sctm.2016-0054

1309. Luo L, Wang CC, Song XP, Wang HM, Zhou H, Sun Y, Wang XK, Hou S, Pei FY. Suppression of SMOC2 reduces bleomycin (BLM)-induced pulmonary fibrosis by inhibition of TGF-β1/SMADs pathway. Biomed Pharmacother. 2018;105:841–847. doi:10.1016/j.biopha.2018.03.058

1310. Mölleken C, Poschmann G, Bonella F, Costabel U, Sitek B, Stühler K, Meyer HE, Schmiegel WH, Marcussen N, Helmer M, et al. MFAP4: a candidate biomarker for hepatic and pulmonary fibrosis?. Sarcoidosis Vasc Diffuse Lung Dis. 2016;33(1):41–50.

1311. Monkley S, Overed-Sayer C, Parfrey H, Rassl D, Crowther D, Escudero-Ibarz L, Davis N, Carruthers A, Berks R, Coetzee M, et al. Sensitization of the UPR by loss of PPP1R15A promotes fibrosis and senescence in IPF. Sci Rep. 2021;11(1):21584. doi:10.1038/s41598-021-00769-7

1312. Li J, Pan C, Tang C, Tan W, Zhang W, Guan J. miR-184 targets TP63 to block idiopathic pulmonary fibrosis by inhibiting proliferation and epithelial-mesenchymal transition of airway epithelial cells. Lab Invest. 2021;101(2):142–154. doi:10.1038/s41374-020-00487-0

1313. Yin Q, Morris GF, Saito S, Zhuang Y, Thannickal VJ, Jazwinski SM, Lasky JA. Enhanced Expression of a Novel Lamin A/C Splice Variant in Idiopathic Pulmonary Fibrosis Lung. Am J Respir Cell Mol Biol. 2023;68(6):625–637. doi:10.1165/rcmb.2022-0222OC

1314. Sha JM, Zhang RQ, Wang XC, Zhou Y, Song K, Sun H, Tu B, Tao H. Epigenetic reader MeCP2 repressed WIF1 boosts lung fibroblast proliferation, migration and pulmonary fibrosis. Toxicol Lett. 2023;381:1–12. doi:10.1016/j.toxlet.2023.04.004

1315. Wei S, Qi F, Wu Y, Liu X. Overexpression of KLF4 Suppresses Pulmonary Fibrosis through the HIF-1α/Endoplasmic Reticulum Stress Signaling Pathway. Int J Mol Sci. 2023;24(18):14008. doi:10.3390/ijms241814008

1316. Barlo NP, van Moorsel CH, Korthagen NM, Heron M, Rijkers GT, Ruven HJ, van den Bosch JM, Grutters JC. Genetic variability in the IL1RN gene and the balance between interleukin (IL)-1 receptor agonist and IL-1β in idiopathic pulmonary fibrosis. Clin Exp Immunol. 2011;166(3):346–351. doi:10.1111/j.1365-2249.2011.04468.x

1317. Fusiak T, Smaldone GC, Condos R. Pulmonary Fibrosis Treated with Inhaled Interferon-gamma (IFN-γ). J Aerosol Med Pulm Drug Deliv. 2015;28(5):406–410. doi:10.1089/jamp.2015.1221

1318. Shi M, Cui H, Shi J, Mei Y. Silencing FHL2 inhibits bleomycin-induced pulmonary fibrosis through the TGF-β1/Smad signaling pathway. Exp Cell Res. 2023;423(2):113470. doi:10.1016/j.yexcr.2023.113470

1319. Rivas-Fuentes S, Herrera I, Salgado-Aguayo A, Buendía-Roldán I, Becerril C, Cisneros J. CX3CL1 and CX3CR1 could be a relevant molecular axis in the pathophysiology of idiopathic pulmonary fibrosis. Int J Med Sci. 2020;17(15):2357–2361. doi:10.7150/ijms.43748

1320. Leem AY, Shin MH, Douglas IS, Song JH, Chung KS, Kim EY, Jung JY, Kang YA, Chang J, Kim YS, et al. All-trans retinoic acid attenuates bleomycin-induced pulmonary fibrosis via downregulating EphA2-EphrinA1 signaling. Biochem Biophys Res Commun. 2017;491(3):721–726. doi:10.1016/j.bbrc.2017.07.122

1321. Dai WJ, Qiu J, Sun J, Ma CL, Huang N, Jiang Y, Zeng J, Ren BC, Li WC, Li YH. Downregulation of microRNA-9 reduces inflammatory response and fibroblast proliferation in mice with idiopathic pulmonary fibrosis through the ANO1-mediated TGF-β-Smad3 pathway. J Cell Physiol. 2019;234(3):2552–2565. doi:10.1002/jcp.26961

1322. Shi X, Chen Y, Liu Q, Mei X, Liu J, Tang Y, Luo R, Sun D, Ma Y, Wu W, et al. LDLR dysfunction induces LDL accumulation and promotes pulmonary fibrosis. Clin Transl Med. 2022;12(1):e711. doi:10.1002/ctm2.711

1323. Pierce EM, Carpenter K, Jakubzick C, Kunkel SL, Flaherty KR, Martinez FJ, Hogaboam CM. Therapeutic targeting of CC ligand 21 or CC chemokine receptor 7 abrogates pulmonary fibrosis induced by the adoptive transfer of human pulmonary fibroblasts to immunodeficient mice. Am J Pathol. 2007;170(4):1152–1164. doi:10.2353/ajpath.2007.060649

1324. Guan R, Yuan L, Li J, Wang J, Li Z, Cai Z, Guo H, Fang Y, Lin R, Liu W, et al. Bone morphogenetic protein 4 inhibits pulmonary fibrosis by modulating cellular senescence and mitophagy in lung fibroblasts. Eur Respir J. 2022;60(6):2102307. doi:10.1183/13993003.02307-2021

1325. Li G, Shen C, Wei D, Yang X, Jiang C, Yang X, Mao W, Zou J, Tan J, Chen J. Deficiency of HtrA3 Attenuates Bleomycin-Induced Pulmonary Fibrosis Via TGF-β1/Smad Signaling Pathway. Lung. 2023;201(2):235–242. doi:10.1007/s00408-023-00608-8

1326. Tsoutsou PG, Gourgoulianis KI, Petinaki E, Mpaka M, Efremidou S, Maniatis A, Molyvdas PA. ICAM-1, ICAM-2 and ICAM-3 in the sera of patients with idiopathic pulmonary fibrosis. Inflammation. 2004;28(6):359–364. doi:10.1007/s10753-004-6647-6

1327. Liu T, Xu P, Ke S, Dong H, Zhan M, Hu Q, Li J. Histone methyltransferase SETDB1 inhibits TGF-β-induced epithelial-mesenchymal transition in pulmonary fibrosis by regulating SNAI1 expression and the ferroptosis signaling pathway. Arch Biochem Biophys. 2022;715:109087. doi:10.1016/j.abb.2021.109087

1328. Ebina M, Shimizukawa M, Shibata N, Kimura Y, Suzuki T, Endo M, Sasano H, Kondo T, Nukiwa T. Heterogeneous increase in CD34-positive alveolar capillaries in idiopathic pulmonary fibrosis. Am J Respir Crit Care Med. 2004;169(11):1203–1208. doi:10.1164/rccm.200308-1111OC

1329. Stephenson KE, Porte J, Kelly A, Wallace WA, Huntington CE, Overed-Sayer CL, Cohen ES, Jenkins RG, John AE. The IL-33:ST2 axis is unlikely to play a central fibrogenic role in idiopathic pulmonary fibrosis. Respir Res. 2023;24(1):89. doi:10.1186/s12931-023-02334-4

1330. Zou M, Hu X, Song W, Gao H, Wu C, Zheng W, Cheng Z. Plasma LTBP2 as a potential biomarker in differential diagnosis of connective tissue disease-associated interstitial lung disease and idiopathic pulmonary fibrosis: a pilot study. Clin Exp Med. 2023;23(8):4809–4816. doi:10.1007/s10238-023-01214-x

1331. Li Y, Liang J, Yang T, Monterrosa Mena J, Huan C, Xie T, Kurkciyan A, Liu N, Jiang D, Noble PW. Hyaluronan synthase 2 regulates fibroblast senescence in pulmonary fibrosis. Matrix Biol. 2016;55:35–48. doi:10.1016/j.matbio.2016.03.004

1332. Im J, Kim K, Hergert P, Nho RS. Idiopathic pulmonary fibrosis fibroblasts become resistant to Fas ligand-dependent apoptosis via the alteration of decoy receptor 3. J Pathol. 2016;240(1):25–37. doi:10.1002/path.4749

1333. Kolb M, Crestani B, Maher TM. Phosphodiesterase 4B inhibition: a potential novel strategy for treating pulmonary fibrosis. Eur Respir Rev. 2023;32(167):220206. doi:10.1183/16000617.0206-2022

1334. Chen A, Sun Z, Sun D, Huang M, Fang H, Zhang J, Qian G. Integrative bioinformatics and validation studies reveal KDM6B and its associated molecules as crucial modulators in Idiopathic Pulmonary Fibrosis. Front Immunol. 2023;14:1183871. doi:10.3389/fimmu.2023.1183871

1335. Monaghan-Benson E, Wittchen ES, Doerschuk CM, Burridge K. A Rnd3/p190RhoGAP pathway regulates RhoA activity in idiopathic pulmonary fibrosis fibroblasts. Mol Biol Cell. 2018;29(18):2165–2175. doi:10.1091/mbc.E17-11-0642

1336. Hou J, Ma T, Cao H, Chen Y, Wang C, Chen X, Xiang Z, Han X. TNF-α-induced NF-κB activation promotes myofibroblast differentiation of LR-MSCs and exacerbates bleomycin-induced pulmonary fibrosis. J Cell Physiol. 2018;233(3):2409–2419. doi:10.1002/jcp.26112

1337. Tsai CF, Chen YC, Li YZ, Wu CT, Chang PC, Yeh WL. Imperatorin ameliorates pulmonary fibrosis via GDF15 expression. Front Pharmacol. 2023;14:1292137. doi:10.3389/fphar.2023.1292137

1338. Yogo Y, Fujishima S, Inoue T, Saito F, Shiomi T, Yamaguchi K, Ishizaka A. Macrophage derived chemokine (CCL22), thymus and activation-regulated chemokine (CCL17), and CCR4 in idiopathic pulmonary fibrosis. Respir Res. 2009;10(1):80. doi:10.1186/1465-9921-10-80

1339. Adegunsoye A, Hrusch CL, Bonham CA, Jaffery MR, Blaine KM, Sullivan M, Churpek MM, Strek ME, Noth I, Sperling AI. Skewed Lung CCR4 to CCR6 CD4+ T Cell Ratio in Idiopathic Pulmonary Fibrosis Is Associated with Pulmonary Function. Front Immunol. 2016;7:516. doi:10.3389/fimmu.2016.00516

1340. Königshoff M, Dumitrascu R, Udalov S, Amarie OV, Reiter R, Grimminger F, Seeger W, Schermuly RT, Eickelberg O. Increased expression of 5-hydroxytryptamine2A/B receptors in idiopathic pulmonary fibrosis: a rationale for therapeutic intervention. Thorax. 2010;65(11):949–955. doi:10.1136/thx.2009.134353

1341. Chen H, Wang Q, Li J, Li Y, Chen A, Zhou J, Zhao J, Mao Z, Zhou Z, Zhang J, et al. IFNγ Transcribed by IRF1 in CD4+ Effector Memory T Cells Promotes Senescence-Associated Pulmonary Fibrosis. Aging Dis. 2023;14(6):2215–2237. doi:10.14336/AD.2023.0320

1342. Pham TX, Lee J, Guan J, Caporarello N, Meridew JA, Jones DL, Tan Q, Huang SK, Tschumperlin DJ, Ligresti G. Transcriptional analysis of lung fibroblasts identifies PIM1 signaling as a driver of aging-associated persistent fibrosis. JCI Insight. 2022;7(6):e153672. doi:10.1172/jci.insight.153672

1343. Jiao B, Zhang Q, Jin C, Yu H, Wu Q. IRF4 Participates in Pulmonary Fibrosis Induced by Silica Particles through Regulating Macrophage Polarization and Fibroblast Activation. Inflammation. 2023. doi:10.1007/s10753-023-01890-7

1344. Yu X, Gu P, Huang Z, Fang X, Jiang Y, Luo Q, Li X, Zhu X, Zhan M, Wang J, et al. Reduced expression of BMP3 contributes to the development of pulmonary fibrosis and predicts the unfavorable prognosis in IIP patients. Oncotarget. 2017;8(46):80531–80544. doi:10.18632/oncotarget.20083

1345. Bernau K, Leet JP, Bruhn EM, Tubbs AJ, Zhu T, Sandbo N. Expression of serum response factor in the lung mesenchyme is essential for development of pulmonary fibrosis. Am J Physiol Lung Cell Mol Physiol. 2021;321(1):L174–L188. doi:10.1152/ajplung.00323.2020

1346. Swigris JJ, Brown KK. The role of endothelin-1 in the pathogenesis of idiopathic pulmonary fibrosis. BioDrugs. 2010;24(1):49–54. doi:10.2165/11319550-000000000-00000

1347. Huang LT, Chou HC, Chen CM. Inhibition of FABP4 attenuates hyperoxia-induced lung injury and fibrosis via inhibiting TGF-β signaling in neonatal rats. J Cell Physiol. 2022;237(2):1509–1520. doi:10.1002/jcp.30622

1348. Derlin T, Jaeger B, Jonigk D, Apel RM, Freise J, Shin HO, Weiberg D, Warnecke G, Ross TL, Wester HJ, et al. Clinical Molecular Imaging of Pulmonary CXCR4 Expression to Predict Outcome of Pirfenidone Treatment in Idiopathic Pulmonary Fibrosis. Chest. 2021;159(3):1094–1106. doi:10.1016/j.chest.2020.08.2043

1349. Oda K, Yatera K, Izumi H, Ishimoto H, Yamada S, Nakao H, Hanaka T, Ogoshi T, Noguchi S, Mukae H. Profibrotic role of WNT10A via TGF-β signaling in idiopathic pulmonary fibrosis. Respir Res. 2016;17:39. doi:10.1186/s12931-016-0357-0

1350. Xylourgidis N, Min K, Ahangari F, Yu G, Herazo-Maya JD, Karampitsakos T, Aidinis V, Binzenhöfer L, Bouros D, Bennett AM, et al. Role of dual-specificity protein phosphatase DUSP10/MKP-5 in pulmonary fibrosis. Am J Physiol Lung Cell Mol Physiol. 2019;317(5):L678–L689. doi:10.1152/ajplung.00264.2018

1351. Wang P, Xie D, Xiao T, Cheng C, Wang D, Sun J, Wu M, Yang Y, Zhang A, Liu Q. H3K18 lactylation promotes the progression of arsenite-related idiopathic pulmonary fibrosis via YTHDF1/m6A/NREP. J Hazard Mater. 2024;461:132582. doi:10.1016/j.jhazmat.2023.132582

1352. Damazo AS, Sampaio AL, Nakata CM, Flower RJ, Perretti M, Oliani SM. Endogenous annexin A1 counter-regulates bleomycin-induced lung fibrosis. BMC Immunol. 2011;12:59. doi:10.1186/1471-2172-12-59

1353. Fan Y, Zheng C, Ma R, Wang J, Yang S, Ye Q. MMP19 Variants in Familial and Sporadic Idiopathic Pulmonary Fibrosis. Lung. 2023;201(6):571–580. doi:10.1007/s00408-023-00652-4

1354. Sokai A, Handa T, Tanizawa K, Oga T, Uno K, Tsuruyama T, Kubo T, Ikezoe K, Nakatsuka Y, Tanimura K, et al. Matrix metalloproteinase-10: a novel biomarker for idiopathic pulmonary fibrosis. Respir Res. 2015;16:120. doi:10.1186/s12931-015-0280-9

1355. Tanaka C, Fujimoto M, Hamaguchi Y, Sato S, Takehara K, Hasegawa M. Inducible costimulator ligand regulates bleomycin-induced lung and skin fibrosis in a mouse model independently of the inducible costimulator/inducible costimulator ligand pathway. Arthritis Rheum. 2010;62(6):1723–1732. doi:10.1002/art.27428

1356. Tsoumakidou M, Karagiannis KP, Bouloukaki I, Zakynthinos S, Tzanakis N, Siafakas NM. Increased bronchoalveolar lavage fluid CD1c expressing dendritic cells in idiopathic pulmonary fibrosis. Respiration. 2009;78(4):446–452. doi:10.1159/000226244

1357. Ye Q, Taleb SJ, Wang H, Parinandi NL, Kass DJ, Rojas M, Wang C, Ma Q, Zhao J, Zhao Y. Molecular Regulation of Heme Oxygenase-1 Expression by E2F Transcription Factor 2 in Lung Fibroblast Cells: Relevance to Idiopathic Pulmonary Fibrosis. Biomolecules. 2022;12(10):1531. doi:10.3390/biom12101531

1358. Galán-Cobo A, Arellano-Orden E, Sánchez Silva R, López-Campos JL, Gutiérrez Rivera C, Gómez Izquierdo L, Suárez-Luna N, Molina-Molina M, Rodríguez Portal JA, Echevarría M.The Expression of AQP1 IS Modified in Lung of Patients With Idiopathic Pulmonary Fibrosis: Addressing a Possible New Target. Front Mol Biosci. 2018;5:43. doi:10.3389/fmolb.2018.00043

1359. Lin K, Wang T, Tang Q, Chen T, Lin M, Jin J, Cao J, Zhang S, Xing Y, Qiao L, et al. IL18R1-Related Molecules as Biomarkers for Asthma Severity and Prognostic Markers for Idiopathic Pulmonary Fibrosis. J Proteome Res. 2023;22(10):3320–3331. doi:10.1021/acs.jproteome.3c00389

1360. Lv X, Liu S, Liu C, Li Y, Zhang T, Qi J, Li K, Hua F, Cui B, Zhang X, et al. TRIB3 promotes pulmonary fibrosis through inhibiting SLUG degradation by physically interacting with MDM2. Acta Pharm Sin B. 2023;13(4):1631–1647. doi:10.1016/j.apsb.2023.01.008

1361. Leslie J, Millar BJ, Del Carpio Pons A, Burgoyne RA, Frost JD, Barksby BS, Luli S, Scott J, Simpson AJ, Gauldie J, et al. FPR-1 is an important regulator of neutrophil recruitment and a tissue-specific driver of pulmonary fibrosis. JCI Insight. 2020;5(4):e125937. doi:10.1172/jci.insight.125937

1362. Yang M, Qian X, Wang N, Ding Y, Li H, Zhao Y, Yao S. Inhibition of MARCO ameliorates silica-induced pulmonary fibrosis by regulating epithelial-mesenchymal transition. Toxicol Lett. 2019;301:64–72. doi:10.1016/j.toxlet.2018.10.031

1363. Huang D, Liu G, Xu Z, Chen S, Wang C, Liu D, Cao J, Cheng J, Wu B, Wu D. The multifaceted role of placental growth factor in the pathogenesis and progression of bronchial asthma and pulmonary fibrosis: Therapeutic implications. Genes Dis. 2022;10(4):1537–1551. doi:10.1016/j.gendis.2022.10.017

1364. Hayek H, Rehbini O, Kosmider B, Brandt T, Chatila W, Marchetti N, Criner GJ, Bolla S, Kishore R, Bowler RP, et al. The Regulation of FASN by Exosomal miR-143-5p and miR-342-5p in Idiopathic Pulmonary Fibrosis. Am J Respir Cell Mol Biol. 2023. doi:10.1165/rcmb.2023-0232OC

1365. Ji H, Dong H, Lan Y, Bi Y, Gu X, Han Y, Yang C, Cheng M, Gao J. Metformin attenuates fibroblast activation during pulmonary fibrosis by targeting S100A4 via AMPK-STAT3 axis. Front Pharmacol. 2023;14:1089812. doi:10.3389/fphar.2023.1089812

1366. Mouawad JE, Sharma S, Renaud L, Pilewski JM, Nadig SN, Feghali-Bostwick C. Reduced Cathepsin L expression and secretion into the extracellular milieu contribute to lung fibrosis in systemic sclerosis. Rheumatology (Oxford). 2023;62(3):1306–1316. doi:10.1093/rheumatology/keac411

1367. Huang J, Cao Y, Li X, Yu F, Han X. E2F1 regulates miR-215-5p to aggravate paraquat-induced pulmonary fibrosis via repressing BMPR2 expression. Toxicol Res (Camb). 2022;11(6):940–950. doi:10.1093/toxres/tfac071

1368. d’Amati A, Ronca R, Maccarinelli F, Turati M, Lorusso L, De Giorgis M, Tamma R, Ribatti D, Annese T. PTX3 shapes profibrotic immune cells and epithelial/fibroblast repair and regeneration in a murine model of pulmonary fibrosis. Pathol Res Pract. 2023;251:154901. doi:10.1016/j.prp.2023.154901

1369. Volpe MC, Ciucci G, Zandomenego G, Vuerich R, Ring NAR, Vodret S, Salton F, Marchesan P, Braga L, Marcuzzo T, et al. Flt1 produced by lung endothelial cells impairs ATII cell transdifferentiation and repair in pulmonary fibrosis. Cell Death Dis. 2023;14(7):437. doi:10.1038/s41419-023-05962-2

1370. Louzada RA, Corre R, Ameziane El Hassani R, Meziani L, Jaillet M, Cazes A, Crestani B, Deutsch E, Dupuy C. NADPH oxidase DUOX1 sustains TGF-β1 signalling and promotes lung fibrosis. Eur Respir J. 2021;57(1):1901949. doi:10.1183/13993003.01949-2019

1371. Zhao W, Yue X, Liu K, Zheng J, Huang R, Zou J, Riemekasten G, Petersen F, Yu X. The status of pulmonary fibrosis in systemic sclerosis is associated with IRF5, STAT4, IRAK1, and CTGF polymorphisms. Rheumatol Int. 2017;37(8):1303–1310. doi:10.1007/s00296-017-3722-5

1372. Barnes JW, Duncan D, Helton S, Hutcheson S, Kurundkar D, Logsdon NJ, Locy M, Garth J, Denson R, Farver C, et al. Role of fibroblast growth factor 23 and klotho cross talk in idiopathic pulmonary fibrosis. Am J Physiol Lung Cell Mol Physiol. 2019;317(1):L141–L154. doi:10.1152/ajplung.00246.2018

1373. Kim SY, Kim JM, Lee SR, Kim HJ, Lee JH, Choi HL, Lee YJ, Lee YS, Cho J. Efferocytosis and enhanced FPR2 expression following apoptotic cell instillation attenuate radiation-induced lung inflammation and fibrosis. Biochem Biophys Res Commun. 2022;601:38–44. doi:10.1016/j.bbrc.2022.02.075

1374. Riehl DR, Sharma A, Roewe J, Murke F, Ruppert C, Eming SA, Bopp T, Kleinert H, Radsak MP, Colucci G, et al. Externalized histones fuel pulmonary fibrosis via a platelet-macrophage circuit of TGFβ1 and IL-27. Proc Natl Acad Sci U S A. 2023;120(40):e2215421120. doi:10.1073/pnas.2215421120

1375. Watanabe T, Minezawa T, Hasegawa M, Goto Y, Okamura T, Sakakibara Y, Niwa Y, Kato A, Hayashi M, Isogai S, et al. Prognosis of pulmonary fibrosis presenting with a usual interstitial pneumonia pattern on computed tomography in patients with myeloperoxidase anti-neutrophil cytoplasmic antibody-related nephritis: a retrospective single-center study. BMC Pulm Med. 2019;19(1):194. doi:10.1186/s12890-019-0969-5

1376. Xiong W, Chen S, Xiang H, Zhao S, Xiao J, Li J, Liu Y, Shu Z, Ouyang J, Zhang J, et al. S1PR1 attenuates pulmonary fibrosis by inhibiting EndMT and improving endothelial barrier function. Pulm Pharmacol Ther. 2023;81:102228. doi:10.1016/j.pupt.2023.102228

1377. Nagahara H, Seno T, Yamamoto A, Obayashi H, Inoue T, Kida T, Nakabayashi A, Kukida Y, Fujioka K, Fujii W, et al. Role of allograft inflammatory factor-1 in bleomycin-induced lung fibrosis. Biochem Biophys Res Commun. 2018;495(2):1901–1907. doi:10.1016/j.bbrc.2017.12.035

1378. Kato K, Zemskova MA, Hanss AD, Kim MM, Summer R, Kim KC. Muc1 deficiency exacerbates pulmonary fibrosis in a mouse model of silicosis. Biochem Biophys Res Commun. 2017;493(3):1230–1235. doi:10.1016/j.bbrc.2017.09.047

1379. Bian F, Lan YW, Zhao S, Deng Z, Shukla S, Acharya A, Donovan J, Le T, Milewski D, Bacchetta M, et al. Lung endothelial cells regulate pulmonary fibrosis through FOXF1/R-Ras signaling. Nat Commun. 2023;14(1):2560. doi:10.1038/s41467-023-38177-2

1380. Safaeian L, Jafarian A, Rabbani M, Sadeghi HM, Torabinia N, Alavi SA. The role of strain variation in BAX and BCL-2 expression in murine bleomycin-induced pulmonary fibrosis. Pak J Biol Sci. 2008;11(23):2606–2612. doi:10.3923/pjbs.2008.2606.2612

1381. Zhu W, Ding Q, Wang L, Xu G, Diao Y, Qu S, Chen S, Shi Y. Vitamin D3 alleviates pulmonary fibrosis by regulating the MAPK pathway via targeting PSAT1 expression in vivo and in vitro. Int Immunopharmacol. 2021;101(Pt B):108212. doi:10.1016/j.intimp.2021.108212

1382. Hamanaka RB, Nigdelioglu R, Meliton AY, Tian Y, Witt LJ, O’Leary E, Sun KA, Woods PS, Wu D, Ansbro B, et al. Inhibition of Phosphoglycerate Dehydrogenase Attenuates Bleomycin-induced Pulmonary Fibrosis. Am J Respir Cell Mol Biol. 2018;58(5):585–593. doi:10.1165/rcmb.2017-0186OC

1383. Edwards CJ, Williams E. The role of interleukin-6 in rheumatoid arthritis-associated osteoporosis. Osteoporos Int. 2010;21(8):1287–1293. doi:10.1007/s00198-010-1192-7

1384. Zhang W, Zhang Y, Hu N, Wang A. Alzheimer’s disease-associated inflammatory pathways might contribute to osteoporosis through the interaction between PROK2 and CSF3. Front Neurol. 2022;13:990779. doi:10.3389/fneur.2022.990779

1385. Zhou Q, Zhou L, Li J. MiR-218-5p-dependent SOCS3 downregulation increases osteoblast differentiation inpostmenopausal osteoporosis. J Orthop Surg Res. 2023;18(1):109. doi:10.1186/s13018-023-03580-4

1386. Wan H, Qian TY, Hu XJ, Huang CY, Yao WF. Correlation of Serum CCL3/MIP-1α Levels with Disease Severity in Postmenopausal Osteoporotic Females. Balkan Med J. 2018;35(4):320–325. doi:10.4274/balkanmedj.2017.1165

1387. Fatehi F, Mollahosseini M, Hassanshahi G, Khanamani Falahati-Pour S, Khorramdelazad H, Ahmadi Z, Noroozi Karimabad M, Farahmand H. CC chemokines CCL2, CCL3, CCL4 and CCL5 are elevated in osteoporosis patients. J Biomed Res. 2017;31(5):468–470. doi:10.7555/JBR.31.20150166

1388. Teng Z, Xie X, Zhu Y, Liu J, Hu X, Na Q, Zhang X, Wei G, Xu S, Liu Y, et al. miR-142-5p in Bone Marrow-Derived Mesenchymal Stem Cells Promotes Osteoporosis Involving Targeting Adhesion Molecule VCAM-1 and Inhibiting Cell Migration. Biomed Res Int. 2018;2018:3274641. doi:10.1155/2018/3274641

1389. Liu H, Yi X, Tu S, Cheng C, Luo J. Kaempferol promotes BMSC osteogenic differentiation and improves osteoporosis by downregulating miR-10a-3p and upregulating CXCL12. Mol Cell Endocrinol. 2021;520:111074. doi:10.1016/j.mce.2020.111074

1390. You M, Zhang L, Zhang X, Fu Y, Dong X. MicroRNA-197-3p Inhibits the Osteogenic Differentiation in Osteoporosis by Down-Regulating KLF 10. Clin Interv Aging. 2021;16:107–117. doi:10.2147/CIA.S269171

1391. Lories RJ, Boonen S, Peeters J, de Vlam K, Luyten FP. Evidence for a differential association of the Arg200Trp single-nucleotide polymorphism in FRZB with hip osteoarthritis and osteoporosis. Rheumatology (Oxford). 2006;45(1):113–114. doi:10.1093/rheumatology/kei148

1392. Yu L, Hu M, Cui X, Bao D, Luo Z, Li D, Li L, Liu N, Wu Y, Luo X, et al. M1 macrophage-derived exosomes aggravate bone loss in postmenopausal osteoporosis via a microRNA-98/DUSP1/JNK axis. Cell Biol Int. 2021;45(12):2452–2463. doi:10.1002/cbin.11690

1393. He Z, Sun Y, Wu J, Xiong Z, Zhang S, Liu J, Liu Y, Li H, Jin T, Yang Y, et al. Evaluation of genetic variants in IL-1B and its interaction with the predisposition of osteoporosis in the northwestern Chinese Han population. J Gene Med. 2020;22(10):e3214. doi:10.1002/jgm.3214

1394. Ahmadi H, Khorramdelazad H, Hassanshahi G, Abbasi Fard M, Ahmadi Z, Noroozi Karimabad M, Mollahosseini M. Involvement of Eotaxins (CCL11, CCL24, CCL26) in Pathogenesis of Osteopenia and Osteoporosis. Iran J Public Health. 2020;49(9):1769–1775. doi:10.18502/ijph.v49i9.4098

1395. Xiong L, Zhao K, Cao Y, Guo HH, Pan JX, Yang X, Ren X, Mei L, Xiong WC. Linking skeletal muscle aging with osteoporosis by lamin A/C deficiency. PLoS Biol. 2020;18(6):e3000731. doi:10.1371/journal.pbio.3000731

1396. Liang J, Chen C, Liu H, Liu X, Zhao H, Hu J. Gossypol Promotes Wnt/β-Catenin Signaling through WIF1 in Ovariectomy-Induced Osteoporosis. Biomed Res Int. 2019;2019:8745487. doi:10.1155/2019/8745487

1397. Zou Z, Liu R, Wang Y, Xing Y, Shi Z, Wang K, Dong D. IL1RN promotes osteoblastic differentiation via interacting with ITGB3 in osteoporosis. Acta Biochim Biophys Sin (Shanghai). 2021;53(3):294–303. doi:10.1093/abbs/gmaa174

1398. Biros E, Malabu UH, Vangaveti VN, Birosova E, Moran CS. The IFN-γ/miniTrpRS signaling axis: An insight into the pathophysiology of osteoporosis and therapeutic potential. Cytokine Growth Factor Rev. 2022;64:7–11. doi:10.1016/j.cytogfr.2022.01.005

1399. Wojdasiewicz P, Turczyn P, Dobies-Krzesniak B, Frasunska J, Tarnacka B. Role of CX3CL1/CX3CR1 Signaling Axis Activity in Osteoporosis. Mediators Inflamm. 2019;2019:7570452. doi:10.1155/2019/7570452

1400. Durbano HW, Halloran D, Nguyen J, Stone V, McTague S, Eskander M, Nohe A. Aberrant BMP2 Signaling in Patients Diagnosed with Osteoporosis. Int J Mol Sci. 2020;21(18):6909. doi:10.3390/ijms21186909

1401. Guo W, Jin P, Li R, Huang L, Liu Z, Li H, Zhou T, Fang B, Xia L. Dynamic network biomarker identifies cdkn1a-mediated bone mineralization in the triggering phase of osteoporosis. Exp Mol Med. 2023;55(1):81–94. doi:10.1038/s12276-022-00915-9

1402. Chen S, Jia L, Zhang S, Zheng Y, Zhou Y. DEPTOR regulates osteogenic differentiation via inhibiting MEG3-mediated activation of BMP4 signaling and is involved in osteoporosis. Stem Cell Res Ther. 2018;9(1):185. doi:10.1186/s13287-018-0935-9

1403. Lavigne P, Benderdour M, Lajeunesse D, Shi Q, Fernandes JC. Expression of ICAM-1 by osteoblasts in healthy individuals and in patients suffering from osteoarthritis and osteoporosis. Bone. 2004;35(2):463–470. doi:10.1016/j.bone.2003.12.030

1404. Aggarwal R, Lu J, Kanji S, Joseph M, Das M, Noble GJ, McMichael BK, Agarwal S, Hart RT, Sun Z, et al. Human umbilical cord blood-derived CD34+ cells reverse osteoporosis in NOD/SCID mice by altering osteoblastic and osteoclastic activities. PLoS One. 2012;7(6):e39365. doi:10.1371/journal.pone.0039365

1405. De Martinis M, Ginaldi L, Sirufo MM, Bassino EM, De Pietro F, Pioggia G, Gangemi S. IL-33/Vitamin D Crosstalk in Psoriasis-Associated Osteoporosis. Front Immunol. 2021;11:604055. doi:10.3389/fimmu.2020.604055

1406. Jones DR. A potential osteoporosis target in the FAS ligand/FAS pathway of osteoblast to osteoclast signaling. Ann Transl Med. 2015;3(14):189. doi:10.3978/j.issn.2305-5839.2015.07.01

1407. Lu X, Zhang Y, Zheng Y, Chen B. The miRNA-15b/USP7/KDM6B axis engages in the initiation of osteoporosis by modulating osteoblast differentiation and autophagy. J Cell Mol Med. 2021;25(4):2069–2081. doi:10.1111/jcmm.16139

1408. Murad R, Shezad Z, Ahmed S, Ashraf M, Qadir M, Rehman R. Serum tumour necrosis factor alpha in osteopenic and osteoporotic postmenopausal females: A cross-sectional study in Pakistan. J Pak Med Assoc. 2018;68(3):428–431.

1409. Teawtrakul N, Chansai S, Yamsri S, Chansung K, Wanitpongpun C, Lanamtieng T, Phiphitaporn P, Fucharoen S, Pongchaiyakul C. The association of growth differentiation factor-15 levels and osteoporosis in patients with thalassemia. Am J Med Sci. 2023;366(2):96–101. doi:10.1016/j.amjms.2023.05.002

1410. Czerny B, Kaminski A, Kurzawski M, Kotrych D, Safranow K, Dziedziejko V, Bohatyrewicz A, Pawlik A. The association of IL-1beta, IL-2, and IL-6 gene polymorphisms with bone mineral density and osteoporosis in postmenopausal women. Eur J Obstet Gynecol Reprod Biol. 2010;149(1):82–85. doi:10.1016/j.ejogrb.2009.12.010

1411. Vacher J. Inpp4b is a novel negative modulator of osteoclast differentiation and a prognostic locus for human osteoporosis. Ann N Y Acad Sci. 2013;1280:52–54. doi:10.1111/nyas.12014

1412. Liu H, Xiong Y, Zhu X, Gao H, Yin S, Wang J, Chen G, Wang C, Xiang L, Wang P, et al. Icariin improves osteoporosis, inhibits the expression of PPARγ, C/EBPα, FABP4 mRNA, N1ICD and jagged1 proteins, and increases Notch2 mRNA in ovariectomized rats. Exp Ther Med. 2017;13(4):1360–1368. doi:10.3892/etm.2017.4128

1413. Liao YJ, Chen YT, Hsiao TH, Lin CH, Wu MF, Hsu CY, Chen YM, Hsu CS. CYP2C19 genotypes and osteoporotic fractures in long-term users of proton pump inhibitors: A hospital-based study. Clin Transl Sci. 2023;16(11):2198–2208. doi:10.1111/cts.13620

1414. Anginot A, Nguyen J, Abou Nader Z, Rondeau V, Bonaud A, Kalogeraki M, Boutin A, Lemos JP, Bisio V, Koenen J, et al. WHIM Syndrome-linked CXCR4 mutations drive osteoporosis. Nat Commun. 2023;14(1):2058. doi:10.1038/s41467-023-37791-4

1415. Wang Y, Zhou X, Wang D. Mesenchymal Stem Cell-Derived Extracellular Vesicles Inhibit Osteoporosis via MicroRNA-27a-Induced Inhibition of DKK2-Mediated Wnt/β-Catenin Pathway. Inflammation. 2022;45(2):780–799. doi:10.1007/s10753-021-01583-z

1416. Tsai DJ, Fang WH, Wu LW, Tai MC, Kao CC, Huang SM, Chen WT, Hsiao PJ, Chiu CC, Su W, et al. The Polymorphism at PLCB4 Promoter (rs6086746) Changes the Binding Affinity of RUNX2 and Affects Osteoporosis Susceptibility: An Analysis of Bioinformatics-Based Case-Control Study and Functional Validation. Front Endocrinol (Lausanne). 2021;12:730686. doi:10.3389/fendo.2021.730686

1417. Dera AA, Ranganath L, Barraclough R, Vinjamuri S, Hamill S, Mandourah AY, Barraclough DL. Altered Levels of mRNAs for Calcium-Binding/Associated Proteins, Annexin A1, S100A4, and TMEM64, in Peripheral Blood Mononuclear Cells Are Associated with Osteoporosis. Dis Markers. 2019;2019:3189520. doi:10.1155/2019/3189520

1418. Zhang B, Yuan P, Xu G, Chen Z, Li Z, Ye H, Wang J, Shi P, Sun X. DUSP6 expression is associated with osteoporosis through the regulation of osteoclast differentiation via ERK2/Smad2 signaling. Cell Death Dis. 2021;12(9):825. doi:10.1038/s41419-021-04110-y

1419. Zhou H, Liu W, Zhu J, Liu M, Fang C, Wu Q, Dong N. Reduced serum corin levels in patients with osteoporosis. Clin Chim Acta. 2013;426:152–156. doi:10.1016/j.cca.2013.09.007

1420. Yen CC, Liu YW, Chang GR, Lan YW, Kao YT, Cheng SN, Chen W, Chen CM. Therapeutic Effects of Kefir Peptides on Hemophilia-Induced Osteoporosis in Mice With Deficient Coagulation Factor VIII. Front Cell Dev Biol. 2022;10:794198. doi:10.3389/fcell.2022.794198

1421. Lai CJ. Pharmacophore-based screening of differentially-expressed PGF, DDIT4, COMP and CHI3L1 from hMSC cell lines reveals five novel therapeutic compounds for primary osteoporosis. J Genet Eng Biotechnol. 2016;14(1):203–210. doi:10.1016/j.jgeb.2015.12.002

1422. Rong K, Liang Z, Xiang W, Wang Z, Wen F, Lu L. IL1R2 polymorphisms and their interaction are associated with osteoporosis susceptibility in the Chinese Han population. Int J Immunogenet. 2021;48(6):510–525. doi:10.1111/iji.12547

1423. Potts W, Bowyer J, Jones H, Tucker D, Freemont AJ, Millest A, Martin C, Vernon W, Neerunjun D, Slynn G, et al. Cathepsin L-deficient mice exhibit abnormal skin and bone development and show increased resistance to osteoporosis following ovariectomy. Int J Exp Pathol. 2004;85(2):85–96. doi:10.1111/j.0959-9673.2004.00373.x

1424. Visconti VV, Greggi C, Fittipaldi S, Casamassima D, Tallarico M, Romano F, Botta A, Tarantino U. The long pentraxin PTX3: a novel serum marker to improve the prediction of osteoporosis and osteoarthritis bone-related phenotypes. J Orthop Surg Res. 2021;16(1):288. doi:10.1186/s13018-021-02440-3

1425. Arthur A, Nguyen TM, Paton S, Klisuric A, Zannettino ACW, Gronthos S. The osteoprogenitor-specific loss of ephrinB1 results in an osteoporotic phenotype affecting the balance between bone formation and resorption. Sci Rep. 2018;8(1):12756.doi:10.1038/s41598-018-31190-2

1426. Mao JH, Sui YX, Ao S, Wang Y, Liu Y, Leng H. miR-140-3p exhibits repressive functions on preosteoblast viability and differentiation by downregulating MCF2L in osteoporosis. In Vitro Cell Dev Biol Anim. 2020;56(1):49–58. doi:10.1007/s11626-019-00405-9

1427. Zhou JG, Hua Y, Liu SW, Hu WQ, Qian R, Xiong L. MicroRNA-1286 inhibits osteogenic differentiation of mesenchymal stem cells to promote the progression of osteoporosis via regulating FZD4 expression. Eur Rev Med Pharmacol Sci. 2020;24(1):1-10. doi:10.26355/eurrev_202001_19889

1428. Ahmadi H, Khorramdelazad H, Hassanshahi G, Abbasi Fard M, Ahmadi Z, Noroozi Karimabad M, Mollahosseini M. Involvement of Eotaxins (CCL11, CCL24, CCL26) in Pathogenesis of Osteopenia and Osteoporosis. Iran J Public Health. 2020;49(9):1769–1775. doi:10.18502/ijph.v49i9.4098

1429. Wang L, Cheng L, Zhang B, Wang N, Wang F. Tanshinone prevents alveolar bone loss in ovariectomized osteoporosis rats by up-regulating phosphoglycerate dehydrogenase. Toxicol Appl Pharmacol. 2019;376:9–16. doi:10.1016/j.taap.2019.05.014

1430. Liu Y, Liu H, Li M, Zhou P, Xing X, Xia W, Zhang Z, Liao E, Chen D, et al. Association of farnesyl diphosphate synthase polymorphisms and response to alendronate treatment in Chinese postmenopausal women with osteoporosis. Chin Med J (Engl). 2014;127(4):662–668.

1431. Akdag A, Dilli D, Erdeve O, Oğuz SS, Dilmen U. Does polycythemia affect interleukin-6 response pattern in early postnatal period?. J Clin Lab Anal. 2010;24(5):340–347. doi:10.1002/jcla.20413

1432. Usenko T, Eskinazi D, Correa PN, Amato D, Ben-David Y, Axelrad AA. Overexpression of SOCS-2 and SOCS-3 genes reverses erythroid overgrowth and IGF-I hypersensitivity of primary polycythemia vera (PV) cells. Leuk Lymphoma. 2007;48(1):134–146. doi:10.1080/10428190601043138

1433. Stetka J, Vyhlidalova P, Lanikova L, Koralkova P, Gursky J, Hlusi A, Flodr P, Hubackova S, Bartek J, Hodny Z, et al. Addiction to DUSP1 protects JAK2V617F-driven polycythemia vera progenitors against inflammatory stress and DNA damage, allowing chronic proliferation. Oncogene. 2019;38(28):5627–5642. doi:10.1038/s41388-019-0813-7

1434. Feng G, Zhang T, Liu J, Ma X, Li B, Yang L, Zhang Y, Xu Z, Qin T, Zhou J, et al. MLF1IP promotes normal erythroid proliferation and is involved in the pathogenesis of polycythemia vera. FEBS Lett. 2017;591(5):760–773. doi:10.1002/1873-3468.12587

1435. Liu L, Zhang Y, Zhang Z, Zhao Y, Fan X, Ma L, Zhang Y, He H, Kang L. Associations of high altitude polycythemia with polymorphisms in EPHA2 and AGT in Chinese Han and Tibetan populations. Oncotarget. 2017;8(32):53234–53243. doi:10.18632/oncotarget.18384

1436. Yigit N, Covey S, Barouk-Fox S, Turker T, Geyer JT, Orazi A. Nuclear factor-erythroid 2, nerve growth factor receptor, and CD34-microvessel density are differentially expressed in primary myelofibrosis, polycythemia vera, and essential thrombocythemia. Hum Pathol. 2015;46(8):1217–1225. doi:10.1016/j.humpath.2015.05.004

1437. Gangemi S, Allegra A, Profita M, Saitta S, Gerace D, Bonanno A, Alonci A, Petrungaro A, Russo S, Musolino C. Decreased plasma levels of IL-33 could contribute to the altered function of Th2 lymphocytes in patients with polycythemia vera and essential thrombocythemia. Cancer Invest. 2013;31(3):212–213. doi:10.3109/07357907.2013.764566

1438. Albayrak C, Tarkun P, Birtaş Ateşoğlu E, Eraldemir C, Özsoy ÖD, Terzi Demirsoy E, Mehtap Ö, Gedük A, Hacıhanefioğlu A. The role of hepcidin, GDF15, and mitoferrin-1 in iron metabolism of polycythemia vera and essential thrombocytosis patients. Turk J Med Sci. 2019;49(1):74–80. doi:10.3906/sag-1803-13

1439. Paul CC, Baumann MA. Impaired interleukin-2 production by T-lymphocytes in polycythemia vera. J Clin Lab Anal. 1989;3(2):84–87. doi:10.1002/jcla.1860030204

1440. Caruccio L, Bettinotti M, Director-Myska AE, Arthur DC, Stroncek D. The gene overexpressed in polycythemia rubra vera, PRV-1, and the gene encoding a neutrophil alloantigen, NB1, are alleles of a single gene, CD177, in chromosome band 19q13.31. Transfusion. 2006;46(3):441–447. doi:10.1111/j.1537-2995.2006.00741.x

1441. Jamwal M, Mallik N, Aravindan AV, Jain A, Sharma P, Malhotra P, Das R. Hemolytic erythrocytosis: an amalgamated phenotype from coinherited Chuvash polycythemia and G6PD Kerala-Kalyan with acquired transient stomatocytosis. Ann Hematol. 2021;100(8):2107–2109. doi:10.1007/s00277-020-04295-w

1442. Hasselbalch HC. A role of NF-E2 in chronic inflammation and clonal evolution in essential thrombocythemia, polycythemia vera and myelofibrosis?. Leuk Res. 2014;38(2):263–266. doi:10.1016/j.leukres.2013.07.002

1443. Lussana F, Carobbio A, Salmoiraghi S, Guglielmelli P, Vannucchi AM, Bottazzi B, Leone R, Mantovani A, Barbui T, Rambaldi A. Driver mutations (JAK2V617F, MPLW515L/K or CALR), pentraxin-3 and C-reactive protein in essential thrombocythemia and polycythemia vera. J Hematol Oncol. 2017;10(1):54. doi:10.1186/s13045-017-0425-z

1444. Tefferi A, Lasho TL, Abdel-Wahab O, Guglielmelli P, Patel J, Caramazza D, Pieri L, Finke CM, Kilpivaara O, Wadleigh M, et al. IDH1 and IDH2 mutation studies in 1473 patients with chronic-, fibrotic-or blast-phase essential thrombocythemia, polycythemia vera or myelofibrosis. Leukemia. 2010;24(7):1302–1309. doi:10.1038/leu.2010.113

1445. Malmquist J. Serum myeloperoxidase in leukaemia and polycythaemia vera. Scand J Haematol. 1972;9(4):311–317. doi:10.1111/j.1600-0609.1972.tb00946.x

1446. Li K, Gesang L, Dan Z, Gusang L. Transcriptome reveals the overexpression of a kallikrein gene cluster (KLK1/3/7/8/12) in the Tibetans with high altitude-associated polycythemia. Int J Mol Med. 2017;39(2):287–296. doi:10.3892/ijmm.2016.2830

1447. Koesoemoprodjo W, Maranatha D. Level of serum IL-33 and emphysema paraseptal in clove cigarette smoker with spontaneous pneumothorax: A case report. Respir Med Case Rep. 2020;30:101133. doi:10.1016/j.rmcr.2020.101133

1448. Kalomenidis I, Moschos C, Kollintza A, Sigala I, Stathopoulos GT, Papiris SA, Light RW, Roussos C. Pneumothorax-associated pleural eosinophilia is tumour necrosis factor-alpha-dependent and attenuated by steroids. Respirology. 2008;13(1):73–78. doi:10.1111/j.1440-1843.2007.01153.x

1449. Chen H, Wang L, Liu J, Wan Z, Zhou L, Liao H, Wan R. LncRNA ITGB2-AS1 promotes cisplatin resistance of non-small cell lung cancer by inhibiting ferroptosis via activating the FOSL2/NAMPT axis. Cancer Biol Ther. 2023;24(1):2223377. doi:10.1080/15384047.2023.2223377

1450. Pan S, Li M, Yu H, Xie Z, Li X, Duan X, Huang G, Zhou Z. microRNA-143-3p contributes to inflammatory reactions by targeting FOSL2 in PBMCs from patients with autoimmune diabetes mellitus. Acta Diabetol. 2021;58(1):63–72. doi:10.1007/s00592-020-01591-9

1451. Liu Y, Wang L, Lin XY, Wang J, Yu JH, Miao Y, Wang EH. The transcription factor DEC1 (BHLHE40/STRA13/SHARP-2) is negatively associated with TNM stage in non-small-cell lung cancer and inhibits the proliferation through cyclin D1 in A549 and BE1 cells. Tumour Biol. 2013;34(3):1641–1650. doi:10.1007/s13277-013-0697-z

1452. Wang Z, Yang MQ, Lei L, Fei LR, Zheng YW, Huang WJ, Li ZH, Liu CC, Xu HT. Overexpression of KRT17 promotes proliferation and invasion of non-small cell lung cancer and indicates poor prognosis. Cancer Manag Res. 2019;11:7485–7497. doi:10.2147/CMAR.S218926

1453. Negrete-Garcia MC, Ramírez-Rodriguez SL, Rangel-Escareño C, Muñoz-Montero S, Kelly-García J, Vázquez-Manríquez ME, Santillán P, Ramírez MM, Ramírez-Martínez G, Ramírez-Venegas A, et al. Deregulated MicroRNAs in Cancer-Associated Fibroblasts from Front Tumor Tissues of Lung Adenocarcinoma as Potential Predictors of Tumor Promotion. Tohoku J Exp Med. 2018;246(2):107–120. doi:10.1620/tjem.246.107

1454. Liu X, Zhou X, Chen Y, Huang Y, He J, Luo H. miR-186-5p targeting SIX1 inhibits cisplatin resistance in non-small-cell lung cancer cells (NSCLCs). Neoplasma. 2020;67(1):147–157. doi:10.4149/neo_2019_190511N420

1455. Su J, Zhou J, Feng Y, Zhang H, Zhang X, Zhao X, Li Y, Guo X. circPTN Promotes the Progression of Non-Small Cell Lung Cancer through Upregulation of E2F2 by Sponging miR-432-5p. Int J Genomics. 2022;2022:6303996. doi:10.1155/2022/6303996

1456. Lee YC, Lee YC, Li CY, Lee YL, Chen BL. BRCA1 and BRCA2 Gene Mutations and Lung Cancer Sisk: A Meta-Analysis. Medicina (Kaunas). 2020;56(5):212. doi:10.3390/medicina56050212

1457. Wang X, Yin Y, Du R. SOX9 dependent FOXA1 expression promotes tumorigenesis in lung carcinoma. Biochem Biophys Res Commun. 2019;516(1):236–244. doi:10.1016/j.bbrc.2019.05.169

1458. Shan G, Bi G, Zhao G, Liang J, Bian Y, Zhang H, Jin X, Hu Z, Yao G, Fan H, et al. Inhibition of PKA/CREB1 pathway confers sensitivity to ferroptosis in non-small cell lung cancer. Respir Res. 2023;24(1):277. doi:10.1186/s12931-023-02567-3

1459. Romero OA, Torres-Diz M, Pros E, Savola S, Gomez A, Moran S, Saez C, Iwakawa R, Villanueva A, Montuenga LM, et al. MAX inactivation in small cell lung cancer disrupts MYC-SWI/SNF programs and is synthetic lethal with BRG1. Cancer Discov. 2014;4(3):292–303. doi:10.1158/2159-8290.CD-13-0799

1460. Li J, Ji Z, Luo X, Li Y, Yuan P, Long J, Shen N, Lu Q, Zeng Q, Zhong R, et al. Urinary bisphenol A and its interaction with ESR1 genetic polymorphism associated with non-small cell lung cancer: findings from a case-control study in Chinese population. Chemosphere. 2020;254:126835. doi:10.1016/j.chemosphere.2020.126835

1461. Zhang H, Luo Z, Tang J, Tian J, Xiao Y, Sun C, Wang T. Transcription factor NFIC functions as a tumor suppressor in lung squamous cell carcinoma progression by modulating lncRNA CASC2. Cell Cycle. 2022;21(1):63–73. doi:10.1080/15384101.2021.1995130

1462. Mohrherr J, Uras IZ, Moll HP, Casanova E. STAT3: Versatile Functions in Non-Small Cell Lung Cancer. Cancers (Basel). 2020;12(5):1107. doi:10.3390/cancers12051107

1463. Xu H, Xiang QY, Li S, Liu YS. High serum Bhlhe40 levels are associated with subclinical atherosclerosis in patients with type 2 diabetes mellitus: A cross-sectional study. Diab Vasc Dis Res. 2023;20(2):14791641231169246. doi:10.1177/14791641231169246

1464. Fogarty MP, Cannon ME, Vadlamudi S, Gaulton KJ, Mohlke KL. Identification of a regulatory variant that binds FOXA1 and FOXA2 at the CDC123/CAMK1D type 2 diabetes GWAS locus. PLoS Genet. 2014;10(9):e1004633. doi:10.1371/journal.pgen.1004633

1465. Xu Y, Song R, Long W, Guo H, Shi W, Yuan S, Xu G, Zhang T. CREB1 functional polymorphisms modulating promoter transcriptional activity are associated with type 2 diabetes mellitus risk in Chinese population. Gene. 2018;665:133–140. doi:10.1016/j.gene.2018.05.002

1466. Galavi H, Noorzehi N, Saravani R, Sargazi S, Mollashahee-Kohkan F, Shahraki H. Association study of SREBF-2 gene polymorphisms and the risk of type 2 diabetes in a sample of Iranian population. Gene. 2018;660:145–150. doi:10.1016/j.gene.2018.03.080

1467. Ereqat S, Cauchi S, Eweidat K, Elqadi M, Nasereddin A. Estrogen receptor 1 gene polymorphisms (PvuII and XbaI) are associated with type 2 diabetes in Palestinian women. PeerJ. 2019;7:e7164. doi:10.7717/peerj.7164

1468. Zhang Y, Lin C, Chen R, Luo L, Huang J, Liu H, Chen W, Xu J, Yu H, Ding Y. Association analysis of SOCS3, JAK2 and STAT3 gene polymorphisms and genetic susceptibility to type 2 diabetes mellitus in Chinese population. Diabetol Metab Syndr. 2022;14(1):4.doi:10.1186/s13098-021-00774-w

1469. Khurana P, Gupta A, Sugadev R, Sharma YK, Varshney R, Ganju L, Kumar B. nSARS-Cov-2, pulmonary edema and thrombosis: possible molecular insights using miRNA-gene circuits in regulatory networks. ExRNA. 2020;2(1):16. doi:10.1186/s41544-020-00057-y

1470. Zhu JY, Wang G, Huang X, Lee H, Lee JG, Yang P, van de Leemput J, Huang W, Kane MA, Yang P, et al. SARS-CoV-2 Nsp6 damages Drosophila heart and mouse cardiomyocytes through MGA/MAX complex-mediated increased glycolysis. Commun Biol. 2022;5(1):1039. doi:10.1038/s42003-022-03986-6

1471. Jafarzadeh A, Nemati M, Jafarzadeh S. Contribution of STAT3 to the pathogenesis of COVID-19. Microb Pathog. 2021;154:104836. doi:10.1016/j.micpath.2021.104836

1472. Kaitsumaru M, Shiota M, Takamatsu D, Blas L, Matsumoto T, Inokuchi J, Oda Y, Eto M. Interstitial pneumonia after regression by olaparib for neuroendocrine prostate cancer with BRCA1 mutation: a case report. Int Cancer Conf J. 2023;12(2):131–136. doi:10.1007/s13691-022-00592-5

1473. Kulkarni VV, Wang Y, Pantaleon Garcia J, Evans SE. Redox-Dependent Activation of Lung Epithelial STAT3 Is Required for Inducible Protection against Bacterial Pneumonia. Am J Respir Cell Mol Biol. 2023;68(6):679–688. doi:10.1165/rcmb.2022-0342OC

1474. Hoch D, Bachbauer M, Pöchlauer C, Algaba-Chueca F, Tandl V, Novakovic B, Megia A, Gauster M, Saffery R, Glasner A, et al. Maternal Obesity Alters Placental Cell Cycle Regulators in the First Trimester of Human Pregnancy: New Insights for BRCA1. Int J Mol Sci. 2020;21(2):468. doi:10.3390/ijms21020468

1475. Vanwong N, Sukasem C, Unaharassamee W, Jiratjintana N, Na Nakorn C, Hongkaew Y, Puangpetch A. Associations of the SREBF2 Gene and INSIG2 Polymorphisms with Obesity and Dyslipidemia in Thai Psychotic Disorder Patients Treated with Risperidone. J Pers Med. 2021;11(10):943. doi:10.3390/jpm11100943

1476. Guclu-Geyik F, Coban N, Can G, Erginel-Unaltuna N. The rs2175898 Polymorphism in the ESR1 Gene has a Significant Sex-Specific Effect on Obesity. Biochem Genet. 2020;58(6):935–952. doi:10.1007/s10528-020-09987-6

1477. Su T, Huang C, Yang C, Jiang T, Su J, Chen M, Fatima S, Gong R, Hu X, Bian Z, et al. Apigenin inhibits STAT3/CD36 signaling axis and reduces visceral obesity. Pharmacol Res. 2020;152:104586. doi:10.1016/j.phrs.2019.104586

1478. Jaeger B, Schupp JC, Plappert L, Terwolbeck O, Artysh N, Kayser G, Engelhard P, Adams TS, Zweigerdt R, Kempf H, et al. Airway basal cells show a dedifferentiated KRT17highPhenotype and promote fibrosis in idiopathic pulmonary fibrosis. Nat Commun. 2022;13(1):5637. doi:10.1038/s41467-022-33193-0

1479. Mu X, Wang H, Li H. Silencing of long noncoding RNA H19 alleviates pulmonary injury, inflammation, and fibrosis of acute respiratory distress syndrome through regulating the microRNA-423-5p/FOXA1 axis. Exp Lung Res. 2021;47(4):183–197. doi:10.1080/01902148.2021.1887967

1480. Guan S, Wu Y, Zhang Q, Zhou J. TGFLβ1 induces CREB1Lmediated miRL1290 upregulation to antagonize lung fibrosis via Napsin A. Int J Mol Med. 2020;46(1):141–148. doi:10.3892/ijmm.2020.4565

1481. Kurumiya E, Iwata M, Kasuya Y, Tatsumi K, Honda T, Murayama T, Nakamura H. Eliglustat exerts anti-fibrotic effects by activating SREBP2 in TGF-β1-treated myofibroblasts derived from patients with idiopathic pulmonary fibrosis. Eur J Pharmacol. 2024. doi:10.1016/j.ejphar.2024.176366

1482. Zhang Y, Lu W, Zhang X, Lu J, Xu S, Chen S, Zhong Z, Zhou T, Wang Q, Chen J, et al. Cryptotanshinone protects against pulmonary fibrosis through inhibiting Smad and STAT3 signaling pathways. Pharmacol Res. 2019;147:104307. doi:10.1016/j.phrs.2019.104307

1483. Jia G, Liang C, Li W, Dai H. MiR-410-3p facilitates Angiotensin II-induced cardiac hypertrophy by targeting Smad7. Bioengineered. 2022;13(1):119–127. doi:10.1080/21655979.2021.2009968

1484. Ragusa R, Di Molfetta A, D’Aurizio R, Del Turco S, Cabiati M, Del Ry S, Basta G, Pitto L, Amodeo A, Trivella MG, et al. Variations of circulating miRNA in paediatric patients with Heart Failure supported with Ventricular Assist Device: a pilot study. Sci Rep. 2020;10(1):5905. doi:10.1038/s41598-020-62757-7

1485. Zhou S, Jin J, Wang J, Zhang Z, Huang S, Zheng Y, Cai L. Effects of Breast Cancer Genes 1 and 2 on Cardiovascular Diseases. Curr Probl Cardiol. 2021;46(3):100421. doi:10.1016/j.cpcardiol.2019.04.001

1486. Sun Y, Cheng Z, Cui M, Chen Y, Xie R, Lu G, Gao C. GAS5/METTL14/ESR1 genetic variants are related to the susceptibility of coronary heart disease. Funct Integr Genomics. 2022;22(3):341–357. doi:10.1007/s10142-022-00831-1

1487. Ye Y, Jin Q, Gong Q, Li A, Sun M, Jiang S, Jin Y, Zhang Z, He J, Zhuang L. Bioinformatics and Experimental Analyses Reveal NFIC as an Upstream Transcriptional Regulator for Ischemic Cardiomyopathy. Genes (Basel). 2022;13(6):1051. doi:10.3390/genes13061051

1488. Lin CC, Chen SY, Lien HY, Lin SZ, Lee TM. Targeting the PI3K/STAT3 axis modulates age-related differences in macrophage phenotype in rats with myocardial infarction. J Cell Mol Med. 2019;23(9):6378–6392. doi:10.1111/jcmm.14526

1489. Zhao H, Gong J, Li L, Zhi S, Yang G, Li P, Li R, Li J. Vitamin E relieves chronic obstructive pulmonary disease by inhibiting COX2-mediated p-STAT3 nuclear translocation through the EGFR/MAPK signaling pathway. Lab Invest. 2022;102(3):272–280. doi:10.1038/s41374-021-00652-z

1490. Dibble KE, Donorfio LKM, Britner PA, Bellizzi KM. Stress, anxiety, and health-related quality of life in BRCA1/2-positive women with and without cancer: A comparison of four US female samples. Gynecol Oncol Rep. 2022;42:101033. doi:10.1016/j.gore.2022.101033

1491. Chen Z, Gu J, Lin S, Xu Z, Xu H, Zhao J, Feng P, Tao Y, Chen S, Wang P. Saffron essential oil ameliorates CUMS-induced depression-like behavior in mice via the MAPK-CREB1-BDNF signaling pathway. J Ethnopharmacol. 2023;300:115719. doi:10.1016/j.jep.2022.115719

1492. Resende LS, Amaral CE, Soares RB, Alves AS, Alves-Dos-Santos L, Britto LR, Chiavegatto S. Social stress in adolescents induces depression and brain-region-specific modulation of the transcription factor MAX. Transl Psychiatry. 2016;6(10):e914. doi:10.1038/tp.2016.202

1493. Ozsoy F, Nursal AF, Karakus N, Demir MO, Yigit S. Estrogen Receptor 1 Gene rs22346939 and rs9340799 Variants are Associated with Major Depressive Disorder and its Clinical Features. Curr Neurovasc Res. 2021;18(1):12–19. doi:10.2174/1567202618666210531122239

1494. Fernandes MF, Lau D, Sharma S, Fulton S. Anxiety-like behavior in female mice is modulated by STAT3 signaling in midbrain dopamine neurons. Brain Behav Immun. 2021;95:391–400. doi:10.1016/j.bbi.2021.04.013

1495. Chida-Nagai A, Shintani M, Sato H, Nakayama T, Nii M, Akagawa H, Furukawa T, Rana A, Furutani Y, Inai K, Nonoyama S, et al. Role of BRCA1-associated protein (BRAP) variant in childhood pulmonary arterial hypertension. PLoS One. 2019;14(1):e0211450. doi:10.1371/journal.pone.0211450

1496. Zhao C, Le X, Li M, Hu Y, Li X, Chen Z, Hu G, Hu L, Li Q.Inhibition of Hsp110-STAT3 interaction in endothelial cells alleviates vascular remodeling in hypoxic pulmonary arterial Hypertension model. Respir Res. 2023;24(1):289. doi:10.1186/s12931-023-02600-5

1497. Garcia C, Lyon L, Conell C, Littell RD, Powell CB. Osteoporosis risk and management in BRCA1 and BRCA2 carriers who undergo risk-reducing salpingo-oophorectomy. Gynecol Oncol. 2015;138(3):723–726. doi:10.1016/j.ygyno.2015.06.020

1498. Huang QY, Li GH, Kung AW. The -9247 T/C polymorphism in the SOST upstream regulatory region that potentially affects C/EBPalpha and FOXA1 binding is associated with osteoporosis. Bone. 2009;45(2):289–294. doi:10.1016/j.bone.2009.03.676

1499. Liu C, Han Y, Zhao X, Li B, Xu L, Li D, Li G. POLR2A blocks osteoclastic bone resorption and protects against osteoporosis by interacting with CREB1. J Cell Physiol. 2021;236(7):5134–5146. doi:10.1002/jcp.30220

1500. Bai XH, Su J, Mu YY, Zhang XQ, Li HZ, He XF, He XF. Association between the ESR1 and ESR2 polymorphisms and osteoporosis risk: An updated meta-analysis. Medicine (Baltimore). 2023;102(41):e35461. doi:10.1097/MD.0000000000035461

1501. Hong L, Yang C. Eupatilin ameliorates postmenopausal osteoporosis via elevating microRNA-211-5p and repressing JAK2/STAT3 pathway. Environ Toxicol. 2023. doi:10.1002/tox.24069

1502. Kim H, Naura AS, Errami Y, Ju J, Boulares AH. Cordycepin blocks lung injury-associated inflammation and promotes BRCA1-deficient breast cancer cell killing by effectively inhibiting PARP. Mol Med. 2011;17(9-10):893–900. doi:10.2119/molmed.2011.00032

1503. Zhang L, Wu Q, Huang Y, Zheng J, Guo S, He L. Formononetin ameliorates airway inflammation by suppressing ESR1/NLRP3/Caspase-1 signaling in asthma. Biomed Pharmacother. 2023;168:115799. doi:10.1016/j.biopha.2023.115799

1504. Zhong Y, Huang T, Huang J, Quan J, Su G, Xiong Z, Lv Y, Li S, Lai X, Xiang Y, et al. The HDAC10 instructs macrophage M2 program via deacetylation of STAT3 and promotes allergic airway inflammation. Theranostics. 2023;13(11):3568–3581. doi:10.7150/thno.82535

1505. Gowri V, Taur P, Chougule A; COE consortia, Desai M. STAT 3 GOF with Polycythemia: a Twist to the Tale-First Case Report from India. J Clin Immunol. 2022;42(4):866–868. doi:10.1007/s10875-022-01232-6

